# Comprehensive species sampling and sophisticated algorithmic approaches refute the monophyly of Arachnida

**DOI:** 10.1101/2021.08.16.456573

**Authors:** Jesús A. Ballesteros, Carlos E. Santibáñez-López, Caitlin M. Baker, Ligia R. Benavides, Tauana J. Cunha, Guilherme Gainett, Andrew Z. Ontano, Emily V.W. Setton, Claudia P. Arango, Efrat Gavish-Regev, Mark S. Harvey, Ward C. Wheeler, Gustavo Hormiga, Gonzalo Giribet, Prashant P. Sharma

## Abstract

Deciphering the evolutionary relationships of Chelicerata (arachnids, horseshoe crabs, and allied taxa) has proven notoriously difficult, due to their ancient rapid radiation and the incidence of elevated evolutionary rates in several lineages. While conflicting hypotheses prevail in morphological and molecular datasets alike, the monophyly of Arachnida is nearly universally accepted. Though a small number of phylotranscriptomic analyses have recovered arachnid monophyly, these did not sample all living chelicerate orders. We generated a dataset of 506 high-quality genomes and transcriptomes, sampling all living orders of Chelicerata with high occupancy and rigorous approaches to orthology inference. Our analyses consistently recovered the nested placement of horseshoe crabs within a paraphyletic Arachnida. This result was insensitive to variation in evolutionary rates of genes, complexity of the substitution models, and alternatives algorithmic approaches to species tree inference. Investigation of systematic bias showed that genes and sites that recover arachnid monophyly are enriched in noise and exhibit low information content. To test the effect of morphological data, we generated a 514-taxon morphological data matrix of extant and fossil Chelicerata, analyzed in tandem with the molecular matrix. Combined analyses recovered the clade Merostomata (the marine orders Xiphosura, Eurypterida, and Chasmataspidida), but nested within Arachnida. Our results suggest that morphological convergence resulting from adaptations to life in terrestrial habitats has driven the historical perception of arachnid monophyly, paralleling the history of numerous other invertebrate terrestrial groups.

## Introduction

Chelicerates are a diverse group of arthropods that have played a major role as predators in ancient and recent ecosystems. United by the eponymous pincer-like appendages (the chelicerae/chelifores), chelicerates comprise the sister group to the remaining arthropods. The most familiar chelicerate orders are members of Arachnida, an assemblage of 12 orders of terrestrial arthropods (e.g., spiders, scorpions, mites). Chelicerates also include two wholly marine clades—the sea spiders (Pycnogonida) and the horseshoe crabs (Xiphosura)—as well as considerable diversity of derived aquatic lineages within mites (Wheeler and Hayashi 1998; Giribet et al. 2001; Dabert et al. 2016; Giribet and Edgecombe 2019). The fossil record of chelicerates also attests to a broader aquatic diversity that includes freshwater horseshoe crabs, sea scorpions (Eurypterida) and chasmataspidids (Dunlop et al. 2007; Dunlop 2010; Lamsdell et al. 2016).

Whereas most higher-level phylogenetic relationships of arthropods have been resolved by the advent of phylogenomic approaches (Giribet and Edgecombe 2019; Edgecombe 2020), the internal phylogeny of chelicerates has remained elusive. The traditional paradigm of chelicerate evolution postulates a single colonization of land by the common ancestor of a monophyletic Arachnida. In this scenario, extinct lineages such as the chasmataspidids and sea-scorpions are thought to represent stepping-stones between horseshoe crabs and the origin of arachnids. Phylogenomic studies have recovered weak support for this scenario, with a considerable majority of analyses supporting a nested placement of Xiphosura as derived arachnids (Sharma et al. 2014; Ballesteros and Sharma 2019; Ballesteros et al. 2019; Ontano et al. 2021), a result also recovered in many older Sanger-based molecular analyses (Wheeler and Hayashi 1998; Colgan et al. 1998; Edgecombe et al. 2000; Giribet et al. 2001, 2002; Mallatt et al. 2004; Mallatt and Giribet 2006; Masta et al. 2009; Pepato et al. 2010; Arabi et al. 2012).

A handful of phylogenomic matrices has recovered arachnid monophyly, attributing this result to (a) the use of slowly-evolving (i.e., less saturated) genes that are less prone to long-branch attraction (LBA) artifacts, or (b) expanded taxonomic sampling (Sharma et al. 2014; Lozano-Fernández et al. 2019; Howard et al. 2020). However, the matrices of these works have been shown to be highly sensitive to model choice and algorithmic approach, and ironically lack representation of all extant chelicerate orders. Upon addition of libraries representing those missing arachnid orders to these same datasets, support for arachnid monophyly collapses (Ballesteros et al. 2019; Ontano et al. 2021). Nevertheless, data quality and quantity remain limited for some groups in phylotranscriptomic datasets, specifically orders like Palpigradi and Solifugae.

Beyond arachnid monophyly, internal relationships within Chelicerata are unstable across phylogenomic analyses, which is in part attributable to the incidence of multiple fast-evolving lineages that incur long branch attraction artefacts, such as Acariformes, Parasitiformes, and Pseudoscorpiones (treated as single orders in this study). Well-resolved parts of the chelicerate phylogeny include the reciprocal monophyly of Pycnogonida and Euchelicerata (the remaining chelicerate orders), the monophyly and internal relationships of Tetrapulmonata (spiders and three other orders that plesiomorphically bear four book lungs), and the monophyly of each chelicerate order. More recently, phylogenomic analyses, together with rare genomic changes, have supported the clade Panscorpiones (Scorpiones + Pseudoscorpiones), in turn sister group to Tetrapulmonata (forming the clade Arachnopulmonata) (Sharma et al. 2014; Ontano et al. 2021) (fig. 1).

**Fig. 1.**
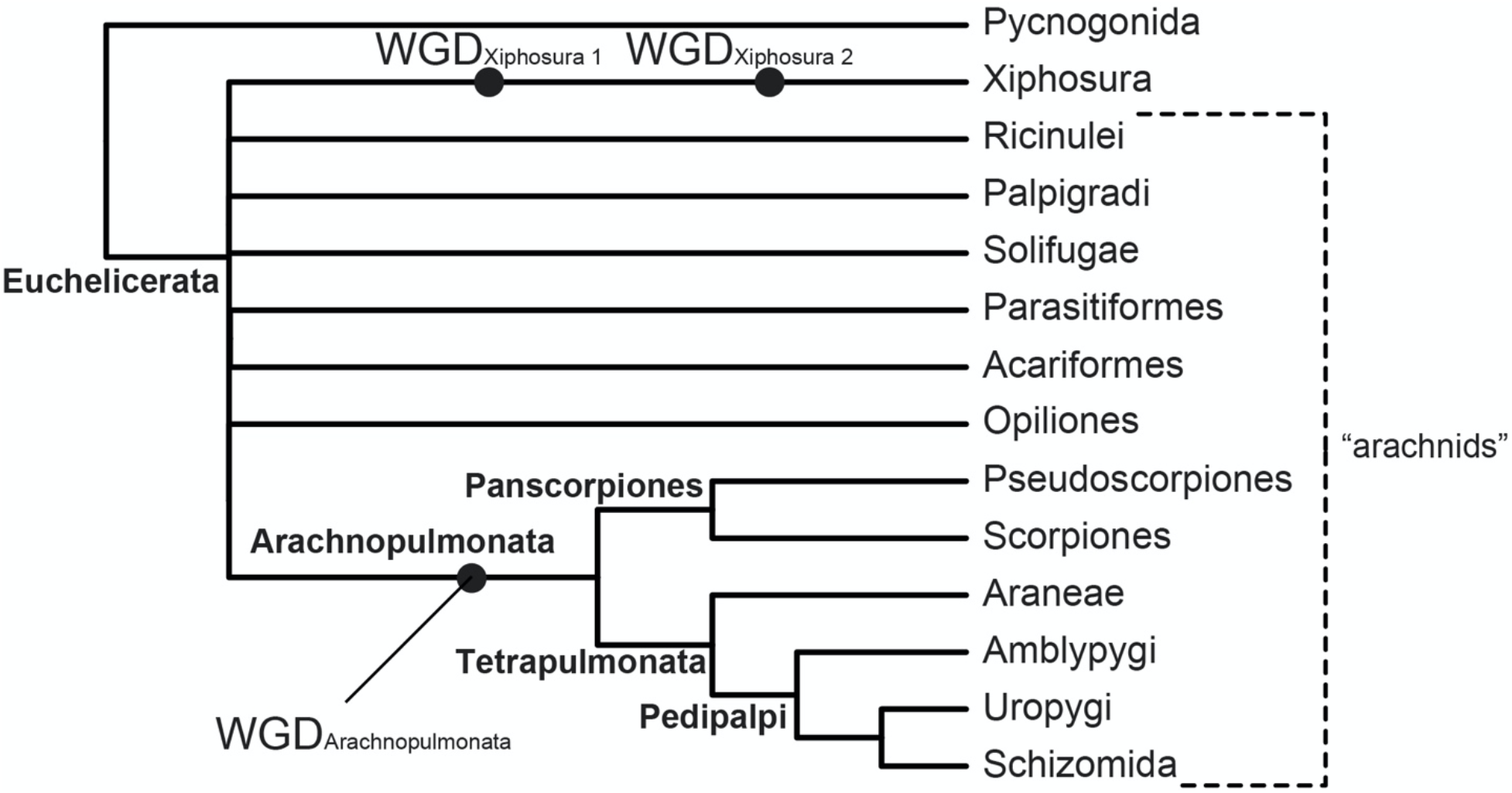
Higher-level phylogeny of Chelicerata showing well-resolved groups (boldface text), based on (10). Circles indicate whole genome duplication events (WGD) subtending specific taxa. Branch lengths are not to scale.

Towards a comprehensive chelicerate phylogeny that can inform the question of arachnid monophyly, we assembled a 506-taxon phylogenomic dataset representing the major lineages of all extant chelicerate orders and densely representing species-rich groups such as Araneae, Scorpiones, Pseudoscorpiones, and Opiliones. Our analyses examined sophisticated strategies to mitigate LBA, such as subsampling loci to minimize saturation, the use of infinite mixture site-heterogeneous models (CAT-GTR), and recently proposed recoding strategies in tandem with site-heterogeneous models applied to partitioned model analyses.

A common feature of phylogenomic studies is the omission of morphological data in an analytical framework, a practice that has been argued to be detrimental to phylogenetic reconstruction (Mongiardino Koch and Parry 2020). Combined analyses of phylogenomics and morphology have been proposed as a means to improve resolution and evaluate congruence between data classes (Mongiardino Koch and Thompson 2021; Neumann et al. 2021). While morphological datasets focusing on relationships among fossil taxa typically recover arachnid monophyly (Lamsdell 2016; Wolfe 2017; but see Aria and Caron 2019; reviewed by Nolan et al. 2020), most of these matrices have historically suffered from minimal sampling of extant arachnid diversity, exhibit marked character conflict, and fail to recover the few relationships that are consistently supported by molecular phylogenies and genomics (e.g., Tetrapulmonata [Wolfe 2017]; Arachnopulmonata [Lamsdell 2016; Wolfe 2017; Aria and Caron 2019]; Euchelicerata [Garwood and Dunlop 2014]). Therefore, toward assessing the impact of fossil taxa and morphological characters on phylogenomic analyses, we assembled a 514-taxon morphological dataset for Chelicerata to complement the phylogenomic dataset. The morphological dataset included extinct taxa (e.g., Chasmataspidida, Eurypterida, Haptopoda, Phalangiotarbida, Synziphosurina, Trigonotarbida, and Uraraneida) as well as key fossils of extant orders.

Here, we show that analyses of molecular datasets alone, as well as combined analyses of morphology and molecules, consistently recover horseshoe crabs as nested within Arachnida. Interrogation of phylogenetic signal across loci showed that genes and sites supporting arachnid monophyly are more prone to systematic error than the remaining loci, suggesting that arachnid monophyly in molecular phylogenies reflects an analytical artifact.

## Results

### Partitioned analyses of phylogenomic datasets

We compiled 506 high-quality transcriptomes or genomes (>95% of libraries generated by us; 80 transcriptomes newly sequenced for this study focused on improving representation of scorpions, palpigrades, and opilioacariforms), sampling 24 outgroup and 482 chelicerate taxa (table s1, Supplementary Material online). Phylogenetically-informed inference of orthologs leveraged a recent *de novo* computation of orthologous genes for Chelicerata (3564 loci identified previously [Ballesteros and Sharma 2019]) using the Unrooted Phylogenetic Orthology (UPhO) pipeline (Ballesteros and Hormiga 2016). As a separate, independent approach to orthology inference, orthologs were drawn from the Benchmarking Universal Single Copy Orthologs loci set for arthropods (BUSCO-Ar) (Simão et al. 2015; Waterhouse et al. 2018). Initial sets of orthologs were filtered based on maximal taxon decisiveness (Steel and Sanderson 2010); we retained only loci that had at least one terminal for all the following clades: Araneae, Pedipalpi (= Uropygi + Schizomida + Amblypygi), Scorpiones, Ricinulei, Xiphosura, Solifugae, Opiliones, Palpigradi, Parasitiformes, Acariformes, Pseudoscorpiones, Pycnogonida, Pancrustacea, Myriapoda, and Onychophora. Applying this criterion, we reduced the UPhO ortholog set to 676 loci (Matrix 1) and the BUSCO set to 399 loci (Matrix 2). Thus, every major lineage (i.e., orders or closely related orders [e.g., Pedipalpi; Acariformes; Parasitiformes]) of chelicerates was represented by at least one terminal for every locus, in all analyses. For both matrices, we implemented (a) the site heterogeneous PMSF model for maximum likelihood search (Wang et al. 2018) (LG+C20+F+*Г*_4_), (b) traditional partitioned model maximum likelihood, and (c) gene tree summary (ASTRAL) approaches.

In all six analyses, we recovered the nested placement of Xiphosura within a paraphyletic Arachnida with support (bootstrap frequency [BS] > 95%; posterior probability > 0.95) and with significance in tests of monophyly (fig. 2; table S2, Supplementary Material online). While relationships of apulmonate arachnid orders varied across topologies, all analyses invariably recovered the monophyly of Tetrapulmonata, Pedipalpi, Euchelicerata, and each chelicerate order. Scorpiones were consistently recovered as the sister group of Tetrapulmonata, whereas Pseudoscorpiones grouped with other long-branched orders (Acariformes and Parasitiformes).

**Fig. 2.**
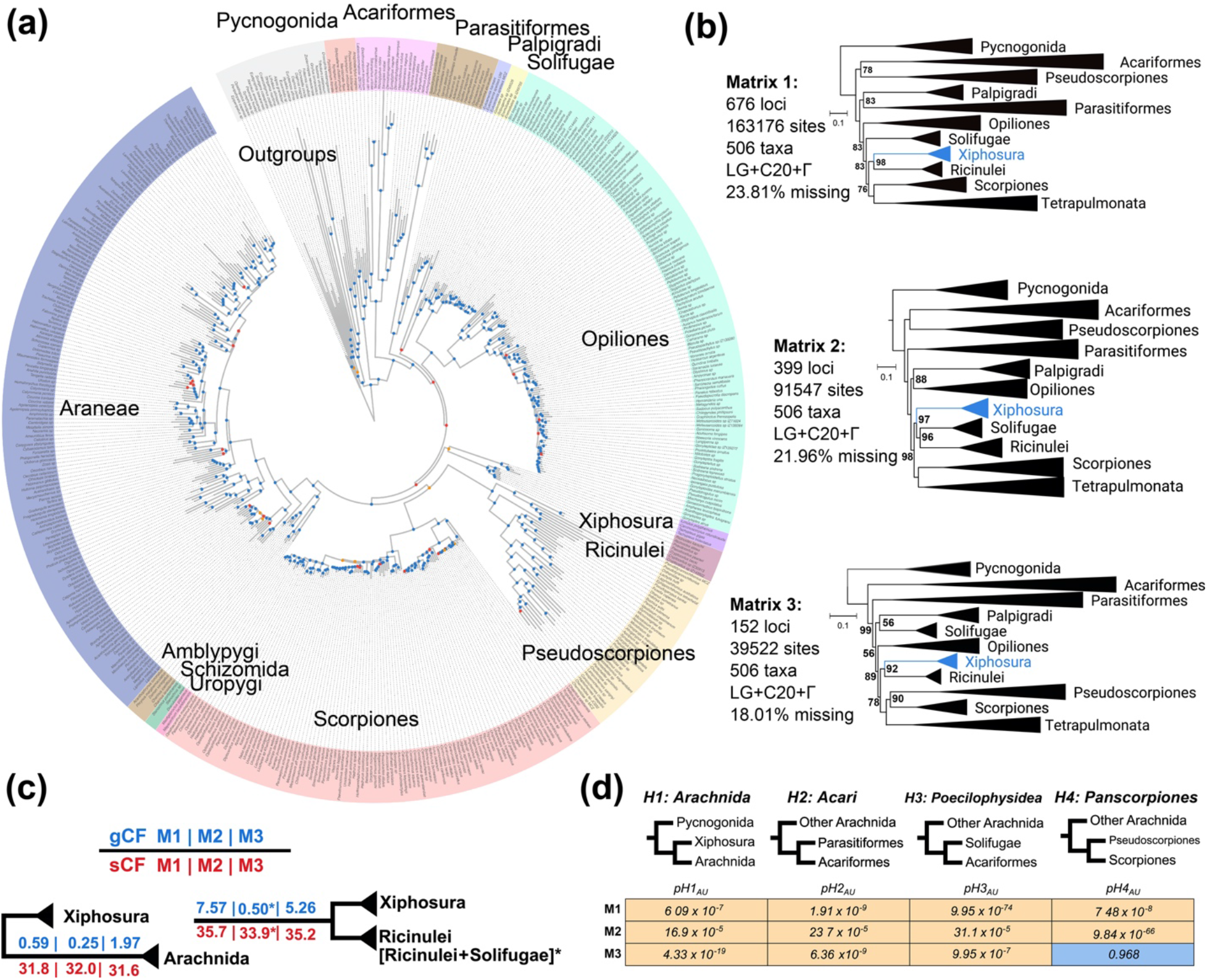
Comprehensive phylogenomic sampling of all extant chelicerate orders recovers horseshoe crabs as derived arachnids. (*a*) Phylogenomic relationships of 506 chelicerate datasets based on maximum likelihood analysis of slowly evolving loci (Matrix 3) and site heterogeneous evolutionary models. Colors correspond to orders; note that Acariformes and Parasitiformes are each treated as orders in this study. Dots on nodes indicate high (>95% bootstrap) or low (<95% boostrap) support. (*b*) Summary of relationships inferred under site heterogeneous models by three matrices. Numbers on nodes correspond to bootstrap resampling frequencies below 100%; all unlabeled nodes are maximally supported. (*c*) Gene (gCF) and site (sCF) concordance factors exhibit higher support for the derived placement of Xiphosura under all three 506-taxon matrices. Asterisks indicate tree topologies wherein Xiphosura was recovered as sister group to Ricinulei + Solifugae. (*d*) Tests of monophyly consistently rejected the monophyly of Arachnida and Acari over the unconstrained topology for Matrices 1–3. Non-significant result for Matrix 3 results from the unconstrained recovery of Panscorpiones in this analysis.

### Analyses of slowly-evolving matrices

In the case of Pseudoscorpiones, an external and independent phylogenetic data class informs the placement of this long-branched order. Specifically, a shared whole genome duplication unites the clade Arachnopulmonata, as evidenced by duplications of Hox clusters, systemic paralogy of developmental patterning genes, and enrichment of microRNA families (Ontano et al. 2021) (fig. 1). As our analyses of Matrices 1 and 2 did not recover a monophyletic Panscorpiones (with pseudoscorpions clustering with other long-branch orders), we reasoned that these datasets remained exposed to LBA.

Several strategies have been proposed to mitigate LBA in arachnid phylogeny, such as the use of site heterogeneous models, the use of slowly-evolving genes, or both (albeit with mixed results across datasets). To mitigate the impact of fast-evolving loci, we generated saturation plots for each locus and isolated a subset of 152 loci with high values for slope (≥0.4) and r^2^ (≥0.95); these loci were concatenated to form Matrix 3 and analyzed using the same approaches as Matrices 1 and 2. Analyses of Matrix 3 with partitioned models, site heterogeneous models, and ASTRAL all recovered the monophyly of Arachnopulmonata (*sensu* Ontano et al. 2021) with maximal nodal support (fig. 2*b*). Maximum likelihood inference under either partitioned or site heterogeneous models also recovered Panscorpiones (BS = 96% and 90%, respectively). All analyses of Matrix 3 rejected arachnid monophyly with support and with significance in tests of monophyly (Fig. 2*d*).

### Bayesian inference analysis with CAT-GTR

Some of the most recalcitrant nodes in the tree of life that are impacted by LBA have been argued to be effectively resolved using analyses under the computationally intensive CAT-GTR infinite mixture model, as implemented in PhyloBayes-mpi. Examples of such nodes include the placement of Chaetognatha, Xenoturbellida, and Porifera (Marlétaz et al. 2019; Kapli and Telford 2020; but see Whelan and Halanych 2016). The PhyloBayes-mpi approach is notoriously difficult to implement for taxon-rich datasets due to the low probability of convergence. We therefore selected 56 representative terminals from the slow-evolving dataset (Matrix 3) such that major taxonomic groups (defined in table s3, Supplementary Material online) were each represented by three to five terminals, major basal splits were represented in each lineage, and the selected taxa exhibited the highest possible data completeness. This dataset was further filtered with BMGE v 1.12 (Criscuolo and Gribaldo 2010) to remove heteropecillous sites, which violate the assumptions of the CAT model (Simion et al. 2017). The resulting matrix (Matrix 4) was comprised of 14,753 sites. Bayesian inference analysis was run on 8 independent chains for >20,000 cycles. To assess the impact of the starting tree on the analysis, two chains (C1 and C2) used the maximum likelihood tree computed for Matrix 4 as starting point (which recovered horseshoe crabs in a derived position). Another two chains (C3 and C4) were started on a maximum likelihood tree for Matrix 4, but constrained to recover arachnid monophyly. Four chains (C5–C8) used random starting trees.

Examination of ESS values and *a posteriori* tree distribution across all eight chains showed that summary statistics broadly exhibited convergence (tables s4 and s5, Supplementary Material online). A high value of the maximum split difference (*maxdiff*) was driven by a soft polytomy at the base of Euchelicerata. We examined estimates both from combined chains as well as summary topologies resulting from each starting tree type (figs. s1 and s2, Supplementary Material online). None of the topologies in the 95% HPD interval of the *a posteriori* distribution supported the monophyly of Arachnida (PP = 0.02814) (fig. 3*a*, 3*b*). Notably, Bayesian analysis using CAT-GTR rejected the monophyly of Acari (Acariformes + Parasitiformes) in favor of Poecilophysidea (Acariformes + Solifugae; PP=1.00) and Cephalosomata (Palpigradi + Poecilophysidea; PP=0.99) (table s2, Supplementary Material online). Moreover, Acari monophyly was supported in 0% of *a posteriori* tree space across the eight chains. These results suggest that Acari reflects another long branch attraction artifact. Notably, PhyloBayes-mpi was able to recover both the monophyly of Arachnopulmonata (PP=1.00) and Panscorpiones (PP≥0.99), regardless of the starting tree topology.

**Fig. 3.**
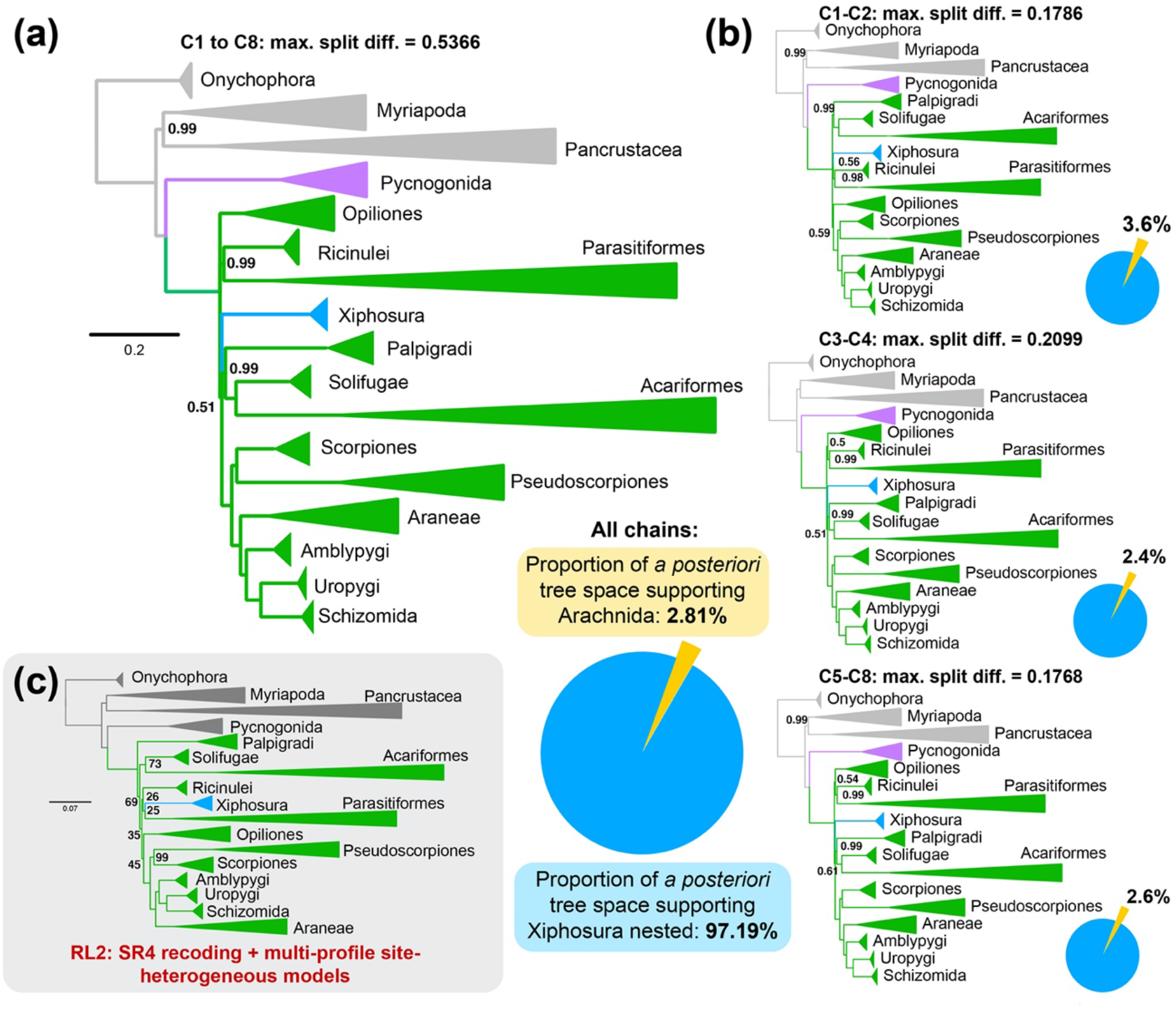
Site heterogeneous model-based approaches using CAT+GTR+*Γ* and SR4 recoding refute the monophyly of Arachnida. (*a*) Summary tree of eight chains from PhyloBayes-mpi analysis of Matrix 4. Numbers on nodes correspond to posterior probabilities below 1.00; all unlabeled nodes are maximally supported. Lower right: Distribution of support across *a posteriori* trees for arachnid monophyly (yellow) versus nested placement of Xiphosura. (*b*) Summary trees from PhyloBayes-mpi analysis separated by starting tree topology. Top: Chains started on maximum likelihood tree topology for Matrix 4 (Xiphosura nested). Middle: Chains started on maximum likelihood tree topology for Matrix 4 with a constraint for arachnid monophyly. Bottom: Chains started on random tree topologies. Nodal support values and pie charts for each summary tree reflect the conventions for (*a*). (*c*) Maximum likelihood tree topology based on SR4 recoding and multi-profile tiered site heterogeneous models (RL2 approach; 38). Numbers on nodes correspond to bootstrap resampling frequencies below 100%; all unlabeled nodes are maximally supported.

### Partitioned analysis with mixture models and recoding

A recently proposed method for reconciling divergent results in partitioned *versus* mixture model studies of recalcitrant nodes makes use of a tiered approach to introduce site-heterogeneous models in tandem with SR4 recoding (RL2, *sensu* Redmond and McLysaght 2021). This approach has been shown to recover consistently the traditional placements of groups like Porifera in empirical datasets.

Upon applying the RL2 strategy to Matrix 3, we recovered yet another tree topology with a nested placement of Xiphosura, as well as Poecilophysidea (BS=73%), Panscorpiones (BS=99%), and Arachnopulmonata (BS=100%) (fig. 3*c*). The backbone of Euchelicerata exhibited negligible support, a result attributable to the loss of information via reducing the peptide alphabet to four states in SR4 recoding. Paralleling this result, previous applications of Dayhoff 6-state recoding to chelicerate datasets have rendered a basal polytomy at the root of Euchelicerata (22). These results are consistent with recent critiques of recoding strategies as solutions to saturation and compositional heterogeneity (Hernandez and Ryan 2021).

### Tests of monophyly and concordance factors

Tests of monophyly were performed using the Approximately Unbiased (AU) test (Shimodaira 2002). The different topologies obtained from Matrices 1–3 were constrained to assess support for the monophyly of Arachnida, Acari, Poecilophysidea (Solifugae + Acariformes), and Panscorpiones (Pseudoscorpiones + Scorpiones). AU tests consistently rejected the monophyly of arachnids over the hypothesis of a derived Xiphosura (fig. 2*d*, table s2, Supplementary Material online).

Traditional measures of nodal support are prone to inflation in phylogenomic datasets. Gene and site concordance factors (gCF and sCF) have been shown to measure phylogenetic signal irrespective of dataset size. We therefore computed values of gCF and sCF both for unconstrained topologies under Matrices 1–3, as well as their counterparts when constrained to recover the monophyly of Arachnida. gCF and sCF values were consistently lower for Arachnida when compared to the hypothesis of a derived Xiphosura (fig. 2*c*).

### Interrogation of phylogenetic signal and systematic bias

To examine whether the derived placement of Xiphosura stemmed from a systematic artifact, we explored phylogenetic signal and properties of genes and sites as a function of support for competing tree topologies (Shen et al. 2017). We found that loci favoring arachnid monophyly were consistently in the minority (39–41%) of genes across our datasets, irrespective of orthology criterion (fig. 4*a*). Proportions of genes supporting arachnid monophyly are comparable to those supporting archaic groupings that have been debunked by phylogenomics and rare genomic changes, such as Dromopoda (=Scorpiones + Opiliones + Solifugae + Pseudoscorpiones; 34–36%) (fig. S3, table s2, Supplementary Material online). Across all matrices, genes exhibited the same distribution of saturation, evolutionary rate and missing data, regardless of support for a monophyletic Arachnida or for Xiphosura nested in Arachnida (fig. 4*b*).

**Fig. 4.**
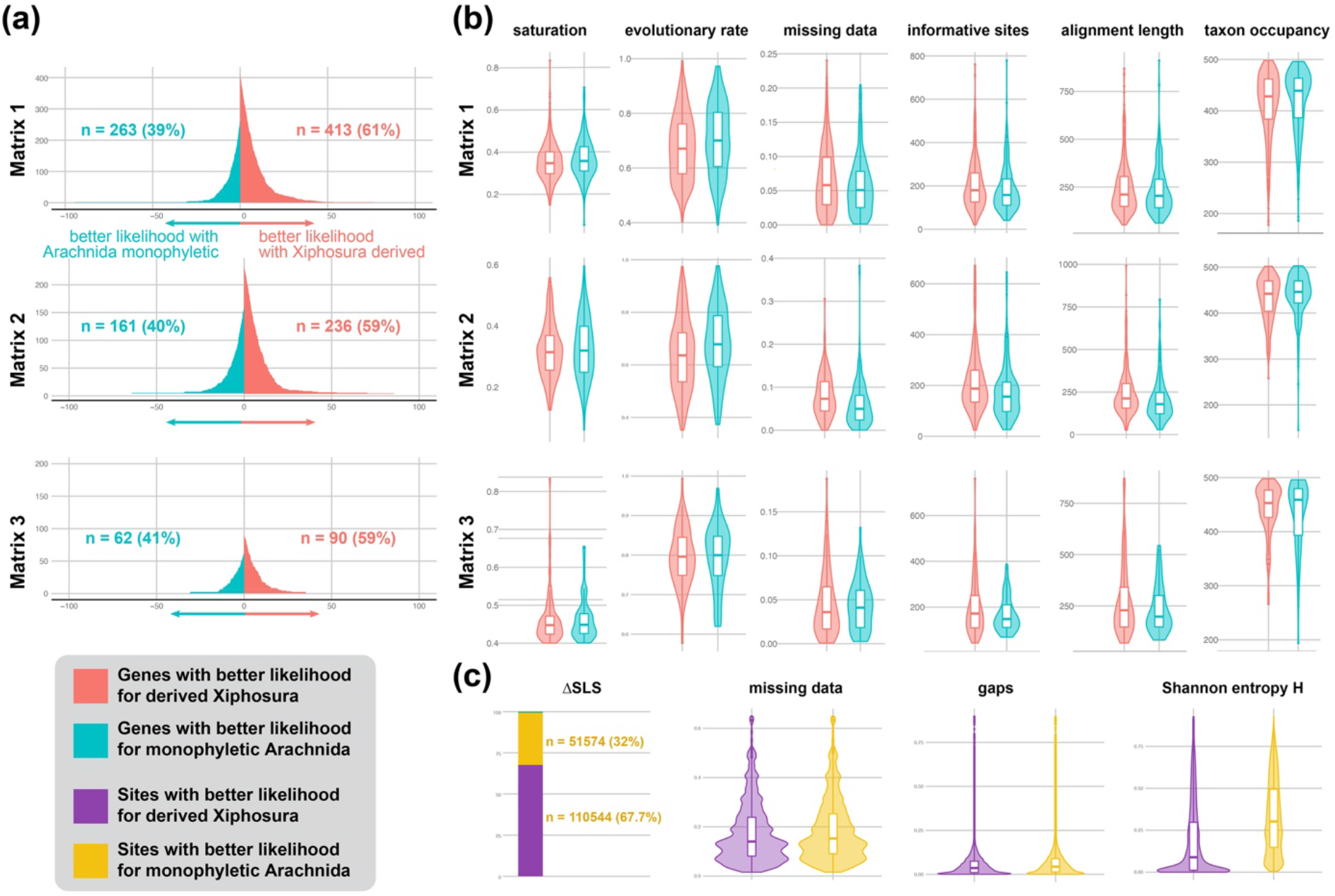
Dissection of phylogenetic signal shows that a minority of artifact-prone genes support arachnid monophyly. (*a*) ΔGLS distributions mapping phylogenetic support for competing hypotheses reveal that a minority of genes (39-41%) support arachnid monophyly, regardless of orthology criterion (Matrices 1 and 2) and filtering of fast-evolving genes (Matrix 3). These proportions are similar to the proportions of genes supporting spurious groupings (*SI Appendix*, Fig. S3). (*b*) Genes supporting the derived placement of Xiphosura exhibit comparable or better metrics of systematic bias (e.g., saturation, evolutionary rate, missing data) than genes supporting Arachnida. (*c*) ΔSLS distributions reveal that the majority of sites (68%) support a derived placement of Xiphosura. Whereas the two categories of sites are similar with respect to missing data, sites supporting arachnid monophyly exhibit high levels of Shannon entropy (exceeding entropy values for spurious groupings; *SI Appendix*, Fig. S4).

Furthermore, we discovered that genes supporting arachnid monophyly were shorter and exhibited fewer parsimony informative sites than genes supporting the unconstrained topology, across all matrices. Short genes with low informativeness have been linked to systematic error across an array of phylogenomic datasets, suggesting that arachnid monophyly may reflect noise rather than true phylogenetic signal. Consistent with this interpretation, we found that sites supporting arachnid monophyly exhibited higher Shannon entropy than sites supporting a nested Xiphosura (fig. 4*c*). Sites supporting arachnid monophyly were fewer in number and had higher Shannon entropy even when compared to sites supporting a debunked grouping that has been falsified by rare genomic changes (Dromopoda; fig. s4, Supplementary Material online).

### Combined analyses of morphology and molecules

To assess the impact of morphological data, we began with the character matrix of Huang *et al.* (2018), the most comprehensively coded morphological matrix of extant chelicerates to date, including recently discovered arachnid fossils that have impacted reconstruction of ancestral states. To this matrix, we added the sea spider *Flagellopantopus* and the extinct order Phalangiotarbida from codings in the literature, as well as all extant chelicerates in the molecular matrix. Errors previously entered in the character coding were corrected. We added new characters from the recent literature pertaining to the neuroanatomy of Xiphosura and several arachnid orders, as well as previously overlooked character systems.

To overcome artefacts stemming from missing and inapplicable character partitions, non-chelicerate outgroup taxa (Onychophora, Mandibulata) were removed from this analysis. For the same reason, we excluded putative chelicerate stem-groups of questionable and controversial placement for which molecular sequence data are inapplicable. Pycnogonida was used to root the Euchelicerata.

When analyzed by itself under equal weights parsimony, the morphological dataset yielded little basal resolution (fig. 5*a*). A strict consensus of equally parsimonious trees recovered a basal polytomy of Euchelicerata. Various interordinal relationships received negligible nodal support, although they accorded closely with recent morphological analyses *viz.* the recovery of Tetrapulmonata (including Trigonotarbida and Haptopoda) and Acaromorpha (Ricinulei + Acari). Under a Bayesian inference approach (fig. 5*b*), the morphological dataset recovered the monophyly of Arachnida, Panscorpiones, Acaromorpha, Acari, and Tetrapulmonata, albeit without support (PP<0.95). Both approaches recovered the monophyly of Merostomata (a grouping of the marine taxa Xiphosura, Synziphosurina, Eurypterida, and Chasmataspidida; BS=90%; PP=1.00).

**Fig. 5.**
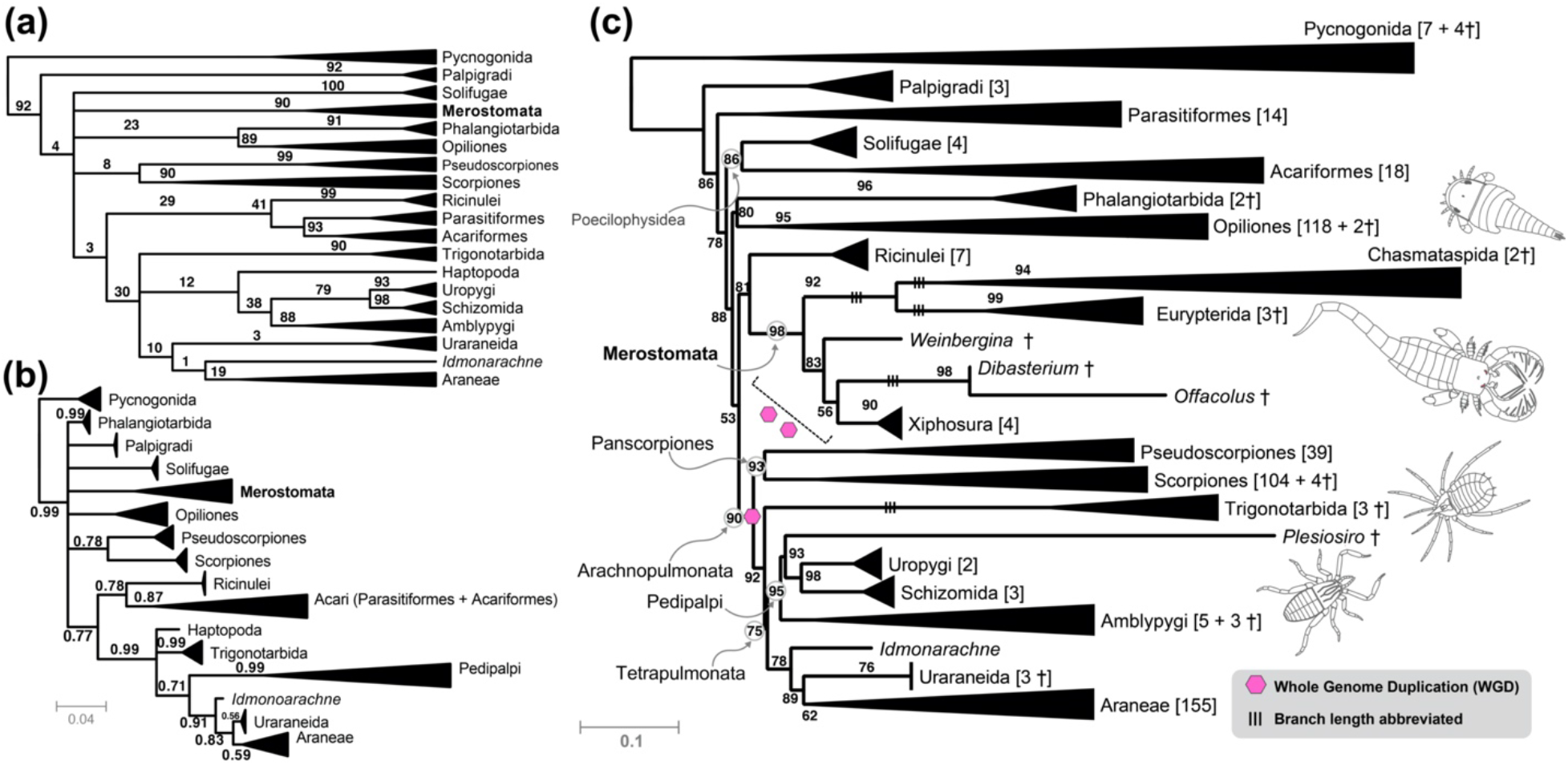
Inclusion of morphology does not rescue arachnid monophyly in total evidence analyses. (*a*) Strict consensus of 1000 equally parsimonious trees inferred for a morphological matrix of 259 morphological characters (482 extant and 32 fossil taxa). Merostomata comprises extinct groups Eurypterida (sea scorpions), Chasmataspidida, and Synziphosurina, as well as Xiphosurida, including the extant Limulidae. Numbers on nodes indicate bootstrap resampling frequencies. (*b*) Summary tree from Bayesian inference analysis of the morphological matrix. Numbers on nodes indicate posterior probabilities below 1.00; unlabeled nodes are maximally supported. (*c*) Maximum likelihood total evidence topology based on 152 slowly-evolving genes and morphological characters. Numbers on nodes represent bootstrap resampling frequencies; unlabeled nodes are maximally supported. Note that the timing of WGD events in Xiphosura cannot be pinpointed on the branches subtending this group.

Maximum likelihood analysis of the combined matrix (fig. 5*c*) also recovered monophyly of Merostomata, which in turn was recovered as the sister group to Ricinulei. We recovered Trigonotarbida as part of the tetrapulmonates, consistent with the presence of two pairs of book lungs in these groups. Phalangiotarbida was recovered as the sister group of Opiliones. Key fossil taxa were recovered in expected placements, such as the harvestman suborder Tetraophthalmi, and the orders Uraraneida and Haptopoda. Palpigradi was recovered as the sister group of the remaining Euchelicerata with moderate support, paralleling the result of the RL2 recoding strategy. Compared to molecular analyses, support values were lowered by the inclusion of morphological data in the combined analysis, a result attributable to the instability incurred by data-poor fossil taxa.

## Discussion

### Arachnid monophyly is not supported by modern phylogenomic approaches

Molecular results that recover non-traditional groupings are often labeled as artifacts, especially when morphological patterns and long-held evolutionary scenarios come under question. Like the basal topology of groups like Metazoa, birds, and angiosperms, the basal topology of Euchelicerata has long defied stability in molecular datasets. Proposals to “correct” the tree and recover arachnid monophyly using molecular datasets have included restricting analyses to slowly-evolving genes (or less saturated genes, a correlate of evolutionary rate) (Sharma et al. 2014; Lozano-Fernández et al. 2019), expansion of taxonomic sampling (Lozano-Fernández et al. 2019), the use of site heterogeneous models (Lozano-Fernández et al. 2019; Howard et al. 2020; but see Sharma et al. 2014; Ontano et al. 2021), or some combination thereof. As our analyses show, the derived placement of Xiphosura (possibly with the other merostomate orders) is consistently recovered despite concomitant application of all these putative solutions. This outcome is consistent with reexaminations of datasets that were previously used to justify arachnid monophyly (Lozano-Fernández et al. 2019; Howard et al. 2020); when reanalyzed with missing groups to achieve the sampling of all extant chelicerate orders, every one of these datasets rejected arachnid monophyly with support (Ballesteros et al. 2019; Ontano et al. 2021).

Why have some recent molecular datasets been able to recover arachnid monophyly (albeit with incomplete sampling of arachnid orders)? As previously shown, the matrices of Lozano et al. (2019) and Howard *et al.* (2020) exhibit a number of bioinformatic and analytical errors in matrix assembly, rendering those matrices flawed and, in one case, unreproducible (supplementary text s2 of Ballesteros et al. 2019; figures S1 and S3 of Ontano et al. 2021). Upon further reexamining those datasets, we additionally found an unexpectedly high number of outliers in root-to-tip distances across gene trees. Using an annotation strategy based on the *Drosophila melanogaster* proteome, we discovered that the cause of this noise was the widescale inclusion of paralogs in these datasets. Specifically, 29% (68/233) of loci in the Lozano et al. Matrix A, and 41% (82/200) of loci in the Howard et al. matrix (algorithm-based orthology inference strategy) included paralogs, often from distantly related multigene families (tables s6, s7, Supplementary Material online).

Could the properties of genes that are able to recover arachnid monophyly inform the selection of “better” loci for chelicerate phylogenomics? To address this, we examined the distribution of phylogenetic signal in our datasets for genes and sites supporting arachnid monophyly, versus the unconstrained topology, using ΔGLS and ΔSLS approaches (Shen et al. 2017). Genes supporting the nested placement of Xiphosura exhibited no evidence of systematic biases compared to the minority, which supported arachnid monophyly (39–41%). Instead, we discovered the opposite trend: genes supporting arachnid monophyly tended to have shorter alignment lengths and fewer informative sites than genes supporting a nested Xiphosura. Short genes and low informativeness are closely associated with phylogenetic error. Consistent with this interpretation, sites supporting arachnid monophyly exhibited higher Shannon entropy and low structure (i.e., greater randomness). For context, the proportions of genes supporting a grouping that has been clearly discredited by genome architecture (i.e., Dromopoda, which historically united two arachnopulmonate orders with two apulmonate orders) are nearly identical to those supporting arachnid monophyly (fig. s3, Supplementary Material online). Moreover, the number of sites supporting this debunked grouping is higher than those supporting arachnid monophyly (fig. s4, Supplementary Material online).

These analyses suggest that support for arachnid monophyly does not reflect hidden signal, so much as noise and error in the datasets that have putatively supported this grouping. We submit that the sum of our analyses, however counterintuitive, may reflect a phylogenetically accurate relationship—Xiphosura (and possibly the other merostomates) may simply constitute derived arachnids.

### Slowly evolving genes and site heterogeneous models overcome LBA artifacts in chelicerate phylogeny

As anticipated, several groups in our phylogeny reflected long root-to-tip distances, constituting lineages prone to LBA artifacts. The inclusion of Opilioacariformes, the slowly-evolving sister group of the remaining Parasitiformes, was previously shown to break up the grouping of Acariformes and Parasitiformes, suggesting that Acari is a long branch artifact (Ontano et al. 2021). In this study, we increased the sampling of Opilioacariformes to three libraries, and concordantly, never obtained the monophyly of Acari, particularly when pursuing approaches best suiting to mitigating LBA. This outcome suggests that the correspondences of mite and tick bauplans (specifically, mouthparts) represent a case of morphological convergence in chelicerates. Similar convergence of mouthparts occurs in the gnathobasic preoral chambers of Opiliones and Scorpiones, which were previously grouped by a subset of morphological analyses (Shultz 2007).

Phylogenomic subsampling for slowly evolving genes did recover Panscorpiones within Arachnopulmonata (fig. 2*b*), a result that is attributable to a marked shift in the proportion of genes supporting this group as a function of evolutionary rate (fig. s5, Supplementary Material online). However, even in maximum likelihood analyses that prioritized slowly evolving genes, we recovered Acariformes and Parasitiformes clustered near the base of the euchelicerate tree, placements that we regarded as possible LBA artifacts. Upon analyzing the slowly evolving matrix with site heterogeneous models in a Bayesian framework (CAT-GTR in PhyloBayes-mpi), not only were Panscorpiones and Arachnopulmonata recovered, but this approach also resolved Acariformes as the sister group of Solifugae (=Poecilophysidea), with Poecilophysidea in turn sister group to Palpigradi (=Cephalosomata) (fig. 3*b*). Four-state recoding in tandem with site heterogeneous models eroded all support from the base of Euchelicerata, but this analysis did recover Poecilophysidea as well (fig. 3*b*).

Intriguingly, these groupings (Poecilophysidea and Cephalosomata, respectively) were previously supported by a minority of phylogenetic analyses and were proposed on the basis of patterns of anterior sclerotization in these orders (Alberti and Peretti 2002; Pepato et al. 2010; Dunlop et al. 2012), potentially validating a subset of morphological character systems in chelicerate higher-level phylogeny. A proximate relationship of Palpigradi and Solifugae is also supported by the anatomy of the coxal gland (Ballesteros et al. 2019). Given the species richness of both Acariformes and Parasitiformes, future efforts to clarify the relative placements of these groups must focus on increasing the representation of basal nodes, a strategy that has been shown to outperform algorithmic and data trimming solutions to resolving the placement of pseudoscorpions (Ontano et al. 2021). Balanced consideration of alternative and overlooked morphological groupings is also warranted in reexaminations of chelicerate phylogeny.

### Morphology is confounded by convergence in chelicerate phylogeny

Unlike in other animal clades (e.g., Koch and Parry 2020), the addition of morphological data to molecular partitions does not ameliorate the discordance with the traditional phylogeny of chelicerates; we found that combining morphological and molecular datasets using model-based approaches recovers Merostomata (the marine group that includes horseshoe crabs) as nested within Arachnida. The notion that morphological synapomorphies of Arachnida can outweigh the dissonance found in molecular data found no support in this study. Furthermore, only in combination with molecular data was morphology able to recover clades supported by rare genomic characters (Panscorpiones and Arachnopulmonata); by itself, morphology has never recovered this arrangement of Arachnopulmonata, either in our analysis or in historical efforts.

One caveat of our combined analysis is that outgroups like putative stem-groups of Chelicerata (e.g., megacheirans) were not included, as their phylogenetic position is controversial even in morphological datasets (Wolfe 2017; Siveter et al. 2017). The exclusion of these groups may prevent character states from being optimized correctly, such as biramous appendages (the presence of exopods), faceted eyes, and gnathobasic mouthparts in marine groups. To assess this possibility, we trialed fusing our molecular dataset (Matrix 3) to two morphological matrices from the literature with widely different taxon sets: a recent, densely sampled matrix of marine crown-group Chelicerata (Bicknell et al. 2019); and a broadly sampled matrix of Panarthropoda (Siveter et al. 2017). Non-overlapping terminals with molecular data only were removed from these analyses to reduce missing data. Upon fusion with Matrix 3, these supplementary datasets featured minimal sampling of extant arachnid fauna (typically, one exemplar per order), as well as greater proportions of missing data than our combined matrix. We found that combining data classes destabilized the traditional relationships previously predicted by those studies, either incurring the non-monophyly of Euchelicerata (fig. s6*a*, s6*b*, Supplementary Material online) or of Chelicerata (fig. s6*c*, s6*d*, Supplementary Material online). Within Euchelicerata, datasets that broadly represented panarthropod diversity (fossil and extant; Siveter et al. 2017) recovered a nested placement of Merostomata within Euchelicerata when combined with molecular data (fig. s6*c*, s6*d*, Supplementary Material online), closely paralleling our results. These outcomes suggest that morphological data partitions seeking to capture deep chelicerate relationships may feature far less robustness of phylogenetic signal than commonly portrayed, specifically in a total evidence framework. Concordantly, a recent paleontological study failed to recover even Tetrapulmonata (Wolfe 2017), the only higher-level group that is consistently recovered by most morphological and molecular datasets. Another recent paleontological study that recovered arachnid non-monophyly took the step of constraining Arachnida *a priori* to ensure the recovery of the traditional topology (Aria and Caron 2019).

Admittedly, the scenario of a nested Xiphosura invites entrenched skepticism, particularly from adherents of paleontology. In addition to an extensive fossil record, horseshoe crabs exhibit an array of putatively plesiomorphic traits that are suggestive of a basally branching placement. The fossil record of merostomates is rich with Xiphosura and Eurypterida species, thought to represent a stepwise colonization of land via internalization of the book gill of these marine groups (for this reason, the position of scorpions at the base of the Arachnida was a central tenet of this evolutionary transformation series). Recent arguments in favor of arachnid monophyly have thus focused on the faceted eye, which is thought to reflect the ancestral condition; the gnathobasic (enditic) mouthparts of merostomates; the biramous condition of merostomate appendages; and the anatomy of the book gill, which shares correspondences with the book lung of large-bodied arachnids (e.g., scorpions; basally branching spiders) (Howard et al. 2020). Moreover, arachnid monophyly has historically been defended on the basis of a series of characters stemming from the musculoskeletal system (Shultz 2001).

However, a comparison with the history of mandibulate arthropod phylogeny offers compelling reasons to doubt the linearity of morphological evolutionary scenarios. Within Chilopoda, only one order of centipedes (Scutigeromorpha) has retained the faceted eye found in fossil outgroups, whereas all other centipedes bear ocelli or are blind, suggesting that faceted eyes are highly prone to discretization and loss in terrestrial habitats in a group at least Devonian in age (Giribet and Edgecombe 2019). Paralleling this trend, various fossil arachnid groups (e.g., fossil scorpions, Trigonotarbida, and fossil Ricinulei) exhibit “semi-compound” eyes (aggregations of ocelli) in head regions positionally homologous to the faceted eyes of Xiphosura and Eurypterida (Dunlop 2010; Howard et al. 2020). The faceted eyes of merostomates may reflect a plesiomorphic condition retained deep in the euchelicerate tree, like the faceted eye of scutigeromorphs within centipedes.

Similarly, discussions of the gnathobasic mouthparts of merostomates echo historical debates over the nature of the gnathobasic mandible of terrestrial mandibulates, as well as other correspondences of head appendages. It was previously thought that Hexapoda and Myriapoda constituted sister groups (the clade Tracheata), a relationship supported by their putatively shared gnathobasic mandible, appendage-free intercalary segment, uniramous appendages, and arrangement of the respiratory organs (tubular tracheae, typically opening as paired spiracles on pleural territories of trunk segments). The gradual overturning of this relationship by molecular phylogenies in favor of the Pancrustacea hypothesis revealed that striking morphological convergences could occur in distantly related taxa as a result of common selection pressures in terrestrial environments (Giribet et al. 2001; Mallatt et al. 2004; Giribet and Mallatt 2006; Giribet and Edgecombe 2019; Edgecombe 2020). In this light, the reduction of gnathobasic mouthparts in terrestrial chelicerate orders could also reflect parallel losses as adaptations to life on land, as evidenced by the uniramous, gnathobasic mandibular architecture of hexapods and myriapods. Parallel losses of secondary rami and simplification of appendages are also broadly observed in terrestrial arthropods, such as arachnids, myriapods, hexapods, and terrestrial malacostracans (e.g., Isopoda, Amphipoda). We submit that the morphology of merostomate appendages is closely tied to evolution in marine habitats and may reflect retention of plesiomorphies; the absence of these structures in terrestrial arthropod groups does not offer compelling evidence uniting Arachnida.

Convergent evolution of tracheal tubules in other terrestrial groups, such as Onychophora, Hexapoda, and Myriapoda, falsifies the interpretation that a lung-like organ is a necessary stepping-stone to the acquisition of tracheal tubules in chelicerates. The conventional and simplistic evolutionary transformation series of book gill to book lung to tracheal tubule is deeply undermined by the complexity of respiratory organ evolution in Chelicerata. This point is underscored by the recent discovery of a eurypterid with trabeculate respiratory organs well after the appearance of arachnids in the fossil record (340 Mya; Lamsdell et al. 2020), secondarily marine scorpions with lamellate gills (*Waeringoscorpio*; Dunlop 2010; Howard et al. 2020), and the diversity of modern aquatic mites (Dabert et al. 2016). The recent recovery of Pseudoscorpiones as a derived member of Arachnopumonata, as well as investigations of respiratory structures across spiders, reveals that book lungs have been frequently lost and repeatedly transformed into tracheal tubules, with loss of book lungs observed in multiple miniaturized arachnopumonate groups (e.g., the posterior book lung pair of Schizomida and most araneomorph spiders; complete loss of book lungs in miniaturized spiders and pseudoscorpions) (Ontano et al. 2021; Ramírez et al. 2021). There is no compelling evidence that evolutionary transitions of respiratory organs have followed a simple, linear series at the base of Arachnida, nor that water-to-land (or the reverse) transitions are rare or irreversible in the arthropod fossil record.

As for the putative musculoskeletal synapomorphies established for Arachnida (musculature and patterns of appendage joints; Shultz 2001), we submit that the evolution of this entire character system may be closely tied to the selective pressures of a terrestrial lifestyle. Arthropod appendages are highly adaptive structures, and biomechanical demands on locomotory appendages differ greatly between aquatic and terrestrial organisms (Boxshall 2004). As with the correspondences of insect and myriapod musculoskeletal anatomy, there is no evident reason why the musculoskeletal system would constitute a homoplasy-free data source for arachnids.

Taken together, morphological character systems that putatively support arachnid monophyly tend to exhibit high levels of homoplasy upon closer examination, especially when examining their counterparts in Mandibulata. Given the remarkable morphological convergence exhibited by Hexapoda and Myriapoda, we postulate that parallel evolution in terrestrial chelicerate orders may confound inferences of homology in morphological datasets. While no morphological characters overtly support a closer relationship of Xiphosura to any subset of arachnid orders (but see Lehmann and Melzer 2013, 2019a, 2019b), the absence of morphological support for numerous, robustly recovered molecular clades is a common feature of ancient invertebrate relationships, as exemplified by the modern higher-level phylogeny of groups like Annelida, Mollusca, and Nematoda (Struck et al. 2015; Smythe et al. 2019; Kocot et al. 2020). Indeed, the discovery of a particular well-supported relationship in molecular datasets typically serves as a catalyst for revitalized morphological study and reinterpretation of previous homology statements, as in the case of Pancrustacea, Ecdysozoa, and Arachnopulmonata (Lehmann and Melzer 2019a, 2019b; Ontano et al. 2021). Given the recovery of Poecilophysidea and Cephalosomata in some analyses (fig. 3), reexamination of previously overlooked interordinal groupings may provide a better understanding of hidden phylogenetic signal in specific chelicerate morphological character systems. A derived placement of merostomates as a group more proximal to Arachnopulmonata could also reconcile the morphology of extinct marine groups like eurypterids with the unambiguously nested position of Scorpiones, a hypothesis that could be tested through functional genetic approaches to understanding the developmental basis for respiratory organ patterning in horseshoe crabs, arachnopulmonates, and apulmonate arachnids.

The nested placement of Xiphosura, together with the reconstruction of multiple terrestrialization events across a grade of arachnid diversity, must be treated as a valid competing hypothesis. Future efforts to integrate new phylogenetic data classes and rare genomic characters (e.g., Ontano et al. 2021) may offer clearer resolution of relationships among the apulmonate arachnid orders and consilience between discordant datasets.

## Conclusion

Analyses of molecular data and total evidence phylogenetic approaches do not support arachnid monophyly. The concept of Arachnida may reflect the antiquated notion that terrestrialization is rare or costly in evolutionary history. As revealed by the history of groups like mandibulate arthropods, nematodes, and Pulmonata (gastropods), terrestrialization has not only evolved many times independently within such taxa, but is also the cause of remarkable and misleading cases of morphological convergence.

The strongest evidence that morphological datasets of Chelicerata may be prone to misinterpretation of homologies is provided by the positions of scorpions and pseudoscorpions, which are united with tetrapulmonates by a rare genomic change (an ancient whole genome duplication event). Morphological datasets, including the dataset we generated, have consistently failed to recover this grouping (with or without the miniaturized Pseudoscorpiones) (Shultz 2007; Garwood and Dunlop 2014; Lamsdell 2016; Wolfe 2017; Siveter et al. 2017; Aria and Caron 2019; Bicknell et al. 2019).

If morphological datasets can falter in the recovery of the only higher-level chelicerate groups robustly resolved by molecular data and genomic architecture, it stands to reason that phylogenetic signal in morphological datasets may not be sufficiently robust to adjudicate other nodes in chelicerate interordinal phylogeny. The traditional placements of Xiphosura, Eurypterida, and various stem-group fossils must then also be regarded as suspect. Given the history of erstwhile morphological groupings like Tracheata, Uniramia, Articulata, Polychaeta, Pulmonata, Opisthobranchia, and numerous others, we postulate that phylogenomic approaches to deep metazoan relationships should treat morphological interpretations with skepticism *prima facie*, especially in the context of selective pressures like terrestrialization that promote morphological convergence and thereby confound inferences of homology.

## Materials and Methods

Details of the methods below are provided in the Supplementary Material.

### Taxon sampling and orthology inference

Taxon selection consisted of 24 outgroup and 482 ingroup terminals; these 506 transcriptomes and genomes (table s1, Supplementary Material online) sampled all extant chelicerate orders with multiple terminals. Eighty new libraries were generated following previously published protocols. Proteomes and peptide sequences were used as inputs. Phylogenetically-informed inference of orthologs leveraged a recent *de novo* computation of orthologous genes for Chelicerata using UPhO (3564 loci identified previously [Ballesteros and Sharma 2019]). For validation, these collections of putative orthologous sequences were BLASTed (blastp v. 2.9.0+ [Camacho et al. 2009]) against the *Drosophila melanogaster* proteome for annotation using the best hit. Sequences not matching the most common annotation were discarded. Separately, orthologs benchmarked using BUSCO-Arthropoda database were analyzed independently. The set of complete, single-copy BUSCOs >100 amino acids in length was retained from each library.

### Matrix construction

Initial sets of orthologs were filtered based on taxon decisiveness (Steel and Sanderson 2010). We retained only loci that had at least one terminal for all the following clades: Araneae, Pedipalpi (Uropygi + Schizomida + Amblypygi), Scorpiones, Ricinulei, Xiphosura, Solifugae, Opiliones, Palpigradi, Parasitiformes (treated here as the order uniting Holothyrida, Ixodida, Mesostigmata, and Opilioacariformes), Acariformes (treated here as the order uniting Sarcoptiformes and Trombidiformes), Pseudoscorpiones, Pycnogonida, Pancrustacea, Myriapoda, and Onychophora. Applying this criterion, we reduced the UPhO ortholog set to 676 loci (Matrix 1) and the BUSCO set to 399 loci (Matrix 2).

To assess the impact of saturation, we generated saturation plots for each locus and isolated a subset of 152 loci with slope ≥ 0.4 and r^2^ ≥ 0.95; these loci were concatenated to form Matrix 3. To operate PhyloBayes-mpi with the computationally demanding CAT+GTR+*Γ* model, we selected 56 representative terminals from the slow-evolving dataset (Matrix 2) such that major taxonomic groups (*SI Appendix*, Table S3) were each represented by three to five terminals, major basal splits were represented in each lineage, and the selected taxa exhibited high data completeness. This dataset was further filtered with BMGE v 1.12 (Criscuolo and Gribaldo 2010) to remove heteropecillous sites to form Matrix 4 for analysis with PhyloBayes-mpi.

### Partitioned analyses

Gene trees were inferred using IQ-TREE v. 1.6.10 (Nguyen et al. 2015) with model-fitting using ModelFinder (Kalyaanamoorthy et al. 2017) and nodal support estimation using the ultrafast bootstrap (Hoang et al. 2018) as follows: *iqtree - mset LG,WAG,JTT,Dayhoff,JTTDCMut,DCMut,PMB -m MFP -bb 1000*. Maximum likelihood analyses of concatenated datasets (Matrices 1–3) were run using a gene partitioning strategy implementing the best substitution models identified during gene tree reconstruction. Tree topologies were inferred using IQ-TREE, with nodal support estimated using ultrafast bootstrapping. Summary coalescent estimates of the species phylogenies were estimated from the individual gene trees using ASTRAL v 3.14.2 (Zhang et al. 2018).

### Mixture model analyses

We computed maximum likelihood analyses with the posterior mean site frequency model (Wang et al. 2018) for Matrices 1-3, using the LG+C20+F+*G* implementation. The use of more site categories (e.g., C60) proved prohibitive for a dataset of this size, with the C20 model demanding 1.1 Tb of RAM to compute site-specific model parameters. Analyses were computed using IQ-TREE v. 1.6.10. Nodal support was estimated using ultrafast bootstrapping. Bayesian inference analysis was performed using PhyloBayes-mpi v 1.8 (Lartillot et al. 2013) and the CAT+GTR+*Γ* model on Matrix 4, which was optimized for this purpose. Bayesian inference analysis was run on 8 independent chains for >20,000 cycles. Convergence of parameters and topologies was assessed using Tracer 1.7.1 (Rambaut et al. 2018) and native PhyloBayes-mpi summary programs. Summary statistics and chain lengths are provided in the *SI Appendix*, respectively. Trace files of parameters and tree files from each run are provided in FigShare. Convergence parameters exhibited differences as a function of combining different chains. Different combinations of chains produced varying maximum split differences. Examination of ESS values and *a posteriori* tree distribution across all eight chains showed that summary statistics broadly exhibited convergence; the high value of the maximum split difference is driven by a soft polytomy at the base of Euchelicerata.

### Recoded mixture model analyses

We implemented a site heterogeneous mixture model approach to partitioned phylogenomics using the RL2 strategy recently proposed (Redmond and McLysaght 2021), which implements 4-state recoding of amino acid data (SR4; Susko and Roger 2007). Analyses were performed in IQ-TREE v. 1.6.10, following the original implementation (Redmond and McLysaght 2021).

### Morphological analysis

We developed a morphological matrix of 259 characters coded for 482 extant and 32 fossil chelicerates. Given the unambiguous recovery of Pycnogonida as the sister group to the remaining chelicerates, Pycnogonida were used to root the tree. Character codings were drawn from previous higher-level analyses of sea spiders (Arango 2002), harvestmen (Giribet et al. 2002; Garwood et al. 2014), scorpions (Sharma et al. 2015), and arachnids (Giribet et al. 2002; Shultz 2007; Garwood and Dunlop 2014; Huang et al. 2018). Errors and discrepancies with previous character codings were modified and we additionally coded new characters informed by recent investigations (Lehmann and Melzer 2013, 2018). Fossil taxa were coded using original descriptions from the literature.

Model-fitting for the morphological dataset was performed in IQ-TREE v. 1.6.10 from the dataset initially partitioned based on the number of character states. Bayesian analyses using the same partitioning scheme were performed in MrBayes v 3.2.7a (Ronquist et al. 2012) using the Mk1 model with unlinked rate and state frequency parameters per partition. The analyses consisted of four independent runs of 50 M cycles. Equal weights and implied weights parsimony analyses were performed using TNT v. 1.5 (Goloboff et al. 2016).

We additionally performed total evidence analyses using two recently published morphological matrices with differing representations of stem-group chelicerate taxa, (Bicknell et al. 2019; Siveter et al. 2017), complemented by molecular data from Matrix 3. In cases of non-overlapping taxa, a chimeric terminal was constructed using the closest related species to a given terminal in the morphological datasets. For each chimeric terminal, character codings were checked to ensure their applicability to their morphological counterpart; no coding changes were required for chimeras (table S8, Supplementary Material online). Due to the degree of missing data in these matrices, analyses were only performed using parsimony (equal and implied weights); model-based analyses consistently failed to converge for these supplementary datasets.

## Data Availability

Raw sequence data that support the findings of this study have been deposited in NCBI Sequence Read Archive. The morphological data matrix and the character list are provided in the files s1 and s2, Supplementary Material online. All transcriptomic assemblies, gene alignments, gene trees, supermatrices, phylogenomic trees, PhyloBayes-mpi trace files, and scripts have been deposited in FigShare.

## Acknowledgments

Sequencing was performed at the Bauer Core Facility (Harvard University) and the BioTechnology Center (UW-Madison). High-throughput computational analyses were performed through the Center for High Throughput Computing (CHTC) and the Bioinformatics Resource Center (BRC) of UW-Madison. J.A.B. was supported by a Guyer postdoctoral fellowship. C.E.S.L. was supported by a CONACYT postdoctoral fellowship (reg. 207146/454834). This material is based on work supported by the National Science Foundation under Grants IOS-1552610 and IOS-2016141 to P.P.S; DEB-1457539 to G. Giribet; and DEB-1457300 to G.H.

