## Supplementary Material for "Comprehensive species sampling and sophisticated algorithmic approaches refute the monophyly of Arachnida"

**Fig. S1.** Summary of *a posteriori* trees from Phylobayes-mpi analysis of Matrix 4 under the CAT-GTR model. Left column: Individual summary trees from chains 1-4. Top right: Summary tree from combining chains 1 and 2 (starting trees using unconstrained ML analysis of Matrix 4). Bottom right: Summary tree from combining chains 3 and 4 (starting trees using ML analysis of Matrix 4, with arachnid monophyly constrained).

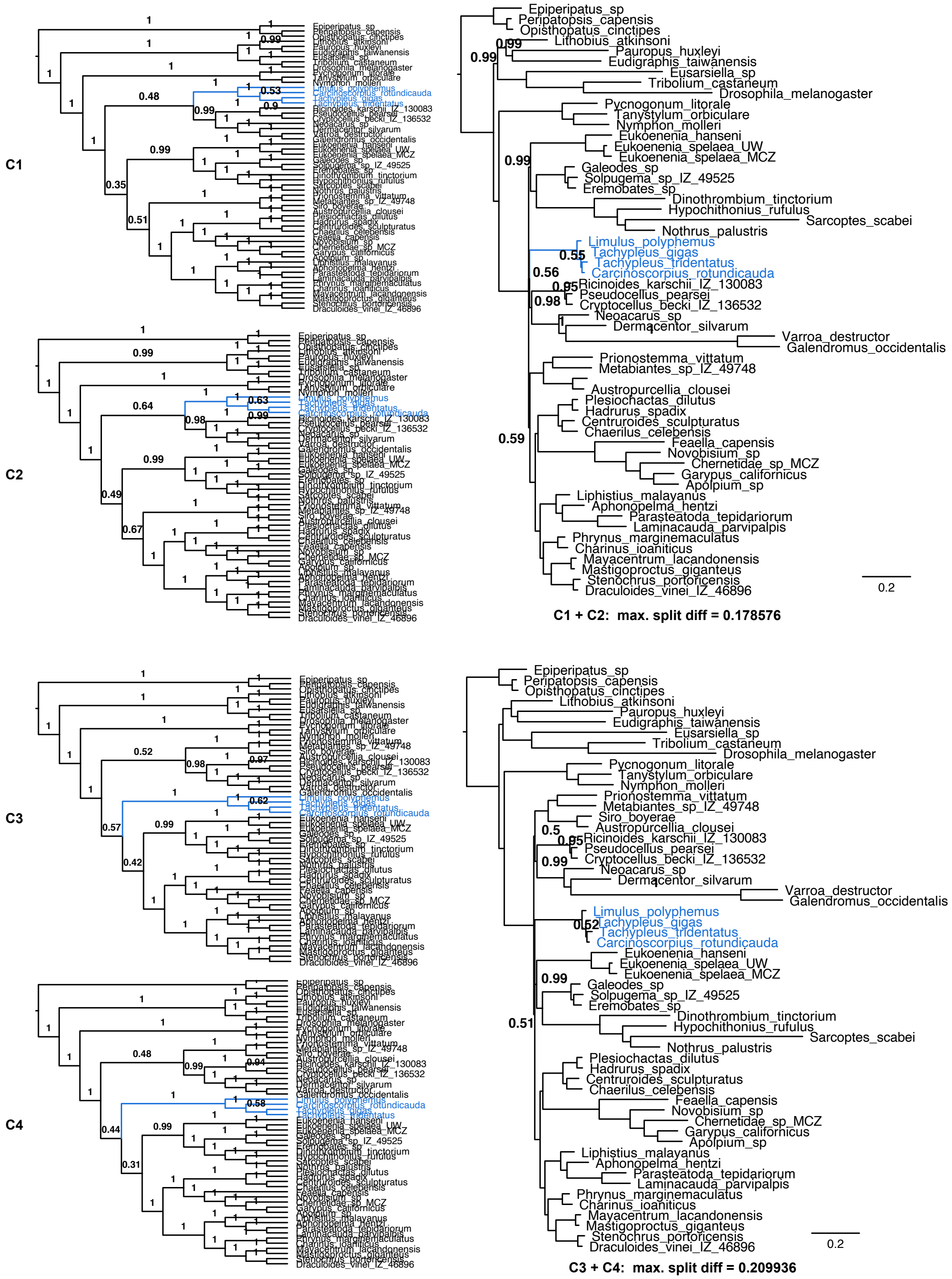

**Fig. S2.** Summary of *a posteriori* trees from Phylobayes-mpi analysis of Matrix 4 under the CAT-GTR model. Left column: Individual summary trees from chains 5-8. Right: Summary tree from combining chains 5-8 (random starting trees).

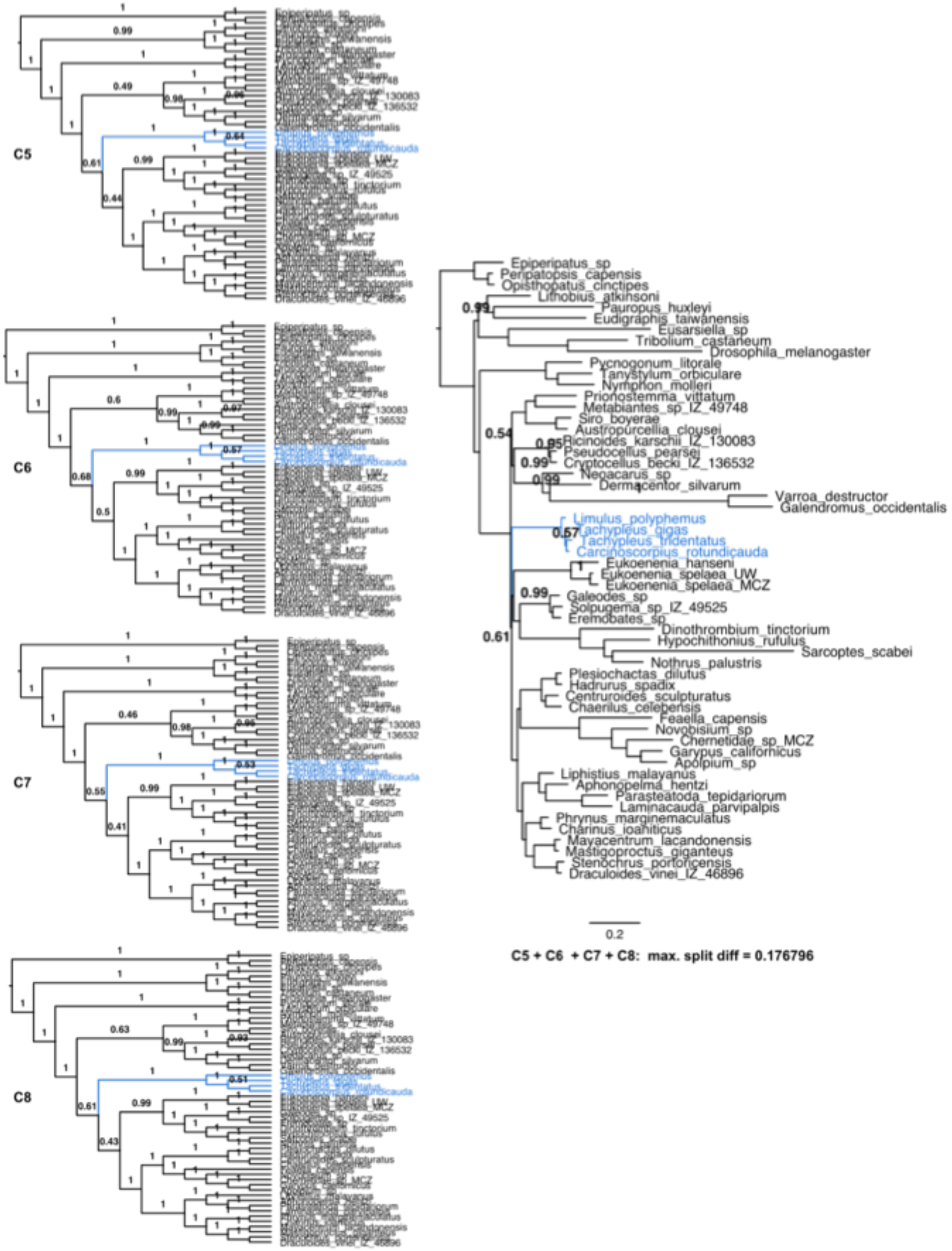

**Fig. S3.** ΔGLS distributions mapping phylogenetic support for the unconstrained tree topology versus trees constrained to recover Dromopoda (Scorpiones + Pseudoscorpiones + Opiliones + Solifugae), for Matrices 1 and 3. A minority of genes (34-36%) support monophyly of Dromopoda, which has been refuted by phylogenomics and rare genomic changes.

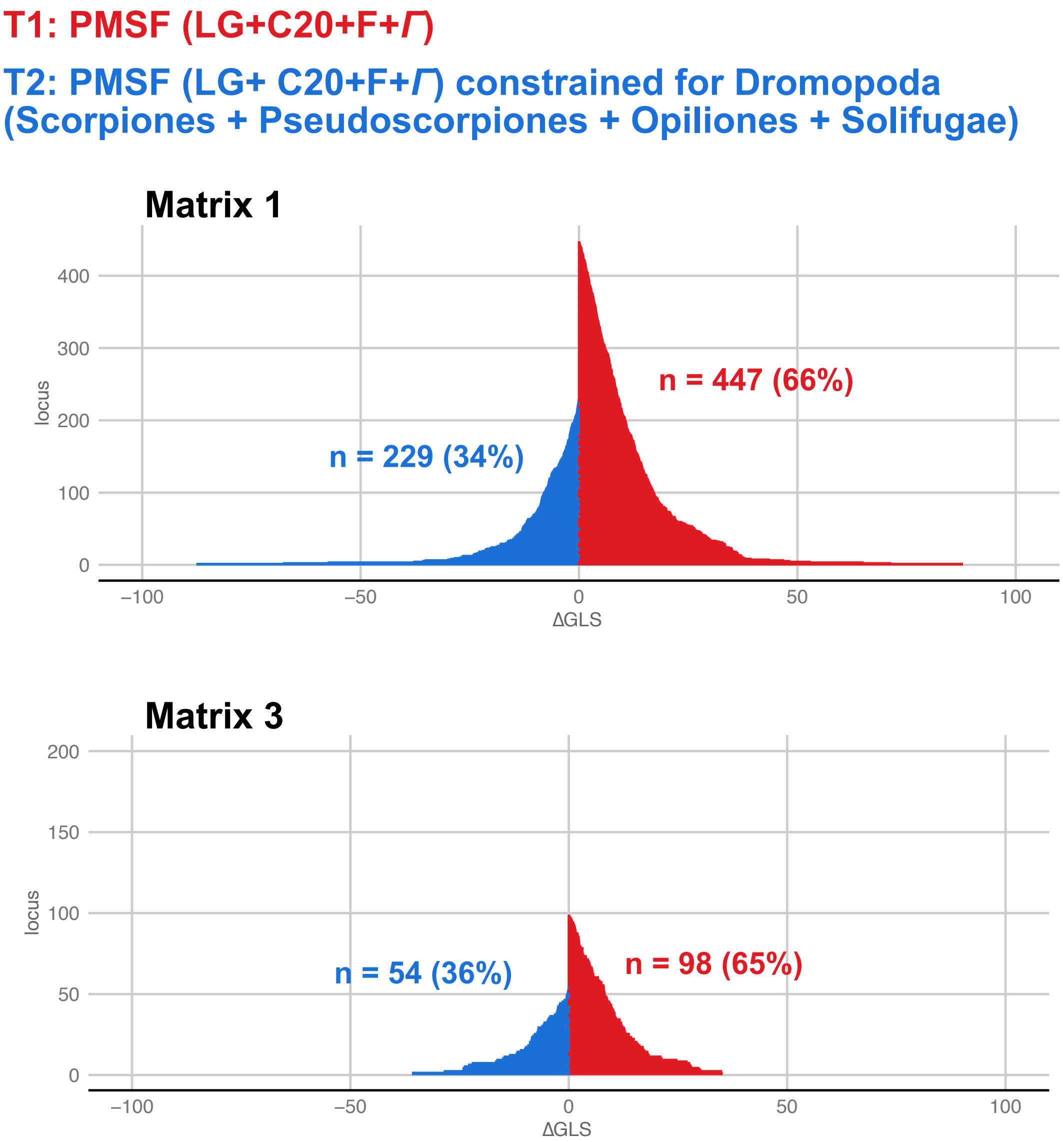

**Fig. S4.** Distributions of Shannon entropy for ∆SLS categories supporting unconstrained topologies (red) versus constrained topologies (blue). Left: Shannon entropy distributions for Dromopoda. Right: Shannon entropy distributions for Arachnida. Note that there are fewer sites supporting Arachnida than Dromopoda, with the latter grouping refuted by phylogenomics and rare genomic changes.

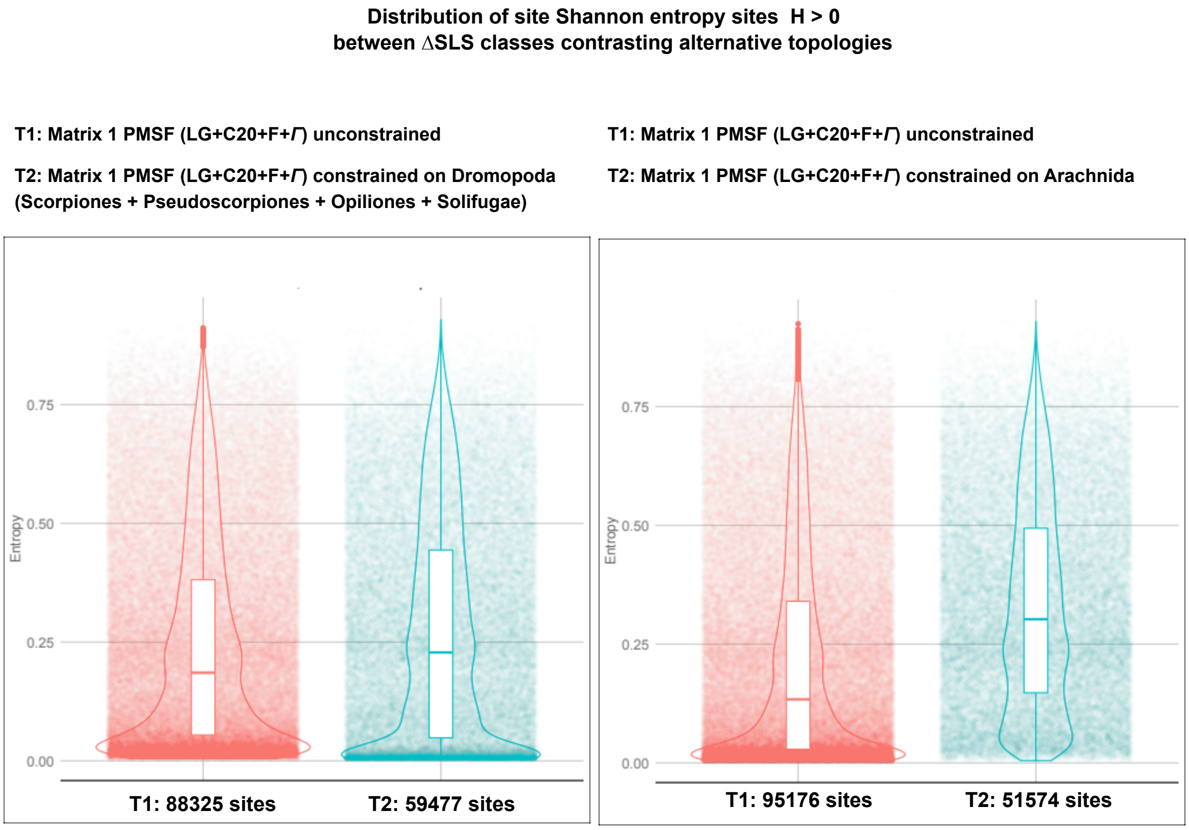

**Fig. S5.** ΔGLS distributions mapping phylogenetic support for the competing hypotheses of pseudoscorpion placement, for Matrices 1 and **3**. Genes filtered only for taxon decisiveness (Matrix 1) have uninformative distributions of gene support for competing topologies. Genes further filtered to retain the least saturated subset (Matrix **3**) show greater support for Panscorpiones (Pseudoscorpiones + Scorpiones), a group that has been validated by rare genomic changes.

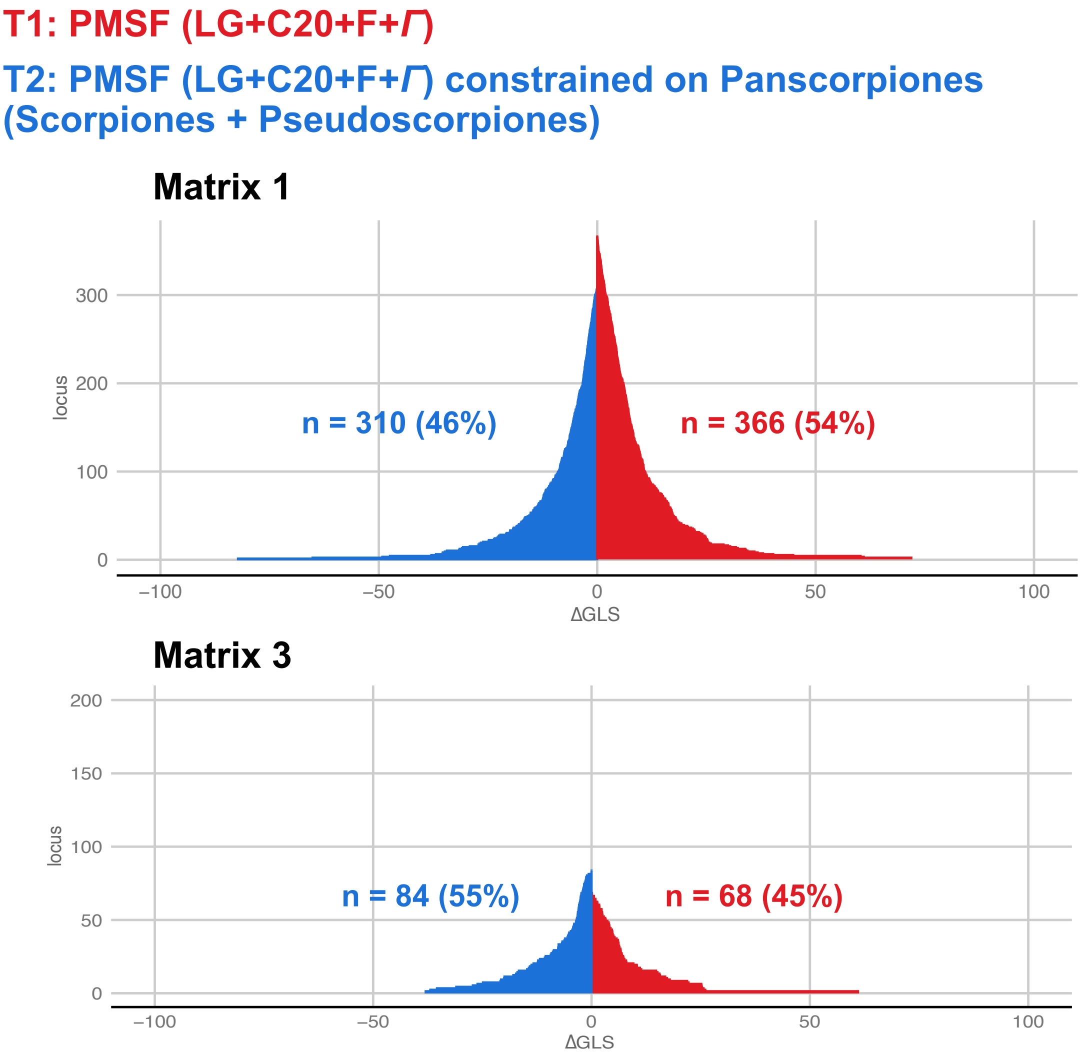

**Fig. S6.** Integration of morphological and molecular datasets refutes arachnid monophyly. (a) Strict consensus tree from equal weights parsimony analysis of Bicknell et al. dataset, combined with Matrix 3. (b) Implied weights parsimony analysis of Bicknell et al. dataset, combined with Matrix 3. Concavity constant *k* = 6. (c) Strict consensus tree from equal weights parsimony analysis of Siveter et al. dataset, combined with Matrix 3. (d) Implied weights parsimony analysis of Siveter et al. dataset, combined with Matrix 3. Concavity constant *k* = 6. Colored branches correspond to Pycnogonida (red), Merostomata (blue), and Arachnida (green). Tree files for other *k* values explored in implied weights framework for both datasets are provided on FigShare.

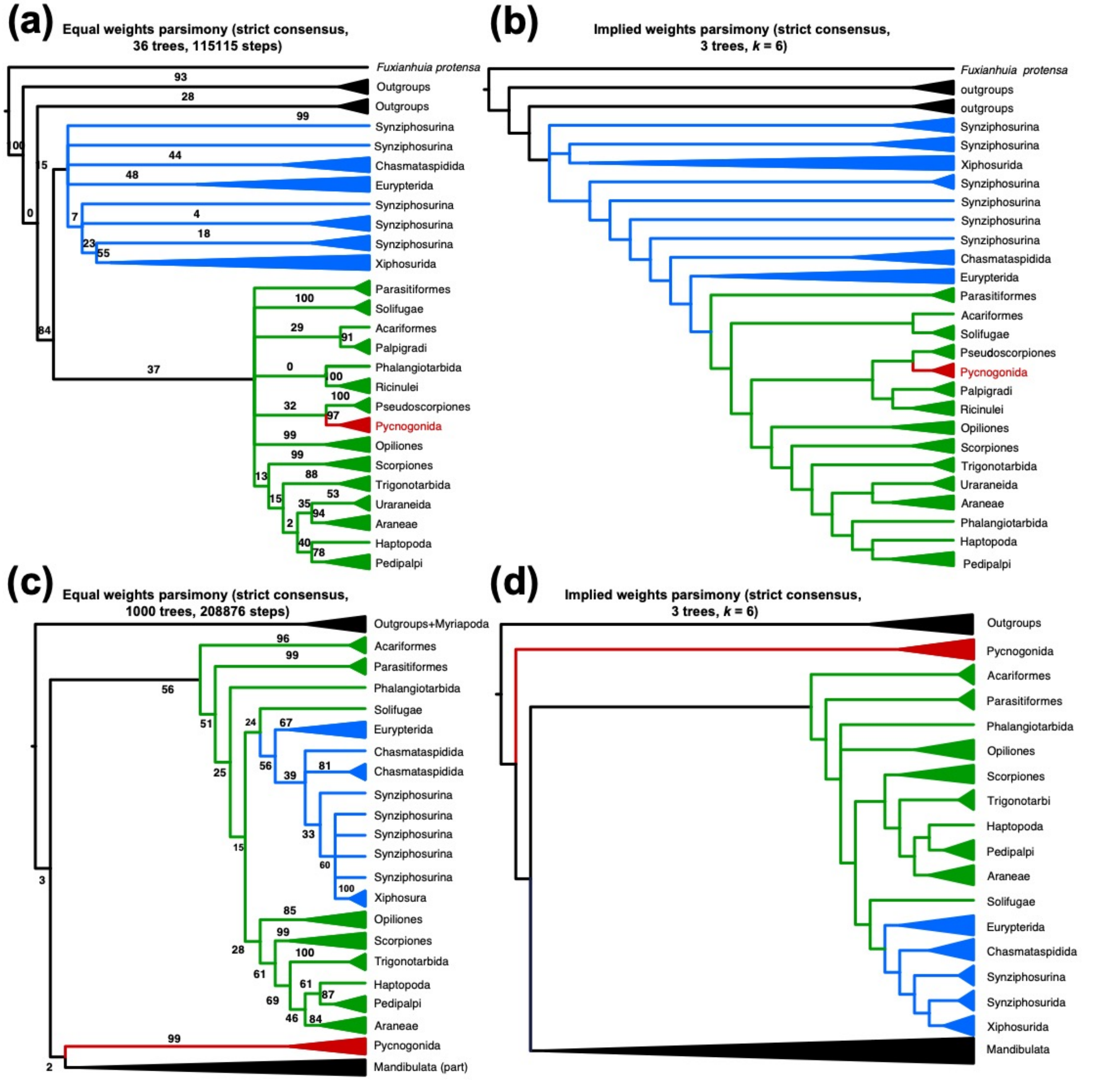

**Table S1.** Species sampling and accession data.

| Terminal | Taxonomy | Accession number | Sampled for Matrix 4 |
|---|---|---|---|
| Epiperipatus sp. MCZ:IZ-141126 | Onychophora | SRR8320992 | X |
| Opisthopatus kwazululandi MCZ:IZ-131434 | Onychophora | SRR8318947 | X |
| Peripatopsis overbergiensis MCZ:IZ-131372 | Onychophora | SRR1145776 | X |
| Eudigraphis taiwaniensis MCZ:IZ-128912 | Myriapoda: Diplopoda | SRR3458640 | X |
| Narceus americanus MCZ:IZ-44069 | Myriapoda: Diplopoda | SRR3233222 | |
| Pauropus huxleyi MCZ:IZ-141222 | Myriapoda: Pauropoda | SRR6145369 | X |
| Symphylella sp. MCZ:IZ-141598 | Myriapoda: Symphyla | SRR6144316 | |
| Scutigera coleoptrata MCZ:IZ-20415 | Myriapoda: Chilopoda | SRR1158078 | |
| Craterostigmus crabilli MCZ:IZ-71256 | Myriapoda: Chilopoda | SRR3232915 | |
| Lithobius atkinsoni | Myriapoda: Chilopoda | SRR7879352 | X |
| Alipes grandidieri MCZ:IZ-130616 | Myriapoda: Chilopoda | SRR619311 | |
| Strigamia maritima | Myriapoda: Chilopoda | GCA_000239455.1 | |
| Eusarsiella sp. | Pancrustacea: Ostracoda | SRR4113497 | X |
| Daphnia magna | Pancrustacea: Branchiopoda | GCA_003990815.1 | |
| Daphnia pulex | Pancrustacea: Branchiopoda | GCA_900092285.2 | |
| Eurytemora affinis | Pancrustacea: Copepoda | GCF000591075.2 | |
| Hyalella azteca | Pancrustacea: Malacostraca | GCA_000764305.3 | |
| Folsomia candida | Pancrustacea: Hexapoda | GCA_002217175.1 | |
| Orchesella cincta | Pancrustacea: Hexapoda | GCA_001718145.1 | |
| Gryllus bimaculatus | Pancrustacea: Hexapoda | pending | |
| Zootermopsis nevadensis | Pancrustacea: Hexapoda | GCF_000696155.1 | |
| Nasonia vitripennis | Pancrustacea: Hexapoda | GCF_009193385.2 | |
| Drosophila melanogaster | Pancrustacea: Hexapoda | GCF_000001215.4 | X |
| Tribolium castaneum | Pancrustacea: Hexapoda | GCA_000002335.3 | X |
| Palaeopantopus | Chelicerata: Pycnogonida | fossil | |
| Palaeoisopus | Chelicerata: Pycnogonida | fossil | |
| Flagellopantopus | Chelicerata: Pycnogonida | fossil | |
| Haliestes dasos | Chelicerata: Pycnogonida | fossil | |
| Tanystylum orbiculare | Chelicerata: Pycnogonida | SRR13153832 | X |
| Nymphon molleri | Chelicerata: Pycnogonida | SRR13153829 | X |
| Anoplodactylus insignis | Chelicerata: Pycnogonida | SRR5237777 | |
| Phoxichilidium femoratum | Chelicerata: Pycnogonida | SRR13153828 | |
| Pycnogonum litorale | Chelicerata: Pycnogonida | SRR7879353 | X |
| Pallenella flava | Chelicerata: Pycnogonida | SRR13153831 | |
| Stylopallene cheilorhynchus | Chelicerata: Pycnogonida | SRR13153830 | |
| Dibasterium | Chelicerata: Synziphosurina | fossil | |
| Offacolus | Chelicerata: Synziphosurina | fossil | |
| Weinbergina | Chelicerata: Synziphosurina | fossil | |
| Limulus polyphemus | Chelicerata: Xiphosura | GCA_000517525.1 | X |
| Carcinoscorpius rotundicauda | Chelicerata: Xiphosura | GCA_011833715.1 | X |
| Tachypleus tridentatus | Chelicerata: Xiphosura | GCA_004210375.1 | X |
| Tachypleus gigas | Chelicerata: Xiphosura | GCA_014155125.1 I | X |
| Parastylonurus | Chelicerata: Eurypterida | fossil | |
| Mixopterus | Chelicerata: Eurypterida | fossil | |
| Eurypterus | Chelicerata: Eurypterida | fossil | |
| Chasmataspis | Chelicerata: Chasmataspidida | fossil | |
| Octoberaspis | Chelicerata: Chasmataspidida | fossil | |
| Eukoenenia hanseni MCZ:IZ-74291 | Chelicerata: Palpigradi | pending | X |
| Eukoenenia spelaea MCZ:IZ-60890 | Chelicerata: Palpigradi | pending | X |
| Eukoenenia spelaea | Chelicerata: Palpigradi | SRR8080647 | X |
| Galeodes sp. | Chelicerata: Solifugae | SRR8080645 | X |
| Solpugema sp. MCZ:IZ-49525 | Chelicerata: Solifugae | pending | X |
| Eremobates sp. MCZ:IZ-49755 | Chelicerata: Solifugae | SRR1146672 | |

| | | | |
|---|---|---|---|
| Eremobates sp. | Chelicerata: Solifugae | pending | X |
| Adenacarus sp. | Chelicerata: Parasitiformes | SRR8080646 | |
| Neocarus sp. | Chelicerata: Parasitiformes | pending | X |
| Ricinoides atewa MCZ:IZ-130073 | Chelicerata: Ricinulei | SRR1145743 | |
| Ricinoides karschii MCZ:IZ-130083 | Chelicerata: Ricinulei | SRR1972991 | X |
| Pseudocellus pearsei MCZ:IZ-16426 | Chelicerata: Ricinulei | SRR1146686 | X |
| Pseudocellus sp. MCZ:IZ-140060 | Chelicerata: Ricinulei | SRR13590369 | |
| Cryptocellus sp. MCZ:IZ-143922 | Chelicerata: Ricinulei | SRR13590368 | |
| Cryptocellus becki MCZ:IZ-136532 | Chelicerata: Ricinulei | SRR1979416 | X |
| Cryptocellus sp. MCZ:IZ-30913 | Chelicerata: Ricinulei | SRR1982218 | |
| Galendromus occidentalis | Chelicerata: Parasitiformes | GCA_000255335.1 | X |
| Gromphadorholaelaps schaeferi MCZ:IZ-71202 | Chelicerata: Parasitiformes | NEW | |
| Varroa destructor | Chelicerata: Parasitiformes | SRR3927486 | X |
| Pneumolaelaps niutirani | Chelicerata: Parasitiformes | SRR10269756 | |
| Tropilaelaps mercedesae | Chelicerata: Parasitiformes | GCA_002081605.1 | |
| Orthinodoros rostratus | Chelicerata: Parasitiformes | SRR1732011 | |
| Ixodes scapularis | Chelicerata: Parasitiformes | GCA_000208615.1 | |
| Haemaphysalis longicornis | Chelicerata: Parasitiformes | SRR7754709 | |
| Amblyomma americanum | Chelicerata: Parasitiformes | SRR4416251 SRR4416250 | |
| Dermacentor silvarum | Chelicerata: Parasitiformes | GCA_013339745.1 | X |
| Hyalomma excavatum | Chelicerata: Parasitiformes | SRR3157672 | |
| Rhipicephalus microplus | Chelicerata: Parasitiformes | SRR1186998 | |
| Labidostommatidae sp. MCZ:IZ-67347 | Chelicerata: Acariformes | pending | |
| Dinothrombium tinctorium | Chelicerata: Acariformes | GCA_003675995.1 | X |
| Panonychus citri | Chelicerata: Acariformes | SRR341928 | |
| Tetranychus cinnabarinus | Chelicerata: Acariformes | SRR519097 | |
| Tetranychus urticae | Chelicerata: Acariformes | GCA_000239435.1 | |
| Rhizoglyphus robini | Chelicerata: Acariformes | SRR946953 | |
| Sarcoptes scabiei | Chelicerata: Acariformes | GCA_014595675.1 | X |
| Dermatophagoides farinae | Chelicerata: Acariformes | SRR1016494 | |
| Dermatophagoides pteronyssinus | Chelicerata: Acariformes | GCA_001901225.2 | |
| Euroglyphus maynei | Chelicerata: Acariformes | GCA_002135145.1 | |
| Hypochthonius rufulus | Chelicerata: Acariformes | SRR4039020 | X |
| Steganacarus magnus | Chelicerata: Acariformes | SRR4039023 | |
| Achipteria coleoptrata | Chelicerata: Acariformes | SRR4039018 | |
| Alaskozetes antarcticus | Chelicerata: Acariformes | SRR6451453, SRR6451455, SRR6451457 | |
| Hermannia gibba | Chelicerata: Acariformes | SRR4039019 | |
| Archegozetes longisetosus | Chelicerata: Acariformes | pending | |
| Nothrus palustris | Chelicerata: Acariformes | SRR4039021 | X |
| Platynothrus peltifer | Chelicerata: Acariformes | SRR4039022 | |
| Idmonarachne | Chelicerata: Uraraneida | fossil | |
| Plesiosiro | Chelicerata: Haptopoda | fossil | |
| Eophrynus | Chelicerata: Trigonotarbida | fossil | |
| Palaeocharinus | Chelicerata: Trigonotarbida | fossil | |
| Anthracomartus | Chelicerata: Trigonotarbida | fossil | |
| Mastigoproctus giganteus | Chelicerata: Uropygi | pending | X |
| Mayacentrum lacandonensis MCZ:IZ-71253 | Chelicerata: Uropygi | SRR9332012 | X |
| Draculoides vinei MCZ:IZ-46896 | Chelicerata: Schizomida | pending | X |
| Stenochrus portoricensis | Chelicerata: Schizomida | SRR6997625 | X |
| Stenochrus sp. MCZ:IZ-74289 | Chelicerata: Schizomida | pending | |
| Graeophonus | Chelicerata: Amblypygi | fossil | |
| Paracharonopsis | Chelicerata: Amblypygi | fossil | |
| Kronocharon | Chelicerata: Amblypygi | fossil | |
| Charinus ioaniticus | Chelicerata: Amblypygi | SRR12389147-148 | X |
| Charinus israelensis | Chelicerata: Amblypygi | SRR12389135-140 | |
| Damon variegatus MCZ:IZ-29740 | Chelicerata: Amblypygi | SRR1145694 | |
| Acanthophrynus coronatus MCZ:IZ-71252 | Chelicerata: Amblypygi | pending | |
| Phrynus marginemaculatus | Chelicerata: Amblypygi | SRR12232018 | X |

| | | | |
|---|---|---|---|
| Attercopus fimbriunguis | Chelicerata: Uraraneida | fossil | |
| Permarachne novokshonovi | Chelicerata: Uraraneida | fossil | |
| Chimerarachne yingi | Chelicerata: Uraraneida | fossil | |
| Liphistius malayanus | Chelicerata: Araneae | SRR1145736 | X |
| Liphistius sp. | Chelicerata: Araneae | SRR1514873 | |
| Sphodros rufipes | Chelicerata: Araneae | SRR1514908 | |
| Megahexura fulva | Chelicerata: Araneae | SRR1514891 | |
| Aliatypus coylei | Chelicerata: Araneae | SRR1514876 | |
| Antrodiaetus unicolor | Chelicerata: Araneae | SRR1514897 | |
| Porrhothele sp. | Chelicerata: Araneae | SRR6997604 | |
| Microhexura montivaga | Chelicerata: Araneae | SRR1514890 | |
| Macrothele calpeiana | Chelicerata: Araneae | SRR6994009 | |
| Paratropis sp. | Chelicerata: Araneae | SRR1514893 | |
| Cyclocosmia truncata | Chelicerata: Araneae | SRR1514884 | |
| Hebestatis theveneti | Chelicerata: Araneae | SRR1514887 | |
| Idiops bersebaensis | Chelicerata: Araneae | SRR1514907 | |
| Aptostichus stephencolberti | Chelicerata: Araneae | SRR1514874 | |
| Promyrmekiaphila clathrata | Chelicerata: Araneae | SRR1514896 | |
| Brachythele longitarsis | Chelicerata: Araneae | SRR1514875 | |
| Pionothele sp. | Chelicerata: Araneae | SRR1514906 | |
| Stanwellia sp. | Chelicerata: Araneae | SRR6997603 | |
| Damarchus sp. | Chelicerata: Araneae | SRR3144092 | |
| Trichopelma laselva | Chelicerata: Araneae | SRR1514881 | |
| Acanthoscurria geniculata | Chelicerata: Araneae | GCA_000661875.1 | |
| Aphonopelma hentzi | Chelicerata: Araneae | pending | X |
| Aphonopelma iviei | Chelicerata: Araneae | SRR1514871 | |
| Filistata insidiatrix | Chelicerata: Araneae | SRR6997865 | |
| Kukulcania hibernalis | Chelicerata: Araneae | SRR1514878 | |
| Hypochilus gertschi | Chelicerata: Araneae | SRR6997860 | |
| Hypochilus pococki | Chelicerata: Araneae | SRR1514889 | |
| Calponia harrisonfordi | Chelicerata: Araneae | SRR3144089 | |
| Segestria sp. | Chelicerata: Araneae | SRR3144084 | |
| Maoriata sp. | Chelicerata: Araneae | SRR6997868 | |
| Orsolobidae sp. | Chelicerata: Araneae | SRR6998651 | |
| Dysdera crocata | Chelicerata: Araneae | SRR1328258 | |
| Ischnothyreus sp. | Chelicerata: Araneae | SRR6997859 | |
| Opopaea sp. | Chelicerata: Araneae | SRR6998659 | |
| Diguetia sp. | Chelicerata: Araneae | SRR3144093 | |
| Pholcus manueli | Chelicerata: Araneae | SRR1365208 | |
| Pholcus phalangioides | Chelicerata: Araneae | SRR1514900 | |
| Loxosceles deserta | Chelicerata: Araneae | SRR3144077 | |
| Drymusa sp. | Chelicerata: Araneae | SRR6997739 | |
| Periegops suterii | Chelicerata: Araneae | SRR6998656 | |
| Ochyrocera sp. | Chelicerata: Araneae | SRR7028536 | |
| Scytodes globula | Chelicerata: Araneae | SRR6998911 | |
| Scytodes thoracica | Chelicerata: Araneae | SRR1514872 | |
| Archoleptoneta sp. | Chelicerata: Araneae | SRR1514872 | |
| Calileptoneta californica | Chelicerata: Araneae | SRR3144085 | |
| Austrochilus forsteri | Chelicerata: Araneae | SRR6997749 | |
| Hickmania troglodytes | Chelicerata: Araneae | SRR6997862 | |
| Progradungula otwayensis | Chelicerata: Araneae | SRR6998916 | |
| Tarlina sp. | Chelicerata: Araneae | SRR6998918 | |
| Gradungula sorenseni | Chelicerata: Araneae | SRR6997863 | |
| Pianoa isolata | Chelicerata: Araneae | SRR6998914 | |
| Austrarchaea sp. | Chelicerata: Araneae | SRR6997750 | |
| Mecysmauchenius sp. | Chelicerata: Araneae | SRR6997871 | |
| Huttonia palpimanoides | Chelicerata: Araneae | SRR6997861 | |
| Otiothops birabeni | Chelicerata: Araneae | SRR6998652 | |

| | | | |
|---|---|---|---|
| Palpimanus gibbulus | Chelicerata: Araneae | SRR6998653 | |
| Stegodyphus mimosarum | Chelicerata: Araneae | GCA_000611955.2 | |
| Megadictyna thilenii | Chelicerata: Araneae | SRR6997870 | |
| Nicodamidae sp. 2 | Chelicerata: Araneae | SRR6998661 | |
| Nicodamidae sp. 1 | Chelicerata: Araneae | SRR7028539 | |
| Mysmenella jobi | Chelicerata: Araneae | SRR6998907 | |
| Novanapis sp. | Chelicerata: Araneae | SRR6998663 | |
| Anelosimus eximius | Chelicerata: Araneae | SRR6997627 | |
| Latrodectus tredecimguttatus | Chelicerata: Araneae | SRR954929 | |
| Euryopis sp. | Chelicerata: Araneae | SRR6997867 | |
| Parasteatoda tepidariorum | Chelicerata: Araneae | GCA_000365465.2 | X |
| Pararchaea alba | Chelicerata: Araneae | SRR6998655 | |
| Malkara sp. | Chelicerata: Araneae | SRR6997869 | |
| Malkaridae sp. | Chelicerata: Araneae | SRR6997874 | |
| Perissopmeros sp. | Chelicerata: Araneae | SRR6998657 | |
| Australomimetus clean | Chelicerata: Araneae | SRR6997751 | |
| Ero leonina | Chelicerata: Araneae | SRR1514886 | |
| Arkys sp. | Chelicerata: Araneae | SRR6997752 | |
| Demadiana sp. | Chelicerata: Araneae | SRR6997740 | |
| Leucauge venusta | Chelicerata: Araneae | SRR1145740 | |
| Meta ovalis | Chelicerata: Araneae | SRR6997876 | |
| Nanometa sp. | Chelicerata: Araneae | SRR6998667 | |
| Tetragnatha tantalus | Chelicerata: Araneae | SRR1427108 | |
| Cepheia longiseta | Chelicerata: Araneae | SRR6997742 | |
| Mysmena leichardti | Chelicerata: Araneae | SRR6998665 | |
| Microdipoena guttata | Chelicerata: Araneae | SRR1333842 | |
| Spinanapis sp. | Chelicerata: Araneae | SRR6998666 | |
| Tekelloides sp. 2 | Chelicerata: Araneae | SRR6998920 | |
| Laminacauda parvipalpis | Chelicerata: Araneae | SRR6997873 | X |
| Frontinella communis | Chelicerata: Araneae | SRR1145739 | |
| Novafroneta gladiatrix | Chelicerata: Araneae | SRR6998662 | |
| Synotaxus cf turbinatus | Chelicerata: Araneae | SRR6998908 | |
| Nesticus bishopi | Chelicerata: Araneae | SRR1655191 | |
| Nesticus cooperi | Chelicerata: Araneae | SRR1514892 | |
| Meringa sp. | Chelicerata: Araneae | SRR6997877 | |
| Runga sp. | Chelicerata: Araneae | SRR6998910 | |
| Physoglenes sp. | Chelicerata: Araneae | SRR6998913 | |
| Physoglenidae sp. | Chelicerata: Araneae | SRR7028534 | |
| Ogulnius sp. | Chelicerata: Araneae | SRR7028537 | |
| Baalzebub sp. | Chelicerata: Araneae | SRR6997748 | |
| Theridiosoma gemmosum | Chelicerata: Araneae | SRR6998922 | |
| Theridiosoma savannum | Chelicerata: Araneae | SRR7028533 | |
| Trichonephila clavipes | Chelicerata: Araneae | GCA_002102615.1 | |
| Micrathena gracilis | Chelicerata: Araneae | SRR1514882 | |
| Verrucosa arenata | Chelicerata: Araneae | SRR3144087 | |
| Cyrtophora sp. | Chelicerata: Araneae | SRR6997738 | |
| Neoscona arabesca | Chelicerata: Araneae | SRR1145741 | |
| Gasteracantha hasselti | Chelicerata: Araneae | SRR1048659 | |
| Macracantha arcuata | Chelicerata: Araneae | SRR1048826 | |
| Menneus sp. | Chelicerata: Araneae | SRR6997878 | |
| Deinopis sp. | Chelicerata: Araneae | SRR6997737 | |
| Deinopis longipes | Chelicerata: Araneae | SRR1514879 | |
| Tamopsis sp. | Chelicerata: Araneae | SRR6998917 | |
| Oecobius navus | Chelicerata: Araneae | SRR1514899 | |
| Oecobius cellariorum | Chelicerata: Araneae | SRR1365089 | |
| Cybaeodamus taim | Chelicerata: Araneae | SRR6997744 | |
| Forsterella sp. | Chelicerata: Araneae | SRR6997864 | |
| Caayguara ybytyriguara | Chelicerata: Araneae | SRR6997747 | |

| | | |
|---|---|---|
| Amaurobius ferox | Chelicerata: Araneae | SRR1329250 |
| Callobius sp. | Chelicerata: Araneae | SRR3144088 |
| Neoramia sp. | Chelicerata: Araneae | SRR6998660 |
| Amphinecta sp. | Chelicerata: Araneae | SRR6997628 |
| Metaltella simoni | Chelicerata: Araneae | SRR3144078 |
| Cambridgea sp. | Chelicerata: Araneae | SRR6997745 |
| Paramatachia sp. | Chelicerata: Araneae | SRR6998654 |
| Agelenopsis emertoni | Chelicerata: Araneae | SRR1514895 |
| Agelenopsis pennsylvanica | Chelicerata: Araneae | SRR1329248 |
| Calymmaria sp. | Chelicerata: Araneae | SRR6997746 |
| Calymmaria persica | Chelicerata: Araneae | SRR3144091 |
| Cicurina travisae | Chelicerata: Araneae | SRR1654705 |
| Cicurina vibora | Chelicerata: Araneae | SRR1514883 |
| Homalonychus theologus | Chelicerata: Araneae | SRR3144075 |
| Tengella radiata | Chelicerata: Araneae | SRR7028532 |
| Uliodon sp. | Chelicerata: Araneae | SRR6998923 |
| Misumenoides formosipes | Chelicerata: Araneae | SRR3144080 |
| Sidymella sp. | Chelicerata: Araneae | SRR6998912 |
| Anahita punctulata | Chelicerata: Araneae | SRR3144072 |
| Peucetia longipalpis | Chelicerata: Araneae | SRR1514898 |
| Dolomedes triton | Chelicerata: Araneae | SRR3144094 |
| Pisaurina mira | Chelicerata: Araneae | SRR1365651 |
| Cupiennius sp. | Chelicerata: Araneae | SRR7028538 |
| Allocosa alticeps | Chelicerata: Araneae | SRR6997626 |
| Schizocosa rovneri | Chelicerata: Araneae | SRR1514894 |
| Habronattus signatus | Chelicerata: Araneae | SRR1514888 |
| Habronattus ustulatus | Chelicerata: Araneae | SRR1656783 |
| Teminius sp. | Chelicerata: Araneae | SRR6998921 |
| Karaops raveni | Chelicerata: Araneae | SRR6997858 |
| Falconina gracilis | Chelicerata: Araneae | SRR6997866 |
| Nyssus sp. | Chelicerata: Araneae | SRR6998658 |
| Clubiona sp. | Chelicerata: Araneae | SRR6997741 |
| Rebilus sp. | Chelicerata: Araneae | SRR6998909 |
| Liocranidae sp. | Chelicerata: Araneae | SRR6997875 |
| Trachelas tranquillus | Chelicerata: Araneae | SRR1329247 |
| Sergiolus capulatus | Chelicerata: Araneae | SRR1514903 |
| Molycria sp. | Chelicerata: Araneae | SRR6998664 |
| Anzacia sp. | Chelicerata: Araneae | SRR6997629 |
| Lampona sp. | Chelicerata: Araneae | SRR6997872 |
| Zosis sp. | Chelicerata: Araneae | SRR6998924 |
| Philoponella herediae | Chelicerata: Araneae | SRR1514880 |
| Uloborus glomosus | Chelicerata: Araneae | SRR1328334 |
| Eophalangium sheari | Chelicerata: Opiliones | fossil |
| Hastocularis argus | Chelicerata: Opiliones | fossil |
| Leptopsalis sp. MCZ:IZ-141287 | Chelicerata: Opiliones | pending |
| Miopsalis sp. MCZ:IZ-140867 | Chelicerata: Opiliones | SRR5234539 |
| Parapurcellia silvicola MCZ:IZ-134743 | Chelicerata: Opiliones | SRX6374208 |
| Purcellia illustrans MCZ:IZ-49518 | Chelicerata: Opiliones | SRX6374206 |
| Chileogovea oedipus MCZ:IZ-138106 | Chelicerata: Opiliones | SRR9611065 |
| Rakaia minutissima MCZ:IZ-143240 | Chelicerata: Opiliones | SRR9611073 |
| Rakaia stewartiensis MCZ:IZ-49801 | Chelicerata: Opiliones | SRR9611075 |
| Rakaia magna australis MCZ:IZ-29212 | Chelicerata: Opiliones | SRR5181449 |
| Rakaia pauli MCZ:IZ-144621 | Chelicerata: Opiliones | SRR9611074 |
| Pettalus thwaitesi MCZ:IZ-132349 | Chelicerata: Opiliones | SRX6374207 |
| Austropurcellia alata MCZ:IZ-141418 | Chelicerata: Opiliones | SRR9611069 |
| Austropurcellia despectata MCZ:IZ-141417 | Chelicerata: Opiliones | SRR9611064 |
| Austropurcellia cf acuta MCZ:IZ-141419 | Chelicerata: Opiliones | SRR9611062 |
| Austropurcellia acuta MCZ:IZ-44060 | Chelicerata: Opiliones | SRR9611068 |

| | | | |
|---|---|---|---|
| Austropurcellia clousei MCZ:IZ-141420 | Chelicerata: Opiliones | SRR9611063 | X |
| Neopurcellia salmoni MCZ:IZ-144625 | Chelicerata: Opiliones | SRR9611061 | |
| Neopurcellia salmoni MCZ:IZ-29310 | Chelicerata: Opiliones | SRR5181448 | |
| Karripurcellia peckorum MCZ:IZ-49989 | Chelicerata: Opiliones | SRR9611060 | |
| Aoraki denticulata Northern MCZ:IZ-141172 | Chelicerata: Opiliones | SRR9611067 | |
| Aoraki denticulata Southern MCZ:IZ-141173 | Chelicerata: Opiliones | SRR9611066 | |
| Aoraki longitarsa MCZ:IZ-137238 | Chelicerata: Opiliones | SRR5181577 | |
| Siro boyerae MCZ:IZ-137238 | Chelicerata: Opiliones | SRR1145699 | X |
| Metagovea oviformis MCZ:IZ-136533 | Chelicerata: Opiliones | SRR5230765 | |
| Metasiro savannahensis MCZ:IZ-133799 | Chelicerata: Opiliones | SRX205297 | |
| Brasilogovea microphaga MCZ:IZ-136559 | Chelicerata: Opiliones | SRR5185916 | |
| Neogovea matawai MCZ:IZ-133922 | Chelicerata: Opiliones | SRX6098501 | |
| Acropsopilio neozealandiae MCZ:IZ-30457 | Chelicerata: Opiliones | SRR5235984 | |
| Hesperonemastoma modestum | Chelicerata: Opiliones | SRX450937 | |
| Ortholasma coronadense | Chelicerata: Opiliones | SRX451776 | |
| Trogulus martensi | Chelicerata: Opiliones | SRX450964 | |
| Caddo agilis MCZ:IZ-40415 | Chelicerata: Opiliones | pending | |
| Thrasychirus modestus MCZ:IZ-49762 | Chelicerata: Opiliones | SRR5234747 | |
| Forsteropsalis pureora MCZ:IZ-29216 | Chelicerata: Opiliones | SRR5234746 | |
| Pantopsalis cheliferoides MCZ:IZ-133328 | Chelicerata: Opiliones | pending | |
| Mangatangi parvum MCZ:IZ-133381 | Chelicerata: Opiliones | pending | |
| Megalopsalis triascuta MCZ:IZ-133420 | Chelicerata: Opiliones | pending | |
| Diguetinus raptator MCZ:IZ-143123 | Chelicerata: Opiliones | pending | |
| Phalangium opilio MCZ:IZ-29189 | Chelicerata: Opiliones | SRR1145735 | |
| Protolophus singularis | Chelicerata: Opiliones | SRX450934 | |
| Prionostemma vittatum MCZ:IZ-144108 | Chelicerata: Opiliones | pending | X |
| Leiobunum verrucosum | Chelicerata: Opiliones | SRX450936 | |
| Paranelima sp. MCZ:IZ-143124 | Chelicerata: Opiliones | pending | |
| Sclerobunus robustus | Chelicerata: Opiliones | SRX647390 | |
| Synthetonychia glacialis MCZ:IZ-137212 | Chelicerata: Opiliones | SRR5236909 | |
| Larifuga capensis MCZ:IZ-49747 | Chelicerata: Opiliones | SRR1145742 | |
| Nuncia sp. MCZ:IZ-133135 | Chelicerata: Opiliones | pending | |
| Pristobunus sp. MCZ:IZ-152391 | Chelicerata: Opiliones | pending | |
| Scotolemon lespesi MCZ:IZ-43672 | Chelicerata: Opiliones | SAMN12670423 | |
| Sitalcina lobata | Chelicerata: Opiliones | SRX451777 | |
| Gnomulus rostratus MCZ:IZ-141286 | Chelicerata: Opiliones | SRX2549521 | |
| Dibunus sp. MCZ:IZ-141285 | Chelicerata: Opiliones | SRX2549524 | |
| Epedanus pinangensis MCZ:IZ-141288 | Chelicerata: Opiliones | pending | |
| Haasus judaeus | Chelicerata: Opiliones | pending | |
| Haasus naasane | Chelicerata: Opiliones | SRX5786020 | |
| Dampetrus sp. MCZ:IZ-140868 | Chelicerata: Opiliones | SRX2549519 | |
| Metabiantes sp. MCZ:IZ-49748 | Chelicerata: Opiliones | SRX451770 | X |
| Stygnomma sp. MCZ:IZ-144060 | Chelicerata: Opiliones | pending | |
| Pellobunus sp. | Chelicerata: Opiliones | SRX2549520 | |
| Kimulidae sp. MCZ:IZ-143960 | Chelicerata: Opiliones | pending | |
| Tegipiolus pachypus MZSP71018 | Chelicerata: Opiliones | SAMN12670426 | |
| Zalmoxidae gen. sp. MCZ:IZ-144055 | Chelicerata: Opiliones | pending | |
| Pachylicus acutus MCZ:IZ-49749 | Chelicerata: Opiliones | SRX451775 | |
| Maracaynatum trinidaense MCZ:IZ-144047 | Chelicerata: Opiliones | pending | |
| Avima matintaperera MCZ:IZ-136560 | Chelicerata: Opiliones | SRX2548330 | |
| Karos barbarikos MCZ:IZ-46444 | Chelicerata: Opiliones | pending | |
| Chapulobunus sp. MCZ:IZ-46443 | Chelicerata: Opiliones | SRX2548334 | |
| Stygnoplus clavotibialis MCZ:IZ-143966 | Chelicerata: Opiliones | pending | |
| Protimesius longipalpis MCZ:IZ-136522 | Chelicerata: Opiliones | SRX2548335 | |
| Auranus hoeferscovitorum MCZ:IZ-136531 | Chelicerata: Opiliones | SAMN12670389 | |
| Pickeliana pickeli MCZ:IZ-139249 | Chelicerata: Opiliones | SAMN12670413 | |
| Gonycranaus pluto MCZ:IZ-139251 | Chelicerata: Opiliones | SAMN12670396 | |
| Camarana flavipalpi MCZ:IZ-139271 | Chelicerata: Opiliones | SAMN12670391 | |

| | | | |
|---|---|---|---|
| Pseudopachylus longipes MCZ:IZ-32171 | Chelicerata: Opiliones | SRR5241471 | |
| Bissula sp. MCZ:IZ-139255 | Chelicerata: Opiliones | SAMN12670390 | |
| Pseudopachylus eximius MCZ:IZ-139280 | Chelicerata: Opiliones | SAMN12670416 | |
| Vonones ornata MCZ:IZ-49750 | Chelicerata: Opiliones | SRR1145738 | |
| Incasarcus argenteus MCZ:IZ-139250 | Chelicerata: Opiliones | SAMN12670404 | |
| Quindina limbata MZSP-71772 | Chelicerata: Opiliones | SAMN12670420 | |
| Saramacia lucasae MCZ:IZ-136518 | Chelicerata: Opiliones | SRR5241473 | |
| Ampycinae sp. MCZ:IZ-46455 | Chelicerata: Opiliones | SAMN12670388 | |
| Glysterus sp. MCZ:IZ-140069 | Chelicerata: Opiliones | SAMN12670394 | |
| Panalus robustus MZSP-71019 | Chelicerata: Opiliones | SAMN12670411 | |
| Phalangodus cottus MZSP-71020 | Chelicerata: Opiliones | SAMN12670412 | |
| Santinezia serratotibialis MCZ:IZ-144078 | Chelicerata: Opiliones | pending | |
| Phareicranaus manauara MCZ:IZ-136554 | Chelicerata: Opiliones | SRX2548333 | |
| Pseudopucrolia discrepans MCZ:IZ-139277 | Chelicerata: Opiliones | SAMN12670417 | |
| Sadocus polyacanthus MCZ:IZ-138113 | Chelicerata: Opiliones | SAMN12670422 | |
| Eubalta planiceps MCZ:IZ-49760 | Chelicerata: Opiliones | SAMN12670392 | |
| Metagyndes innata MCZ:IZ-29745 | Chelicerata: Opiliones | SRR5241474 | |
| Hernandaria una MCZ:IZ-139258 | Chelicerata: Opiliones | SAMN12670403 | |
| Graphinotus therezopolis MCZ:IZ-139273 | Chelicerata: Opiliones | SAMN12670402 | |
| Meteusarcoides sp. MCZ:IZ-139264 | Chelicerata: Opiliones | pending | |
| Meteusarcoides caudatus MZSP-71024 | Chelicerata: Opiliones | SAMN12670407 | |
| Acutisoma longipes MCZ:IZ-139261 | Chelicerata: Opiliones | SAMN12670386 | |
| Goniosoma sp. MCZ:IZ-139266 | Chelicerata: Opiliones | SAMN12670395 | |
| Roeweria virescens MCZ:IZ-139279 | Chelicerata: Opiliones | PRJNA556673 | |
| Longiperna concolor MCZ:IZ-139260 | Chelicerata: Opiliones | SAMN12670406 | |
| Gonyleptidae sp. MCZ IZ-139272 MCZ:IZ-139272 | Chelicerata: Opiliones | pending | |
| Mitobates conspersus MCZ:IZ-139263 | Chelicerata: Opiliones | SAMN12670409 | |
| Promitobates ornatus MZSP-71022 | Chelicerata: Opiliones | SAMN12670415 | |
| Gonyleptellus pustulosus MCZ:IZ-139269 | Chelicerata: Opiliones | SAMN12670397 | |
| Gonyleptes fragilis MZSP-71023 | Chelicerata: Opiliones | SAMN12670399 | |
| Pseudotrogulus mirim MCZ:IZ-139270 | Chelicerata: Opiliones | SAMN12670418 | |
| Pseudotrogulus trostkyi MCZ:IZ-139278 | Chelicerata: Opiliones | SAMN12670419 | |
| Ampheres leucopheus MCZ:IZ-139267 | Chelicerata: Opiliones | SAMN12670387 | |
| Acanthogonyleptes aff fulvigranulatus MZSP-71027 | Chelicerata: Opiliones | SAMN12670385 | |
| Gonyleptes atrus MZSP-71025 | Chelicerata: Opiliones | SAMN12670398 | |
| Gonyleptes sp. MCZ:IZ-139262 | Chelicerata: Opiliones | SAMN12670400 | |
| Geraecormobius bispinifrons MCZ:IZ-139274 | Chelicerata: Opiliones | SAMN12670393 | |
| Mischonyx cuspidatus MCZ:IZ-139265 | Chelicerata: Opiliones | SAMN12670408 | |
| Sodreana leprevosti MCZ:IZ-139256 | Chelicerata: Opiliones | SAMN12670424 | |
| Sodreana sodrena MCZ:IZ-139268 | Chelicerata: Opiliones | SAMN12670425 | |
| Progonyleptoidellus striatus MZSP-71021 | Chelicerata: Opiliones | SAMN12670414 | |
| Iporangaia pustulosa MCZ:IZ-139275 | Chelicerata: Opiliones | SAMN12670405 | |
| Gonyleptoides marumbiensis MCZ:IZ-139254 | Chelicerata: Opiliones | SAMN12670401 | |
| Neosadocus maximus MCZ:IZ-139257 | Chelicerata: Opiliones | SAMN12670410 | |
| Pseudotyrannochthonius sp. B MCZ:IZ-144011 | Chelicerata: Pseudoscorpiones | SRR9331997 | |
| Pseudotyrannochthonius sp. MCZ:IZ-149275 | Chelicerata: Pseudoscorpiones | SRR9331998 | |
| Lechytia hoffi MCZ:IZ-139248 | Chelicerata: Pseudoscorpiones | SRR9331999 | |
| Lagynochthonius australicus MCZ:IZ-143136 | Chelicerata: Pseudoscorpiones | SRR9332016 | |
| Ephippiochthonius tetrachelatus MCZ:IZ-140293 | Chelicerata: Pseudoscorpiones | SRR9331983 | |
| Lechytia hoffi MCZ:IZ-139248 | Chelicerata: Pseudoscorpiones | SRR9332011 | |
| Feaella capensis MCZ:IZ-139240 | Chelicerata: Pseudoscorpiones | SRR9331986 | X |
| Pseudogarypus banksi MCZ:IZ-140301 | Chelicerata: Pseudoscorpiones | SRR9331993 | |

| | | | |
|---|---|---|---|
| Bochica withi MCZ:IZ-144332 | Chelicerata: Pseudoscorpiones | SRR9332002 | |
| Dhanus sumatranus MCZ:IZ-49986 | Chelicerata: Pseudoscorpiones | SRR9331984 | |
| Indohya sp. MCZ:IZ-151267 | Chelicerata: Pseudoscorpiones | SRR9332015 | |
| Ideobisium crassimanum MCZ:IZ-144356 | Chelicerata: Pseudoscorpiones | SRR9331979 | |
| Parahya submersa MCZ:IZ-139241 | Chelicerata: Pseudoscorpiones | SRR9331989 | |
| Gymnobisium sp. MCZ:IZ-139238 | Chelicerata: Pseudoscorpiones | SRR9331987 | |
| Microcreagrinae sp. MCZ:IZ-60626 | Chelicerata: Pseudoscorpiones | SRR9332009 | |
| Microbisium brunneum MCZ:IZ-141276 | Chelicerata: Pseudoscorpiones | SRR9332013 | |
| Microbisium sp. MCZ:IZ-60599 | Chelicerata: Pseudoscorpiones | SRR9332014 | |
| Novobisium sp. MCZ:IZ-141855 | Chelicerata: Pseudoscorpiones | SRR9331992 | X |
| Afrogarypus subimpressus MCZ:IZ-139239 | Chelicerata: Pseudoscorpiones | SRR9332004 | |
| Geogarypus maculatus MCZ:IZ-144335 | Chelicerata: Pseudoscorpiones | SRR9331988 | |
| Apolpium sp. MCZ:IZ-144360 | Chelicerata: Pseudoscorpiones | SRR9332003 | X |
| Anchigarypus californicus MCZ:IZ-149276 | Chelicerata: Pseudoscorpiones | SRR9331985 | X |
| Synsphyronus apimelus | Chelicerata: Pseudoscorpiones | SRR7062201 | |
| Pseudogarypinus frontalis MCZ:IZ-139246 | Chelicerata: Pseudoscorpiones | SRR9331994 | |
| Larca granulata MCZ:IZ-140297 | Chelicerata: Pseudoscorpiones | SRR9332018 | |
| Protogarypinus giganteus | Chelicerata: Pseudoscorpiones | SRR9331995 | |
| Cheiridiidae sp. MCZ:IZ-140294 | Chelicerata: Pseudoscorpiones | SRR9332008 | |
| Sternophoridae sp. MCZ:IZ-133496 | Chelicerata: Pseudoscorpiones | SRR9332020 | |
| Cacodemonius segmentidentatus MCZ:IZ-144363 | Chelicerata: Pseudoscorpiones | SRR9332001 | |
| Oratemnus curtus MCZ:IZ-49983 | Chelicerata: Pseudoscorpiones | SRR9331991 | |
| Protochelifer sp. MCZ:IZ-49984 | Chelicerata: Pseudoscorpiones | SRR9331996 | |
| Cheliferidae sp. MCZ:IZ-40982 | Chelicerata: Pseudoscorpiones | SRR9332007 | |
| Parachelifer persimilis MCZ:IZ-139247 | Chelicerata: Pseudoscorpiones | SRR9331990 | |
| Lamprochernes savignyi MCZ:IZ-149278 | Chelicerata: Pseudoscorpiones | SRR9332017 | |
| Conicochernes crassus MCZ:IZ-49985 | Chelicerata: Pseudoscorpiones | SRR9331982 | |
| Progonyleptoidellus striatus | Chelicerata: Pseudoscorpiones | SRR1514877 | |
| Haplochernes kraepelini | Chelicerata: Pseudoscorpiones | SRR1767661 | |
| Chernetidae sp. MCZ:IZ-142323 | Chelicerata: Pseudoscorpiones | SRR9332006 | X |
| Chernetidae sp. JC032 | Chelicerata: Pseudoscorpiones | SRR9332005 | |
| Waeringoscorpio | Chelicerata: Scorpiones | fossil | |
| Pulmonoscorpius | Chelicerata: Scorpiones | fossil | |
| Palaeoscorpius | Chelicerata: Scorpiones | fossil | |
| Compsoscorpius | Chelicerata: Scorpiones | fossil | |
| Proscorpius | Chelicerata: Scorpiones | fossil | |
| Chaerilus celebensis | Chelicerata: Scorpiones | SRR1721804 | X |
| Lychas variegatus | Chelicerata: Scorpiones | pending | |

| | | | |
|---|---|---|---|
| Mesobuthus martensii | Chelicerata: Scorpiones | GCA_000484575.1 | |
| Compsobuthus levyi | Chelicerata: Scorpiones | pending | |
| Birulatus israelensis | Chelicerata: Scorpiones | pending | |
| Compsobuthus sp. | Chelicerata: Scorpiones | pending | |
| Compsobuthus schmiedeknechti | Chelicerata: Scorpiones | pending | |
| Androctonus amoreuxi | Chelicerata: Scorpiones | pending | |
| Androctonus australis | Chelicerata: Scorpiones | SRR1724216 | |
| Hottentotta trilineatus | Chelicerata: Scorpiones | SRR1721800 | |
| Orthochirus scrobiculosus | Chelicerata: Scorpiones | pending | |
| Androctonus crassicauda | Chelicerata: Scorpiones | pending | |
| Buthus israelensis | Chelicerata: Scorpiones | pending | |
| Buthacus cf arenicola | Chelicerata: Scorpiones | pending | |
| Leirus hebraeus | Chelicerata: Scorpiones | pending | |
| Leirus quinquestriatus | Chelicerata: Scorpiones | pending | |
| Ananteris balzani | Chelicerata: Scorpiones | pending | |
| Babycurus gigas | Chelicerata: Scorpiones | pending | |
| Grosphus grandidieri | Chelicerata: Scorpiones | pending | |
| Parabuthus transvaalicus | Chelicerata: Scorpiones | SRR1721799 | |
| Uroplectes olivaceus | Chelicerata: Scorpiones | pending | |
| Uropolectes vittatus | Chelicerata: Scorpiones | pending | |
| Tityus smithi | Chelicerata: Scorpiones | pending | |
| Tityus marahensis | Chelicerata: Scorpiones | pending | |
| Tityus costatus | Chelicerata: Scorpiones | pending | |
| Tityus serrulatus | Chelicerata: Scorpiones | pending | |
| Troglorhopalurus lacrau | Chelicerata: Scorpiones | pending | |
| Physoctonus debilis | Chelicerata: Scorpiones | pending | |
| Jaguajir agammeneon | Chelicerata: Scorpiones | pending | |
| Jaguajir rochai | Chelicerata: Scorpiones | pending | |
| Heteroctenus junceus | Chelicerata: Scorpiones | pending | |
| Rhopalurus garridoi | Chelicerata: Scorpiones | pending | |
| Centruroides caribensis | Chelicerata: Scorpiones | pending | |
| Centruroides hentzi | Chelicerata: Scorpiones | pending | |
| Centruroides vittatus | Chelicerata: Scorpiones | SRR1515193 | |
| Centruroides sculpturatus | Chelicerata: Scorpiones | SRR1515193 | X |
| Iurus dekanum | Chelicerata: Scorpiones | SRR1721734 | |
| Bothriurus coriaceus | Chelicerata: Scorpiones | SRR6467511 | |
| Bothriurus burmeisteri | Chelicerata: Scorpiones | SRR1721670 | |
| Centromachetes sp. | Chelicerata: Scorpiones | SRR6467879 | |
| Cercophonius squama | Chelicerata: Scorpiones | SRR6470446 | |
| Cercophomius queenslandae | Chelicerata: Scorpiones | SRR6466561 | |
| Cercophonius sulcatus | Chelicerata: Scorpiones | SRR6470146 | |
| Caraboctonus keyserlingii | Chelicerata: Scorpiones | SRX5829265 | |
| Superstitiona donensis | Chelicerata: Scorpiones | SRR1721951 | |
| Anuroctonus phaiodactylus | Chelicerata: Scorpiones | SRR1721879 | |
| Brotheas granulatus | Chelicerata: Scorpiones | SRR1721887 | |
| Scorpiops sp. | Chelicerata: Scorpiones | SRR1767662 | |
| Euscorpius italicus | Chelicerata: Scorpiones | SRR1721892 | |
| Plesiochactas dilutus | Chelicerata: Scorpiones | SRR7250103 | X |
| Megacormus gertschi | Chelicerata: Scorpiones | SRR3657526 | |
| Megacormus sp. | Chelicerata: Scorpiones | SRR1767669 | |
| Uroctonus mordax | Chelicerata: Scorpiones | SRR8518581 | |
| Belisarius xambeui | Chelicerata: Scorpiones | SRR1721953 | |
| Hoffmannihadrurus aztecus | Chelicerata: Scorpiones | ERX3551392 | |
| Hadrurus arizonensis | Chelicerata: Scorpiones | SRR1721733 | |
| Hadrurus concolorous | Chelicerata: Scorpiones | ERX3560276 | |
| Hadrurus spadix | Chelicerata: Scorpiones | SRX2056046 | X |
| Uroctonites huachuca | Chelicerata: Scorpiones | SRR8518582 | |
| Kovarikia boggerti | Chelicerata: Scorpiones | SRX5322100 | |

| | | |
|---|---|---|
| Pseudouroctonus apacheanus | Chelicerata: Scorpiones | SRR8518585 |
| Vaejovis cashi | Chelicerata: Scorpiones | SRR8518583 |
| Parauroctonus baergi | Chelicerata: Scorpiones | SRR7443668 |
| Smeringurus mesaensis | Chelicerata: Scorpiones | SRR7473845 |
| Smeringurus vachoni | Chelicerata: Scorpiones | SRR7474136 |
| Vaejovis mexicanus | Chelicerata: Scorpiones | SRR7421527 |
| Serradigitus gertschi | Chelicerata: Scorpiones | ERX2850109 |
| Konetontil acapulco | Chelicerata: Scorpiones | SRR7422029 |
| Konetontli chamelaensis | Chelicerata: Scorpiones | SRR7427084 |
| Mesomexovis aff variegatus | Chelicerata: Scorpiones | SRR7439652 |
| Paravaejovis schwenkmeyeri | Chelicerata: Scorpiones | ERX2670405 |
| Paravaejovis spinigerus | Chelicerata: Scorpiones | SRR1721954 |
| Kuarapu purepecha | Chelicerata: Scorpiones | SRR7439043 |
| Chihuahuanus coahuilae | Chelicerata: Scorpiones | SRR7439185 |
| Mesovexovis occidentalis | Chelicerata: Scorpiones | SRR7439610 |
| Thorellius intrepidus | Chelicerata: Scorpiones | SRR7427141 |
| Urodacus yaschenkoi | Chelicerata: Scorpiones | SRX685583 |
| Urodacus elongatus | Chelicerata: Scorpiones | SRX4723123 |
| Urodacus planimanus | Chelicerata: Scorpiones | pending |
| Hormiops davidovi | Chelicerata: Scorpiones | pending |
| Hormurus weberi | Chelicerata: Scorpiones | pending |
| Liocheles australasiae | Chelicerata: Scorpiones | SRR1721664 |
| Liocheles sp. KhaoDeng | Chelicerata: Scorpiones | pending |
| Nebo hierochonticus | Chelicerata: Scorpiones | pending |
| Diplocentrus diablo | Chelicerata: Scorpiones | SRR1721672 |
| Diplocentrus zacatecanus | Chelicerata: Scorpiones | pending |
| Kolotl magnus | Chelicerata: Scorpiones | SRX4717881 |
| Kolotl poncei | Chelicerata: Scorpiones | pending |
| Iomachus politus | Chelicerata: Scorpiones | pending |
| Hadogenes troglodytes | Chelicerata: Scorpiones | SRR1721665 |
| Hadogenes paucidens | Chelicerata: Scorpiones | pending |
| Opistacanthus asper | Chelicerata: Scorpiones | pending |
| Opistacanthus validus | Chelicerata: Scorpiones | pending |
| Chiromachus ochropus | Chelicerata: Scorpiones | pending |
| Opisthacanthus madagascarensis | Chelicerata: Scorpiones | SRR1721668 |
| Palaeocheloctonus pauliani | Chelicerata: Scorpiones | pending |
| Opistophthalmus glabrifrons | Chelicerata: Scorpiones | pending |
| Pandinus imperator | Chelicerata: Scorpiones | SRR1721600 |
| Opistophthalmus walhbergi | Chelicerata: Scorpiones | pending |
| Opistophthalmus boehmi | Chelicerata: Scorpiones | pending |
| Scorpio fuscus | Chelicerata: Scorpiones | pending |
| Troglokhammouanus steineri | Chelicerata: Scorpiones | SRR1721739 |
| Vietbocap lao | Chelicerata: Scorpiones | SRR1721740 |
| Goniotarbus | Chelicerata: Phalangiotarbida | fossil |
| Bonatarbus | Chelicerata: Phalangiotarbida | fossil |

**Table S2.** Tests of monophyly for Matrices 1-4.

| | | Arachnida | Acari | Panscorpiones | Poecilophysidea | Dromopoda |
|---|---|---|---|---|---|---|
| AU test | Matrix 1 PMSF | 6.09E-07 | 1.91E-09 | 7.48E-08 | 9.95E-74 | 5.65E-06 |
| | Matrix 2 PMSF | 1.69E-04 | 0.000237 | 0.000178 | 0.000311 | 0.000144 |
| | Matrix 3 PMSF | 4.33E-19 | 6.36E-09 | 0.968 | 1.10E-07 | 3.68E-54 |
| Posterior probability | PhyloBayes-mpi | 0.02813567 | 0 | 1 | 1 | 0 |

**Table S3.** Taxonomic groupings used for defining taxon decisiveness.

| ***Taxon*** | ***Number of exemplars*** |
|---|---|
| Onychophora | 3 |
| Pancrustacea | 12 |
| Myriapoda | 9 |
| Pycnogonida | 7 |
| Acariformes | 18 |
| Parasitiformes | 14 |
| Palpigradi | 3 |
| Opiliones | 118 |
| Solifugae | 4 |
| Xiphosura | 4 |
| Ricinulei | 7 |
| Pseudoscorpiones | 39 |
| Scorpiones | 103 |
| Amblypygi | 5 |
| Thelyphonida | 2 |
| Schizomida | 3 |
| Araneae | 155 |

**Table S4.** Chain lengths for Bayesian inference analyses using PhyloBayes-mpi.

| ***chain*** | ***states*** |
|---|---|
| C1 | 20374 |
| C2 | 20181 |
| C3 | 20284 |
| C4 | 20393 |
| C5 | 20820 |
| C6 | 20669 |
| C7 | 20029 |
| C8 | 20244 |

**Table S5.** Summary statistics for Bayesian inference analyses using Phylobayes-mpi, based on all eight chains.

| ***parameter*** | ***mean*** | ***ESS*** |
|---|---|---|
| loglik | -3.101E5 | 206 |
| length | 11.873 | 985 |
| alpha | 0.689 | 162 |
| Nmode | 602.002 | 2794 |
| statent | 1.537 | 172 |
| statalpha | 5.822 | 543 |
| rrent | 4.223 | 405 |
| rrmean | 1.002 | 1.437E5 |

**Table S6.** Detection of paralogs in the 233-locus Matrix A of Lozano *et al.* (2019).

| ***Locus*** | ***Number of sequences with at least one hit*** | ***Number of paralogs detected*** | ***Mappings in D. melanogaster proteome*** |
|---|---|---|---|
| ar21 | 84 | 2 | +Drosophila_melanogasterlG022873lGNP_650498.3+Drosophila_melanogasterlG028454lGNP_732075.3:81 |
| arp23 | 82 | 1 | +Drosophila_melanogasterlG013712lGNP_476596.1+Drosophila_melanogasterlG024891lGNP_723845.1:82 |
| atpsynthalphaamt | 87 | 1 | +Drosophila_melanogasterlG026041lGNP_726243.1:87 |
| cctA | 88 | 2 | +Drosophila_melanogasterlG015590lGNP_524450.2+Drosophila_melanogasterlG028743lGNP_732748.1:87 |
| cctB | 87 | 2 | +Drosophila_melanogasterlG016727lGNP_572524.1:85 |
| cctD | 87 | 2 | +Drosophila_melanogasterlG018473lGNP_609579.1:86 |
| cctE | 87 | 3 | +Drosophila_melanogasterlG014897lGNP_523707.1+Drosophila_melanogasterlG025524lGNP_725107.1:80 |
| cctG | 88 | 1 | +Drosophila_melanogasterlG004565lGNP_001189236.1+Drosophila_melanogasterlG004566lGNP_001189237.1+Drosophila_melanogasterlG022932lGNP_650572.2+Drosophila_melanogasterlG028499lGNP_732167.1:88 |
| cctN | 87 | 1 | +Drosophila_melanogasterlG022361lGNP_649835.1:87 |
| cctZ | 87 | 1 | +Drosophila_melanogasterlG017140lGNP_573066.1:87 |
| cpn60mt | 83 | 3 | +Drosophila_melanogasterlG014526lGNP_511115.2+Drosophila_melanogasterlG015992lGNP_524925.1+Drosophila_melanogasterlG017955lGNP_608948.2+Drosophila_melanogasterlG024572lGNP_723104.2+Drosophila_melanogasterlG024573lGNP_723105.2+Drosophila_melanogasterlG026590lGNP_727489.1:65 |
| crfg | 84 | 1 | +Drosophila_melanogasterlG011931lGNP_001286247.1+Drosophila_melanogasterlG019222lGNP_610484.1:84 |
| ef1EF1 | 89 | 3 | +Drosophila_melanogasterlG012005lGNP_001286321.1+Drosophila_melanogasterlG012006lGNP_001286322.1+Drosophila_melanogasterlG014343lGNP_477375.1+Drosophila_melanogasterlG015723lGNP_524611.1+Drosophila_melanogasterlG025509lGNP_725085.1+Drosophila_melanogasterlG029180lGNP_733449.1+Drosophila_melanogasterlG030286lGNP_996315.1+Drosophila_melanogasterlG030287lGNP_996316.1:82 |
| ef1RF3 | 87 | 5 | +Drosophila_melanogasterlG007742lGNP_001260415.1+Drosophila_melanogasterlG007743lGNP_001260416.1+Drosophila_melanogasterlG011558lGNP_001285874.1+Drosophila_melanogasterlG014251lGNP_477259.1:67 |
| ef2EF2 | 88 | 2 | +Drosophila_melanogasterlG016124lGNP_525105.2+Drosophila_melanogasterlG023741lGNP_651605.1+Drosophila_melanogasterlG025154lGNP_724357.2+Drosophila_melanogasterlG025155lGNP_724358.2:71 |
| eif5a | 82 | 1 | +Drosophila_melanogasterlG020399lGNP_611878.1+Drosophila_melanogasterlG026124lGNP_726411.1:82 |

| | | | |
|---|---|---|---|
| fibri | 76 | 1 | +Drosophila_melanogasterlG015002lGNP_523817.1:76 |
| fpps | 74 | 1 | +Drosophila_melanogasterlG014348lGNP_477380.1:74 |
| g122 | 79 | 1 | +Drosophila_melanogasterlG014329lGNP_477356.1:79 |
| g125 | 20 | 1 | +Drosophila_melanogasterlG018135lGNP_609162.1:20 |
| g127 | 69 | 1 | +Drosophila_melanogasterlG020455lGNP_611940.1+Drosophila_melanogasterlG026143lGNP_726463.1:69 |
| g143 | 79 | 1 | +Drosophila_melanogasterlG023971lGNP_651885.1:79 |
| g46 | 77 | 1 | +Drosophila_melanogasterlG007218lGNP_001259877.1+Drosophila_melanogasterlG014046lGNP_477005.1:77 |
| g51 | 7 | 1 | +Drosophila_melanogasterlG004466lGNP_001189124.1+Drosophila_melanogasterlG004467lGNP_001189125.1+Drosophila_melanogasterlG009255lGNP_001261987.1+Drosophila_melanogasterlG021687lGNP_648985.1:7 |
| g53 | 10 | 1 | +Drosophila_melanogasterlG024158lGNP_652341.1:10 |
| g65 | 73 | 1 | +Drosophila_melanogasterlG008182lGNP_001260877.1+Drosophila_melanogasterlG019335lGNP_610629.1:73 |
| g7 | 72 | 1 | +Drosophila_melanogasterlG021580lGNP_648848.1:72 |
| g70 | 69 | 1 | +Drosophila_melanogasterlG020884lGNP_647961.1:69 |
| g78 | 81 | 1 | +Drosophila_melanogasterlG021751lGNP_649059.1:81 |
| g82 | 15 | 3 | +Drosophila_melanogasterlG010811lGNP_001285120.1+Drosophila_melanogasterlG016862lGNP_572713.1:5 |
| grc5 | 86 | 1 | +Drosophila_melanogasterlG009494lGNP_001262233.1+Drosophila_melanogasterlG024018lGNP_651954.3+Drosophila_melanogasterlG027922lGNP_730773.4:86 |
| hsp70E | 88 | 4 | +Drosophila_melanogasterlG009839lGNP_001262586.1+Drosophila_melanogasterlG010830lGNP_001285139.1+Drosophila_melanogasterlG012098lGNP_001286414.1+Drosophila_melanogasterlG014542lGNP_511132.2+Drosophila_melanogasterlG014928lGNP_523741.2+Drosophila_melanogasterlG015233lGNP_524063.1+Drosophila_melanogasterlG015486lGNP_524339.1+Drosophila_melanogasterlG015502lGNP_524356.1+Drosophila_melanogasterlG015611lGNP_524474.1+Drosophila_melanogasterlG015883lGNP_524798.2+Drosophila_melanogasterlG015993lGNP_524927.2+Drosophila_melanogasterlG022659lGNP_650209.1+Drosophila_melanogasterlG026626lGNP_727563.1+Drosophila_melanogasterlG026627lGNP_727564.1+Drosophila_melanogasterlG026628lGNP_727565.1+Drosophila_melanogasterlG027609lGNP_729940.1+Drosophila_melanogasterlG027610lGNP_729941.1+Drosophila_melanogasterlG028271lGNP_731651.1+Drosophila_melanogasterlG028296lGNP_731716.1+Drosophila_melanogasterlG028415lGNP_731987.1+Drosophila_melanogasterlG028416lGNP_731988.1+Drosophila_melanogasterlG028417lGNP_731989.1+Drosophila_melanogasterlG029548lGNP_788663.1+Drosophila_melanogasterlG029562lGNP_788679.1+Drosophila_melanogasterlG029563lGNP_788680.1:79 |

| | | | |
|---|---|---|---|
| hsp70mt | 88 | 10 | +Drosophila_melanogasterlG012098lGNP_001286414.1+Drosophila_melanogasterlG014928lGNP_523741.2:15 |
| hsp90C | 83 | 3 | +Drosophila_melanogasterlG008645lGNP_001261362.1+Drosophila_melanogasterlG015079lGNP_523899.1:81 |
| hsp90E | 87 | 3 | +Drosophila_melanogasterlG023737lGNP_651601.1:19 |
| if1a | 83 | 1 | +Drosophila_melanogasterlG013059lGNP_001287386.1+Drosophila_melanogasterlG015820lGNP_524728.2+Drosophila_melanogasterlG030219lGNP_996231.1:83 |
| if2b | 83 | 1 | +Drosophila_melanogasterlG015215lGNP_524043.1:83 |
| if2g | 84 | 2 | +Drosophila_melanogasterlG009840lGNP_001262587.1+Drosophila_melanogasterlG028418lGNP_731993.1+Drosophila_melanogasterlG028419lGNP_731994.1:70 |
| if4aa | 88 | 3 | +Drosophila_melanogasterlG005102lGNP_001245907.1+Drosophila_melanogasterlG007455lGNP_001260117.1+Drosophila_melanogasterlG013711lGNP_476595.1+Drosophila_melanogasterlG022324lGNP_649788.2+Drosophila_melanogasterlG024583lGNP_723137.1+Drosophila_melanogasterlG024584lGNP_723138.1+Drosophila_melanogasterlG024585lGNP_723139.1:72 |
| if4ab | 86 | 3 | +Drosophila_melanogasterlG005102lGNP_001245907.1+Drosophila_melanogasterlG007455lGNP_001260117.1+Drosophila_melanogasterlG013711lGNP_476595.1+Drosophila_melanogasterlG022324lGNP_649788.2+Drosophila_melanogasterlG024583lGNP_723137.1+Drosophila_melanogasterlG024584lGNP_723138.1+Drosophila_melanogasterlG024585lGNP_723139.1:79 |
| if6 | 79 | 1 | +Drosophila_melanogasterlG024321lGNP_659573.1:79 |
| l12eA | 84 | 1 | +Drosophila_melanogasterlG027584lGNP_729866.1:84 |
| l12eC | 79 | 1 | +Drosophila_melanogasterlG007624lGNP_001260293.1+Drosophila_melanogasterlG015810lGNP_524714.1:79 |
| l12eD | 84 | 1 | +Drosophila_melanogasterlG010637lGNP_001284944.1+Drosophila_melanogasterlG010638lGNP_001284945.1+Drosophila_melanogasterlG014478lGNP_511063.1+Drosophila_melanogasterlG026444lGNP_727094.1+Drosophila_melanogasterlG026445lGNP_727096.1:84 |
| mcmA | 77 | 9 | +Drosophila_melanogasterlG015455lGNP_524308.2:58 |
| mcmD | 80 | 7 | +Drosophila_melanogasterlG015162lGNP_523984.1:66 |
| nsf1C | 88 | 5 | +Drosophila_melanogasterlG011085lGNP_001285396.1+Drosophila_melanogasterlG017298lGNP_573258.1:82 |
| nsf1G | 90 | 3 | +Drosophila_melanogasterlG017549lGNP_608447.1+Drosophila_melanogasterlG023909lGNP_651811.1:78 |
| nsf1J | 91 | 6 | +Drosophila_melanogasterlG016547lGNP_572308.3:9 |
| nsf1K | 89 | 3 | +Drosophila_melanogasterlG015602lGNP_524464.1:83 |
| nsf1L | 90 | 4 | +Drosophila_melanogasterlG010799lGNP_001285108.1+Drosophila_melanogasterlG016839lGNP_572686.1+Drosophila_melanogasterlG022448lGNP_649938.2+Drosophila_melanogasterlG028180lGNP_731401.3:75 |

| | | | |
|---|---|---|---|
| nsf1M | 89 | 3 | +Drosophila_melanogaster\|G013165\|GNP_001287493.1+Drosophila_melanogaster\|G015606\|GNP_524469.2:84 |
| nsf2A | 90 | 5 | +Drosophila_melanogaster\|G001631\|GNP_001097249.1+Drosophila_melanogaster\|G001632\|GNP_001097250.1+Drosophila_melanogaster\|G011945\|GNP_001286261.1+Drosophila_melanogaster\|G014337\|GNP_477369.1:81 |
| orf2 | 82 | 1 | +Drosophila_melanogaster\|G017510\|GNP_608387.1:82 |
| ornamtransa | 82 | 1 | +Drosophila_melanogaster\|G021817\|GNP_649139.1:82 |
| pace5A | 82 | 1 | +Drosophila_melanogaster\|G008763\|GNP_001261482.1+Drosophila_melanogaster\|G020960\|GNP_648057.2:82 |
| psmaA | 85 | 2 | +Drosophila_melanogaster\|G014202\|GNP_477202.2+Drosophila_melanogaster\|G025806\|GNP_725669.1:83 |
| psmaB | 85 | 2 | +Drosophila_melanogaster\|G016112\|GNP_525092.1:83 |
| psmaC | 85 | 3 | +Drosophila_melanogaster\|G013789\|GNP_476691.1:80 |
| psmaD | 86 | 2 | +Drosophila_melanogaster\|G015475\|GNP_524328.1:83 |
| psmaE | 86 | 3 | +Drosophila_melanogaster\|G011484\|GNP_001285798.1+Drosophila_melanogaster\|G014738\|GNP_523532.1:81 |
| psmaF | 84 | 2 | +Drosophila_melanogaster\|G025407\|GNP_724834.1:83 |
| psmaG | 87 | 1 | +Drosophila_melanogaster\|G003238\|GNP_001163078.1+Drosophila_melanogaster\|G015916\|GNP_524837.2+Drosophila_melanogaster\|G025299\|GNP_724614.1+Drosophila_melanogaster\|G025301\|GNP_724616.1:87 |
| psmbH | 82 | 1 | +Drosophila_melanogaster\|G007836\|GNP_001260511.1+Drosophila_melanogaster\|G018665\|GNP_609804.1:82 |
| psmbI | 83 | 2 | +Drosophila_melanogaster\|G022377\|GNP_649858.1:82 |
| psmbJ | 85 | 1 | +Drosophila_melanogaster\|G015280\|GNP_524115.1:85 |
| psmbK | 84 | 3 | +Drosophila_melanogaster\|G016509\|GNP_572267.1:10 |
| psmbL | 83 | 1 | +Drosophila_melanogaster\|G024076\|GNP_652031.2:83 |
| psmbM | 84 | 1 | +Drosophila_melanogaster\|G024061\|GNP_652014.1+Drosophila_melanogaster\|G025477\|GNP_724974.1:84 |
| psmbN | 80 | 1 | +Drosophila_melanogaster\|G022121\|GNP_649529.1+Drosophila_melanogaster\|G027990\|GNP_730922.1:80 |
| pyrdehydroe1bmt | 84 | 2 | +Drosophila_melanogaster\|G023790\|GNP_651668.1+Drosophila_melanogaster\|G029027\|GNP_733265.1:78 |
| rad23 | 79 | 2 | +Drosophila_melanogaster\|G026196\|GNP_726561.1:75 |
| rf1 | 83 | 4 | +Drosophila_melanogaster\|G009368\|GNP_001262104.1+Drosophila_melanogaster\|G009369\|GNP_001262105.1+Drosophila_melanogaster\|G021874\|GNP_649210.1+Drosophila_melanogaster\|G027810\|GNP_730517.1+Drosophila_melanogaster\|G027811\|GNP_730518.1+Drosophila_melanogaster\|G027812\|GNP_730519.1+Drosophila_melanogaster\|G027813\|GNP_730520.1+Drosophila_melanogaster\|G029459\|GNP_788547.1:71 |

| | | | |
|---|---|---|---|
| rla2A | 80 | 1 | +Drosophila_melanogasterIG014951IGNP_523764.1:80 |
| rla2B | 85 | 1 | +Drosophila_melanogasterIG007165IGNP_001259822.1+Drosophila_melanogasterIG013745IGNP_476630.1:85 |
| rpl1 | 88 | 1 | +Drosophila_melanogasterIG021309IGNP_648514.1:88 |
| rpl11b | 85 | 1 | +Drosophila_melanogasterIG012294IGNP_001286614.1+Drosophila_melanogasterIG014089IGNP_477054.1:85 |
| rpl12b | 86 | 1 | +Drosophila_melanogasterIG015898IGNP_524819.1+Drosophila_melanogasterIG026125IGNP_726412.1+Drosophila_melanogasterIG026126IGNP_726413.1:86 |
| rpl13 | 82 | 1 | +Drosophila_melanogasterIG000791IGNP_001033891.1+Drosophila_melanogasterIG007643IGNP_001260312.1+Drosophila_melanogasterIG014736IGNP_523530.1:82 |
| rpl14a | 81 | 1 | +Drosophila_melanogasterIG015153IGNP_523975.1:81 |
| rpl15a | 82 | 1 | +Drosophila_melanogasterIG000295IGNP_001015155.1+Drosophila_melanogasterIG000296IGNP_001015156.1+Drosophila_melanogasterIG002267IGNP_001104429.2+Drosophila_melanogasterIG002268IGNP_001104430.1+Drosophila_melanogasterIG004644IGNP_001245395.1+Drosophila_melanogasterIG004645IGNP_001245396.1+Drosophila_melanogasterIG004646IGNP_001245397.1+Drosophila_melanogasterIG004647IGNP_001245398.1:82 |
| rpl16b | 82 | 2 | +Drosophila_melanogasterIG022145IGNP_649560.1:80 |
| rpl17 | 85 | 1 | +Drosophila_melanogasterIG010654IGNP_001284961.1+Drosophila_melanogasterIG016578IGNP_572346.1+Drosophila_melanogasterIG026454IGNP_727118.1+Drosophila_melanogasterIG026455IGNP_727119.1+Drosophila_melanogasterIG026456IGNP_727120.1:85 |
| rpl18 | 85 | 1 | +Drosophila_melanogasterIG008782IGNP_001261501.1+Drosophila_melanogasterIG012644IGNP_001286966.1+Drosophila_melanogasterIG020985IGNP_648091.1:85 |
| rpl19a | 84 | 1 | +Drosophila_melanogasterIG013746IGNP_476631.1+Drosophila_melanogasterIG029972IGNP_995941.1:84 |
| rpl2 | 84 | 1 | +Drosophila_melanogasterIG015819IGNP_524726.1+Drosophila_melanogasterIG027101IGNP_728756.1:84 |
| rpl20 | 85 | 1 | +Drosophila_melanogasterIG014960IGNP_523774.1:85 |
| rpl21 | 80 | 1 | +Drosophila_melanogasterIG008006IGNP_001260689.1+Drosophila_melanogasterIG011817IGNP_001286133.1+Drosophila_melanogasterIG018941IGNP_610144.1:80 |
| rpl22 | 74 | 2 | +Drosophila_melanogasterIG014147IGNP_477134.1:70 |
| rpl23a | 80 | 1 | +Drosophila_melanogasterIG014998IGNP_523813.1:80 |
| rpl24A | 84 | 1 | +Drosophila_melanogasterIG011579IGNP_001285895.1+Drosophila_melanogasterIG011580IGNP_001285896.1+Drosophila_melanogasterIG018540IGNP_609649.1:84 |
| rpl24B | 85 | 2 | +Drosophila_melanogasterIG022556IGNP_650073.1:80 |
| rpl25 | 81 | 1 | +Drosophila_melanogasterIG015066IGNP_523886.1:81 |

| | | | |
|---|---|---|---|
| rpl26 | 80 | 1 | +Drosophila_melanogaster\|G009293\|GNP_001262025.1+Drosophila_melanogaster\|G013388\|GNP_001303372.1+Drosophila_melanogaster\|G021760\|GNP_649070.1:80 |
| rpl27 | 80 | 1 | +Drosophila_melanogaster\|G004071\|GNP_001188689.2+Drosophila_melanogaster\|G011288\|GNP_001285601.1+Drosophila_melanogaster\|G014009\|GNP_476963.1:80 |
| rpl3 | 88 | 1 | +Drosophila_melanogaster\|G015463\|GNP_524316.1+Drosophila_melanogaster\|G028229\|GNP_731548.2:88 |
| rpl30 | 78 | 1 | +Drosophila_melanogaster\|G007871\|GNP_001260550.2+Drosophila_melanogaster\|G015787\|GNP_524687.1+Drosophila_melanogaster\|G025054\|GNP_724149.1:78 |
| rpl31 | 83 | 1 | +Drosophila_melanogaster\|G019233\|GNP_610503.1+Drosophila_melanogaster\|G025392\|GNP_724804.1+Drosophila_melanogaster\|G025393\|GNP_724805.1:83 |
| rpl32 | 80 | 3 | +Drosophila_melanogaster\|G029085\|GNP_733339.1:23 |
| rpl33a | 76 | 1 | +Drosophila_melanogaster\|G022130\|GNP_649539.1:76 |
| rpl34 | 74 | 2 | +Drosophila_melanogaster\|G012924\|GNP_001287249.1+Drosophila_melanogaster\|G012925\|GNP_001287250.1+Drosophila_melanogaster\|G022403\|GNP_649887.1+Drosophila_melanogaster\|G028158\|GNP_731345.1:67 |
| rpl35 | 74 | 1 | +Drosophila_melanogaster\|G010593\|GNP_001284900.1+Drosophila_melanogaster\|G016491\|GNP_572243.1+Drosophila_melanogaster\|G026428\|GNP_727016.1:74 |
| rpl36 | 72 | 1 | +Drosophila_melanogaster\|G010443\|GNP_001284750.1+Drosophila_melanogaster\|G013744\|GNP_476629.1+Drosophila_melanogaster\|G026269\|GNP_726686.1+Drosophila_melanogaster\|G026270\|GNP_726687.1+Drosophila_melanogaster\|G026271\|GNP_726688.1:72 |
| rpl37a | 57 | 1 | +Drosophila_melanogaster\|G006918\|GNP_001259561.1+Drosophila_melanogaster\|G017091\|GNP_573005.1:57 |
| rpl42 | 73 | 1 | +Drosophila_melanogaster\|G011423\|GNP_001285737.1+Drosophila_melanogaster\|G018149\|GNP_609179.2:73 |
| rpl43b | 17 | 1 | +Drosophila_melanogaster\|G004073\|GNP_001188691.1+Drosophila_melanogaster\|G015868\|GNP_524781.1+Drosophila_melanogaster\|G024557\|GNP_723060.1:17 |
| rpl4B | 86 | 1 | +Drosophila_melanogaster\|G015671\|GNP_524538.2:86 |
| rpl6 | 80 | 1 | +Drosophila_melanogaster\|G029168\|GNP_733433.1:80 |
| rpl7A | 83 | 1 | +Drosophila_melanogaster\|G011480\|GNP_001285794.1+Drosophila_melanogaster\|G014737\|GNP_523531.1:83 |
| rpl9 | 84 | 1 | +Drosophila_melanogaster\|G011510\|GNP_001285826.1+Drosophila_melanogaster\|G014169\|GNP_477161.1+Drosophila_melanogaster\|G024814\|GNP_723644.1:84 |
| rpp0 | 85 | 1 | +Drosophila_melanogaster\|G009463\|GNP_001262202.1+Drosophila_melanogaster\|G015364\|GNP_524211.1:85 |
| rps1 | 88 | 2 | +Drosophila_melanogaster\|G004652\|GNP_001245404.1+Drosophila_melanogaster\|G015730\|GNP_524618.1:85 |

| | | | |
|---|---|---|---|
| rps10 | 82 | 1 | +Drosophila_melanogasterlG004973lGNP_001245758.1+Drosophila_melanogasterlG011134lGNP_001285445.1+Drosophila_melanogasterlG017453lGNP_608324.1+Drosophila_melanogasterlG026905lGNP_728273.1:82 |
| rps11 | 84 | 1 | +Drosophila_melanogasterlG025525lGNP_725114.1:84 |
| rps13a | 82 | 1 | +Drosophila_melanogasterlG000790lGNP_001033885.1+Drosophila_melanogasterlG011433lGNP_001285747.1+Drosophila_melanogasterlG013986lGNP_476938.1:82 |
| rps14 | 87 | 1 | +Drosophila_melanogasterlG010687lGNP_001284995.1+Drosophila_melanogasterlG015957lGNP_524884.1+Drosophila_melanogasterlG016143lGNP_536352.1+Drosophila_melanogasterlG026490lGNP_727218.1:87 |
| rps15 | 83 | 2 | +Drosophila_melanogasterlG025778lGNP_725591.1:29 |
| rps16 | 83 | 1 | +Drosophila_melanogasterlG012411lGNP_001286731.1+Drosophila_melanogasterlG020236lGNP_611685.1:83 |
| rps17 | 81 | 1 | +Drosophila_melanogasterlG015178lGNP_524002.1:81 |
| rps18 | 82 | 1 | +Drosophila_melanogasterlG014010lGNP_476964.1+Drosophila_melanogasterlG025922lGNP_725943.1:82 |
| rps19 | 82 | 1 | +Drosophila_melanogasterlG011028lGNP_001285338.1+Drosophila_melanogasterlG011029lGNP_001285339.1+Drosophila_melanogasterlG014600lGNP_523376.1+Drosophila_melanogasterlG026800lGNP_727992.1+Drosophila_melanogasterlG026801lGNP_727993.1:82 |
| rps2 | 86 | 1 | +Drosophila_melanogasterlG007634lGNP_001260303.1+Drosophila_melanogasterlG011471lGNP_001285785.1+Drosophila_melanogasterlG013939lGNP_476874.1:86 |
| rps20 | 77 | 1 | +Drosophila_melanogasterlG015562lGNP_524421.1:77 |
| rps22a | 83 | 1 | +Drosophila_melanogasterlG002938lGNP_001162738.1+Drosophila_melanogasterlG015806lGNP_524709.1+Drosophila_melanogasterlG026672lGNP_727690.1+Drosophila_melanogasterlG026673lGNP_727692.1+Drosophila_melanogasterlG026674lGNP_727693.1:83 |
| rps23 | 85 | 1 | +Drosophila_melanogasterlG012094lGNP_001286410.1+Drosophila_melanogasterlG019604lGNP_610939.2:85 |
| rps24 | 81 | 1 | +Drosophila_melanogasterlG020242lGNP_611693.1:81 |
| rps25 | 79 | 1 | +Drosophila_melanogasterlG015462lGNP_524315.2+Drosophila_melanogasterlG028227lGNP_731544.1:79 |
| rps26 | 83 | 1 | +Drosophila_melanogasterlG007859lGNP_001260537.1+Drosophila_melanogasterlG011726lGNP_001286042.1+Drosophila_melanogasterlG014796lGNP_523595.1+Drosophila_melanogasterlG025038lGNP_724109.1+Drosophila_melanogasterlG025039lGNP_724110.1:83 |
| rps27 | 19 | 1 | +Drosophila_melanogasterlG006339lGNP_001247301.1+Drosophila_melanogasterlG023548lGNP_651359.1:19 |
| rps27a | 88 | 2 | +Drosophila_melanogasterlG007358lGNP_001260018.1+Drosophila_melanogasterlG013862lGNP_476776.1:28 |

| | | | |
|---|---|---|---|
| rps3 | 85 | 1 | +Drosophila_melanogaster\|G013153\|GNP_001287481.1+Drosophila_melanogaster\|G013747\|GNP_476632.1:85 |
| rps4 | 88 | 1 | +Drosophila_melanogaster\|G012732\|GNP_001287055.1+Drosophila_melanogaster\|G015223\|GNP_524053.2+Drosophila_melanogaster\|G027585\|GNP_729871.1:88 |
| rps5 | 86 | 1 | +Drosophila_melanogaster\|G011049\|GNP_001285360.1+Drosophila_melanogaster\|G011050\|GNP_001285361.1+Drosophila_melanogaster\|G011051\|GNP_001285362.1+Drosophila_melanogaster\|G014605\|GNP_523382.1:86 |
| rps6 | 83 | 2 | +Drosophila_melanogaster\|G014487\|GNP_511073.1:76 |
| rps7 | 85 | 1 | +Drosophila_melanogaster\|G023887\|GNP_651782.1+Drosophila_melanogaster\|G029096\|GNP_733355.1+Drosophila_melanogaster\|G029097\|GNP_733356.2+Drosophila_melanogaster\|G030283\|GNP_996312.1:85 |
| rps8 | 86 | 1 | +Drosophila_melanogaster\|G006396\|GNP_001247362.1+Drosophila_melanogaster\|G013266\|GNP_001287596.1+Drosophila_melanogaster\|G013267\|GNP_001287597.1+Drosophila_melanogaster\|G023853\|GNP_651740.1:86 |
| rps9 | 83 | 2 | +Drosophila_melanogaster\|G012686\|GNP_001287008.1+Drosophila_melanogaster\|G015180\|GNP_524004.2+Drosophila_melanogaster\|G027431\|GNP_729506.1:78 |
| sadhchydrolaseE1 | 86 | 4 | +Drosophila_melanogaster\|G006927\|GNP_001259573.1+Drosophila_melanogaster\|G010959\|GNP_001285268.1+Drosophila_melanogaster\|G014572\|GNP_511164.2:70 |
| sap40 | 83 | 1 | +Drosophila_melanogaster\|G010478\|GNP_001284785.1+Drosophila_melanogaster\|G013837\|GNP_476750.1+Drosophila_melanogaster\|G026303\|GNP_726744.1+Drosophila_melanogaster\|G026304\|GNP_726745.3:83 |
| srp54 | 83 | 2 | +Drosophila_melanogaster\|G015110\|GNP_523931.1:78 |
| srs | 82 | 1 | +Drosophila_melanogaster\|G007323\|GNP_001259982.1+Drosophila_melanogaster\|G017783\|GNP_608743.2:82 |
| stbproptase2ab | 85 | 2 | +Drosophila_melanogaster\|G011411\|GNP_001285724.1+Drosophila_melanogaster\|G011412\|GNP_001285725.1+Drosophila_melanogaster\|G013884\|GNP_476805.1:78 |
| stcproptase2ac | 85 | 3 | +Drosophila_melanogaster\|G006647\|GNP_001259272.1+Drosophila_melanogaster\|G006648\|GNP_001259273.1+Drosophila_melanogaster\|G006649\|GNP_001259274.1+Drosophila_melanogaster\|G014476\|GNP_511061.1:57 |
| suca | 86 | 3 | +Drosophila_melanogaster\|G004391\|GNP_001189044.1+Drosophila_melanogaster\|G015085\|GNP_523905.2:80 |
| tif2a | 81 | 1 | +Drosophila_melanogaster\|G011019\|GNP_001285329.1+Drosophila_melanogaster\|G017192\|GNP_573130.1:81 |
| tribe1006 | 67 | 1 | +Drosophila_melanogaster\|G023403\|GNP_651181.1:67 |
| tribe1007 | 77 | 1 | +Drosophila_melanogaster\|G007218\|GNP_001259877.1+Drosophila_melanogaster\|G014046\|GNP_477005.1:77 |
| tribe1009 | 82 | 2 | +Drosophila_melanogaster\|G022273\|GNP_649725.2:49 |
| tribe1015 | 79 | 2 | +Drosophila_melanogaster\|G023176\|GNP_650887.1:48 |

| | | | |
|---|---|---|---|
| tribe1026 | 82 | 1 | +Drosophila_melanogaster\|G023164\|GNP_650864.1:82 |
| tribe1046 | 46 | 1 | +Drosophila_melanogaster\|G009727\|GNP_001262472.1+Drosophila_melanogaster\|G022532\|GNP_650047.3:46 |
| tribe1047 | 77 | 2 | +Drosophila_melanogaster\|G027357\|GNP_729358.1:74 |
| tribe1050 | 76 | 1 | +Drosophila_melanogaster\|G010636\|GNP_001284943.1+Drosophila_melanogaster\|G013574\|GNP_001303559.1+Drosophila_melanogaster\|G016555\|GNP_572318.1:76 |
| tribe1054 | 79 | 1 | +Drosophila_melanogaster\|G022119\|GNP_649527.1:79 |
| tribe1081 | 81 | 1 | +Drosophila_melanogaster\|G009380\|GNP_001262116.1+Drosophila_melanogaster\|G021894\|GNP_649234.1:81 |
| tribe1099 | 71 | 1 | +Drosophila_melanogaster\|G014015\|GNP_476969.1:71 |
| tribe1118 | 80 | 1 | +Drosophila_melanogaster\|G018369\|GNP_609444.1:80 |
| tribe1121 | 14 | 1 | +Drosophila_melanogaster\|G024118\|GNP_652189.1:14 |
| tribe1132 | 80 | 1 | +Drosophila_melanogaster\|G003428\|GNP_001163296.1+Drosophila_melanogaster\|G020469\|GNP_611954.2+Drosophila_melanogaster\|G026145\|GNP_726470.1+Drosophila_melanogaster\|G026146\|GNP_726471.1:80 |
| tribe1136 | 73 | 1 | +Drosophila_melanogaster\|G008182\|GNP_001260877.1+Drosophila_melanogaster\|G019335\|GNP_610629.1:73 |
| tribe1165 | 73 | 1 | +Drosophila_melanogaster\|G021227\|GNP_648409.1:73 |
| tribe1169 | 81 | 2 | +Drosophila_melanogaster\|G005655\|GNP_001246538.1:80 |
| tribe1170 | 78 | 1 | +Drosophila_melanogaster\|G011573\|GNP_001285889.1+Drosophila_melanogaster\|G018533\|GNP_609642.1:78 |
| tribe1181 | 79 | 1 | +Drosophila_melanogaster\|G013404\|GNP_001303388.1+Drosophila_melanogaster\|G022038\|GNP_649424.1:79 |
| tribe1190 | 77 | 1 | +Drosophila_melanogaster\|G015557\|GNP_524416.1:77 |
| tribe1200 | 75 | 1 | +Drosophila_melanogaster\|G015351\|GNP_524197.1:75 |
| tribe1204 | 77 | 1 | +Drosophila_melanogaster\|G003783\|GNP_001163688.1+Drosophila_melanogaster\|G023303\|GNP_651053.1:77 |
| tribe1244 | 79 | 1 | +Drosophila_melanogaster\|G017650\|GNP_608575.1:79 |
| tribe1245 | 81 | 2 | +Drosophila_melanogaster\|G006072\|GNP_001247010.1+Drosophila_melanogaster\|G022438\|GNP_649925.1:80 |
| tribe1294 | 79 | 1 | +Drosophila_melanogaster\|G022551\|GNP_650067.1:79 |
| tribe1335 | 78 | 1 | +Drosophila_melanogaster\|G006320\|GNP_001247279.1+Drosophila_melanogaster\|G023451\|GNP_651234.1:78 |
| tribe1378 | 75 | 1 | +Drosophila_melanogaster\|G019718\|GNP_611078.1:75 |
| tribe1381A | 80 | 1 | +Drosophila_melanogaster\|G014381\|GNP_477419.1:80 |
| tribe1409 | 82 | 1 | +Drosophila_melanogaster\|G009017\|GNP_001261742.1+Drosophila_melanogaster\|G024087\|GNP_652042.1:82 |
| tribe281 | 77 | 1 | +Drosophila_melanogaster\|G009743\|GNP_001262488.1+Drosophila_melanogaster\|G013501\|GNP_001 |

| | | | |
|---|---|---|---|
| | | | 303486.1+Drosophila_melanogaster\|G014217\|GNP_477217.1+Drosophila_melanogaster\|G028266\|GNP_731641.1:77 |
| tribe320 | 86 | 1 | +Drosophila_melanogaster\|G013106\|GNP_001287433.1+Drosophila_melanogaster\|G023202\|GNP_650922.1+Drosophila_melanogaster\|G028666\|GNP_732566.1:86 |
| tribe333 | 81 | 1 | +Drosophila_melanogaster\|G011668\|GNP_001285984.1+Drosophila_melanogaster\|G014183\|GNP_477176.1:81 |
| tribe340 | 14 | 2 | +Drosophila_melanogaster\|G007311\|GNP_001259970.1+Drosophila_melanogaster\|G024469\|GNP_722853.1+Drosophila_melanogaster\|G024470\|GNP_722854.1+Drosophila_melanogaster\|G024471\|GNP_722855.1:12 |
| tribe351 | 29 | 1 | +Drosophila_melanogaster\|G003466\|GNP_001163336.1+Drosophila_melanogaster\|G003467\|GNP_001163337.1+Drosophila_melanogaster\|G020751\|GNP_647791.2+Drosophila_melanogaster\|G027146\|GNP_728839.1+Drosophila_melanogaster\|G027147\|GNP_728840.1+Drosophila_melanogaster\|G027148\|GNP_728841.2:29 |
| tribe368 | 23 | 1 | +Drosophila_melanogaster\|G030477\|GYP_009047273.1:23 |
| tribe369 | 22 | 1 | +Drosophila_melanogaster\|G030474\|GYP_009047270.1:22 |
| tribe460 | 81 | 1 | +Drosophila_melanogaster\|G013446\|GNP_001303431.1+Drosophila_melanogaster\|G022533\|GNP_650048.1:81 |
| tribe495 | 82 | 1 | +Drosophila_melanogaster\|G015683\|GNP_524550.1+Drosophila_melanogaster\|G029059\|GNP_733304.1+Drosophila_melanogaster\|G029060\|GNP_733305.1:82 |
| tribe508 | 81 | 1 | +Drosophila_melanogaster\|G029626\|GNP_788765.1+Drosophila_melanogaster\|G029627\|GNP_788766.1:81 |
| tribe532 | 83 | 1 | +Drosophila_melanogaster\|G015504\|GNP_524358.2:83 |
| tribe542 | 79 | 1 | +Drosophila_melanogaster\|G020752\|GNP_647792.1:79 |
| tribe550 | 81 | 3 | +Drosophila_melanogaster\|G008021\|GNP_001260708.1:5 |
| tribe572 | 81 | 1 | +Drosophila_melanogaster\|G007217\|GNP_001259876.1+Drosophila_melanogaster\|G015839\|GNP_524748.2+Drosophila_melanogaster\|G024401\|GNP_722715.1:81 |
| tribe585A | 85 | 5 | +Drosophila_melanogaster\|G014677\|GNP_523463.2:80 |
| tribe586 | 84 | 1 | +Drosophila_melanogaster\|G015268\|GNP_524103.2:84 |
| tribe593 | 85 | 1 | +Drosophila_melanogaster\|G006762\|GNP_001259397.1+Drosophila_melanogaster\|G010765\|GNP_001285074.1+Drosophila_melanogaster\|G016785\|GNP_572610.1:85 |
| tribe613 | 14 | 1 | +Drosophila_melanogaster\|G023745\|GNP_651614.1:14 |
| tribe619 | 16 | 1 | +Drosophila_melanogaster\|G024082\|GNP_652037.1:16 |
| tribe622 | 81 | 1 | +Drosophila_melanogaster\|G014020\|GNP_476975.1+Drosophila_melanogaster\|G025507\|GNP_725073.1:81 |
| tribe629A | 80 | 1 | +Drosophila_melanogaster\|G025523\|GNP_725106.1:80 |
| tribe632 | 82 | 1 | +Drosophila_melanogaster\|G000743\|GNP_001033818.1+Drosophila_melanogaster\|G010448\|GNP_001284755.1+Drosophila_melanogaster\|G014357\|GNP |

| | | | |
|---|---|---|---|
| | | | _477390.1+Drosophila_melanogasterlG026272lGNP_726690.1+Drosophila_melanogasterlG026273lGNP_726691.1+Drosophila_melanogasterlG026274lGNP_726692.1+Drosophila_melanogasterlG026275lGNP_726693.1+Drosophila_melanogasterlG026276lGNP_726694.1+Drosophila_melanogasterlG026277lGNP_726695.1:82 |
| tribe646 | 74 | 1 | +Drosophila_melanogasterlG011300lGNP_001285613.1+Drosophila_melanogasterlG011301lGNP_001285614.1+Drosophila_melanogasterlG017921lGNP_608909.1:74 |
| tribe647 | 75 | 1 | +Drosophila_melanogasterlG003594lGNP_001163480.1+Drosophila_melanogasterlG003595lGNP_001163481.1+Drosophila_melanogasterlG009366lGNP_001262101.1+Drosophila_melanogasterlG015334lGNP_524178.2:75 |
| tribe683 | 27 | 1 | +Drosophila_melanogasterlG007791lGNP_001260465.1+Drosophila_melanogasterlG018615lGNP_609747.1:27 |
| tribe686 | 58 | 2 | +Drosophila_melanogasterlG025733lGNP_725504.1:24 |
| tribe700 | 79 | 1 | +Drosophila_melanogasterlG005875lGNP_001246795.1+Drosophila_melanogasterlG005876lGNP_001246796.1+Drosophila_melanogasterlG015275lGNP_524110.1+Drosophila_melanogasterlG027677lGNP_730178.1:79 |
| tribe716 | 78 | 1 | +Drosophila_melanogasterlG015186lGNP_524011.1:78 |
| tribe717 | 78 | 1 | +Drosophila_melanogasterlG021397lGNP_648622.1:78 |
| tribe726 | 11 | 2 | +Drosophila_melanogasterlG004981lGNP_001245767.1+Drosophila_melanogasterlG026911lGNP_728295.1+Drosophila_melanogasterlG026912lGNP_728296.1:10 |
| tribe739 | 80 | 1 | +Drosophila_melanogasterlG011365lGNP_001285678.1+Drosophila_melanogasterlG018038lGNP_609046.1:80 |
| tribe740 | 56 | 1 | +Drosophila_melanogasterlG018191lGNP_609222.1:56 |
| tribe756 | 76 | 2 | +Drosophila_melanogasterlG012731lGNP_001287054.1+Drosophila_melanogasterlG021399lGNP_648624.1:73 |
| tribe764 | 67 | 1 | +Drosophila_melanogasterlG011999lGNP_001286315.1+Drosophila_melanogasterlG019380lGNP_610679.1:67 |
| tribe765 | 20 | 1 | +Drosophila_melanogasterlG018135lGNP_609162.1:20 |
| tribe768 | 82 | 1 | +Drosophila_melanogasterlG022365lGNP_649840.1:82 |
| tribe801mt | 78 | 1 | +Drosophila_melanogasterlG011003lGNP_001285313.1+Drosophila_melanogasterlG017164lGNP_573097.1+Drosophila_melanogasterlG026754lGNP_727921.1:78 |
| tribe831 | 86 | 1 | +Drosophila_melanogasterlG010749lGNP_001285058.1+Drosophila_melanogasterlG016754lGNP_572567.1:86 |
| tribe832 | 84 | 1 | +Drosophila_melanogasterlG013966lGNP_476905.1+Drosophila_melanogasterlG029944lGNP_995904.1:84 |
| tribe850 | 79 | 1 | +Drosophila_melanogasterlG000115lGNP_001014579.1+Drosophila_melanogasterlG000116lGNP_001014580.1+Drosophila_melanogasterlG015168lGNP_523990.1:79 |
| tribe852 | 78 | 1 | +Drosophila_melanogasterlG017397lGNP_573385.1:78 |

| | | | |
|---|---|---|---|
| tribe858 | 81 | 2 | +Drosophila_melanogaster\|G014044\|GNP_477003.1:76 |
| tribe893 | 78 | 1 | +Drosophila_melanogaster\|G022210\|GNP_649645.1:78 |
| tribe895 | 82 | 1 | +Drosophila_melanogaster\|G020168\|GNP_611604.1:82 |
| tribe896 | 86 | 1 | +Drosophila_melanogaster\|G020738\|GNP_647775.1:86 |
| tribe905 | 65 | 1 | +Drosophila_melanogaster\|G000992\|GNP_001036353.1+Drosophila_melanogaster\|G000993\|GNP_001036354.1+Drosophila_melanogaster\|G024080\|GNP_652035.1:65 |
| tribe906 | 74 | 2 | +Drosophila_melanogaster\|G009351\|GNP_001262085.1+Drosophila_melanogaster\|G021841\|GNP_649169.3:72 |
| tribe927 | 79 | 1 | +Drosophila_melanogaster\|G023971\|GNP_651885.1:79 |
| tribe930 | 80 | 1 | +Drosophila_melanogaster\|G014980\|GNP_523794.1+Drosophila_melanogaster\|G025907\|GNP_725894.1:80 |
| tribe942 | 81 | 1 | +Drosophila_melanogaster\|G011067\|GNP_001285378.1+Drosophila_melanogaster\|G017273\|GNP_573228.1:81 |
| u2snrnp | 81 | 1 | +Drosophila_melanogaster\|G007157\|GNP_001259814.1+Drosophila_melanogaster\|G014208\|GNP_477208.1:81 |
| vacaatpasepl21a | 78 | 1 | +Drosophila_melanogaster\|G006166\|GNP_001247111.1+Drosophila_melanogaster\|G024057\|GNP_652010.1:78 |
| vata | 82 | 5 | +Drosophila_melanogaster\|G005196\|GNP_001246015.1+Drosophila_melanogaster\|G007751\|GNP_001260424.1+Drosophila_melanogaster\|G007752\|GNP_001260425.1+Drosophila_melanogaster\|G007753\|GNP_001260426.1+Drosophila_melanogaster\|G014762\|GNP_523560.2+Drosophila_melanogaster\|G018489\|GNP_609595.1+Drosophila_melanogaster\|G024055\|GNP_652004.2+Drosophila_melanogaster\|G024862\|GNP_723775.1+Drosophila_melanogaster\|G024863\|GNP_723776.1:69 |
| vatb | 85 | 2 | +Drosophila_melanogaster\|G003698\|GNP_001163597.1+Drosophila_melanogaster\|G013969\|GNP_476908.1+Drosophila_melanogaster\|G028297\|GNP_731726.1:84 |
| vatc | 82 | 2 | +Drosophila_melanogaster\|G014257\|GNP_477266.1+Drosophila_melanogaster\|G017436\|GNP_599140.1:80 |
| vate | 81 | 1 | +Drosophila_melanogaster\|G012858\|GNP_001287182.1+Drosophila_melanogaster\|G015389\|GNP_524237.1+Drosophila_melanogaster\|G028010\|GNP_730957.1:81 |
| vatpased | 82 | 1 | +Drosophila_melanogaster\|G024045\|GNP_651987.1:82 |
| vdac2 | 85 | 1 | +Drosophila_melanogaster\|G000798\|GNP_001033899.1+Drosophila_melanogaster\|G005153\|GNP_001245961.1+Drosophila_melanogaster\|G007695\|GNP_001260365.1+Drosophila_melanogaster\|G013890\|GNP_476813.1+Drosophila_melanogaster\|G017413\|GNP_599110.1:85 |
| w09c | 83 | 1 | +Drosophila_melanogaster\|G007677\|GNP_001260347.1+Drosophila_melanogaster\|G014347\|GNP_477379.1:83 |

**Table S7.** Detection of paralogs in the 200 slowest-evolving gene matrix of Howard *et al.* (2020).

<table>
<tr><th>Locus</th><th>Number of sequences with at least one hit</th><th>Number of paralogs detected</th><th>Mappings in D. melanogaster proteome</th></tr>
<tr><td>OG10224</td><td>83</td><td>3</td><td>+Drosophila_melanogaster|G005983|GNP_001246916.1+Drosophila_melanogaster|G015382|GNP_524230.2:78</td></tr>
<tr><td>OG1026</td><td>77</td><td>1</td><td>+Drosophila_melanogaster|G009653|GNP_001262397.1+Drosophila_melanogaster|G028140|GNP_731308.1:77</td></tr>
<tr><td>OG10391</td><td>70</td><td>1</td><td>+Drosophila_melanogaster|G028412|GNP_731983.1:70</td></tr>
<tr><td>OG10592</td><td>83</td><td>3</td><td>+Drosophila_melanogaster|G005344|GNP_001246192.1+Drosophila_melanogaster|G015947|GNP_524874.2:67</td></tr>
<tr><td>OG10605</td><td>78</td><td>1</td><td>+Drosophila_melanogaster|G000743|GNP_001033818.1+Drosophila_melanogaster|G010448|GNP_001284755.1+Drosophila_melanogaster|G014357|GNP_477390.1+Drosophila_melanogaster|G026272|GNP_726690.1+Drosophila_melanogaster|G026273|GNP_726691.1+Drosophila_melanogaster|G026274|GNP_726692.1+Drosophila_melanogaster|G026275|GNP_726693.1+Drosophila_melanogaster|G026276|GNP_726694.1+Drosophila_melanogaster|G026277|GNP_726695.1:78</td></tr>
<tr><td>OG10726</td><td>75</td><td>1</td><td>+Drosophila_melanogaster|G011003|GNP_001285313.1+Drosophila_melanogaster|G017164|GNP_573097.1+Drosophila_melanogaster|G026754|GNP_727921.1:75</td></tr>
<tr><td>OG10928</td><td>74</td><td>1</td><td>+Drosophila_melanogaster|G017999|GNP_608996.1:74</td></tr>
<tr><td>OG11399</td><td>73</td><td>1</td><td>+Drosophila_melanogaster|G015866|GNP_524779.1:73</td></tr>
<tr><td>OG1161</td><td>83</td><td>3</td><td>+Drosophila_melanogaster|G003840|GNP_001163753.1+Drosophila_melanogaster|G012515|GNP_001286835.1+Drosophila_melanogaster|G014981|GNP_523795.2+Drosophila_melanogaster|G015025|GNP_523842.2+Drosophila_melanogaster|G015438|GNP_524290.2+Drosophila_melanogaster|G023742|GNP_651606.2+Drosophila_melanogaster|G025908|GNP_725896.1:79</td></tr>
<tr><td>OG11675</td><td>68</td><td>1</td><td>+Drosophila_melanogaster|G023689|GNP_651538.1:68</td></tr>
<tr><td>OG11754</td><td>80</td><td>2</td><td>+Drosophila_melanogaster|G013864|GNP_476778.1:74</td></tr>
<tr><td>OG1177</td><td>79</td><td>1</td><td>+Drosophila_melanogaster|G015110|GNP_523931.1:79</td></tr>
<tr><td>OG1184</td><td>82</td><td>2</td><td>+Drosophila_melanogaster|G013858|GNP_476772.1+Drosophila_melanogaster|G015184|GNP_524009.2+Drosophila_melanogaster|G015414|GNP_524264.1+Drosophila_melanogaster|G015445|GNP_524297.1:79</td></tr>
<tr><td>OG11903</td><td>80</td><td>1</td><td>+Drosophila_melanogaster|G022661|GNP_650214.1+Drosophila_melanogaster|G028298|GNP_731727.2:80</td></tr>
<tr><td>OG12034</td><td>82</td><td>2</td><td>+Drosophila_melanogaster|G009831|GNP_001262578.1+Drosophila_melanogaster|G009832|GNP_001262579.1+Drosophila_melanogaster|G028398|GNP_731941.1:78</td></tr>
<tr><td>OG12043</td><td>83</td><td>3</td><td>+Drosophila_melanogaster|G009831|GNP_001262578.1+Drosophila_melanogaster|G009832|GNP_001262579.1+Drosophila_melanogaster|G028398|GNP_731941.1:78</td></tr>
</table>

| | | | |
|---|---|---|---|
| OG12103 | 75 | 1 | +Drosophila_melanogasterlG015813lGNP_524719.1:75 |
| OG1215 | 81 | 1 | +Drosophila_melanogasterlG026041lGNP_726243.1:81 |
| OG12175 | 83 | 1 | +Drosophila_melanogasterlG005421lGNP_001246276.1+Drosophila_melanogasterlG005422lGNP_001246277.1+Drosophila_melanogasterlG012021lGNP_001286337.1+Drosophila_melanogasterlG014899lGNP_523710.1+Drosophila_melanogasterlG025527lGNP_725120.1:83 |
| OG12223 | 77 | 1 | +Drosophila_melanogasterlG020399lGNP_611878.1+Drosophila_melanogasterlG026124lGNP_726411.1:77 |
| OG12247 | 78 | 1 | +Drosophila_melanogasterlG014010lGNP_476964.1+Drosophila_melanogasterlG025922lGNP_725943.1:78 |
| OG1234 | 67 | 3 | +Drosophila_melanogasterlG002702lGNP_001138070.1+Drosophila_melanogasterlG002703lGNP_001138071.1+Drosophila_melanogasterlG002704lGNP_001138072.1+Drosophila_melanogasterlG006224lGNP_001247176.1+Drosophila_melanogasterlG006225lGNP_001247177.1+Drosophila_melanogasterlG006226lGNP_001247178.1+Drosophila_melanogasterlG006227lGNP_001247179.1+Drosophila_melanogasterlG009944lGNP_001262694.1+Drosophila_melanogasterlG009945lGNP_001262695.1+Drosophila_melanogasterlG023017lGNP_650681.2+Drosophila_melanogasterlG023715lGNP_651569.1+Drosophila_melanogasterlG028543lGNP_732293.1+Drosophila_melanogasterlG028544lGNP_732294.1+Drosophila_melanogasterlG028545lGNP_732295.1+Drosophila_melanogasterlG028546lGNP_732296.1+Drosophila_melanogasterlG028990lGNP_733215.1+Drosophila_melanogasterlG028991lGNP_733216.1+Drosophila_melanogasterlG028992lGNP_733217.1+Drosophila_melanogasterlG028993lGNP_733218.1+Drosophila_melanogasterlG028994lGNP_733219.1:53 |
| OG12579 | 80 | 1 | +Drosophila_melanogasterlG010687lGNP_001284995.1+Drosophila_melanogasterlG015957lGNP_524884.1+Drosophila_melanogasterlG016143lGNP_536352.1+Drosophila_melanogasterlG026490lGNP_727218.1:80 |
| OG1269 | 76 | 4 | +Drosophila_melanogasterlG004933lGNP_001245715.1+Drosophila_melanogasterlG004934lGNP_001245716.1+Drosophila_melanogasterlG004935lGNP_001245717.1+Drosophila_melanogasterlG006411lGNP_001247377.1+Drosophila_melanogasterlG006412lGNP_001247378.2+Drosophila_melanogasterlG006974lGNP_001259622.1+Drosophila_melanogasterlG014597lGNP_523373.2+Drosophila_melanogasterlG015712lGNP_524600.3+Drosophila_melanogasterlG026798lGNP_727985.2+Drosophila_melanogasterlG026799lGNP_727986.2:64 |
| OG12776 | 73 | 1 | +Drosophila_melanogasterlG017857lGNP_608830.3:73 |
| OG1279 | 82 | 3 | +Drosophila_melanogasterlG012005lGNP_001286321.1+Drosophila_melanogasterlG012006lGNP_001286322.1+Drosophila_melanogasterlG014343lGNP_477375.1+Drosophila_melanogasterlG015723lGNP_524611.1+Drosophila_melanogasterlG025509lGNP_725085.1+Drosophila_melanogasterlG029180lGNP_733449.1+Drosophila_melanogasterlG030286lGNP_996315.1+Drosophila_melanogasterlG030287lGNP_996316.1:78 |
| OG12847 | 83 | 4 | +Drosophila_melanogasterlG023236lGNP_650961.2:73 |

| | | | |
|---|---|---|---|
| OG12881 | 80 | 1 | +Drosophila_melanogasterlG012094lGNP_001286410.1+Drosophila_melanogasterlG019604lGNP_610939.2:80 |
| OG12931 | 78 | 1 | +Drosophila_melanogasterlG000790lGNP_001033885.1+Drosophila_melanogasterlG011433lGNP_001285747.1+Drosophila_melanogasterlG013986lGNP_476938.1:78 |
| OG13008 | 75 | 1 | +Drosophila_melanogasterlG014528lGNP_511117.1:75 |
| OG13036 | 76 | 1 | +Drosophila_melanogasterlG013750lGNP_476636.1:76 |
| OG13068 | 75 | 1 | +Drosophila_melanogasterlG017146lGNP_573076.1+Drosophila_melanogasterlG026743lGNP_727896.1+Drosophila_melanogasterlG026744lGNP_727897.1+Drosophila_melanogasterlG030404lGNP_996448.1+Drosophila_melanogasterlG030405lGNP_996449.1:75 |
| OG13076 | 76 | 1 | +Drosophila_melanogasterlG019715lGNP_611074.1:76 |
| OG13084 | 81 | 1 | +Drosophila_melanogasterlG013921lGNP_476853.1+Drosophila_melanogasterlG025591lGNP_725235.1:81 |
| OG13130 | 77 | 3 | +Drosophila_melanogasterlG019763lGNP_611136.1:32 |
| OG13134 | 79 | 2 | +Drosophila_melanogasterlG014464lGNP_511049.1:76 |
| OG13144 | 83 | 1 | +Drosophila_melanogasterlG005421lGNP_001246276.1+Drosophila_melanogasterlG005422lGNP_001246277.1+Drosophila_melanogasterlG012021lGNP_001286337.1+Drosophila_melanogasterlG014899lGNP_523710.1+Drosophila_melanogasterlG025527lGNP_725120.1:83 |
| OG1317 | 83 | 3 | +Drosophila_melanogasterlG003840lGNP_001163753.1+Drosophila_melanogasterlG012515lGNP_001286835.1+Drosophila_melanogasterlG014981lGNP_523795.2+Drosophila_melanogasterlG015025lGNP_523842.2+Drosophila_melanogasterlG015438lGNP_524290.2+Drosophila_melanogasterlG023742lGNP_651606.2+Drosophila_melanogasterlG025908lGNP_725896.1:79 |
| OG1323 | 79 | 1 | +Drosophila_melanogasterlG018799lGNP_609965.1:79 |
| OG1325 | 78 | 1 | +Drosophila_melanogasterlG003781lGNP_001163686.1+Drosophila_melanogasterlG023299lGNP_651049.3+Drosophila_melanogasterlG028742lGNP_732744.1:78 |
| OG13309 | 83 | 3 | +Drosophila_melanogasterlG014590lGNP_523366.2:80 |
| OG13352 | 81 | 1 | +Drosophila_melanogasterlG013059lGNP_001287386.1+Drosophila_melanogasterlG015820lGNP_524728.2+Drosophila_melanogasterlG030219lGNP_996231.1:81 |
| OG1337 | 83 | 3 | +Drosophila_melanogasterlG003840lGNP_001163753.1+Drosophila_melanogasterlG012515lGNP_001286835.1+Drosophila_melanogasterlG014981lGNP_523795.2+Drosophila_melanogasterlG015025lGNP_523842.2+Drosophila_melanogasterlG015438lGNP_524290.2+Drosophila_melanogasterlG023742lGNP_651606.2+Drosophila_melanogasterlG025908lGNP_725896.1:79 |
| OG1348 | 81 | 4 | +Drosophila_melanogasterlG009368lGNP_001262104.1+Drosophila_melanogasterlG009369lGNP_001262105.1+Drosophila_melanogasterlG021874lGNP_649210.1+Drosophila_melanogasterlG027810lGNP_730517.1+Drosophila_melanogasterlG027811lGNP_730518.1+Drosophila_melanogasterlG027812lGNP_730519. |

| | | | |
|---|---|---|---|
| | | | 1+Drosophila_melanogaster\|G027813\|GNP_730520.1+Drosophila_melanogaster\|G029459\|GNP_788547.1:70 |
| OG13634 | 79 | 2 | +Drosophila_melanogaster\|G012619\|GNP_001286940.1+Drosophila_melanogaster\|G020882\|GNP_647959.1:76 |
| OG13832 | 83 | 2 | +Drosophila_melanogaster\|G013858\|GNP_476772.1:82 |
| OG13968 | 78 | 1 | +Drosophila_melanogaster\|G009293\|GNP_001262025.1+Drosophila_melanogaster\|G013388\|GNP_001303372.1+Drosophila_melanogaster\|G021760\|GNP_649070.1:78 |
| OG14123 | 75 | 1 | +Drosophila_melanogaster\|G014998\|GNP_523813.1:75 |
| OG1423 | 80 | 1 | +Drosophila_melanogaster\|G013165\|GNP_001287493.1+Drosophila_melanogaster\|G015606\|GNP_524469.2:80 |
| OG1427 | 69 | 2 | +Drosophila_melanogaster\|G020715\|GNP_647746.1+Drosophila_melanogaster\|G030211\|GNP_996221.1+Drosophila_melanogaster\|G030212\|GNP_996222.1:62 |
| OG14312 | 76 | 1 | +Drosophila_melanogaster\|G012411\|GNP_001286731.1+Drosophila_melanogaster\|G020236\|GNP_611685.1:76 |
| OG14521 | 73 | 1 | +Drosophila_melanogaster\|G000167\|GNP_001014633.2:73 |
| OG14607 | 68 | 1 | +Drosophila_melanogaster\|G018211\|GNP_609246.1:68 |
| OG1480 | 78 | 3 | +Drosophila_melanogaster\|G000152\|GNP_001014617.1+Drosophila_melanogaster\|G009775\|GNP_001262520.1+Drosophila_melanogaster\|G009776\|GNP_001262521.1+Drosophila_melanogaster\|G009777\|GNP_001262522.1+Drosophila_melanogaster\|G009778\|GNP_001262523.1+Drosophila_melanogaster\|G009779\|GNP_001262524.1+Drosophila_melanogaster\|G015483\|GNP_524336.2+Drosophila_melanogaster\|G028310\|GNP_731762.1+Drosophila_melanogaster\|G028311\|GNP_731763.1+Drosophila_melanogaster\|G028312\|GNP_731764.1:58 |
| OG15046 | 70 | 3 | +Drosophila_melanogaster\|G004610\|GNP_001189293.1+Drosophila_melanogaster\|G015649\|GNP_524514.2+Drosophila_melanogaster\|G028926\|GNP_733133.1+Drosophila_melanogaster\|G028927\|GNP_733134.1+Drosophila_melanogaster\|G028928\|GNP_733135.1:10 |
| OG1506 | 82 | 2 | +Drosophila_melanogaster\|G009840\|GNP_001262587.1+Drosophila_melanogaster\|G028418\|GNP_731993.1+Drosophila_melanogaster\|G028419\|GNP_731994.1:70 |
| OG15319 | 77 | 1 | +Drosophila_melanogaster\|G002938\|GNP_001162738.1+Drosophila_melanogaster\|G015806\|GNP_524709.1+Drosophila_melanogaster\|G026672\|GNP_727690.1+Drosophila_melanogaster\|G026673\|GNP_727692.1+Drosophila_melanogaster\|G026674\|GNP_727693.1:77 |
| OG15504 | 80 | 1 | +Drosophila_melanogaster\|G007358\|GNP_001260018.1+Drosophila_melanogaster\|G013862\|GNP_476776.1:80 |
| OG1569 | 82 | 5 | +Drosophila_melanogaster\|G029994\|GNP_995971.2:1 |
| OG1603 | 62 | 3 | +Drosophila_melanogaster\|G007886\|GNP_001260565.1+Drosophila_melanogaster\|G013883\|GNP_476804.1+Drosophila_melanogaster\|G013973\|GNP_476922.1:41 |
| OG1605 | 79 | 1 | +Drosophila_melanogaster\|G022419\|GNP_649906.1:79 |

| | | | |
|---|---|---|---|
| OG16210 | 78 | 1 | +Drosophila_melanogasterlG000565lGNP_001027283.1+Drosophila_melanogasterlG000569lGNP_001027287.1+Drosophila_melanogasterlG000573lGNP_001027291.1+Drosophila_melanogasterlG000578lGNP_001027296.1+Drosophila_melanogasterlG000582lGNP_001027300.1+Drosophila_melanogasterlG000587lGNP_001027305.1+Drosophila_melanogasterlG000592lGNP_001027310.1+Drosophila_melanogasterlG000597lGNP_001027315.1+Drosophila_melanogasterlG000602lGNP_001027320.1+Drosophila_melanogasterlG000607lGNP_001027325.1+Drosophila_melanogasterlG000612lGNP_001027330.1+Drosophila_melanogasterlG000617lGNP_001027335.1+Drosophila_melanogasterlG000622lGNP_001027340.1+Drosophila_melanogasterlG000627lGNP_001027345.1+Drosophila_melanogasterlG000632lGNP_001027350.1+Drosophila_melanogasterlG000637lGNP_001027355.1+Drosophila_melanogasterlG000642lGNP_001027360.1+Drosophila_melanogasterlG000647lGNP_001027365.1+Drosophila_melanogasterlG000652lGNP_001027370.1+Drosophila_melanogasterlG000657lGNP_001027375.1+Drosophila_melanogasterlG000662lGNP_001027380.1+Drosophila_melanogasterlG000667lGNP_001027385.1+Drosophila_melanogasterlG025143lGNP_724342.1:78 |
| OG1624 | 80 | 1 | +Drosophila_melanogasterlG007612lGNP_001260276.1+Drosophila_melanogasterlG011457lGNP_001285771.1+Drosophila_melanogasterlG014723lGNP_523517.1+Drosophila_melanogasterlG024731lGNP_723438.1:80 |
| OG1625 | 73 | 3 | +Drosophila_melanogasterlG020852lGNP_647918.2+Drosophila_melanogasterlG024039lGNP_651978.2:52 |
| OG16557 | 80 | 1 | +Drosophila_melanogasterlG027584lGNP_729866.1:80 |
| OG1684 | 82 | 1 | +Drosophila_melanogasterlG008840lGNP_001261559.1+Drosophila_melanogasterlG015146lGNP_523968.1:82 |
| OG17040 | 78 | 1 | +Drosophila_melanogasterlG000565lGNP_001027283.1+Drosophila_melanogasterlG000569lGNP_001027287.1+Drosophila_melanogasterlG000573lGNP_001027291.1+Drosophila_melanogasterlG000578lGNP_001027296.1+Drosophila_melanogasterlG000582lGNP_001027300.1+Drosophila_melanogasterlG000587lGNP_001027305.1+Drosophila_melanogasterlG000592lGNP_001027310.1+Drosophila_melanogasterlG000597lGNP_001027315.1+Drosophila_melanogasterlG000602lGNP_001027320.1+Drosophila_melanogasterlG000607lGNP_001027325.1+Drosophila_melanogasterlG000612lGNP_001027330.1+Drosophila_melanogasterlG000617lGNP_001027335.1+Drosophila_melanogasterlG000622lGNP_001027340.1+Drosophila_melanogasterlG000627lGNP_001027345.1+Drosophila_melanogasterlG000632lGNP_001027350.1+Drosophila_melanogasterlG000637lGNP_001027355.1+Drosophila_melanogasterlG000642lGNP_001027360.1+Drosophila_melanogasterlG000647lGNP_001027365.1+Drosophila_melanogasterlG000652lGNP_001027370.1+Drosophila_melanogasterlG000657lGNP_001027375.1+Drosophila_melanogasterlG000662lGNP_001027380.1+Drosophila_melanogasterlG000667lGNP_001027385.1+Drosophila_melanogasterlG025143lGNP_724342.1:78 |
| OG17078 | 64 | 1 | +Drosophila_melanogasterlG001374lGNP_001096936.1+Drosophila_melanogasterlG006759lGNP_001259394.1+Drosophila_melanogasterlG006760lGNP_001259395.1+Drosophila_melanogasterlG006761lGNP_00 |

| | | | |
|---|---|---|---|
| | | | 1259396.1+Drosophila_melanogasterlG016784lGNP_572609.1:64 |
| OG17091 | 73 | 1 | +Drosophila_melanogasterlG022210lGNP_649645.1:73 |
| OG17098 | 82 | 3 | +Drosophila_melanogasterlG000574lGNP_001027292.1+Drosophila_melanogasterlG000583lGNP_001027301.1+Drosophila_melanogasterlG000588lGNP_001027306.1+Drosophila_melanogasterlG000593lGNP_001027311.1+Drosophila_melanogasterlG000598lGNP_001027316.1+Drosophila_melanogasterlG000603lGNP_001027321.1+Drosophila_melanogasterlG000608lGNP_001027326.1+Drosophila_melanogasterlG000613lGNP_001027331.1+Drosophila_melanogasterlG000618lGNP_001027336.1+Drosophila_melanogasterlG000623lGNP_001027341.1+Drosophila_melanogasterlG000628lGNP_001027346.1+Drosophila_melanogasterlG000633lGNP_001027351.1+Drosophila_melanogasterlG000638lGNP_001027356.1+Drosophila_melanogasterlG000643lGNP_001027361.1+Drosophila_melanogasterlG000663lGNP_001027381.1+Drosophila_melanogasterlG000668lGNP_001027386.1+Drosophila_melanogasterlG025144lGNP_724343.1:73 |
| OG1726 | 81 | 2 | +Drosophila_melanogasterlG024565lGNP_723089.1+Drosophila_melanogasterlG024566lGNP_723090.1+Drosophila_melanogasterlG024567lGNP_723091.1:77 |
| OG1727 | 73 | 2 | +Drosophila_melanogasterlG012943lGNP_001287269.1+Drosophila_melanogasterlG013943lGNP_476880.1+Drosophila_melanogasterlG013944lGNP_476881.1+Drosophila_melanogasterlG017414lGNP_599111.1+Drosophila_melanogasterlG017415lGNP_599112.1+Drosophila_melanogasterlG017416lGNP_599113.1+Drosophila_melanogasterlG028193lGNP_731451.1+Drosophila_melanogasterlG028194lGNP_731452.1+Drosophila_melanogasterlG028195lGNP_731453.1:58 |
| OG17291 | 77 | 1 | +Drosophila_melanogasterlG010447lGNP_001284754.1+Drosophila_melanogasterlG016215lGNP_569852.1:77 |
| OG17319 | 76 | 1 | +Drosophila_melanogasterlG026570lGNP_727447.1:76 |
| OG17437 | 73 | 1 | +Drosophila_melanogasterlG020942lGNP_648037.1:73 |
| OG1753 | 83 | 2 | +Drosophila_melanogasterlG005102lGNP_001245907.1+Drosophila_melanogasterlG007455lGNP_001260117.1+Drosophila_melanogasterlG013711lGNP_476595.1+Drosophila_melanogasterlG024583lGNP_723137.1+Drosophila_melanogasterlG024584lGNP_723138.1+Drosophila_melanogasterlG024585lGNP_723139.1:72 |
| OG17636 | 70 | 1 | +Drosophila_melanogasterlG009199lGNP_001261931.1+Drosophila_melanogasterlG021581lGNP_648849.3:70 |
| OG17794 | 67 | 1 | +Drosophila_melanogasterlG015861lGNP_524774.1:67 |
| OG18162 | 75 | 1 | +Drosophila_melanogasterlG007859lGNP_001260537.1+Drosophila_melanogasterlG011726lGNP_001286042.1+Drosophila_melanogasterlG014796lGNP_523595.1+Drosophila_melanogasterlG025038lGNP_724109.1+Drosophila_melanogasterlG025039lGNP_724110.1:75 |
| OG18359 | 77 | 1 | +Drosophila_melanogasterlG014980lGNP_523794.1+Drosophila_melanogasterlG025907lGNP_725894.1:77 |
| OG19046 | 72 | 1 | +Drosophila_melanogasterlG020752lGNP_647792.1:72 |

| | | | |
|---|---|---|---|
| OG1937 | 69 | 2 | +Drosophila_melanogasterlG020715lGNP_647746.1+Drosophila_melanogasterlG030211lGNP_996221.1+Drosophila_melanogasterlG030212lGNP_996222.1:62 |
| OG19418 | 72 | 1 | +Drosophila_melanogasterlG018848lGNP_610024.1:72 |
| OG1953 | 83 | 1 | +Drosophila_melanogasterlG015463lGNP_524316.1+Drosophila_melanogasterlG028229lGNP_731548.2:83 |
| OG1954 | 79 | 2 | +Drosophila_melanogasterlG010799lGNP_001285108.1+Drosophila_melanogasterlG016839lGNP_572686.1+Drosophila_melanogasterlG022448lGNP_649938.2+Drosophila_melanogasterlG028180lGNP_731401.3:68 |
| OG19655 | 63 | 4 | +Drosophila_melanogasterlG005975lGNP_001246908.1+Drosophila_melanogasterlG022084lGNP_649483.1+Drosophila_melanogasterlG027963lGNP_730861.1:29 |
| OG197 | 80 | 1 | +Drosophila_melanogasterlG003921lGNP_001188532.1+Drosophila_melanogasterlG003922lGNP_001188533.1+Drosophila_melanogasterlG006547lGNP_001259164.1+Drosophila_melanogasterlG006548lGNP_001259165.1+Drosophila_melanogasterlG006549lGNP_001259166.1+Drosophila_melanogasterlG006550lGNP_001259167.1+Drosophila_melanogasterlG014434lGNP_477484.2+Drosophila_melanogasterlG014435lGNP_477485.1+Drosophila_melanogasterlG026326lGNP_726784.1:80 |
| OG19715 | 71 | 1 | +Drosophila_melanogasterlG011423lGNP_001285737.1+Drosophila_melanogasterlG018149lGNP_609179.2:71 |
| OG1983 | 80 | 2 | +Drosophila_melanogasterlG011344lGNP_001285657.1+Drosophila_melanogasterlG017991lGNP_608987.1+Drosophila_melanogasterlG019846lGNP_611238.2:71 |
| OG19839 | 66 | 1 | +Drosophila_melanogasterlG000531lGNP_001027236.1+Drosophila_melanogasterlG004152lGNP_001188778.1:66 |
| OG19883 | 73 | 1 | +Drosophila_melanogasterlG015605lGNP_524467.1:73 |
| OG19978 | 78 | 1 | +Drosophila_melanogasterlG022365lGNP_649840.1:78 |
| OG19983 | 59 | 1 | +Drosophila_melanogasterlG003273lGNP_001163119.1+Drosophila_melanogasterlG019390lGNP_610691.1:59 |
| OG20031 | 74 | 1 | +Drosophila_melanogasterlG007871lGNP_001260550.2+Drosophila_melanogasterlG015787lGNP_524687.1+Drosophila_melanogasterlG025054lGNP_724149.1:74 |
| OG2006 | 82 | 3 | +Drosophila_melanogasterlG005102lGNP_001245907.1+Drosophila_melanogasterlG007455lGNP_001260117.1+Drosophila_melanogasterlG013711lGNP_476595.1+Drosophila_melanogasterlG022324lGNP_649788.2+Drosophila_melanogasterlG024583lGNP_723137.1+Drosophila_melanogasterlG024584lGNP_723138.1+Drosophila_melanogasterlG024585lGNP_723139.1:69 |
| OG20114 | 72 | 1 | +Drosophila_melanogasterlG015562lGNP_524421.1:72 |
| OG20215 | 77 | 1 | +Drosophila_melanogasterlG015462lGNP_524315.2+Drosophila_melanogasterlG028227lGNP_731544.1:77 |
| OG2044 | 78 | 1 | +Drosophila_melanogasterlG001197lGNP_001036624.1+Drosophila_melanogasterlG012827lGNP_001287151.1+Drosophila_melanogasterlG012828lGNP_001287152.1+Drosophila_melanogasterlG012829lGNP_00 |

| | | | |
|---|---|---|---|
| | | | 1287153.1+Drosophila_melanogasterlG015988lGNP_524918.1+Drosophila_melanogasterlG027923lGNP_730774.1+Drosophila_melanogasterlG027924lGNP_730775.1:78 |
| OG20658 | 74 | 1 | +Drosophila_melanogasterlG015980lGNP_524910.1:74 |
| OG20743 | 70 | 1 | +Drosophila_melanogasterlG018689lGNP_609836.1:70 |
| OG2110 | 81 | 2 | +Drosophila_melanogasterlG012716lGNP_001287039.1+Drosophila_melanogasterlG016547lGNP_572308.3+Drosophila_melanogasterlG021317lGNP_648525.1:71 |
| OG21140 | 65 | 1 | +Drosophila_melanogasterlG006731lGNP_001259363.1+Drosophila_melanogasterlG006732lGNP_001259364.1+Drosophila_melanogasterlG006733lGNP_001259365.1+Drosophila_melanogasterlG015766lGNP_524662.1+Drosophila_melanogasterlG026514lGNP_727293.1:65 |
| OG21314 | 82 | 2 | +Drosophila_melanogasterlG028218lGNP_731510.1:74 |
| OG2243 | 64 | 1 | +Drosophila_melanogasterlG022081lGNP_649476.1:64 |
| OG22471 | 80 | 1 | +Drosophila_melanogasterlG011668lGNP_001285984.1+Drosophila_melanogasterlG014183lGNP_477176.1:80 |
| OG22573 | 79 | 5 | +Drosophila_melanogasterlG008553lGNP_001261269.1:62 |
| OG2280 | 75 | 4 | +Drosophila_melanogasterlG030145lGNP_996145.1:49 |
| OG2332 | 78 | 5 | +Drosophila_melanogasterlG003862lGNP_001163778.1+Drosophila_melanogasterlG004633lGNP_001189321.1+Drosophila_melanogasterlG004634lGNP_001189322.1+Drosophila_melanogasterlG004635lGNP_001189323.1+Drosophila_melanogasterlG004636lGNP_001189324.1+Drosophila_melanogasterlG004637lGNP_001189325.1+Drosophila_melanogasterlG004638lGNP_001189326.1+Drosophila_melanogasterlG010346lGNP_001263115.1+Drosophila_melanogasterlG010347lGNP_001263116.1+Drosophila_melanogasterlG010348lGNP_001263117.1+Drosophila_melanogasterlG023911lGNP_651813.2+Drosophila_melanogasterlG029125lGNP_733386.1+Drosophila_melanogasterlG029126lGNP_733387.1+Drosophila_melanogasterlG029127lGNP_733388.1+Drosophila_melanogasterlG029128lGNP_733389.2+Drosophila_melanogasterlG029129lGNP_733390.1:59 |
| OG2366 | 61 | 1 | +Drosophila_melanogasterlG015478lGNP_524331.2:61 |
| OG2424 | 74 | 1 | +Drosophila_melanogasterlG027996lGNP_730935.1:74 |
| OG2429 | 82 | 3 | +Drosophila_melanogasterlG013582lGNP_001303567.1+Drosophila_melanogasterlG016111lGNP_525090.1+Drosophila_melanogasterlG030414lGNP_996459.1+Drosophila_melanogasterlG030415lGNP_996460.1+Drosophila_melanogasterlG030416lGNP_996461.1+Drosophila_melanogasterlG030417lGNP_996462.1:78 |
| OG24382 | 67 | 1 | +Drosophila_melanogasterlG023553lGNP_651364.1:67 |
| OG2478 | 82 | 3 | +Drosophila_melanogasterlG016669lGNP_572452.1:48 |
| OG2501 | 72 | 1 | +Drosophila_melanogasterlG014473lGNP_511058.1:72 |
| OG2506 | 75 | 4 | +Drosophila_melanogasterlG009600lGNP_001262340.1:56 |

| | | | |
|---|---|---|---|
| OG2545 | 75 | 5 | +Drosophila_melanogaster\|G010113\|GNP_001262872.1:46 |
| OG2575 | 78 | 1 | +Drosophila_melanogaster\|G009226\|GNP_001261958.1+Drosophila_melanogaster\|G015282\|GNP_524117.1:78 |
| OG2599 | 80 | 3 | +Drosophila_melanogaster\|G024039\|GNP_651978.2:44 |
| OG26549 | 60 | 1 | +Drosophila_melanogaster\|G001537\|GNP_001097135.1:60 |
| OG2685 | 82 | 6 | +Drosophila_melanogaster\|G006772\|GNP_001259407.1+Drosophila_melanogaster\|G010160\|GNP_001262919.1+Drosophila_melanogaster\|G015621\|GNP_524484.1+Drosophila_melanogaster\|G015830\|GNP_524738.1+Drosophila_melanogaster\|G015990\|GNP_524921.1+Drosophila_melanogaster\|G016002\|GNP_524937.1+Drosophila_melanogaster\|G026557\|GNP_727418.1:72 |
| OG2704 | 82 | 2 | +Drosophila_melanogaster\|G016146\|GNP_536733.1:77 |
| OG2774 | 82 | 3 | +Drosophila_melanogaster\|G010363\|GNP_001263132.1+Drosophila_melanogaster\|G015714\|GNP_524602.1+Drosophila_melanogaster\|G029150\|GNP_733414.1+Drosophila_melanogaster\|G029151\|GNP_733415.1:52 |
| OG2986 | 81 | 3 | +Drosophila_melanogaster\|G011411\|GNP_001285724.1+Drosophila_melanogaster\|G011412\|GNP_001285725.1+Drosophila_melanogaster\|G013884\|GNP_476805.1:72 |
| OG3062 | 82 | 3 | +Drosophila_melanogaster\|G002937\|GNP_001162737.1+Drosophila_melanogaster\|G010854\|GNP_001285163.1+Drosophila_melanogaster\|G010855\|GNP_001285164.1+Drosophila_melanogaster\|G014549\|GNP_511140.1+Drosophila_melanogaster\|G026647\|GNP_727631.1+Drosophila_melanogaster\|G026648\|GNP_727632.1:75 |
| OG3080 | 83 | 1 | +Drosophila_melanogaster\|G007554\|GNP_001260218.1+Drosophila_melanogaster\|G011410\|GNP_001285723.1+Drosophila_melanogaster\|G014259\|GNP_477269.1:83 |
| OG3144 | 68 | 1 | +Drosophila_melanogaster\|G005010\|GNP_001245797.1+Drosophila_melanogaster\|G005011\|GNP_001245798.1+Drosophila_melanogaster\|G011214\|GNP_001285526.1+Drosophila_melanogaster\|G017560\|GNP_608461.1:68 |
| OG3185 | 83 | 1 | +Drosophila_melanogaster\|G017917\|GNP_608905.1:83 |
| OG3204 | 82 | 2 | +Drosophila_melanogaster\|G011177\|GNP_001285489.1+Drosophila_melanogaster\|G011178\|GNP_001285490.1+Drosophila_melanogaster\|G013585\|GNP_001303570.1+Drosophila_melanogaster\|G015886\|GNP_524803.1+Drosophila_melanogaster\|G026924\|GNP_728342.1:71 |
| OG3318 | 77 | 4 | +Drosophila_melanogaster\|G008920\|GNP_001261640.1+Drosophila_melanogaster\|G008921\|GNP_001261641.1+Drosophila_melanogaster\|G008922\|GNP_001261643.1+Drosophila_melanogaster\|G008923\|GNP_001261644.1+Drosophila_melanogaster\|G012688\|GNP_001287010.1+Drosophila_melanogaster\|G012689\|GNP_001287011.1+Drosophila_melanogaster\|G012690\|GNP_001287012.1+Drosophila_melanogaster\|G021184\|GNP_648345.1+Drosophila_melanogaster\|G027439\|GNP_729528.2:65 |
| OG3411 | 82 | 3 | +Drosophila_melanogaster\|G009890\|GNP_001262639.1+Drosophila_melanogaster\|G014403\|GNP_477442.1:54 |

| | | | |
|---|---|---|---|
| OG3672 | 81 | 3 | +Drosophila_melanogaster\|G006647\|GNP_001259272.1+Drosophila_melanogaster\|G006648\|GNP_001259273.1+Drosophila_melanogaster\|G006649\|GNP_001259274.1+Drosophila_melanogaster\|G014476\|GNP_511061.1:63 |
| OG375 | 62 | 1 | +Drosophila_melanogaster\|G009696\|GNP_001262441.1+Drosophila_melanogaster\|G022462\|GNP_649954.1:62 |
| OG38 | 72 | 2 | +Drosophila_melanogaster\|G001411\|GNP_001096987.2+Drosophila_melanogaster\|G014535\|GNP_511124.1:29 |
| OG39 | 76 | 1 | +Drosophila_melanogaster\|G001417\|GNP_001096993.1+Drosophila_melanogaster\|G001418\|GNP_001096994.1+Drosophila_melanogaster\|G010991\|GNP_001285300.1+Drosophila_melanogaster\|G014078\|GNP_477042.1+Drosophila_melanogaster\|G026747\|GNP_727901.1+Drosophila_melanogaster\|G030407\|GNP_996451.1+Drosophila_melanogaster\|G030408\|GNP_996452.1:76 |
| OG4231 | 67 | 1 | +Drosophila_melanogaster\|G023732\|GNP_651595.1:67 |
| OG4438 | 71 | 1 | +Drosophila_melanogaster\|G013916\|GNP_476848.1:71 |
| OG4500 | 75 | 1 | +Drosophila_melanogaster\|G007679\|GNP_001260349.1+Drosophila_melanogaster\|G013950\|GNP_476888.1:75 |
| OG4615 | 71 | 9 | +Drosophila_melanogaster\|G009046\|GNP_001261772.1:4 |
| OG4636 | 69 | 1 | +Drosophila_melanogaster\|G012193\|GNP_001286511.1+Drosophila_melanogaster\|G013909\|GNP_476838.1:69 |
| OG4733 | 76 | 2 | +Drosophila_melanogaster\|G028553\|GNP_732312.1:74 |
| OG4754 | 73 | 1 | +Drosophila_melanogaster\|G007218\|GNP_001259877.1+Drosophila_melanogaster\|G014046\|GNP_477005.1:73 |
| OG4810 | 72 | 3 | +Drosophila_melanogaster\|G017464\|GNP_608335.1+Drosophila_melanogaster\|G026908\|GNP_728284.1:63 |
| OG4960 | 80 | 1 | +Drosophila_melanogaster\|G012732\|GNP_001287055.1+Drosophila_melanogaster\|G015223\|GNP_524053.2+Drosophila_melanogaster\|G027585\|GNP_729871.1:80 |
| OG5235 | 80 | 3 | +Drosophila_melanogaster\|G025435\|GNP_724887.2+Drosophila_melanogaster\|G025436\|GNP_724888.2:21 |
| OG5237 | 81 | 1 | +Drosophila_melanogaster\|G015819\|GNP_524726.1+Drosophila_melanogaster\|G027101\|GNP_728756.1:81 |
| OG5248 | 76 | 2 | +Drosophila_melanogaster\|G028553\|GNP_732312.1:74 |
| OG5265 | 82 | 1 | +Drosophila_melanogaster\|G007634\|GNP_001260303.1+Drosophila_melanogaster\|G011471\|GNP_001285785.1+Drosophila_melanogaster\|G013939\|GNP_476874.1:82 |
| OG5408 | 79 | 1 | +Drosophila_melanogaster\|G015215\|GNP_524043.1:79 |
| OG5777 | 81 | 2 | +Drosophila_melanogaster\|G014487\|GNP_511073.1:76 |
| OG5834 | 76 | 5 | +Drosophila_melanogaster\|G011117\|GNP_001285428.1:62 |
| OG5895 | 79 | 2 | +Drosophila_melanogaster\|G014991\|GNP_523805.1:77 |
| OG6030 | 72 | 2 | +Drosophila_melanogaster\|G006454\|GNP_001259063.1:70 |

| | | | |
|---|---|---|---|
| OG6174 | 76 | 1 | +Drosophila_melanogasterlG018369lGNP_609444.1:76 |
| OG627 | 79 | 6 | +Drosophila_melanogasterlG009839lGNP_001262586.1+Drosophila_melanogasterlG010830lGNP_001285139.1+Drosophila_melanogasterlG012098lGNP_001286414.1+Drosophila_melanogasterlG014542lGNP_511132.2+Drosophila_melanogasterlG014928lGNP_523741.2+Drosophila_melanogasterlG015233lGNP_524063.1+Drosophila_melanogasterlG015486lGNP_524339.1+Drosophila_melanogasterlG015502lGNP_524356.1+Drosophila_melanogasterlG015611lGNP_524474.1+Drosophila_melanogasterlG015883lGNP_524798.2+Drosophila_melanogasterlG015993lGNP_524927.2+Drosophila_melanogasterlG022659lGNP_650209.1+Drosophila_melanogasterlG026626lGNP_727563.1+Drosophila_melanogasterlG026627lGNP_727564.1+Drosophila_melanogasterlG026628lGNP_727565.1+Drosophila_melanogasterlG027609lGNP_729940.1+Drosophila_melanogasterlG027610lGNP_729941.1+Drosophila_melanogasterlG028271lGNP_731651.1+Drosophila_melanogasterlG028296lGNP_731716.1+Drosophila_melanogasterlG028415lGNP_731987.1+Drosophila_melanogasterlG028416lGNP_731988.1+Drosophila_melanogasterlG028417lGNP_731989.1+Drosophila_melanogasterlG029548lGNP_788663.1+Drosophila_melanogasterlG029562lGNP_788679.1+Drosophila_melanogasterlG029563lGNP_788680.1:69 |
| OG6367 | 83 | 1 | +Drosophila_melanogasterlG010802lGNP_001285111.1+Drosophila_melanogasterlG010803lGNP_001285112.1+Drosophila_melanogasterlG024031lGNP_651969.1+Drosophila_melanogasterlG026596lGNP_727499.1:83 |
| OG6853 | 83 | 3 | +Drosophila_melanogasterlG014179lGNP_477171.1+Drosophila_melanogasterlG029262lGNP_788057.2:72 |
| OG6865 | 77 | 1 | +Drosophila_melanogasterlG004154lGNP_001188780.1:77 |
| OG7 | 62 | 1 | +Drosophila_melanogasterlG019427lGNP_610735.1:62 |
| OG7027 | 79 | 1 | +Drosophila_melanogasterlG014178lGNP_477170.1+Drosophila_melanogasterlG017433lGNP_599137.1:79 |
| OG7268 | 77 | 1 | +Drosophila_melanogasterlG007157lGNP_001259814.1+Drosophila_melanogasterlG014208lGNP_477208.1:77 |
| OG7313 | 76 | 4 | +Drosophila_melanogasterlG003705lGNP_001163604.1+Drosophila_melanogasterlG029555lGNP_788670.1:52 |
| OG7350 | 79 | 1 | +Drosophila_melanogasterlG008043lGNP_001260732.1+Drosophila_melanogasterlG014119lGNP_477090.1:79 |
| OG7467 | 68 | 1 | +Drosophila_melanogasterlG028983lGNP_733208.1:68 |
| OG7628 | 78 | 1 | +Drosophila_melanogasterlG011553lGNP_001285869.1+Drosophila_melanogasterlG014180lGNP_477172.1:78 |
| OG7696 | 80 | 2 | +Drosophila_melanogasterlG011049lGNP_001285360.1+Drosophila_melanogasterlG011050lGNP_001285361.1+Drosophila_melanogasterlG011051lGNP_001285362.1+Drosophila_melanogasterlG014605lGNP_523382.1:79 |
| OG7796 | 79 | 1 | +Drosophila_melanogasterlG006396lGNP_001247362.1+Drosophila_melanogasterlG013266lGNP_001287596.1+Drosophila_melanogasterlG013267lGNP_0012 |

| | | | |
|---|---|---|---|
| | | | 87597.1+Drosophila_melanogaster\|G023853\|GNP_651740.1:79 |
| OG7828 | 81 | 1 | +Drosophila_melanogaster\|G008183\|GNP_001260878.1+Drosophila_melanogaster\|G019336\|GNP_610630.1:81 |
| OG7843 | 78 | 2 | +Drosophila_melanogaster\|G021155\|GNP_648308.1:7 |
| OG791 | 68 | 3 | +Drosophila_melanogaster\|G009839\|GNP_001262586.1+Drosophila_melanogaster\|G010830\|GNP_001285139.1+Drosophila_melanogaster\|G012098\|GNP_001286414.1+Drosophila_melanogaster\|G014542\|GNP_511132.2+Drosophila_melanogaster\|G014928\|GNP_523741.2+Drosophila_melanogaster\|G015233\|GNP_524063.1+Drosophila_melanogaster\|G015486\|GNP_524339.1+Drosophila_melanogaster\|G015502\|GNP_524356.1+Drosophila_melanogaster\|G015611\|GNP_524474.1+Drosophila_melanogaster\|G015883\|GNP_524798.2+Drosophila_melanogaster\|G015993\|GNP_524927.2+Drosophila_melanogaster\|G022659\|GNP_650209.1+Drosophila_melanogaster\|G026626\|GNP_727563.1+Drosophila_melanogaster\|G026627\|GNP_727564.1+Drosophila_melanogaster\|G026628\|GNP_727565.1+Drosophila_melanogaster\|G027609\|GNP_729940.1+Drosophila_melanogaster\|G027610\|GNP_729941.1+Drosophila_melanogaster\|G028271\|GNP_731651.1+Drosophila_melanogaster\|G028296\|GNP_731716.1+Drosophila_melanogaster\|G028415\|GNP_731987.1+Drosophila_melanogaster\|G028416\|GNP_731988.1+Drosophila_melanogaster\|G028417\|GNP_731989.1+Drosophila_melanogaster\|G029548\|GNP_788663.1+Drosophila_melanogaster\|G029562\|GNP_788679.1+Drosophila_melanogaster\|G029563\|GNP_788680.1:61 |
| OG8046 | 83 | 1 | +Drosophila_melanogaster\|G010802\|GNP_001285111.1+Drosophila_melanogaster\|G010803\|GNP_001285112.1+Drosophila_melanogaster\|G024031\|GNP_651969.1+Drosophila_melanogaster\|G026596\|GNP_727499.1:83 |
| OG8056 | 81 | 3 | +Drosophila_melanogaster\|G015666\|GNP_524533.1+Drosophila_melanogaster\|G028995\|GNP_733222.1+Drosophila_melanogaster\|G028996\|GNP_733223.1:46 |
| OG8066 | 81 | 3 | +Drosophila_melanogaster\|G008531\|GNP_001261247.1+Drosophila_melanogaster\|G013998\|GNP_476950.1:68 |
| OG8215 | 72 | 1 | +Drosophila_melanogaster\|G021380\|GNP_648603.1:72 |
| OG8244 | 82 | 1 | +Drosophila_melanogaster\|G013746\|GNP_476631.1+Drosophila_melanogaster\|G029972\|GNP_995941.1:82 |
| OG8257 | 68 | 1 | +Drosophila_melanogaster\|G018336\|GNP_609402.1:68 |
| OG845 | 79 | 4 | +Drosophila_melanogaster\|G005196\|GNP_001246015.1+Drosophila_melanogaster\|G007751\|GNP_001260424.1+Drosophila_melanogaster\|G007752\|GNP_001260425.1+Drosophila_melanogaster\|G007753\|GNP_001260426.1+Drosophila_melanogaster\|G014762\|GNP_523560.2+Drosophila_melanogaster\|G018489\|GNP_609595.1+Drosophila_melanogaster\|G024055\|GNP_652004.2+Drosophila_melanogaster\|G024862\|GNP_723775.1+Drosophila_melanogaster\|G024863\|GNP_723776.1:67 |
| OG847 | 58 | 1 | +Drosophila_melanogaster\|G014529\|GNP_511118.2:58 |
| OG8481 | 81 | 1 | +Drosophila_melanogaster\|G014124\|GNP_477098.1+Drosophila_melanogaster\|G017431\|GNP_599135.1+ |

| | | | |
|---|---|---|---|
| | | | Drosophila_melanogaster\|G017432\|GNP_599136.1+Drosophila_melanogaster\|G025743\|GNP_725524.1+Drosophila_melanogaster\|G029906\|GNP_995849.1+Drosophila_melanogaster\|G029907\|GNP_995850.1:81 |
| OG85 | 78 | 1 | +Drosophila_melanogaster\|G007180\|GNP_001259837.1+Drosophila_melanogaster\|G017613\|GNP_608534.2:78 |
| OG8590 | 74 | 2 | +Drosophila_melanogaster\|G023530\|GNP_651330.1:72 |
| OG8624 | 80 | 1 | +Drosophila_melanogaster\|G009555\|GNP_001262295.1+Drosophila_melanogaster\|G022149\|GNP_649568.1:80 |
| OG8657 | 77 | 1 | +Drosophila_melanogaster\|G012900\|GNP_001287225.1+Drosophila_melanogaster\|G012901\|GNP_001287226.1+Drosophila_melanogaster\|G022307\|GNP_649769.1:77 |
| OG8718 | 72 | 1 | +Drosophila_melanogaster\|G016656\|GNP_572438.1+Drosophila_melanogaster\|G026496\|GNP_727233.1+Drosophila_melanogaster\|G026497\|GNP_727234.1:72 |
| OG8850 | 81 | 1 | +Drosophila_melanogaster\|G014124\|GNP_477098.1+Drosophila_melanogaster\|G017431\|GNP_599135.1+Drosophila_melanogaster\|G017432\|GNP_599136.1+Drosophila_melanogaster\|G025743\|GNP_725524.1+Drosophila_melanogaster\|G029906\|GNP_995849.1+Drosophila_melanogaster\|G029907\|GNP_995850.1:81 |
| OG8895 | 83 | 3 | +Drosophila_melanogaster\|G001973\|GNP_001097667.1+Drosophila_melanogaster\|G001974\|GNP_001097668.2+Drosophila_melanogaster\|G009486\|GNP_001262225.1+Drosophila_melanogaster\|G014003\|GNP_476955.1+Drosophila_melanogaster\|G027909\|GNP_730757.1+Drosophila_melanogaster\|G027910\|GNP_730758.1+Drosophila_melanogaster\|G027911\|GNP_730759.1+Drosophila_melanogaster\|G027912\|GNP_730760.1:80 |
| OG9020 | 80 | 2 | +Drosophila_melanogaster\|G004370\|GNP_001189023.1+Drosophila_melanogaster\|G008579\|GNP_001261295.1+Drosophila_melanogaster\|G013925\|GNP_476857.1:77 |
| OG9104 | 79 | 2 | +Drosophila_melanogaster\|G012686\|GNP_001287008.1+Drosophila_melanogaster\|G015180\|GNP_524004.2+Drosophila_melanogaster\|G027431\|GNP_729506.1:75 |
| OG9191 | 58 | 1 | +Drosophila_melanogaster\|G014457\|GNP_477509.1:58 |
| OG9251 | 67 | 1 | +Drosophila_melanogaster\|G014280\|GNP_477296.1+Drosophila_melanogaster\|G024794\|GNP_723586.1+Drosophila_melanogaster\|G024795\|GNP_723587.1:67 |
| OG9287 | 83 | 4 | +Drosophila_melanogaster\|G014938\|GNP_523751.2+Drosophila_melanogaster\|G025703\|GNP_725452.1+Drosophila_melanogaster\|G025704\|GNP_725453.1+Drosophila_melanogaster\|G025705\|GNP_725454.1+Drosophila_melanogaster\|G025706\|GNP_725455.1:58 |
| OG9408 | 83 | 3 | +Drosophila_melanogaster\|G006478\|GNP_001259087.1+Drosophila_melanogaster\|G015740\|GNP_524631.1:61 |
| OG9477 | 76 | 2 | +Drosophila_melanogaster\|G015096\|GNP_523917.2:2 |
| OG9573 | 76 | 1 | +Drosophila_melanogaster\|G018775\|GNP_609936.1:76 |
| OG9600 | 83 | 3 | +Drosophila_melanogaster\|G007692\|GNP_001260362.1+Drosophila_melanogaster\|G011498\|GNP_001285 |

| | | | |
|---|---|---|---|
| | | | 814.1+Drosophila_melanogasterIG014150IGNP_477137.1+Drosophila_melanogasterIG024808IGNP_723616.1:72 |
| OG9650 | 75 | 1 | +Drosophila_melanogasterIG028731IGNP_732717.1+Drosophila_melanogasterIG028732IGNP_732718.1+Drosophila_melanogasterIG028733IGNP_732719.1+Drosophila_melanogasterIG030243IGNP_996265.1:75 |
| OG9660 | 79 | 1 | +Drosophila_melanogasterIG012294IGNP_001286614.1+Drosophila_melanogasterIG014089IGNP_477054.1:79 |
| OG9704 | 73 | 1 | +Drosophila_melanogasterIG021066IGNP_648201.1:73 |
| OG9778 | 70 | 1 | +Drosophila_melanogasterIG005301IGNP_001246149.1+Drosophila_melanogasterIG018997IGNP_610224.1:70 |
| OG9785 | 83 | 3 | +Drosophila_melanogasterIG015263IGNP_524098.2:65 |
| OG9939 | 79 | 1 | +Drosophila_melanogasterIG017650IGNP_608575.1:79 |

**Table S8.** Chimeric terminals created for fusion of Matrix 3 with published morphological matrices.

*Morphological matrix of Bicknell et al. (46):*

Pycnogonum_litorale = Pycnogonum_litorale
Limulus_polyphemus = Limulus_polyphemus
Tachypleus_gigas = Tachypleus_gigas
Tachypleus_tridentatus = Tachypleus_tridentatus
Carcinoscorpius_rotundicauda = Carcinoscorpius_rotundicauda
Centruroides_vittatus = Centruroides_sculpturatus
Androctonus_australis = Androctonus_australis
Caddo_agilis = Caddo agilis
Chileogovea_oedipus = Chileogovea_oedipus
Cyphophthalmus_duricorius = Siro_boyerae
Leiobunum_rotundum = Leiobunum_verrucosum
Gonyleptes_horridus = Gonyleptes_fragilis
Sclerobunus_robustus = Sclerobunus_robustus
Neobisium_maritimum = Novobisium_sp
Chelifer_cancroides = Cheliferidae_sp
Eremocosta_striata = Eremobates_sp
Galeodes_armeniacus = Galeodes_sp
Eukoenenia_mirabilis = Espelaea_UWisc
Neocarus_texanus = Neoacarus_sp
Amblyomma_americanum = Amblyomma_americanum
Allothrombium_simoni = Dinothrombium_tinctorium
Cryptocellus_goodnighti = Cryptocellus_sp_IZ143922
Ricinoides_atewa = Ricinoides_atewa
Liphistius_malayanus = Liphistius_malayanus
Aphonopelma_hentzi = Aphonopelma_hentzi
Hypochilus_pococki = Hypochilus_pococki
Cupienius_getazi = Cupiennius_sp
Phrynus_marginemaculatus = Phrynus_marginemaculatus
Charinus_victori = Charinus_ioaniticus
Mastigoproctus_giganteus = Mastigoproctus_giganteus
Schizomus_crassicaudatus = Stenochrus_portoricensis

*Morphological matrix of Siveter et al. (47):*

Peripatus = Epiperipatus sp.
Euperipatoides = Opisthopatus kwazululandi
Ammothella = Tanystylum orbiculare
Limulus = Limulus polyphemus
Carcinoscorpius = Carcinoscorpius rotundicauda

Centruroides = (Centruroides sculpturatus)
Heterometrus = Pandinus imperator
Caddo = Caddo agilis
Nipponopsalis = Trogulus
Siro = Siro
Equitius = Larifuga
Idiogaryos = Sternophoridae
Eremocosta = Eremobates_sp
Amblyomma = Amblyomma_americanum
Archegozetes = Archogozetes_longisetosus
Opilioacarus = Adenacarus_sp
Cryptocellus = Cryptocellus_sp_IZ-143922
Prokoenenia = Espelaea_UWisc
Liphistius = Liphistius_malayanus
Aphonopelma = Aphonopelma_hentzi
Cupienius = Cupiennius_sp
Stenochrus = Stenochrus_portoricensis
Mastigoproctus = Mastigoproctus_giganteus
Phrynus = Phrynus marginemaculatus
Scutigera = Scutigera
Lithobius = Lithobius
Craterostigmus = Craterostigmus
Scolopendra = Alipes
Pachymerium = Strigamia
Scutigerella = Symphylella
Pauropus = Pauropus
Narceus = Narceus
Vargula = Eusarsiella
Daphnia = Daphnia_pulex
Calanus = Eurytemora
Orchestia = Hyalella
Tomocerus = Folsomia
Locusta = Gryllus
Drosophila = Drosophila

**File S1. Morphological characters and states.**

1 Number of head shield segments
states: five [cephalosoma/proterosoma]; seven [prosomal shield]
2 Ophthalmic ridges
states: absent; present.
3 Pleural margin of prosomal shield
states: absent; present
4 Cardiac lobe
states: absent; present
5 Prosomal repugnatorial glands
states: absent present,
6 Ozophores
states: absent; present
7 Cucullus
states: absent; present
8 Width of sternal region
states: broad throughout; narrow anteriorly; narrow posteriorly; narrow throughout
9 Division of prosomal sternum
states: undivided; divided
10 Cephalic doublure
states: absent; present
11 Lines demarcating meso- and metapeltidium on the prosomal shield
states: absent; present
12 Genal spines
states: absent; present
13 Anterior median projection of prosomal dorsal shield
states: absent; present
14 Trapezoidal projection of anterior carapace margin
states: absent; present
15 Keels between the median and lateral eyes on the prosomal dorsal shield
states: absent; present
16 Proboscis
states: absent; present,
17 Direction of mouth opening
states: directed anteroventrally; directed posteroventrally
18 Labium or tritosternum
states: absent; present
19 Epistomal-labral plate
states: absent; present
20 Formation of the ventroposterior wall of pre-oral chamber
states: formed by labium; formed by palpal coxae
21 Pedipalpal endite outgrown,

states: absent; present
22 Fused pedipalpal endites
states: absent; present
23 Leg 1 endite outgrown
states: absent; present
24 Fused leg 1 endites
states: absent; present
25 Leg 2 endite outgrown
states: absent; present
26 Fused leg 2 endites
states: absent; present
27 Palate plate
states: absent; present
28 Filtering preoral setae
states: absent; present
29 Three-branched epistomal skeleton
states: absent; present
30 Intercheliceral epipharyngeal sclerite
states: absent; present
31 Epipharyngeal sclerite
states: absent; large, projecting posteriorly
32 Metasoma
states: absent; present
33 Prosoma and opisthosoma form a single functional unit
states: absent; present
34 Metasoma length
states: pygidium; five segments; nine segments
35 Well-developed post-anal telson
states: absent; present,
36 Pygidium transformed into anal tubercle
states: absent; present,
37 Presence absence of flagellate telson
states: absent; present,
38 Vesicle and aculeus on the telson
states: absent; present
39 Specialized male postanal flagellum
states: absent; present,
40 Number_of_deutocerebral_appendage_articles,
states: more than 3; 3; 2
41 Basal suture in proximal cheliceral segment
states: absent; present
42 Cheliceral_PD_axis_with_point_of_inflection
states: absent; present
43 Cheliceral teeth

states: absent; present
44 Distal-most cheliceral tooth
states: 1; 2
45 Position of the cheliceral apotele articulation
states: articulates ventrally; articulates dorsally; articulates laterally
46 Distal chelicera type
states: chelate 'clasp-knife' 'type'; Prostigmata styliform or *Anystys*-like chelicerae,
47 Diaphanous cheliceral teeth
states: absent; present
48 Angle of cheliceral articulation
states: parallel to sagittal plane; oblique to sagittal plane; parallel to transverse plane; parallel to coronal plane
49 Chelicerae project beyond anterior prosomal shield margin
states: absent; present
50 Naked cheliceral fang
states: absent; present
51 Plagula ventralis
states: absent; present
52 Cheliceral venom gland;
states: absent; present
53 Endocephalic spinning apparatus
states: absent; present
54 Cheliceral flagellum
states: absent; present
55 Chelicero-carapacal articulation
states: absent; present
56 Mesal fusion of chelicerae
states: absent; chelicerae proximally fused
57 Shape of movable digit of chelicerae
states: *Anystis* type (Acarifomes); styliform,
58 Cheliceral serrula
states: absent; present
59 Morphology of the cheliceral serrula
states: rounded; tooth-like; hyaline
60 Presence/absence cheliceral brush
states: absent; present
61 Fusion of palpal coxae
states: free; fused medially
62 Gnathosoma
states: absent; present
63 Subcapitular rutella
states: absent; present
64 Morphology of palpal chelae

states: leg like; subraptorial chelate; scorpionoid
65 Apophyses on patella and tibia of pedipalp
states: absent; present
66 Modified patellar apophysis of male pedipalp
states: absent; present
67 Plane of motion of pedipalps
states: vertical; horizontal
68 Prominent dorsal flange on the pedipalp trochanter
states: absent; present
69 Dorsal row of femoral spines
states: absent; present
70 Presence/absence ventral apophysis on palpal trochanter
states: absent; present
71 Spine-like ventral apophysis on palpal trochanter
states: absent; present
72 Palpal cleaning organ
states: absent; present
73 Pedipalpal venom glands
states: absent; present
74 Fusion level of palpal apotele
states: differentiated from tarsus; not differentiated from tarsus,
75 Adhesive palpal organ
states: absent; present
76 Dorsal row of spines on pedipalp patella
states: absent; present
77 Three principal spines on pedipalp patella with spine 1 large
states: absent; present
78 Patellar spines decreasing in length proximally
states: absent; present
79 Reduced proximal spine from three-spined palpal patella'
states: absent; present
80 Pedipalpal megaspines forming a distal catching basket
states: absent; present (phrynicid hand)
81 Spines on palpal tarsus
states: absent; present
82 Fusion of palpal tibia and tarsus
states: unfused; fused
83 Ovigers
states: absent present,
84 Gnathobases
states: present absent,
85 Elongated (antenniform) first leg
states: absent present,
86 Leg 1 sternocoxal articulation

states: absent present,
87 Leg 1 apotele
states: present; absent
88 Segmentation of leg 1 tibia
states: unmodified; 16 segments ;up to 23 segments; 25 segments; more than 25 segments
89 Elongation of leg 2
states: unmodified; elongate
90 Enlargement of coxa 2
states: unmodified; enlarged
91 Presence of exopods
states: retained on more than one prosomal limb; on sixth prosomal limb; absent,
92 Morphology of coxo-trochanteral joint
states: simple; complex
93 Femoral division in legs 3 and 4
states: present; absent
94 Types of femoropatella articulation
states: transverse hinge; bicondylar articulation; monocondylar articulation,
95 Type of patellotibial articulation
states: monocondylar; hinge bicondylar
96 Auxiliary posterior articulation on patellatibial articulation
states: absent; present
97 Appendages of post-oral somites III–V with fused tibia and tarsus
states: absent; present
98 Division of tarsus
states: tarsus divided into basi- and telotarsus; tarsus undivided
99 Metatarsus at least ca. 1.5 times tarsus length
states: absent; present,
100 Telotarsi of walking legs 2–4 with three tarsomeres
states: absent; present
101 Apotele morphology
states: a simple cone or blade; a medial piece, comprising a claw or pulvillus; more frequently bearing a pair of lateral claws
102 Pulvillus
states: absent; present
103 Chelate legs
states: absent; present
104 Divided claws
states: absent; present
105 Single tarsal claws on legs 1 and 2 and double tarsal claws on legs 3 and 4
states: absent; present
106 Plate-like opisthosomal appendages
states: absent; present
107 Limb VI

states: unmodified; pusher; paddle
108 Number of body segments
states: twenty; nineteen; eighteen; seventeen; sixteen; fifteen; fourteen; thirteen; twelve; eight segments or less
109 Width of prosoma-opisthosoma junction
states: broad xiphosuran cephalothorax; narrow (pedicel)
110 Length of first opisthosomal tergite
states: unmodified; very short
111 Six abbreviated opisthosomal tergites
states: absent; present
112 Opisthosomal sternite 1
states: present; absent
113 Triangular opisthosomal sternite 1
states: present; absent
114 Locking mechanism between opisthosoma and prosoma
states: absent; present
115 Fusion of opisthosomal tergites 7-10
states: separate; fused into a single plate
116 Opisthosomal tergites with a distinct axial region (cf. trilobite)
states: absent; present
117 Division on posterior opisthosomal tergites
states: absent; present
118 Fused opisthosomal tergites
states: absent; present
119 Fused opisthosomal tergites medially divided
states: absent; present
120 Number of diplotergites
states: absent; one; more than one
121 Fused opisthosomal segments
states: absent; buckler thoracatron,
122 Appendages on opisthosomal segment 1
states: present; absent
123 Median abdominal appendage
states: absent; present
124 Genital operculum formed by the fusion of the two anterior-most abdominal appendages
states: absent; present
125 Appendages on opisthosomal segments at any developmental stage
states: absent; present
126 Overlap between the genital operculum and the sternite of third opisthosomal segment
states: absent; present
127 Opisthosomal silk glands and spigots
states: absent; present

128 Opisthosomal spinnerets
states: absent; present
129 Pygidial defensive secretions
states: absent; present
130 Genital acetabula
states: absent present
131 Number of genital acetabula
states: three; two; more than three
132 Presence/absence dorsal anal operculum
states: absent; present
133 Lateral eyes
states: absent; present
134 Morphology of lateral eye lenses
states: compound; five or more pairs of lenses; three primary pairs [excluding any microlenses]; two pairs; one pair; no lenses
135 Arrangement of lateral eye rhabdomeres in cross-section
states: net-like; star-shaped
136 Number of median eyes
states: four; two or three; absent
137 Organization of retinula cells of median eyes
states: organized into closed rhabdomes; organized into a network of rhabdomeres; disorganized inverse retina
138 Slit sense organs
states: absent; present
139 Trichobothria
states: absent; present
140 Tibial trichobothria with 2-1-1-1 distribution
states: absent; present
141 Pro-dorsal trichobothria
states: absent; present,
142 Pectines
states: absent; present
143 Malleoli
states: absent; present
144 Tarsal organ on leg I
states: absent; present
145 Tarsal organ on leg II
states: absent; present
146 Opisthosomal ganglia in adults
states: absent; present
147 Perineural membrane enveloping arterial sinus
states: present; absent
148 Intercheliceral median organ
states: absent; present

149 Filamentous gills
states: absent; present
150 Lamellate breathing structures; book gills or lungs.
states: absent; present
151 Tubular tracheae
states: absent; present
152 Book lung/gill on 2nd (i.e. genital) opisthosomal segment
states: present; absent
153 Book lung/gill on 3rd (i.e. postgenital) opisthosomal segment
states: present; absent,
154 Book lung/gill on 4th to 7th opisthosomal segment
states: present absent,
155 Stigmata
states: absent; present
156 Spines on book lung lamellar margins
states: absent; present
157 Shape of pillars of the haemolymph spaces inside the gill/lung lamellae'
states: at least two perikarya meeting midway in the haemolymph space; pillars including a strong axis of microtubules
158 Location of prosomal spiracles
states: absent; between the coxae of the second and third walking legs; associated with coxae of third and fourth walking leg; between cheliceral basis; brachypyline oribatid tracheal system
159 Location of opisthosomal spiracles
states: absent; paired ventral stigmata on genital segment; paired ventral stigmata on 3rd and 4th opisthosomal segments; four pairs of dorsal stigmata on the anterior opisthosoma
160 Kiemenplatten
states: absent; present
161 Postcerebral crop and proventriculus
states: reduced; present
162 Well-developed sucking stomach
states: absent; present
163 Endosternite
states: absent; present
164 Termination of anterior endosternal horn
states: terminating in muscular attachment to labrum; terminating in muscular attachment to palpal coxa
165 Fenestrate endosternite
states: absent; present
166 Malpighian tubules
states: absent; present
167 Coxal glands opening on proximal podomere of chelifore
states: absent; present

168 Presence/absenece of sacculus associated to coxal gland
states: absent; present
169 Coxal gland opening at the base of pedipalp,
states: absent; present
170 Coxal glands opening at base of leg 1
states: absent; present
171 Coxal glands opening on leg 3 segment
states: absent; present
172 Contribution of the coxal gland to saliva
states: absent; Buxtons group II coxal gland; coxal glands and saliva converging into the pre-oral chamber through external taenidia or gutters; podocephalic channel
173 Dorsomedian excretory organ
states: absent; present
174 Origin of lateral extrinsic precerebral pharyngeal muscle
states: arising from anterior endosternal horns; arising from medial surface of palpal coxae
175 Ventral extrinsic precerebral pharyngeal muscle and tergopharyngeal muscle of precerebral pharynx
states: absent; present
176 cheliceral tergal-deutomerite muscle
states: absent; present
177 Number of heads of lateral tergocheliceral muscle
states: one head; three heads
178 Paired muscle arising from posterior margin of anterior carapacal doublure and inserting on prosomal shield
states: absent; present
179 Posterior oblique muscles of box-truss axial muscle system (BTAMS) of postoral somites I–V
states: absent; present in one or more somites
180 Anterior oblique muscles of BTAMS posterior to postoral somite VI
states: absent; present
181 Intercoxal endosternal extensor muscles
states: absent; present
182 Endosternal dorsal suspensors of somites I and II
states: absent; present
183 Endosternal dorsal suspensor muscles in somite four with anterolateral carapacal insertion
states: absent; present
184 Endosternal dorsal suspensor muscle of somite five
states: absent; present
185 Attachment of ventral endosternal suspensor muscles
states: attaching primarily to sternum; attaching primarily to coxa of appendage of anteriorly adjacent somite

186 Palpal posteromedial tergocoxal muscle
states: present absent
187 Origin of palpal posteromedial endosternocoxal muscle
states: originates on endosternite, inserts on coxa; originates and inserts on coxa
188 Intracoxal muscle
states: absent; present
189 Size of insertion process of anteromedial tergocoxal muscle
states: weakly developed; large, well developed
190 Symmetry of femoropatellar flexor
states: symmetrical; asymmetrical
191 Insertion of pedal anterior femur–patella muscle
states: inserting primarily on patellar margin; inserting primarily on patellar plagula
192 Pedal posterior femeropatella-tibia muscle
states: present; absent
193 Pedal patellotibia-tarsus muscle
states: present; absent
194 Posterior transpatellar muscles insertion
states: 'dorsoposterior femur/posterior patella'; distal process of femur absent
195 Atellotibial extensor
states: absent; present
196 Position of the insertion of anterior transpatellar muscle
states: anterior/anteroventral; ventral/posteroventral; absent
197 Position of the anterior patellotibial muscle on tibia
states: anterior; ventral; absent
198 Posterior patellotibial muscle
states: absent; present
199 Origin of apotele depressor
states: tarsus; tibia
200 Patellar head of apotele depressor
states: absent; present
201 Origin on posterior patellar wall of the patellar head of apotele depressor
states: absent; present
202 Position of the attachments of opisthosomal posterior oblique axial muscles
states: tergal; pleural
203 Organization of opisthosomal pleural muscle
states: continuous; dorsoventral sheet divided into three components
204 Dorsal and ventral longitudinal muscles
states: spanning full length of opisthosoma; spanning first and last four opisthosomal somites
205 Internal fertilization
states: absent; present
206 Gonopore position
states: on limb bases; on second opisthosomal segment,

207 Gonopores on second opisthosomal segment
states: paired; unpaired
208 Anteriorly positioned gonopore
states: absent; present
209 Intromittent organ associated with genital region
states: absent; present
210 Penis/spermatopositor
states: spermatopositor; penis; acariforme aedagus
211 Male palpal organ
states: absent; present
212 Sperm transfer organ on leg 3
states: absent; present
213 Ovipositor
States: absent; present
214 Stalked spermatophore
states: absent; present
215 Spermatophore uptake
states: without mating; face-to-face uptake; mating parade,
216 Testis
states: glandular area unspecialised; glandular area distinctly larger
217 Tubular genital accessory glands
states: absent; present
218 Brood sac
states: absent; present
219 Sperm: Flagellum in the mature sperm
states: absent; present
220 Sperm: Coiled sperm cells
states: absent; present
221 Sperm: Acrosomal filament
states: absent; present
222 Sperm: Coiled acrosomal filament
states: absent; present
223 Sperm: Acrosomal filament piercing nucleus
ates: present; absent
224 Sperm: Microtubule arrangement in axoneme
states: 9+0; 9+1; 9+2; 9+3
225 Sperm: Helical nucleus
states: absent; present
226 Sperm: Shape of postcentriolar nucleus
states: unmodified; asymmetrical; elongate
227 Sperm: Depth of fossa implantation
states: shallow; deep
228 Sperm: Manchette of microtubules
states: absent; present

229 Sperm: Nuclear envelope
states: absent; present
230 Sperm: Persisting flagellar tunnel
states: absent; present
231 Sperm: Vacuolated sperm cells
states: absent; present
232 Yolk distribution of eggs
states: centrolecithal; isolecithal; telolecithal
233 Anatomical fate of embryonic growth zone
states: gives rise to prosoma and opisthosoma; gives rise to opisthosoma only
234 Hexapodal first instar
states: absent; present
235 Egg teeth on pedipalpal coxae
states: absent; present
236 Embryonic lateral organs
states: absent; present
237 Heteromorphic parasitic larvae
states: absent; present
238 Modified dispersal larval stage (hypopus)
states: absent; present
239 Anamorphic development with protonymphal stage
states: absent; present
240 Male oviger segments
states: 11; 10; 9; 7; 6; 4
241 Female oviger
states: absent; present
242 Terminal claw of oviger
states: absent; present
243 Auxiliary claws of propodus
states: absent; present
244 Cement glands
states: absent; present on femora and other podomeres
245 Abdominal rudiment
states: horizontal, parallel to trunk; erect diagonally or dorsally
246 Shape of proboscis
states: cylindrical; inflated proximally, acute distally; inflated distally; tapering or pipette-like
247 Proboscis position
states: frontal and fixed; in angle and movable; ventral and movable
248 Gonostome
states: closed; open
249 Ocularium
states: absent; present
250 Oviposition and copulatory ducts separate

states: absent; separated
251 Segmentation of anterior lateral spinnerets
states: indistinct; segments clearly defined
252 Segmentation of posterior lateral spinnerets
states: indistinct; up to four distinct segments; more than four distinct segments
253 Pedipalpal trichobothria type
states: Type A; Type B; Type C; Type D
254 Egg laying
states: absent; present
255 Vivaparous embryogenesis
states: apoikogenic; katoikogenic
256 Hemolymph vascular system
states: absent; present
257 Perineural vascular sheath
states: absent; present
258 R-cell terminals from lateral eyes and median eyes overlapping
states: absent; present
259 Arcuate body located apart from neuropils
states: absent; present

**File S2.** Morphological character matrix in Nexus format.

#NEXUS

BEGIN TAXA;
TITLE Taxa;
DIMENSIONS NTAX=517;
TAXLABELS
Haliestes Palaeopantopus Palaeoisopus Flagellopantopus Dibasterium Offacolus Weinbergina Attercopus_fimbriunguis Permarachne_novokshonovi Chimerarachne_yingi Idmonarachne Parastylonurus Mixopterus Eurypterus Chasmataspis Octoberaspis Waeringoscorpio Pulmonoscorpius Palaeoscorpius Compsoscorpius Proscorpius Palaeocharinus Anthracomartus Eophrynus Plesiosiro Graeophonus Paracharonopsis Kronocharon Eophalangium Hastocularis Pycno 'Tanystylum_orbiculare_P02' Anoplo 'Phoxichilidium_tubilariae_P07' 'Nymphon_molleri_P06' 'Meridionale_flava_P04' 'Stylopallene_cheilorynchus_P05' 'Labidostommatidae_IZ-67347' 'Dinothrombium_tinctorium' 'Panonychus_citri_SRR341928' 'Tetranychus_cinnabarinus_SRR519097' 'Tetranychus_urticae' 'Rhizoglyphus_robini_SRR946953' 'Sarcoptes_scabei' 'Dermatophagoides_farinae_SRR1016494' 'Dermatophagoides_pteronyssinus' 'Euroglyphus_maynei' 'Hypochithonius_rufulus_SRR4039020' 'Steganacarus_magnus_SRR4039023' 'Achipteria_coleoptrata_SRR4039018' 'Alaskozetes_antarcticus' 'Hermannia_gibba_SRR4039019' 'Archegozetes_longisetosus_MCZ' 'Nothrus_palustris_SRR4039021' 'Platynothrus_peltifer_SRR4039022' 'Pseudotyrannochthonius_B_144011' 'Pseudotyrannochthonius_MCZ' 'Chthoniidae_JC022' 'Lagynochthonius_australicus_MCZ' 'Ephippiochthonius_tetrachelatus_MCZ_140293' 'Lechytia_hoffi_MCZ139248' 'Feaella_capensis_139240' 'Pseudogarypus_banksi_140301' 'Bochica_MCZ144332' 'Dhanus_sumatranus_MCZ49986' 'Indohya_Australia_151267' 'Ideobisium_crassimanum_MCZ' 'Parahya_submersa_139241' 'Gymnobisium_139238' Microcreaginae 'Microbisium_brunneum_MCZ' 'Microbisium_60599' 'Novobisium_141855' 'Afrogarypus_MCZ139239' 'Geogarypus_maculatus_MCZ' 'Apolpium_MCZ144360' 'Garypus_californicus_MCZ' Synsphyronus 'Pseudogarypinus_frontalis_MCZ139246' 'Larca_granulata_MCZ' 'Protogarypinus_giganteus' 'Cheiridiidae_MCZ140294' 'Sternophoridae_133496' 'Cacodemonius_MCZ144363' 'Oratemnus_curtus_49983' 'Protochelifer_sp_MCZ' 'Cheliferidae_JC031_MCZ40982' 'Parachelifer_persimilis_MCZ139247' 'Lamprochernes_savignyi_MCZ' 'Conicochernes_crassus_MCZ49985' 'Hesperochernes_sp' 'Haplochernes_kraepeli_reassaembly' 'Chernetidae_MCZ142323' 'Chernetidae_sp_JC032' 'Ehan_MCZ74291' 'Espe_MCZ60890' 'Espelaea_UWisc' 'Galendromus_occidentalis' 'Gromphadorholaelaps_schaeferi_MCZ71202' 'Varroa_destructor_SRR3927486' 'Pneumolaelaps_niutirani_SRR10269756' 'Tropilaelaps_mercedesae' 'Adenacarus_sp' 'Neoacarus_sp_MCZ'

'Orthinodoros_rostratus_SRR1732011' 'IscaW_1_6' 'Haemaphysalis_longicornis_SRR7754709' 'Amblyomma_americanum' 'Dermacentor_silvarum' 'Hyalomma_excavatum_SRR3157672' 'Rhipicephalus_micropilus_SRR1186998' 'Siro_boyerae_SRR1145699' 'Leptopsalis_141287' 'Miopsalis_SRR5234539_peptides_iso' 'Metagovea_oviformis_MCZ136533' 'Metasiro_savannahensis_IZ-133799' 'Brasilogovea_microphaga_MCZ136559' 'Neogovea_matawai_MCZ133922' 'Parapurcellia_silvicola_MCZ134743' 'Purcellia_illustrans_MCZ49518' 'Chileogovea_oedipus_MCZ138106' 'Rakaia_minutissima_MCZ143240' 'Rakaia_stewartiensis_MCZ49801' 'Rakaia_magna_australis_MCZ29212' 'Rakaia_pauli_MCZ144621' 'Rakaia_pauli_Trini_CDhit' 'Pettalus_thwaitsei_MCZ132349' 'Austropurcellia_alata_MCZ141418' 'Austropurcellia_despectata_MCZ141417' 'Austropurcellia_cf_acuta_MCZ141419' 'Austropurcellia_acuta_MCZ44060' 'Austropurcellia_clousei_MCZ141420' 'Neopurcellia_salmoni_MCZ144625' 'Neopurcellia_salmoni_MCZ29310' 'Karripurcellia_peckorum_MCZ49989' 'Aoraki_141173' 'Aoraki_denticulata_Northern_MCZ141173' 'Aoraki_denticulata_Southern_MCZ141172' 'Aoraki_longitarsa_MCZ137238' 'Acropsopilio_neozealandiae_MCZ30457' 'Hesperonemastoma_modestum' 'Ortholasma_coronadense' 'Trogulus_martensi' 'Caddo_agilis' 'thrasychirus_49762' 'Forsteropsalis_pureora_MCZ29216' 'Pantopsalis_nz2016' 'mangatangi_133381' 'megalopsalis_133420' 'Diguetinus_143123' 'Phalangium_opilio_IZ-29189' 'Protolophus_singularis' 'prionostemma_144108' 'Leiobunum_verrucosum' 'paranelima_143124' 'Sclerobunus_robustus' 'Synthetonychia_glacialis_MCZ137212' 'Larifuga_capensis_IZ-49747' 'Nuncia_nz1211' Pristobunus 'Scotolemon_lespesi_MCZ43672' 'Sitalcina_lobata' 'Gnomulus_141286' 'Dibunus_141285' 'Epedanus_141288' 'Haasus_judaeus' 'Haasus_naasane' Dampetrus 'Metabiantes_sp_IZ-49748' 'Stygnomma_144060' pellobunus 'Kimulidae_143960' 'Tegipiolus_pachypus_MZSP71018' 'pellobunus_144055' 'Pachylicus_acutus_IZ-49749' 'maracaynatum_144047' Avima Karos chapolobunus 'Stygnoplus_143966' Protimesius 'Auranus_hoeferscovitorum_MCZ136531' 'Pickeliana_pickeli_MCZ139249' 'Gonycranaus_pluto_MCZ139251' 'Camarana_sp_MCZ139271' Pseudopachylus 'Bissula_spnov_MCZ139255' 'Pseudopachylus_sp_MCZ139280' 'Vonones_ornata_IZ-49750' 'Incasarcus_argenteus_MCZ139250' 'Quindina_limbata_MZSP71772' Saramacia 'Ampycinae_sp_MCZ46455' 'Glysterus_sp_MCZ140069' 'Panalus_robustus_MZSP71019' 'Phalangodus_cottus_MZSP71020' 'Santinezia_144078' 'phareicranaus_136554' 'Pseudopucrolia_discrepans_MCZ139277' 'Sadocus_polyacanthus_MCZ138113' 'Chilegyndes_phillipsoni_MCZ49760' metagyndes 'Hernandaria_una_MCZ139258' 'Graphinotus_therezopolis_MCZ139273' 'Meteusarcoides_139264' 'Meteusarcoides_sp_MZSP71024' 'Acutisoma_longipes_MCZ139261' 'Goniosoma_sp_MCZ139266' 'Roeweria_virescens_MCZ139279' 'Longiperna_sp_MCZ139260' iz139272 'Mitobates_sp_MCZ139263' 'Promitobates_ornatus_MZSP71022' 'Gonyleptellus_sp_MCZ139269' 'Gonyleptes_fragilis_MZSP71023' 'Pseudotrogulus_mirim_MCZ139270' 'Pseudotrogulus_sp_MCZ139278'

'Ampheres_leucopheus_MCZ139267'
'Acanthogonyleptes_aff_fulvigranulatus_MZSP71027' 'Gonyleptes_atrus_MZSP71025'
'Gonyleptes_sp_MCZ139262' 'Geraecormobius_bispinifrons_MCZ139274'
'Mischonyx_cuspidatus_MCZ139265' 'Sodreana_leprevosti_MCZ139256'
'Sodreana_sodrena_MCZ139268' 'Progonyleptoidellus_striatus_MZSP71021'
'Iporangaia_pustulosa_MCZ139275' 'Gonyleptoides_marumbiensis_MCZ139254'
'Neosadocus_sp_MCZ139257' 'Galeodes_sp' 'Solpugema_sp_IZ-49525'
'Eremobates_sp_IZ-49755' 'Eremobates_sp_new' Lpol Crot
'Tachypleus_tridentatus_2019G' Tgig 'Ricinoides_atewa' 'Ricinoides_karschii_IZ-130083' 'Pseudocellus_pearsei' 'Pseudocellus_sp_IZ-140060' 'Cryptocellus_sp_IZ-143922' 'Cryptocellus_becki_IZ-136532' 'Cryptocellus_sp_IZ-30913' ChaerilusSS
TroglokhammouanusSS Vietbocap Lvar 'Mesobuthus_martensii' Cley BiisraFull
CompsoI 'Csch_trinity' Aamox 'Androctonus_australisSS' HottentottaSS 'Oscr_trinity'
AcraII 'Bisra_trinity' Bare Lheb LquiE Abal Bgig Grosphus ParabuthusSS Uoli Uvit Tsmi
Tarc Tcos Tserrulatus Tlac Pdeb Jaga Jroc Hjun 'Rgar_trinity' Ccar Chentzi
'Centruroides_vittatus' CscuSS Iurus Bcor 'Bothriurus_burmeisteriSS' Centro Cerc Cque
Csul Ckeyserlingii SuperSS AnuroctonusSS Brotheas ScorpiopsSS EuscorpiusSS
Pdilutus 'Mega_gertschi' MegacormusSS Umordax BelisariusSS Haztecus HadrSS
Hconcolorous Hspadix Uhuachuca Kboggerti Papacheanus Vcashi Pbae Smes
Svachoni Vmexicanus Sgertschi Kacapulco Kchamela Mvariegatus Pschwenkmeyeri
Pspinigerus Kpurepecha Ccoahuilae Moccidentalis Tintrepidus Uyaschenkoi
'Uro_elongatus' Urodacus 'Hormiops_davidovi' 'Hormurus_weberi' Liocheles
'Liocheles_sp_KhaoDeng' 'Nebo_hierochonticus' DiplocentrusSS
'Diplocentrus_zacatecanus' Kolotl 'Kolotl_poncei' 'Iomachus_politus' HadogenesSS
'Hadogenes_paucidens' 'Opistacanthus_asper' 'Opistacanthus_validus'
'Chiromachus_ochropus' OpisthacanthusSS 'Palaeocheloctonus_pauliani'
'Opistophthalmus_glabrifrons' PandinusSS 'Opistophthalmus_walhbergi'
'Opistophthalmus_boehmi' Sfuscus 'Cioa_Trinity' 'Cisr_Trinity' 'Dvar_IZ29740'
'Acor_IZ71252' 'Pmar_Trinity' 'Mastigoproctus_giganteus_new' 'Mlac_IZ71253'
'Draculoides_vinei_IZ-46896' 'Stenochrus_portoricensis_MCZ' 'Stenochrus_sp_IZ-74289' Liphistius 'Liphistius_sp2' 'Sphodros_rufipes' 'Megahexura_fulva'
'Aliatypus_coylei' 'Antrodiaetus_unicolor' 'Porrhothele_sp' 'Microhexura_montivaga'
'Macrothele_calpeiana' 'Paratropis_sp' 'Cyclocosmia_truncata' 'Hebestatis_theveneti'
'Idiops_bersebaensis' 'Aptostichus_stephencolberti' 'Promyrmekiaphila_clathrata'
'Brachythele_longitarsis' 'Pionothele_sp' 'Stanwellia_sp' 'Damarchus_sp'
'Trichopelma_laselva' 'Acanthoscurria_geniculata' 'A_hentzi_pooled'
'Aphonopelma_iviei' 'Filistata_insidiatrix' Kukulcaniahibernalis2 'Hypochilus_gertschi'
'Hypochilus_pococki' 'Calponia_harrisonfordi' 'Segestria_sp' 'Maoriata_sp'
'Orsolobidae_sp' Dysdera 'Ischnothyreus_sp' 'Opopaea_sp' 'Diguetia_sp'
'Pholcus_manueli' 'Pholcus_phalangioides' 'Loxosceles_deserta' 'Drymusa_sp'
'Periegops_suterii' 'Ochyrocera_sp' 'Scytodes_globula' 'Scytodes_thoracica'
Archoleptoneta Calileptoneta Austrochilus 'Hickmania_clean' 'Progradungula_GOOD'
'Tarlina_sp' 'Gradungula_sorenseni' 'Pianoa_isolata' 'Austrarchaea_sp'
'Mecysmauchenius_clean' 'Huttonia_palpimanoides' 'Othiotops_birabeni'

'Palpimanus_gibbulus' 'Stegodyphus_sp' 'Megadictyna_thilenii' Nicodamidae2 'Nicodamidae_sp1' 'Mysmenella_sp' 'Novanapis_sp' 'Anelosimus_clean' Latrodectus 'Euryopis_sp' 'Parasteatoda_tepidariorum' 'Pararchaea_alba' 'Malkara_clean' 'Malkaridae_GH09' 'Perissopmeros_sp' 'Australomimetus_clean' 'Ero_leonina' 'Arkys_sp' Demadiana Leucage 'Meta_ovalis_good' 'Nanometa_sp' 'Tetragnatha_tantalus' 'Cepheia_clean' 'Mysmena_leichardti' 'Microdipoena_guttata' 'Spinanapis_sp' 'Tekelloides_sp2' 'Laminacauda_clean' Frontinella 'Novafroneta_sp' 'Synotaxus_sp' 'Nesticus_bishopi' 'Nesticus_cooperi' 'Meringa_sp' 'Runga_sp' 'Physoglenes_clean' 'Physoglenidae_clean' 'Ogulnius_sp' 'Baalzebub_sp' 'Theridiosoma_gemmosum' 'Theridiosoma_savannum' 'Trichonephila_clavipes_Genome' 'Micrathena_gracilis' 'Verrucosa_arenata' 'Cyrtophora_sp' Neoscona 'Gasteracantha_hasselti' 'Macracantha_arcuata' 'Menneus_sp' 'Deinopis_clean' 'Deinopis_longipes' 'Tamopsis_sp' 'Oecobius_navus' 'Oecobius_sp' 'Zosis_sp' 'Philoponella_herediae' 'Uloborus_glomosus' 'Cybaeodamus_taim' 'Forsterella_sp' 'Caayguara_clean' Amaurobius 'Callobius_sp' 'Neoramia_sp' Amphinecta 'Metaltella_simoni' Cambridgea 'Paramatachia_sp' 'Agelenopsis_emertoni_cdhitest.fasta.transdecoder' 'Agelenopsis_pennsylvanica' 'Calymmaria_clean' 'Calymmaria_persica' 'Cicurina_travisae' 'Cicurina_vobora' 'Homalonychus_theologus' 'Tengella_sp' 'Uliodon_sp' 'Misumenoides_formosipes' 'Sidymella_sp' 'Anahita_punctulata' 'Peucetia_longipalpis' 'Dolomedes_trinity' 'Pisaurina_mira' 'Cupiennius_sp' Allocosa 'Schizocosa_rovneri' 'Habronattus_signatus' 'Habronattus_ustulatus' 'Teminius_sp' 'Karaops_raveni' 'Falconina_gracilis' 'Nyssus_sp' 'Clubiona_clean' 'Rebilus_sp' 'Liocranidae_GOOD' 'Trachelas_tranquillus' 'Sergiolus_capulatus' 'Molycria_sp' Anzacia 'Lampona_sp' Goniotarbus Bornatarbus
;

END;

BEGIN CHARACTERS;
DIMENSIONS NCHAR=259;
FORMAT DATATYPE = STANDARD RESPECTCASE GAP = - MISSING = ? SYMBOLS = "  0 1 2 3 4 5 6 7 8 9 A";

MATRIX
Haliestes 00000000-0-000-1000-??????0?0??00-0-00000---200-1??0?0?0-0-00000000000-0??00-----?110000002?0???000100000090------------1--0---?--00--??????0000??00201110-----??????????????????-------?-?????????????----??----00-????????????????????????????10?101?1?---??????
Palaeopantopus 0000?000-0-000-1200-???????????00-1-0000??--??????????????????????????????????????????????2???????????????????????------00---

???0?????????00?????0201110??????????????????????????????????????????
??????????-000????????????????????????????????????????????????

Palaeoisopus 0000?000-0-000-1200-???????????00-1-0?000---200-1????0?0-0-00000000000-0??00-----?110000002?0???0000100000800-010-----001000000???00--0?????0000??00201110----0?????????????????????????????????????????????-0000???????????????????????????0000?021?0?---??????

Flagellopantopus 0??0???0-0-0??-1200-???????????00-1-1?0-0---------???0------?000000000-0??00-----?1100?0002?0???0?00??0??00800-010-----001000000???0?--??????00?????0201110----0?????????????????????????????????????????????-0000???????????????????????????????010?0?---??????

Dibasterium 1000?000-00000-010?????????????00-1-000001---00-0????0?0-0-000?2000000-0?000-----0000?00000?1????1-00?100102??0000000--000001000???00--2-????00???000000000?--0?????????????????????????????????????????????-0??0??????????????????????????----?????0?---??????

Offacolus 1000?000-00110-0?0?????????????00-1-000001---00-0????0?0-0-000?2000000-0?000-----00?0?000?0?1????1-000100105??0000000--000001000???00--2-????00???000000000?--0?????????????????????????????????????????????-0??0??????????????????????????----?????0?---??????

Weinbergina 1101?000-1000000?0?????????????00-1-000101--200??????0?0-0-000?0000000-0?000-----0000?0000??1???01-0000001031?0000000--000001000???010???????00?????00000001?--0?????????????????????????????????????????????-0??0??????????????????????????----?????0?---??????

Attercopus_fimbriunguis
1????0?????????????????????????1?10?2?-??110??1?1?0??-0-0????????????????????????????????????????????????????????????????1?????????????????0??????0????????????????????????????????????????????????????????????????????????????????????????????????----?????0?---??????

Permarachne_novokshonovi
1????0?????0???0????????????????10?1?10????????????????????????????????????????????????0??00?2?????00?0??0???0?2?0???0??0-?010????????????????????00?????0???????????????????????????????????????????????????????????????????????????????????????????????????----?????0???-??????

Chimerarachne_yingi
100??00100000000?1?0????????????100101002?-??11?11???00?0-??000000-0000-0??00----000100000021100000000010000?0?2000-00000-001001111?0-012????1??00?????000?????????????????????????????????????????????????????????10?????????????????????????????----?????0?22-??????

Idmonarachne
100000010000?000???0???????????010000---20---11011???0000-0-00000000000-

0?000----00010000002?1???00101000010?200?0?0100-
?0100???0?????????????00?????0?????????????????????????????????????????
???????????????????-0?0???????????????????????????----?????0?---??????
Parastylonurus
1000?00101000000?00????????????1011-000101--100-00???00-0-00000000000-
0?000-----0000?000020000001-0000--1000100000000-00011?000???010?1?00--
0000??00001000??--???????????????????????????????????????????????11?0-
0000???????????????????????????----?????0?---??????
Mixopterus 1000?00101000000?00????????????1011-
000101--100-00???00-0-00001000000-0?000-----0000?000020000001-0000--
1200100000100-00011?000???010?1?00--0000??00001000??--
???????????????????????????????????????????????11?0-
0000???????????????????????????----?????0?---??????
Eurypterus 1000?00101000000?00????????????1011-
000101--100-00???00-0-00000000000-0?000-----0000?000020000001-0000--
1200100000000-000111000???010?1?00--0000??00001000??--
1??????????1???????????????????????????????????11?0-
00001??????????????????????????----?????0?---??????
Chasmataspis
1001?00???010000???????????????1021-000???--????????????-
???????????????????-----?00?????01??????1-0001001000100000100-
01???0??0???010?1?????00????????????????0????????????????????????????
???????????????????-0??0??????????????????????????----?????0?---??????
Octoberaspis
1100?00???010000???????????????1021-000???--????????????-
???????????????????-----?0?????????????????000001200100000000-
0001000?0???010?1?????00????????????????0????????????????????????????
???????????????????-0??0??????????????????????????----?????0?---??????
Waeringoscorpio
1000?0010?000000???????????????1011-010???--????????????-???0003001000-
0?100----?0010???002?1???00?0100001000000000000-
001001??0???0???1??????0?????1???????????????????????????????????????
???????????????????-0000???????????????????????????----?????0?--???????
Pulmonoscorpius
1000?00100000000??????1111?????1011-010100--200310???0?0-0-00003001000-
0?100----?0010?00002?1???0000100001000000000000-0010010?0???011?1??0--
10?????0001101??--0??????????????????????????????????????????????1?0-
0000???????????????????????????----?????0?--???????
Palaeoscorpius
1000?0010??00000???????????????101?-?1?100--2??31????0?0-???0003001000-
??100----?0010?00002?1???0000100001000000000000-
001001??0???0?????????00?????000??01???????????????????????????????????
??????????????????0-0000???????????????????????????----?????0?--???????

Compsoscorpius
1000?001001000000?????1111?????1011-010100--20031????0?0-0-00003001000-??100----?0010???002?1???00?0??0??1000000000000-0010010?0???011?1??10?00?????000?1?1??????????????????????????????????????????????????????0-0000??????????????????????????----?????0?--???????

Proscorpius 1000?0010?000000???????????????1011-010100--20031????0?0-0-00003001000-??100----?0010?00002?1???00001000010000000000000-0010010?0???010?1?????00??????????????--0???????????????????????????????????????????0-0000??????????????????????????----?????0?--???????

Palaeocharinus 1000?0010100100001100010??0101000-0----20---110000000000-0-00002000000-00000----00010000002111000000100001010001-10100-101001?00???011?1?10-?00?????0000011100200??????????????-????????????????????????????????-0000??????????????????????????----?????0?---??????

Anthracomartus 1000?00100001000????0010???????00-0----20---11000????000-0-00002000000-0?000----00010?00002?1???0000100001010001-10100-10100???0???01??1?????00?????0??????????????????????????-????????????????????????????????-0000??????????????????????????----?????00---??????

Eophrynus 1000?0010?0010-0???????????????00-0----20---11000????000-0-0000?000000-0?0?0----?0010???002?1???0000??0??1010001-10100-00100???0???00--1?????00?????0??????????????????????????-????????????????????????????????-00?0??????????????????????????----?????00---??????

Plesiosiro 1100?0011?001000??0?????????????00-0----20---11000????000-0-00000000000-0???0----?0011??0002?1???0001?00??1010100100000-00100?1?0???01??1?????00?????0?????1???????????????????????????????????????????????????????-0000??????????????????????????----?????00---??????

Graeophonus
1000?00110000100????????????????10000---2?-0011001????000-???00010000010??0010---00011???002?1???00011?0??1012000000000-00100?1?0??-01??1?????0??????0????????????????????????????????????????????????????????????????00??????????????????????????----?????00---??????

Paracharonopsis
1000?00110000100????????????????10000---20-??110010???000-1?00001000000111?001100010011??2002?10??00011100010120000000000-001001100??-

012?1??1?00000???0?????????????????????????????????????????????????
?????????00???????????????????????????----?????00---??????
Kronocharon
1000?00110000000?1?????????????10000---20-??110010???000-
1?0000100101101?101111100011???002?10??0001110001012000000000-
001001100??-
01??1??1?00000???0?????????????????????????????????????????????????
?????????00???????????????????????????----?????00---??????
Eophalangium 1000??0??1100000???????????????01-
0----101--200-00?000?0-0-0000?????????????----
??010????020??????????????040000000000-0010-000000-
0???1?????0??????0211111--
??0??????????????????????????????????????????111101??1??????????????
????????????----????110---??????
Hastocularis 1000110??00000000???1?0010?????01-0--
--101--200-0????0?0-0-0000?000000-????0----
0?010?00102?1???00001?0???040000000000-0010-00?000-
01??1?????0??????0?????????????????????????????????????????????????
???111101?0???????????????????????????----????11?---??????
Pycno 00000000-0-000-1200-000000000??---0-000--
-----------0-------0000000000-00000-----011000000200???000010000-090------------1--0--
-0--00--0300---000-000201110-----000--0-00??00-????-------?-???????????00----00----
00-0-000010--?00000?000000000001101010-10---0-00??
'Tanystylum_orbiculare_P02' 00000000-0-000-1000-000000000??-
--0-00010---2-0-10000000-0-00000000000-00000-----011000000200???000010000-
090------------1--0---0--00--0300---000-000201110-----000--0100??00-????-------?-
???????????00----00----00-0-000010--?00000?000000000100101 30-10---0-00??
Anoplo 00000000-0-000-1000-000000000??---0-
00010---200-10000000-0-00000000000-00000-----011000000200???000010000-090---
---------1--0---0--00--0300---000-000201110-----000--0100??00-????-------?-
???????????00----00----00-0-000010--?00000?000000000021000100-10---0-00??
'Phoxichilidium_tubilariae_P07' 00000000-0-000-1000-000000000??-
--0-00010---200-10000000-0-00000000000-00000-----011000000200???000010000-
090------------1--0---0--00--0300---000-000201110-----000--0100??00-????-------?-
???????????00----00----00-0-000010--?00000?000000000021000100-10---0-00??
'Nymphon_molleri_P06' 00000000-0-000-1000-000000000??---
0-00010---200-10000000-0-00000000000-00000-----011000000200???000010000-090-
-----------1--0---0--00--0300---000-000201110-----000--0100??00-????-------?-
???????????00----00----00-0-000010--?00000?000000000010110100-10---0-00??
'Meridionale_flava_P04' 00000000-0-000-1000-000000000??---
0-00010---200-10000000-0-00000000000-00000-----011000000200???000010000-090-
-----------1--0---0--00--0300---000-000201110-----000--0100??00-????-------?-
???????????00----00----00-0-000010--?00000?000000000010101011-10---0-00??

'Stylopallene_cheilorynchus_P05' 00000000-0-000-1000-000000000??---0-00010---200-10000000-0-00000000000-00000-----011000000200???000010000-090------------1--0---0--00--0300---000-000201110-----000--0100??00-????-------?-??????????00----00----00-0-000010--?00000?00000000010101011-10---0-00??
'Labidostommatidae_IZ-67347' 00000000-0-00000001-1000000000-01-0----20---000--0000000-0-01100000000-00100----00010000020001001-010000?070000000000100010-000001101401?11010000000211111--300001000000103100--0???????0??000000000010-1??1110-00011?1001-101-0--01-00110?001---------00---0-00??
'Dinothrombium_tinctorium' 00000000-0-000-0001-1000000000-01-0----20---020--000000010-01104000000-00100----00010000020001001-010000?080000000000100010-000000-01402?11010000000211111--300001000000103100--0???????0??000000000010-1??1110-00001?1001-101-0--01-001100101---------00---0-00??
'Panonychus_citri_SRR341928' 00000000-0-000-0001-1000000000-01-0----20---020--000000110-01104000000-00100----00010000020101001-010000?080000000000100010-000000-01302?10--00000000211111--300001000000103100--0???????0??000000000010-1??1110-20000?1001-101-0--01-001100001---------00---0-00??
'Tetranychus_cinnabarinus_SRR519097' 00000000-0-000-0001-1000000000-01-0----20---020--000000110-01104000000-00100----00010000020101001-010000?080000000000100010-000000-01302?10--00000000211111--300001000000103100--0???????0??000000000010-1??1110-20000?1001-101-0--01-001100001---------00---0-00??
'Tetranychus_urticae' 00000000-0-000-0001-1000000000-01-0----20---020--000000110-01104000000-00100----00010000020101001-010000?080000000000100010-000000-01302?10--00000000211111--300001000000103100--0???????0??000000000010-1??1110-20000?1001-101-0--01-001100001---------00---0-00??
'Rhizoglyphus_robini_SRR946953' 00000000-0-000-0001-1000000000-01-0----20---000--0000000-0-01100000000-00100----00010000020101001-010000?060000000000100010-000001100--2-10--00000000201110--000001001000103000--0???????0??000000000010-1??1110-20000?1001-101-0--01-001101011---------00---0-00??
'Sarcoptes_scabei' 00000000-0-000-0001-1000000000-01-0----20---000--0000000-0-01100000000-00100----00010000020101001-010000?060000000000100010-000001100--2-10--00000000201110--000001001000103000--0???????0??000000000010-1??1110-20000?1001-101-0--01-001101011---------00---0-00??
'Dermatophagoides_farinae_SRR1016494' 00000000-0-000-0001-1000000000-01-0----20---000--0000000-0-01100000000-00100----00010000020101001-010000?060000000000100010-000001100--2-10--

00000000201110--000001001000103000--0???????0??000000000010-1??1110-20000?1001-101-0--01-001101011---------00---0-00??
'Dermatophagoides_pteronyssinus' 00000000-0-000-0001-1000000000-01-0----20---000--0000000-0-01100000000-00100----00010000020101001-010000?0600000000100010-000001100--2-10--00000000201110--000001001000103000--0???????0??000000000010-1??1110-20000?1001-101-0--01-001101011---------00---0-00??
'Euroglyphus_maynei' 00000000-0-000-0001-1000000000-01-0----20---000--0000000-0-01100000000-00100----00010000020101001-010000?0600000000100010-000001100--2-10--00000000201110--000001001000103000--0???????0??000000000010-1??1110-20000?1001-101-0--01-001101011---------00---0-00??
'Hypochithonius_rufulus_SRR4039020' 00000000-0-00000001-1000000000-01-0----20---000--0000000-0-01110000000-00100----00010000020001001-010000?0600000000100010-000001001301?1101000000000201110--000001000000103000--0???????0??000000000010-1??1110-10011?1001-101-0--01-001101001---------00---0-00??
'Steganacarus_magnus_SRR4039023' 00000000-0-00000001-1000000000-01-0----20---000--0000000-0-01110000000-00100----00010000020001001-010000?0600000000100010-000001001301?1101000000000201110--000001000000103000--0???????0??000000000010-1??1110-10011?1001-101-0--01-001101001---------00---0-00??
'Achipteria_coleoptrata_SRR4039018' 00000000-0-00000001-1000000000-01-0----20---000--0000000-0-01110000000-00100----00010000020001001-010000?0600000000100010-000001001301?1101000000000201110--000001000000103000--0???????0??000000000010-1??1110-10011?1001-101-0--01-001101001---------00---0-00??
'Alaskozetes_antarcticus' 00000000-0-00000001-1000000000-01-0----20---000--0000000-0-01110000000-00100----00010000020001001-010000?0600000000100010-000001001301?1101000000000201110--000001000000103000--0???????0??000000000010-1??1110-10011?1001-101-0--01-001101001---------00---0-00??
'Hermannia_gibba_SRR4039019' 00000000-0-00000001-1000000000-01-0----20---000--0000000-0-01110000000-00100----00010000020001001-010000?0600000000100010-000001001301?1101000000000201110--000001000000103000--0???????0??000000000010-1??1110-10011?1001-101-0--01-001101001---------00---0-00??
'Archegozetes_longisetosus_MCZ' 00000000-0-00000001-1000000000-01-0----20---000--0000000-0-01110000000-00100----00010000020001001-010000?0600000000100010-

000001001301?1101000000000201110--000001000000103000--
0???????0??000000000010-1??1110-10011?1001-101-0--01-001101001---------00---0-
00??
'Nothrus_palustris_SRR4039021' 00000000-0-00000001-
1000000000-01-0----20---000--0000000-0-01110000000-00100----
0001000000020001001-010000?0600000000000100010-
000001001301?1101000000000201110--000001000000103000--
0???????0??000000000010-1??1110-10011?1001-101-0--01-001101001---------00---0-
00??
'Platynothrus_peltifer_SRR4039022' 00000000-0-00000001-
1000000000-01-0----20---000--0000000-0-01110000000-00100----
0001000000020001001-010000?0600000000000100010-
000001001301?1101000000000201110--000001000000103000--
0???????0??000000000010-1??1110-10011?1001-101-0--01-001101001---------00---0-
00??
'Pseudotyrannochthonius_B_144011' 10000000-00000-0001-
1100000000-00-0----20---000010001100-0-00003001000-00100----
0001000000020112002?011000?010000000000-0010-000000-01302-
110000001000211111--020001000000010000--0??011000000000011121110100111 0-
00001000100111210000101100000 1---------00---0-0001
'Pseudotyrannochthonius_MCZ' 10000000-00000-0001-
1100000000-00-0----20---000010001100-0-00003001000-00100----
0001000000020112002?011000?010000000000-0010-000000-01302-
110000001000211111--020001000000010000--0??011000000000011121110100111 0-
00001000100111210000101100000 1---------00---0-0001
'Chthoniidae_JC022' 10000000-00000-0001-1100000000-00-
0----20---000010001100-0-00003001000-00100----
0001000000020112002?011000?010000000000-0010-000000-01302-
110000001000211111--020001000000010000--0??011000000000011121110100111 0-
00001000100111210000101100000 1---------00---0-0001
'Lagynochthonius_australicus_MCZ' 10000000-00000-0001-
1100000000-00-0----20---000010001100-0-00003001000-00100----
0001000000020112002?011000?010000000000-0010-000000-01302-
110000001000211111--020001000000010000--0??011000000000011121110100111 0-
00001000100111210000101100000 1---------00---0-0001
'Ephippiochthonius_tetrachelatus_MCZ_140293' 10000000-00000-0001-
1100000000-00-0----20---000010001100-0-00003001000-00100----
0001000000020112002?011000?010000000000-0010-000000-01302-
110000001000211111--020001000000010000--0??011000000000011121110100111 0-
00001000100111210000101100000 1---------00---0-0001
'Lechytia_hoffi_MCZ139248' 10000000-00000-0001-1100000000-
00-0----20---000010001100-0-00003001000-00100----
0001000000020112002?011000?010000000000-0010-000000-01302-

110000001000211111--020001000000010000--0??011000000000011211101001110-
000010001001112100001011000001---------00---0-0001
'Feaella_capensis_139240' 10000000-00010-0001-1100000000-00-0----20---000010001100-0-00003001000-00100----00010000020112001-011000?010000000000-0010-000000-01302-110000001000211111--020001000000010000--0??011000000000011211101001110-00001?001001112100001011000001---------00---0-0001
'Pseudogarypus_banksi_140301' 10000000-00010-0001-1100000000-00-0----20---000010001100-0-00003001000-00100----00010000020112001-011000?010000000000-0010-000000-01302-110000001000211111--020001000000010000--0??011000000000011211101001110-000010001001112100001011000001---------00---0-0001
'Bochica_MCZ144332' 10000000-00000-0001-1100000000-00-0----20---000010001110-0-00003001000-01100----000100000201120000011000?010000000000-0010-000000-01302-110000001000211111--020001000000010000--0??011000000000011211101001110-00001?001001112100001011000001---------00---0-0001
'Dhanus_sumatranus_MCZ49986' 10000000-00000-0001-1100000000-00-0----20---000010001110-0-00003001000-01100----000100000201120000011000?010000000000-0010-000000-01302-110000001000211111--020001000000010000--0??011000000000011211101001110-00001?001001112100001011000001---------00---0-0001
'Indohya_Australia_151267' 10000000-00000-0001-1100000000-00-0----20---000010001110-0-00003001000-01100----000100000201120000011000?010000000000-0010-000000-01302-110000001000211111--020001000000010000--0??011000000000011211101001110-00001?001001112100001011000001---------00---0-0001
'Ideobisium_crassimanum_MCZ' 10000000-00000-0001-1100000000-00-0----20---000010001110-0-00003001000-01100----000100000201120000011000?010000000000-0010-000000-01302-110000001000211111--020001000000010000--0??011000000000011211101001110-00001?001001112100001011000001---------00---0-0001
'Parahya_submersa_139241' 10000000-00000-0001-1100000000-00-0----20---000010001110-0-00003001000-01100----000100000201120000011000?010000000000-0010-000000-01302-110000001000211111--020001000000010000--0??011000000000011211101001110-00001?001001112100001011000001---------00---0-0001
'Gymnobisium_139238' 10000000-00000-0001-1100000000-00-0----20---000010001110-0-00003001000-01100----000100000201120000011000?010000000000-0010-000000-01302-110000001000211111--020001000000010000--0??011000000000011211101001110-00001?001001112100001011000001---------00---0-0001
Microcreaginae 10000000-00000-0001-1100000000-00-0----20---000010001110-0-00003001000-01100----

0001000000201120000011000?010000000000-0010-000000-01302-
1100000001000211111--020001000000010000--0??0110000000000111211101001110-
00001000100111210000101100000 1---------00---0-0001
'Microbisium_brunneum_MCZ' 10000000-00000-0001-
1100000000-00-0----20---000010001110-0-00003001000-01100----
0001000000201120000011000?010000000000-0010-000000-01302-
1100000001000211111--020001000000010000--0??0110000000000111211101001110-
000010001001112100001011000001---------00---0-0001
'Microbisium_60599' 10000000-00000-0001-1100000000-00-
0----20---000010001110-0-00003001000-01100----
0001000000201120000011000?010000000000-0010-000000-01302-
1100000001000211111--020001000000010000--0??0110000000000111211101001110-
000010001001112100001011000001---------00---0-0001
'Novobisium_141855' 10000000-00000-0001-1100000000-00-
0----20---000010001110-0-00003001000-01100----
0001000000201120000011000?010000000000-0010-000000-01302-
1100000001000211111--020001000000010000--0??0110000000000111211101001110-
000010001001112100001011000001---------00---0-0001
'Afrogarypus_MCZ139239' 10000000-00000-0001-1100000000-
00-0----20---000010001110-0-00003001000-01100----
0001000000201120000011000?010000000000-0010-000000-01302-
1100000001000211111--020001000000010000--0??0110000000000111211101001110-
00001?001001112100001011000001---------00---0-0001
'Geogarypus_maculatus_MCZ' 10000000-00000-0001-
1100000000-00-0----20---000010001110-0-00003001000-01100----
0001000000201120000011000?010000000000-0010-000000-01302-
1100000001000211111--020001000000010000--0??0110000000000111211101001110-
00001?001001112100001011000001---------00---0-0001
'Apolpium_MCZ144360' 10000000-00000-0001-1100000000-
00-0----20---000010001110-0-00003001000-01100----
0001000000201120000011000?010000000000-0010-000000-01302-
1100000001000211111--020001000000010000--0??0110000000000111211101001110-
00001?001001112100001011000001---------00---0-0001
'Garypus_californicus_MCZ' 10000000-00000-0001-1100000000-
00-0----20---000010001110-0-00003001000-01100----
0001000000201120000011000?010000000000-0010-000000-01302-
1100000001000211111--020001000000010000--0??0110000000000111211101001110-
000010001001112100001011000001---------00---0-0001
Synsphyronus 10000000-00000-0001-1100000000-00-0--
--20---000010001110-0-00003001000-01100----
0001000000201120000011000?010000000000-0010-000000-01302-
1100000001000211111--020001000000010000--0??0110000000000111211101001110-
000010001001112100001011000001---------00---0-0001

'Pseudogarypinus_frontalis_MCZ139246' 10000000-00000-0001-1100000000-00-0----20---000010001110-0-00003001000-01100----0001000000201120000011000?010000000000-0010-000000-01302-11000001000211111--020001000000010000--0??011000000000011121110100111O-000010001001112100001011000001---------00---0-0001
'Larca_granulata_MCZ' 10000000-00000-0001-1100000000-00-0----20---000010001110-0-00003001000-01100----0001000000201120000011000?010000000000-0010-000000-01302-11000001000211111--020001000000010000--0??011000000000011121110100111O-00001?001001112100001011000001---------00---0-0001
'Protogarypinus_giganteus' 10000000-00000-0001-1100000000-00-0----20---000010001110-0-00003001000-01100----0001000000201120000011000?010000000000-0010-000000-01302-11000001000211111--020001000000010000--0??011000000000011121110100111O-00001?001001112100001011000001---------00---0-0001
'Cheiridiidae_MCZ140294' 10000000-00000-0001-1100000000-00-0----20---000010001110-0-00003001000-01100----0001000000201120000011000?010000000000-0010-000000-01302-11000001000211111--020001000000010000--0??011000000000011121110100111O-000010001001112100001011000001---------00---0-0001
'Sternophoridae_133496' 10000000-00000-0001-1100000000-00-0----20---000010001110-0-00003001000-01100----0001000000201120000011000?010000000000-0010-000000-01302-11000001000211111--020001000000010000--0??011000000000011121110100111O-00001?001001112100001011000001---------00---0-0001
'Cacodemonius_MCZ144363' 10000000-00000-0001-1100000000-00-0----20---000010001110-0-00003001000-01100----0001000000201120000011000?010000000000-0010-000000-01302-11000001000211111--020001000000010000--0??011000000000011121110100111O-00001?001001112100001011000001---------00---0-0001
'Oratemnus_curtus_49983' 10000000-00000-0001-1100000000-00-0----20---000010001110-0-00003001000-01100----0001000000201120000011000?010000000000-0010-000000-01302-11000001000211111--020001000000010000--0??011000000000011121110100111O-000011001001112100001011000001---------00---0-0001
'Protochelifer_sp_MCZ' 10000000-00000-0001-1100000000-00-0----20---000010001110-0-00003001000-01100----0001000000201120000011000?010000000000-0010-000000-01302-11000001000211111--020001000000010000--0??011000000000011121110100111O-000011001001112100001011000001---------00---0-0001
'Cheliferidae_JC031_MCZ40982' 10000000-00000-0001-1100000000-00-0----20---000010001110-0-00003001000-01100----0001000000201120000011000?010000000000-0010-000000-01302-

11000000100021111--0200010000000010000--0??011000000000011121110100111O-
0000110010011121000010110000001---------00---0-0001
'Parachelifer_persimilis_MCZ139247' 10000000-00000-0001-
1100000000-00-0----20---000010001110-0-00003001000-01100----
00010000002011200000110000?010000000000-0010-000000-01302-
11000000100021111--0200010000000010000--0??011000000000011121110100111O-
0000110010011121000010110000001---------00---0-0001
'Lamprochernes_savignyi_MCZ' 10000000-00000-0001-
1100000000-00-0----20---000010001110-0-00003001000-01100----
00010000002011200000110000?010000000000-0010-000000-00--2-
11000000100021111--0200010000000010000--0??011000000000011121110100111O-
0000110010011121000010110000001---------00---0-00--
'Conicochernes_crassus_MCZ49985' 10000000-00000-0001-
1100000000-00-0----20---000010001110-0-00003001000-01100----
00010000002011200000110000?010000000000-0010-000000-00--2-
11000000100021111--0200010000000010000--0??011000000000011121110100111O-
0000110010011121000010110000001---------00---0-00--
'Hesperochernes_sp' 10000000-00000-0001-1100000000-00-
0----20---000010001110-0-00003001000-01100----
00010000002011200000110000?010000000000-0010-000000-00--2-
11000000100021111--0200010000000010000--0??011000000000011121110100111O-
0000110010011121000010110000001---------00---0-00--
'Haplochernes_kraepeli_reassaembly' 10000000-00000-0001-
1100000000-00-0----20---000010001110-0-00003001000-01100----
00010000002011200000110000?010000000000-0010-000000-00--2-
11000000100021111--0200010000000010000--0??011000000000011121110100111O-
0000110010011121000010110000001---------00---0-00--
'Chernetidae_MCZ142323' 10000000-00000-0001-1100000000-
00-0----20---000010001110-0-00003001000-01100----
00010000002011200000110000?010000000000-0010-000000-00--2-
11000000100021111--0200010000000010000--0??011000000000011121110100111O-
0000110010011121000010110000001---------00---0-00--
'Chernetidae_sp_JC032' 10000000-00000-0001-1100000000-
00-0----20---000010001110-0-00003001000-01100----
00010000002011200000110000?010000000000-0010-000000-00--2-
11000000100021111--0200010000000010000--0??011000000000011121110100111O-
0000110010011121000010110000001---------00---0-00--
'Ehan_MCZ74291' 0000000110-000-0000-
00000000010100101001111--200-10000000-0-00000000000-00000----
000110000020100000000100000?022000000000-00100?00000-00--2-01--0000101020---
-----00010000110010000001?011000000000000000010-1001110-0000??000100---0--
00-0??0??001---------00---0-00--
'Espe_MCZ60890' 0000000110-000-0000-
00000000010100101001111--200-10000000-0-00000000000-00000----

00011000002010000000010000?022000000000-00100?00000-00--2-01--0000101020---
-----00010000110010000001?01100000000000000010-1001110-0000??000100---0--
00-0??0??001---------00---0-00--
'Espelaea_UWisc' 0000000110-000-0000-
0000000001010010100111--200-10000000-0-00000000000-00000----
00011000002010000000010000?022000000000-00100?00000-00--2-01--0000101020---
-----00010000110010000001?01100000000000000010-1001110-0000??000100---0--
00-0??0??001---------00---0-00--
'Galendromus_occidentalis' 10000001000000-0001-1000000000-
01-0----100--000--0000000-0-01100000000-00000----
00010000002010100??011000?0800000000100010-000000-01402-10--
0010000021111--2000010010001000000-0??011000??000000000010-1001111-
00000-00010101-0--00-101100001---------00---0-00??
'Gromphadorholaelaps_schaeferi_MCZ71202' 10000001000000-0001-
1000000000-01-0----100--000--0000000-0-01100000000-00000----
00010000002010100??011000?0800000000100010-000000-01402-10--
0010000021111--2000010010001000000-0??011000??000000000010-1001111-
00000-00010101-0--00-101100001---------00---0-00??
'Varroa_destructor_SRR3927486' 10000001000000-0001-
1000000000-01-0----100--000--0000000-0-01100000000-00000----
00010000002010100??011000?0800000000100010-000000-01402-10--
0010000021111--2000010010001000000-0??011000??000000000010-1001111-
00000-00010101-0--00-101100001---------00---0-00??
'Pneumolaelaps_niutirani_SRR10269756' 10000001000000-0001-
1000000000-01-0----100--000--0000000-0-01100000000-00000----
00010000002010100??011000?0800000000100010-000000-01402-10--
0010000021111--2000010010001000000-0??011000??000000000010-1001111-
00000-00010101-0--00-101100001---------00---0-00??
'Tropilaelaps_mercedesae' 10000001000000-0001-1000000000-
01-0----100--000--0000000-0-01100000000-00000----
00010000002010100??011000?0800000000100010-000000-01402-10--
0010000021111--2000010010001000000-0??011000??000000000010-1001111-
00000-00010101-0--00-101100001---------00---0-00??
'Adenacarus_sp' 10000001000000-001101000000000-01-
0----100--000-00000000-0-01110000000-00000----
00011000002000100??011000?010000000000-0010-000000-01302-10--
0010000021111--0300010010001020000-0??011000??00000?00001?-1001110-
0001??00010101-0--00-101100001---------00---0-00??
'Neoacarus_sp_MCZ' 10000001000000-001101000000000-
01-0----100--000-00000000-0-01110000000-00000----
00011000002000100??011000?010000000000-0010-000000-01302-10--
0010000021111--0300010010001020000-0??011000??00000?00001?-1001110-
0001??00010101-0--00-101100001---------00---0-00??

'Orthinodoros_rostratus_SRR1732011' 10000001000000-0001-1000000000-01-0----100--000--0000000-0-01100000000-00000----00010000002010100??011000?0800000000100010-000000-01402-10--00100000211111--20000100100010000000-0??011000??000000000010-1001111-00000-00010101-0--00-101100001---------00---0-00??

'IscaW_1_6' 10000001000000-0001-1000000000-01-0----100--000--0000000-0-01100000000-00000----00010000002010100??011000?0800000000100010-000000-01402-10--00100000211111--20000100100010000000-0??011000??000000000010-1001111-00000-00010101-0--00-101100001---------00---0-00??

'Haemaphysalis_longicornis_SRR7754709' 10000001000000-0001-1000000000-01-0----100--000--0000000-0-01100000000-00000----00010000002010100??011000?0800000000100010-000000-01402-10--00100000211111--20000100100010000000-0??011000??000000000010-1001111-00000-00010101-0--00-101100001---------00---0-00??

'Amblyomma_americanum' 10000001000000-0001-1000000000-01-0----100--000--0000000-0-01100000000-00000----00010000002010100??011000?0800000000100010-000000-01402-10--00100000211111--20000100100010000000-0??011000??000000000010-1001111-00000-00010101-0--00-101100001---------00---0-00??

'Dermacentor_silvarum' 10000001000000-0001-1000000000-01-0----100--000--0000000-0-01100000000-00000----00010000002010100??011000?0800000000100010-000000-01402-10--00100000211111--20000100100010000000-0??011000??000000000010-1001111-00000-00010101-0--00-101100001---------00---0-00??

'Hyalomma_excavatum_SRR3157672' 10000001000000-0001-1000000000-01-0----100--000--0000000-0-01100000000-00000----00010000002010100??011000?0800000000100010-000000-01402-10--00100000211111--20000100100010000000-0??011000??000000000010-1001111-00000-00010101-0--00-101100001---------00---0-00??

'Rhipicephalus_micropilus_SRR1186998' 10000001000000-0001-1000000000-01-0----100--000--0000000-0-01100000000-00000----00010000002010100??011000?0800000000100010-000000-01402-10--00100000211111--20000100100010000000-0??011000??000000000010-1001111-00000-00010101-0--00-101100001---------00---0-00??

'Siro_boyerae_SRR1145699' 10001101000000-0000-1111000010-01-0----101--200-10000000-0-00000000000-00000----0001000000201120000010000?040000000000-0010-000000-00-?2-10--00001000211111--01000100000000100001-0??11100000000002-00011010011111110010-00011101000000-001000001--------100---0-00??

'Leptopsalis_141287' 10001101000000000000-1111000010-01-0----101--200-10000000-0-00000000000-00000----0001000000201120000010000?040000000000-0010-000000-014?2-10--

00001000211111--0100010000000100001-0??11100000000002-
000110100111111110010-00011101000000-001000001--------100---0-00??
'Miopsalis_SRR5234539_peptides_iso' 10001101000000000000-
1111000010-01-0----101--200-10000000-0-00000000000-00000----
0001000000201120000010000?040000000000-0010-000000-014?2-10--
00001000211111--0100010000000100001-0??11100000000002-
000110100111111110010-00011101000000-001000001--------100---0-00??
'Metagovea_oviformis_MCZ136533' 10001101000000-0000-
1111000010-01-0----101--200-10000000-0-00000000000-00000----
0001000000201120000010000?040000000000-0010-000000-00-?2-10--
00001000211111--0100010000000100001-0??11100000000002-
000110100111111110010-00011101000000-001000001--------100---0-00??
'Metasiro_savannahensis_IZ-133799' 10001101000000000000-
1111000010-01-0----101--200-10000000-0-00000000000-00000----
0001000000201120000010000?040000000000-0010-000000-004?2-10--
00001000211111--0100010000000100001-0??11100000000002-
000110100111111110010-00011101000000-001000001--------100---0-00??
'Brasilogovea_microphaga_MCZ136559' 10001101000000000000-
1111000010-01-0----101--200-10000000-0-00000000000-00000----
0001000000201120000010000?040000000000-0010-000000-004?2-10--
00001000211111--0100010000000100001-0??11100000000002-
000110100111111110010-00011101000000-001000001--------100---0-00??
'Neogovea_matawai_MCZ133922' 10001101000000-0000-
1111000010-01-0----101--200-10000000-0-00000000000-00000----
0001000000201120000010000?040000000000-0010-000000-00-?2-10--
00001000211111--0100010000000100001-0??11100000000002-
000110100111111110010-00011101000000-001000001--------100---0-00??
'Parapurcellia_silvicola_MCZ134743' 10001101000000000000-
1111000010-01-0----101--200-10000000-0-00000000000-00000----
0001000000201120000010000?040000000000-0010-000000-014?2-10--
00001000211111--0100010000000100001-0??11100000000002-
000110100111111110010-00011101000000-001000001--------100---0-00??
'Purcellia_illustrans_MCZ49518' 10001101000000000000-
1111000010-01-0----101--200-10000000-0-00000000000-00000----
0001000000201120000010000?040000000000-0010-000000-014?2-10--
00001000211111--0100010000000100001-0??11100000000002-
000110100111111110010-00011101000000-001000001--------100---0-00??
'Chileogovea_oedipus_MCZ138106' 10001101000000-0000-
1111000010-01-0----101--200-10000000-0-00000000000-00000----
0001000000201120000010000?040000000000-0010-000000-01-?2-10--
00001000211111--0100010000000100001-0??11100000000002-
000110100111111110010-00011101000000-001000001--------100---0-00??
'Rakaia_minutissima_MCZ143240' 10001101000000000000-
1111000010-01-0----101--200-10000000-0-00000000000-00000----

0001000000201120000010000?040000000000-0010-000000-014?2-10--
00001000211111--010001000000100001-0??11100000000002-
00011010011111110010-00011101000000-001000001--------100---0-00??
'Rakaia_stewartiensis_MCZ49801' 1000110100000000000-
1111000010-01-0----101--200-10000000-0-00000000000-00000----
0001000000201120000010000?040000000000-0010-000000-014?2-10--
00001000211111--010001000000100001-0??11100000000002-
00011010011111110010-00011101000000-001000001--------100---0-00??
'Rakaia_magna_australis_MCZ29212' 10001101000000-0000-
1111000010-01-0----101--200-10000000-0-00000000000-00000----
0001000000201120000010000?040000000000-0010-000000-01-?2-10--
00001000211111--010001000000100001-0??11100000000002-
00011010011111110010-00011101000000-001000001--------100---0-00??
'Rakaia_pauli_MCZ144621' 1000110100000000000-
1111000010-01-0----101--200-10000000-0-00000000000-00000----
0001000000201120000010000?040000000000-0010-000000-014?2-10--
00001000211111--010001000000100001-0??11100000000002-
00011010011111110010-00011101000000-001000001--------100---0-00??
'Rakaia_pauli_Trini_CDhit' 1000110100000000000-1111000010-
01-0----101--200-10000000-0-00000000000-00000----
0001000000201120000010000?040000000000-0010-000000-014?2-10--
00001000211111--010001000000100001-0??11100000000002-
00011010011111110010-00011101000000-001000001--------100---0-00??
'Pettalus_thwaitsei_MCZ132349' 10001101000000-0000-
1111000010-01-0----101--200-10000000-0-00000000000-00000----
0001000000201120000010000?040000000000-0010-000000-01-?2-10--
00001000211111--010001000000100001-0??11100000000002-
00011010011111110010-00011101000000-001000001--------100---0-00??
'Austropurcellia_alata_MCZ141418' 1000110100000000000-
1111000010-01-0----101--200-10000000-0-00000000000-00000----
0001000000201120000010000?040000000000-0010-000000-014?2-10--
00001000211111--010001000000100001-0??11100000000002-
00011010011111110010-00011101000000-001000001--------100---0-00??
'Austropurcellia_despectata_MCZ141417' 1000110100000000000-
1111000010-01-0----101--200-10000000-0-00000000000-00000----
0001000000201120000010000?040000000000-0010-000000-014?2-10--
00001000211111--010001000000100001-0??11100000000002-
00011010011111110010-00011101000000-001000001--------100---0-00??
'Austropurcellia_cf_acuta_MCZ141419' 10001101000000-0000-
1111000010-01-0----101--200-10000000-0-00000000000-00000----
0001000000201120000010000?040000000000-0010-000000-01-?2-10--
00001000211111--010001000000100001-0??11100000000002-
00011010011111110010-00011101000000-001000001--------100---0-00??

'Austropurcellia_acuta_MCZ44060' 10001101000000000-
1111000010-01-0----101--200-10000000-0-00000000000-00000----
00010000002011200000010000?040000000000-0010-000000-014?2-10--
00001000211111--0100010000000100001-0??11100000000002-
0001101001111110010-00011101000000-001000001--------100---0-00??
'Austropurcellia_clousei_MCZ141420' 10001101000000000-
1111000010-01-0----101--200-10000000-0-00000000000-00000----
00010000002011200000010000?040000000000-0010-000000-014?2-10--
00001000211111--0100010000000100001-0??11100000000002-
0001101001111110010-00011101000000-001000001--------100---0-00??
'Neopurcellia_salmoni_MCZ144625' 10001101000000-0000-
1111000010-01-0----101--200-10000000-0-00000000000-00000----
00010000002011200000010000?040000000000-0010-000000-01-?2-10--
00001000211111--0100010000000100001-0??11100000000002-
0001101001111110010-00011101000000-001000001--------100---0-00??
'Neopurcellia_salmoni_MCZ29310' 10001101000000-0000-
1111000010-01-0----101--200-10000000-0-00000000000-00000----
00010000002011200000010000?040000000000-0010-000000-01-?2-10--
00001000211111--0100010000000100001-0??11100000000002-
0001101001111110010-00011101000000-001000001--------100---0-00??
'Karripurcellia_peckorum_MCZ49989' 10001101000000000-
1111000010-01-0----101--200-10000000-0-00000000000-00000----
00010000002011200000010000?040000000000-0010-000000-014?2-10--
00001000211111--0100010000000100001-0??11100000000002-
0001101001111110010-00011101000000-001000001--------100---0-00??
'Aoraki_141173' 10001101000000000-1111000010-01-0-
---101--200-10000000-0-00000000000-00000----
00010000002011200000010000?040000000000-0010-000000-014?2-10--
00001000211111--0100010000000100001-0??11100000000002-
0001101001111110010-00011101000000-001000001--------100---0-00??
'Aoraki_denticulata_Northern_MCZ141173' 10001101000000-0000-
1111000010-01-0----101--200-10000000-0-00000000000-00000----
00010000002011200000010000?040000000000-0010-000000-01-?2-10--
00001000211111--0100010000000100001-0??11100000000002-
0001101001111110010-00011101000000-001000001--------100---0-00??
'Aoraki_denticulata_Southern_MCZ141172' 10001101000000000-
1111000010-01-0----101--200-10000000-0-00000000000-00000----
00010000002011200000010000?040000000000-0010-000000-014?2-10--
00001000211111--0100010000000100001-0??11100000000002-
0001101001111110010-00011101000000-001000001--------100---0-00??
'Aoraki_longitarsa_MCZ137238' 10001101000000000-
1111000010-01-0----101--200-10000000-0-00000000000-00000----
00010000002011200000010000?040000000000-0010-000000-014?2-10--

00001000211111--0100010000000100001-0??11100000000002-
0001101001111110010-00011101000000-001000001--------100---0-00??
'Acropsopilio_neozealandiae_MCZ30457' 10001001000000-0000-
1111000011001-0----101--200-10000000-0-00000000000-00100----
00010000102011200000010000?040000000000-0010-000000-00--1010--
00001000211111--0100010000000100001-
0??1110000000000100001101001111010010-00011100-0--00-001000001--------010---
0-0000
'Hesperonemastoma_modestum' 10001001000000-0000-
1111000011001-0----101--201-10000000-0-00000000000-00100----
00010000102011200000010000?040000000000-0010-000000-00--1010--
00001000211111--0100010000000100001-
0??1110000000000100001101001111010010-00011100-0--00-001000001--------010---
0-0000
'Ortholasma_coronadense' 10001001000000-0000-
1111000011001-0----101--201-10000000-0-00000000000-00100----
00010000102011200000010000?040000000000-0010-000000-00--1010--
00001000211111--0100010000000100001-
0??1110000000000100001101001111010010-00011100-0--00-001000001--------010---
0-0000
'Trogulus_martensi' 10001001000000-0000-1111000011001-
0----101--201-10000000-0-00000000000-00100----
00010000102011200000010000?040000000000-0010-000000-00--1010--
00001000211111--0100010000000100001-
0??1110000000000100001101001111010010-00011100-0--00-001000001--------010---
0-0000
'Caddo_agilis' 10001001001000-0000-1111000011001-0---
-101--200-10000000-0-00000000000-00000----
00010000102011200000010000?040000000000-0010-000000-00--1010--
00001000211111--0100010000000100001-
0??1110000000000100001101001111010010-00011100-0--00-001000001--------010---
0-0000
'thrasychirus_49762' 10001001001000-0000-1111000011001-
0----101--200-10000000-0-00000000000-00000----
00010000102011200000010000?040000000000-0010-000000-00--1010--
00001000211111--0100010000000100001-
0??1110000000000100001101001111010010-00011100-0--00-001000001--------010---
0-0000
'Forsteropsalis_pureora_MCZ29216' 10001001001000-0000-
1111000011001-0----101--200-10000000-0-00000000000-00000----
00010000102011200000010000?040000000000-0010-000000-00--1010--
00001000211111--0100010000000100001-
0??1110000000000100001101001111010010-00011100-0--00-001000001--------010---
0-0000

'Pantopsalis_nz2016' 10001001001000-0000-1111000011001-
0----101--200-10000000-0-00000000000-00000----
0001000010201120000010000?040000000000-0010-000000-00--1010--
00001000211111--0100010000000100001-
0??1110000000000100001101001111010010-00011100-0--00-001000001--------010---
0-0000
'mangatangi_133381' 10001001001000-0000-
1111000011001-0----101--200-10000000-0-00000000000-00000----
0001000010201120000010000?040000000000-0010-000000-00--1010--
00001000211111--0100010000000100001-
0??1110000000000100001101001111010010-00011100-0--00-001000001--------010---
0-0000
'megalopsalis_133420' 10001001001000-0000-
1111000011001-0----101--200-10000000-0-00000000000-00000----
0001000010201120000010000?040000000000-0010-000000-00--1010--
00001000211111--0100010000000100001-
0??1110000000000100001101001111010010-00011100-0--00-001000001--------010---
0-0000
'Diguetinus_143123' 10001001001000-0000-1111000011001-
0----101--200-10000000-0-00000000000-00000----
0001000010201120000010000?040000000000-0010-000000-00--1010--
00001000211111--0100010000000100001-
0??1110000000000100001101001111010010-00011100-0--00-001000001--------010---
0-0000
'Phalangium_opilio_IZ-29189' 10001001001000-0000-
1111000011001-0----101--200-10000000-0-00000000000-00000----
0001000010201120000010000?040000000000-0010-000000-00--1010--
00001000211111--0100010000000100001-
0??1110000000000100001101001111010010-00011100-0--00-001000001--------010---
0-0000
'Protolophus_singularis' 10001001001000-0000-1111000011001-
0----101--200-10000000-0-00000000000-00000----
0001000010201120000010000?040000000000-0010-000000-00--1010--
00001000211111--0100010000000100001-
0??1110000000000100001101001111010010-00011100-0--00-001000001--------010---
0-0000
'prionostemma_144108' 10001001001000-0000-
1111000011001-0----101--200-10000000-0-00000000000-00000----
0001000010201120000010000?040000000000-0010-000000-00--1010--
00001000211111--0100010000000100001-
0??1110000000000100001101001111010010-00011100-0--00-001000001--------010---
0-0000
'Leiobunum_verrucosum' 10001001001000-0000-
1111000011001-0----101--200-10000000-0-00000000000-00000----

0001000010201120000010000?040000000000-0010-000000-00--1010--
00001000211111--0100010000000100001-
0??1110000000000100001101001111010010-00011100-0--00-001000001--------010---
0-0000
'paranelima_143124' 10001001001000-0000-
1111000011001-0----101--200-10000000-0-00000000000-00000----
0001000010201120000010000?040000000000-0010-000000-00--1010--
00001000211111--0100010000000100001-
0??1110000000000100001101001111010010-00011100-0--00-001000001--------010---
0-0000
'Sclerobunus_robustus' 10001001000000-0000-
1111000011001-0----101--200-10000000-0-00001000000-00000----
0001000010201120000010001?040000000000-0010-000000-00--1010--
00001000211111--0100010000000100001-
0??1110000000000100001101001111010010-00011100-0--00-001000001--------010---
0-0000
'Synthetonychia_glacialis_MCZ137212' 10001001000000-0000-
1111000011001-0----101--200-10000000-0-00001000000-00000----
0001000010201120000010001?040000000000-0010-000000-00--1010--
00001000211111--0100010000000100001-
0??1110000000000100001101001111010010-00011100-0--00-001000001--------000---
0-0000
'Larifuga_capensis_IZ-49747' 10001001000000-0000-
1111000011001-0----101--200-10000000-0-00001000000-00000----
0001000010201120000010001?040000000000-0010-000000-00--1010--
00001000211111--0100010000000100001-
0??1110000000000100001101001111010010-00011100-0--00-001000001--------010---
0-0000
'Nuncia_nz1211' 10001001000000-0000-1111000011001-
0----101--200-10000000-0-00001000000-00000----
0001000010201120000010001?040000000000-0010-000000-00--1010--
00001000211111--0100010000000100001-
0??1110000000000100001101001111010010-00011100-0--00-001000001--------010---
0-0000
Pristobunus 10001001000000-0000-1111000011001-0---
-101--200-10000000-0-00001000000-00000----
0001000010201120000010001?040000000000-0010-000000-00--1010--
00001000211111--0100010000000100001-
0??1110000000000100001101001111010010-00011100-0--00-001000001--------010---
0-0000
'Scotolemon_lespesi_MCZ43672' 10001001000000-0000-
1111000011001-0----101--200-10000000-0-00001000000-00000----
0001000010201120000010001?040000000000-0010-000000-00--1010--
00001000211111--0100010000000100001-

0??111000000000010000110100111010010-00011100-0--00-001000001--------010---0-0000
'Sitalcina_lobata' 10001001000000-0000-1111000011001-0----101--200-10000000-0-00001000000-00000----00010000102011200000010001?040000000000-0010-000000-00--1010--00001000211111--0100010000000100001-0??111000000000010000110100111010010-00011100-0--00-001000001--------010---0-0000
'Gnomulus_141286' 10001001000000-0000-1111000011001-0----101--200-10000000-0-00001000000-00000----00010000102011200000010001?040000000000-0010-000000-00--1010--00001000211111--0100010000000100001-0??111000000000010000110100111010010-00011100-0--00-001000001--------010---0-0000
'Dibunus_141285' 10001001000000-0000-1111000011001-0----101--200-10000000-0-00001000000-00000----00010000102011200000010001?040000000000-0010-000000-00--1010--00001000211111--0100010000000100001-0??111000000000010000110100111010010-00011100-0--00-001000001--------000---0-0000
'Epedanus_141288' 10001001000000-0000-1111000011001-0----101--200-10000000-0-00001000000-00000----00010000102011200000010001?040000000000-0010-000000-00--1010--00001000211111--0100010000000100001-0??111000000000010000110100111010010-00011100-0--00-001000001--------010---0-0000
'Haasus_judaeus' 10001001000000-0000-1111000011001-0----101--200-10000000-0-00001000000-00000----00010000102011200000010001?040000000000-0010-000000-00--1010--00001000211111--0100010000000100001-0??111000000000010000110100111010010-00011100-0--00-001000001--------010---0-0000
'Haasus_naasane' 10001001000000-0000-1111000011001-0----101--200-10000000-0-00001000000-00000----00010000102011200000010001?040000000000-0010-000000-00--2-10--00001000211111--0100010000000100001-0??111000000000010000110100111010010-00011100-0--00-001000001--------010---0-00--
Dampetrus 10001001000000-0000-1111000011001-0----101--200-10000000-0-00001000000-00000----00010000102011200000010001?040000000000-0010-000000-00--1010--00001000211111--0100010000000100001-0??111000000000010000110100111010010-00011100-0--00-001000001--------010---0-0000

'Metabiantes_sp_IZ-49748' 10001001000000-0000-1111000011001-0----101--200-10000000-0-00001000000-00000----00010000102011200000100001?040000000000-0010-000000-00--1010--00001000211111--0100010000000100001-0??1110000000000100001101001111010010-00011100-0--00-001000001--------000---0-0000
'Stygnomma_144060' 10001001000000-0000-1111000011001-0----101--200-10000000-0-00001000000-00000----00010000102011200000100001?040000000000-0010-000000-00--1010--00001000211111--0100010000000100001-0??1110000000000100001101001111010010-00011100-0--00-001000001--------000---0-0000
pellobunus 10001001000000-0000-1111000011001-0----101--200-10000000-0-00001000000-00000----00010000102011200000100001?040000000000-0010-000000-00--1010--00001000211111--0100010000000100001-0??1110000000000100001101001111010010-00011100-0--00-001000001--------010---0-0000
'Kimulidae_143960' 10001001000000-0000-1111000011001-0----101--200-10000000-0-00001000000-00000----00010000102011200000100001?040000000000-0010-000000-00--1010--00001000211111--0100010000000100001-0??1110000000000100001101001111010010-00011100-0--00-001000001--------010---0-0000
'Tegipiolus_pachypus_MZSP71018' 10001001000000-0000-1111000011001-0----101--200-10000000-0-00001000000-00000----00010000102011200000100001?040000000000-0010-000000-00--1010--00001000211111--0100010000000100001-0??1110000000000100001101001111010010-00011100-0--00-001000001--------010---0-0000
'pellobunus_144055' 10001001000000-0000-1111000011001-0----101--200-10000000-0-00001000000-00000----00010000102011200000100001?040000000000-0010-000000-00--1010--00001000211111--0100010000000100001-0??1110000000000100001101001111010010-00011100-0--00-001000001--------010---0-0000
'Pachylicus_acutus_IZ-49749' 10001001000000-0000-1111000011001-0----101--200-10000000-0-00001000000-00000----00010000102011200000100001?040000000000-0010-000000-00--1010--00001000211111--0100010000000100001-0??1110000000000100001101001111010010-00011100-0--00-001000001--------010---0-0000
'maracaynatum_144047' 10001001000000-0000-1111000011001-0----101--200-10000000-0-00001000000-00000----

0001000010201120000010001?040000000000-0010-000000-00--1010--
00001000211111--0100010000000100001-
0??1110000000000100001101001111010010-00011100-0--00-001000001--------010---
0-0000
Avima 10001001000000-0000-1111000011001-0----
101--200-10000000-0-00001000000-00000----
0001000010201120000010001?040000000000-0010-000000-00--1010--
00001000211111--0100010000000100001-
0??1110000000000100001101001111010010-00011100-0--00-001000001--------010---
0-0000
Karos 10001001000000-0000-1111000011001-0----
101--200-10000000-0-00001000000-00000----
0001000010201120000010001?040000000000-0010-000000-00--1010--
00001000211111--0100010000000100001-
0??1110000000000100001101001111010010-00011100-0--00-001000001--------010---
0-0000
chapolobunus 10001001000000-0000-1111000011001-0-
---101--200-10000000-0-00001000000-00000----
0001000010201120000010001?040000000000-0010-000000-00--1010--
00001000211111--0100010000000100001-
0??1110000000000100001101001111010010-00011100-0--00-001000001--------010---
0-0000
'Stygnoplus_143966' 10001001000000-0000-
1111000011001-0----101--200-10000000-0-00001000000-00000----
0001000010201120000010001?040000000000-0010-000000-00--1010--
00001000211111--0100010000000100001-
0??1110000000000100001101001111010010-00011100-0--00-001000001--------000---
0-0000
Protimesius 10001001000000-0000-1111000011001-0---
-101--200-10000000-0-00001000000-00000----
0001000010201120000010001?040000000000-0010-000000-00--1010--
00001000211111--0100010000000100001-
0??1110000000000100001101001111010010-00011100-0--00-001000001--------000---
0-0000
'Auranus_hoeferscovitorum_MCZ136531' 10001001000000-0000-
1111000011001-0----101--200-10000000-0-00001000000-00000----
0001000010201120000010001?040000000000-0010-000000-00--1010--
00001000211111--0100010000000100001-
0??1110000000000100001101001111010010-00011100-0--00-001000001--------010---
0-0000
'Pickeliana_pickeli_MCZ139249' 10001001000000-0000-
1111000011001-0----101--200-10000000-0-00001000000-00000----
0001000010201120000010001?040000000000-0010-000000-00--1010--
00001000211111--0100010000000100001-

0??1110000000000010000110100111010010-00011100-0--00-001000001--------010---
0-0000
'Gonycranaus_pluto_MCZ139251' 10001001000000-0000-
1111000011001-0----101--200-10000000-0-00001000000-00000----
0001000010201120000010001?040000000000-0010-000000-00--1010--
00001000211111--0100010000000100001-
0??1110000000000010000110100111010010-00011100-0--00-001000001--------010---
0-0000
'Camarana_sp_MCZ139271' 10001001000000-0000-
1111000011001-0----101--200-10000000-0-00001000000-00000----
0001000010201120000010001?040000000000-0010-000000-00--1010--
00001000211111--0100010000000100001-
0??1110000000000010000110100111010010-00011100-0--00-001000001--------010---
0-0000
Pseudopachylus 10001001000000-0000-1111000011001-
0----101--200-10000000-0-00001000000-00000----
0001000010201120000010001?040000000000-0010-000000-00--1010--
00001000211111--0100010000000100001-
0??1110000000000010000110100111010010-00011100-0--00-001000001--------010---
0-0000
'Bissula_spnov_MCZ139255' 10001001000000-0000-
1111000011001-0----101--200-10000000-0-00001000000-00000----
0001000010201120000010001?040000000000-0010-000000-00--1010--
00001000211111--0100010000000100001-
0??1110000000000010000110100111010010-00011100-0--00-001000001--------010---
0-0000
'Pseudopachylus_sp_MCZ139280' 10001001000000-0000-
1111000011001-0----101--200-10000000-0-00001000000-00000----
0001000010201120000010001?040000000000-0010-000000-00--1010--
00001000211111--0100010000000100001-
0??1110000000000010000110100111010010-00011100-0--00-001000001--------010---
0-0000
'Vonones_ornata_IZ-49750' 10001001000000-0000-
1111000011001-0----101--200-10000000-0-00001000000-00000----
0001000010201120000010001?040000000000-0010-000000-00--1010--
00001000211111--0100010000000100001-
0??1110000000000010000110100111010010-00011100-0--00-001000001--------010---
0-0000
'Incasarcus_argenteus_MCZ139250' 10001001000000-0000-
1111000011001-0----101--200-10000000-0-00001000000-00000----
0001000010201120000010001?040000000000-0010-000000-00--1010--
00001000211111--0100010000000100001-
0??1110000000000010000110100111010010-00011100-0--00-001000001--------010---
0-0000

'Quindina_limbata_MZSP71772'	10001001000000-0000-1111000011001-0----101--200-10000000-0-00001000000-00000----00010000102011200000010001?040000000000-0010-000000-00--1010--00001000211111--0100010000000100001-0??111000000000010000110100111010010-00011100-0--00-001000001--------010---0-0000
Saramacia	10001001000000-0000-1111000011001-0----101--200-10000000-0-00001000000-00000----00010000102011200000010001?040000000000-0010-000000-00--1010--00001000211111--0100010000000100001-0??111000000000010000110100111010010-00011100-0--00-001000001--------010---0-0000
'Ampycinae_sp_MCZ46455'	10001001000000-0000-1111000011001-0----101--200-10000000-0-00001000000-00000----00010000102011200000010001?040000000000-0010-000000-00--1010--00001000211111--0100010000000100001-0??111000000000010000110100111010010-00011100-0--00-001000001--------010---0-0000
'Glysterus_sp_MCZ140069'	10001001000000-0000-1111000011001-0----101--200-10000000-0-00001000000-00000----00010000102011200000010001?040000000000-0010-000000-00--1010--00001000211111--0100010000000100001-0??111000000000010000110100111010010-00011100-0--00-001000001--------010---0-0000
'Panalus_robustus_MZSP71019'	10001001000000-0000-1111000011001-0----101--200-10000000-0-00001000000-00000----00010000102011200000010001?040000000000-0010-000000-00--1010--00001000211111--0100010000000100001-0??111000000000010000110100111010010-00011100-0--00-001000001--------010---0-0000
'Phalangodus_cottus_MZSP71020'	10001001000000-0000-1111000011001-0----101--200-10000000-0-00001000000-00000----00010000102011200000010001?040000000000-0010-000000-00--1010--00001000211111--0100010000000100001-0??111000000000010000110100111010010-00011100-0--00-001000001--------010---0-0000
'Santinezia_144078'	10001001000000-0000-1111000011001-0----101--200-10000000-0-00001000000-00000----00010000102011200000010001?040000000000-0010-000000-00--1010--00001000211111--0100010000000100001-0??111000000000010000110100111010010-00011100-0--00-001000001--------010---0-0000
'phareicranaus_136554'	10001001000000-0000-1111000011001-0----101--200-10000000-0-00001000000-00000----

00010000102011200000010001?040000000000-0010-000000-00--1010--
00001000211111--0100010000000100001-
0??1110000000000100001101001111010010-00011100-0--00-001000001--------010---
0-0000
'Pseudopucrolia_discrepans_MCZ139277' 10001001000000-0000-
1111000011001-0----101--200-10000000-0-00001000000-00000----
00010000102011200000010001?040000000000-0010-000000-00--1010--
00001000211111--0100010000000100001-
0??1110000000000100001101001111010010-00011100-0--00-001000001--------010---
0-0000
'Sadocus_polyacanthus_MCZ138113' 10001001000000-0000-
1111000011001-0----101--200-10000000-0-00001000000-00000----
00010000102011200000010001?040000000000-0010-000000-00--1010--
00001000211111--0100010000000100001-
0??1110000000000100001101001111010010-00011100-0--00-001000001--------010---
0-0000
'Chilegyndes_phillipsoni_MCZ49760' 10001001000000-0000-
1111000011001-0----101--200-10000000-0-00001000000-00000----
00010000102011200000010001?040000000000-0010-000000-00--1010--
00001000211111--0100010000000100001-
0??1110000000000100001101001111010010-00011100-0--00-001000001--------010---
0-0000
metagyndes 10001001000000-0000-1111000011001-0-
---101--200-10000000-0-00001000000-00000----
00010000102011200000010001?040000000000-0010-000000-00--1010--
00001000211111--0100010000000100001-
0??1110000000000100001101001111010010-00011100-0--00-001000001--------010---
0-0000
'Hernandaria_una_MCZ139258' 10001001000000-0000-
1111000011001-0----101--200-10000000-0-00001000000-00000----
00010000102011200000010001?040000000000-0010-000000-00--1010--
00001000211111--0100010000000100001-
0??1110000000000100001101001111010010-00011100-0--00-001000001--------010---
0-0000
'Graphinotus_therezopolis_MCZ139273' 10001001000000-0000-
1111000011001-0----101--200-10000000-0-00001000000-00000----
00010000102011200000010001?040000000000-0010-000000-00--1010--
00001000211111--0100010000000100001-
0??1110000000000100001101001111010010-00011100-0--00-001000001--------010---
0-0000
'Meteusarcoides_139264' 10001001000000-0000-
1111000011001-0----101--200-10000000-0-00001000000-00000----
00010000102011200000010001?040000000000-0010-000000-00--1010--
00001000211111--0100010000000100001-

0??1110000000000100001101001111010010-00011100-0--00-001000001--------010---0-0000
'Meteusarcoides_sp_MZSP71024' 10001001000000-0000-1111000011001-0----101--200-10000000-0-00001000000-00000----0001000010201120000010001?040000000000-0010-000000-00--1010--00001000211111--0100010000000100001-0??1110000000000100001101001111010010-00011100-0--00-001000001--------010---0-0000
'Acutisoma_longipes_MCZ139261' 10001001000000-0000-1111000011001-0----101--200-10000000-0-00001000000-00000----0001000010201120000010001?040000000000-0010-000000-00--1010--00001000211111--0100010000000100001-0??1110000000000100001101001111010010-00011100-0--00-001000001--------010---0-0000
'Goniosoma_sp_MCZ139266' 10001001000000-0000-1111000011001-0----101--200-10000000-0-00001000000-00000----0001000010201120000010001?040000000000-0010-000000-00--1010--00001000211111--0100010000000100001-0??1110000000000100001101001111010010-00011100-0--00-001000001--------010---0-0000
'Roeweria_virescens_MCZ139279' 10001001000000-0000-1111000011001-0----101--200-10000000-0-00001000000-00000----0001000010201120000010001?040000000000-0010-000000-00--1010--00001000211111--0100010000000100001-0??1110000000000100001101001111010010-00011100-0--00-001000001--------010---0-0000
'Longiperna_sp_MCZ139260' 10001001000000-0000-1111000011001-0----101--200-10000000-0-00001000000-00000----0001000010201120000010001?040000000000-0010-000000-00--1010--00001000211111--0100010000000100001-0??1110000000000100001101001111010010-00011100-0--00-001000001--------010---0-0000
iz139272 10001001000000-0000-1111000011001-0----101--200-10000000-0-00001000000-00000----0001000010201120000010001?040000000000-0010-000000-00--1010--00001000211111--0100010000000100001-0??1110000000000100001101001111010010-00011100-0--00-001000001--------010---0-0000
'Mitobates_sp_MCZ139263' 10001001000000-0000-1111000011001-0----101--200-10000000-0-00001000000-00000----0001000010201120000010001?040000000000-0010-000000-00--1010--00001000211111--0100010000000100001-0??1110000000000100001101001111010010-00011100-0--00-001000001--------010---0-0000

'Promitobates_ornatus_MZSP71022' 10001001000000-0000-
1111000011001-0----101--200-10000000-0-00001000000-00000----
00010000102011200000010001?040000000000-0010-000000-00--1010--
00001000211111--010001000000010001-
0??1110000000000100001101001111010010-00011100-0--00-001000001--------010---
0-0000
'Gonyleptellus_sp_MCZ139269' 10001001000000-0000-
1111000011001-0----101--200-10000000-0-00001000000-00000----
00010000102011200000010001?040000000000-0010-000000-00--1010--
00001000211111--010001000000010001-
0??1110000000000100001101001111010010-00011100-0--00-001000001--------010---
0-0000
'Gonyleptes_fragilis_MZSP71023' 10001001000000-0000-
1111000011001-0----101--200-10000000-0-00001000000-00000----
00010000102011200000010001?040000000000-0010-000000-00--1010--
00001000211111--010001000000010001-
0??1110000000000100001101001111010010-00011100-0--00-001000001--------010---
0-0000
'Pseudotrogulus_mirim_MCZ139270' 10001001000000-0000-
1111000011001-0----101--200-10000000-0-00001000000-00000----
00010000102011200000010001?040000000000-0010-000000-00--1010--
00001000211111--010001000000010001-
0??1110000000000100001101001111010010-00011100-0--00-001000001--------010---
0-0000
'Pseudotrogulus_sp_MCZ139278' 10001001000000-0000-
1111000011001-0----101--200-10000000-0-00001000000-00000----
00010000102011200000010001?040000000000-0010-000000-00--1010--
00001000211111--010001000000010001-
0??1110000000000100001101001111010010-00011100-0--00-001000001--------010---
0-0000
'Ampheres_leucopheus_MCZ139267' 10001001000000-0000-
1111000011001-0----101--200-10000000-0-00001000000-00000----
00010000102011200000010001?040000000000-0010-000000-00--1010--
00001000211111--010001000000010001-
0??1110000000000100001101001111010010-00011100-0--00-001000001--------010---
0-0000
'Acanthogonyleptes_aff_fulvigranulatus_MZSP71027' 10001001000000-0000-
1111000011001-0----101--200-10000000-0-00001000000-00000----
00010000102011200000010001?040000000000-0010-000000-00--1010--
00001000211111--010001000000010001-
0??1110000000000100001101001111010010-00011100-0--00-001000001--------010---
0-0000
'Gonyleptes_atrus_MZSP71025' 10001001000000-0000-
1111000011001-0----101--200-10000000-0-00001000000-00000----

00010000102011200000010001?040000000000-0010-000000-00--1010--
00001000211111--0100010000000100001-
0??1110000000000100001101001111010010-00011100-0--00-001000001--------010---
0-0000
'Gonyleptes_sp_MCZ139262' 10001001000000-0000-
1111000011001-0----101--200-10000000-0-00001000000-00000----
00010000102011200000010001?040000000000-0010-000000-00--1010--
00001000211111--0100010000000100001-
0??1110000000000100001101001111010010-00011100-0--00-001000001--------010---
0-0000
'Geraecormobius_bispinifrons_MCZ139274' 10001001000000-0000-
1111000011001-0----101--200-10000000-0-00001000000-00000----
00010000102011200000010001?040000000000-0010-000000-00--1010--
00001000211111--0100010000000100001-
0??1110000000000100001101001111010010-00011100-0--00-001000001--------010---
0-0000
'Mischonyx_cuspidatus_MCZ139265' 10001001000000-0000-
1111000011001-0----101--200-10000000-0-00001000000-00000----
00010000102011200000010001?040000000000-0010-000000-00--1010--
00001000211111--0100010000000100001-
0??1110000000000100001101001111010010-00011100-0--00-001000001--------010---
0-0000
'Sodreana_leprevosti_MCZ139256' 10001001000000-0000-
1111000011001-0----101--200-10000000-0-00001000000-00000----
00010000102011200000010001?040000000000-0010-000000-00--1010--
00001000211111--0100010000000100001-
0??1110000000000100001101001111010010-00011100-0--00-001000001--------010---
0-0000
'Sodreana_sodrena_MCZ139268' 10001001000000-0000-
1111000011001-0----101--200-10000000-0-00001000000-00000----
00010000102011200000010001?040000000000-0010-000000-00--1010--
00001000211111--0100010000000100001-
0??1110000000000100001101001111010010-00011100-0--00-001000001--------010---
0-0000
'Progonyleptoidellus_striatus_MZSP71021' 10001001000000-0000-
1111000011001-0----101--200-10000000-0-00001000000-00000----
00010000102011200000010001?040000000000-0010-000000-00--1010--
00001000211111--0100010000000100001-
0??1110000000000100001101001111010010-00011100-0--00-001000001--------010---
0-0000
'Iporangaia_pustulosa_MCZ139275' 10001001000000-0000-
1111000011001-0----101--200-10000000-0-00001000000-00000----
00010000102011200000010001?040000000000-0010-000000-00--1010--
00001000211111--0100010000000100001-

0??1110000000000100001101001111010010-00011100-0--00-001000001--------010---0-0000
'Gonyleptoides_marumbiensis_MCZ139254' 10001001000000-0000-1111000011001-0----101--200-10000000-0-00001000000-00000----00010000102011200000010001?040000000000-0010-000000-00--1010--00001000211111--010001000000010000l-
0??1110000000000100001101001111010010-00011100-0--00-001000001--------010---0-0000
'Neosadocus_sp_MCZ139257' 10001001000000-0000-1111000011001-0----101--200-10000000-0-00001000000-00000----00010000102011200000010001?040000000000-0010-000000-00--1010--00001000211111--0100010000000100001-
0??1110000000000100001101001111010010-00011100-0--00-001000001--------010---0-0000
'Galeodes_sp' 00000000-0-00000001-1000000100-00-0---20---000010000010-0-00000000000-00010----0001100000200210000011010?022000000000-0010-100000-01301110--01001000211111--120000--10110010-0--0-------0-?000002-1111101001110-00000-10010101-0--01-001001001---------00---0-0000
'Solpugema_sp_IZ-49525' 00000000-0-00000001-1000000100-00-0----20---000010000010-0-00000000000-00010----0001101000200210000011010?022000000000-0010-100000-01301110--01001000211111--120000--10110010-0--0-------0-?000002-1111101001110-00000-10010101-0--01-001001001---------00---0-0000
'Eremobates_sp_IZ-49755' 00000000-0-00000001-1000000100-00-0----20---000010000010-0-00000000000-00010----0001100000200210000011010?022000000000-0010-100000-01301110--01001000211111--120000--10110010-0--0-------0-?000002-1111101001110-00000-10010101-0--01-001001001---------00---0-0000
'Eremobates_sp_new' 00000000-0-00000001-1000000100-00-0----20---000010000010-0-00000000000-00010----0001100000200210000011010?022000000000-0010-100000-01301110--01001000211111--120000--10110010-0--0-------0-?000002-1111101001110-00000-10010101-0--01-001001001---------00---0-0000
Lpol 11110000-1010000100-1010100100-00-1-000101--200-00000000-0-00002000000-00000-----0000000010100011-0001001131-0000000--02000100000-01011200--0000100000100001--01010000000100000-0010000000?000000000000-0000100-00000-00001111000000?001000000---------00---0-1010
Crot 11110000-1010000100-1010100100-00-1-000101--200-00000000-0-00002000000-00000-----0000000010100011-0001001131-0000000--02000100000-01011200--0000100000100001--01010000000100000-0010000000?000000000000-0000100-00000-00001111000000?001000000---------00---0-1010

'Tachypleus_tridentatus_2019G' 11110000-1010000100-1010100100-00-1-000101--200-00000000-0-00002000000-00000-----00000000010100011-0001001131-0000000--02000100000-01011200--0000100000100001--01010000000100000-0010000000?000000000000-0000100-00000-00001111000000?001000000---------00---0-1010
Tgig 11110000-1010000100-1010100100-00-1-000101--200-00000000-0-00002000000-00000-----00000000010100011-0001001131-0000000--02000100000-01011200--0000100000100001--01010000000100000-0010000000?000000000000-0000100-00000-00001111000000?001000000---------00---0-1010
'Ricinoides_atewa' 10000011000000-001000000000100-10000---20---100200000000-0-01002000000-00000----0101000011200010000010000?010000010100-20100000000-015-2-10--00110000211111--200001001000102000--0??0100?0?0000002-21010-1001110-00100-0000011?200010000?100001---------00---0-00??
'Ricinoides_karschii_IZ-130083' 10000011000000-001000000000100-10000---20---100200000000-0-01002000000-00000----0101000011200010000010000?010000010100-20100000000-015-2-10--00110000211111--200001001000102000--0??0100?0?0000002-21010-1001110-00100-0000011?200010000?100001---------00---0-00??
'Pseudocellus_pearsei' 10000011000000-001000000000100-10000---20---100200000000-0-01002000000-00000----0101000011200010000010000?010000010100-20100000000-015-2-10--00110000211111--200001001000102000--0??0100?0?0000002-21010-1001110-00100-0000011?200010000?100001---------00---0-00??
'Pseudocellus_sp_IZ-140060' 10000011000000-001000000000100-10000---20---100200000000-0-01002000000-00000----0101000011200010000010000?010000010100-20100000000-015-2-10--00110000211111--200001001000102000--0??0100?0?0000002-21010-1001110-00100-0000011?200010000?100001---------00---0-00??
'Cryptocellus_sp_IZ-143922' 10000011000000-001000000000100-10000---20---100200000000-0-01002000000-00000----0101000011200010000010000?010000010100-20100000000-015-2-10--00110000211111--200001001000102000--0??0100?0?0000002-21010-1001110-00100-0000011?200010000?100001---------00---0-00??
'Cryptocellus_becki_IZ-136532' 10000011000000-001000000000100-10000---20---100200000000-0-01002000000-00000----0101000011200010000010000?010000010100-20100000000-015-2-10--00110000211111--200001001000102000--0??0100?0?0000002-21010-1001110-00100-0000011?200010000?100001---------00---0-00??
'Cryptocellus_sp_IZ-30913' 10000011000000-001000000000100-10000---20---100200000000-0-01002000000-00000----0101000011200010000010000?010000010100-20100000000-015-2-10--

00110000211111--200001001000102000--0??0100?0?0000002-21010-1001110-00100-0000011?200010000?100001---------00---0-00??
ChaerilusSS 1000000100100000000-0010110010-1011-010100--200310000000-0-00003001000-00100----?0010000002011200000100001000000000000-00100100000-011110110010??110000110110--0001001000010000 1-010110000000000011111101001110-00001100001101210000101000000 1---------00--2101111
TroglokhammouanusSS 1000000100100000000-0010110010-1011-010100--200310000000-0-00003001000-00100----?0010000002011200000100001000000000000-00100100000-014110110010??110000110110--0001001000010000 1-010110000000000011111101001110-00001100001101210000101000000 1---------00--3101111
Vietbocap 1000000100100000000-0010110010-1011-010100--200310000000-0-00003001000-00100----?0010000002011200000100001000000000000-00100100000-014110110010??110000110110--0001001000010000 1-010110000000000011111101001110-00001100001101210000101000000 1---------00--3101111
Lvar 1000000100100000000-0010110010-1011-010100--200310000000-0-00003001000-00100----?0010000002011200000100001000000000000-00100100000-011110110010??110000110110--0001001000010000 1-010110000000000011111101001110-00001100001101210000101000000 1---------00--0101111
'Mesobuthus_martensii' 1000000100100000000-0010110010-1011-010100--200310000000-0-00003001000-00100----?0010000002011200000100001000000000000-00100100000-011110110010??110000110110--0001001000010000 1-010110000000000011111101001110-00001100001101210000101000000 1---------00--0101111
Cley 1000000100100000000-0010110010-1011-010100--200310000000-0-00003001000-00100----?0010000002011200000100001000000000000-00100100000-011110110010??110000110110--0001001000010000 1-010110000000000011111101001110-00001100001101210000101000000 1---------00--0101111
BiisraFull 1000000100100000000-0010110010-1011-010100--200310000000-0-00003001000-00100----?0010000002011200000100001000000000000-00100100000-011110110010??110000110110--0001001000010000 1-010110000000000011111101001110-00001100001101210000101000000 1---------00--0101111

Compsol 1000000100100000000-0010110010-1011-010100--200310000000-0-00003001000-00100----?0010000002011200000100001000000000000-00100100000-011110110010??110000110110--00010010000100001-0101100000000000011111101001110-0000110000110121000010100000001---------00--0101111
'Csch_trinity' 1000000100100000000-0010110010-1011-010100--200310000000-0-00003001000-00100----?0010000002011200000100001000000000000-00100100000-011110110010??110000110110--00010010000100001-0101100000000000011111101001110-0000110000110121000010100000001---------00--0101111
Aamox 1000000100100000000-0010110010-1011-010100--200310000000-0-00003001000-00100----?0010000002011200000100001000000000000-00100100000-011110110010??110000110110--00010010000100001-0101100000000000011111101001110-0000110000110121000010100000001---------00--0101111
'Androctonus_australisSS' 1000000100100000000-0010110010-1011-010100--200310000000-0-00003001000-00100----?0010000002011200000100001000000000000-00100100000-011110110010??110000110110--00010010000100001-0101100000000000011111101001110-0000110000110121000010100000001---------00--0101111
HottentottaSS 1000000100100000000-0010110010-1011-010100--200310000000-0-00003001000-00100----?0010000002011200000100001000000000000-00100100000-011110110010??110000110110--00010010000100001-0101100000000000011111101001110-0000110000110121000010100000001---------00--0101111
'Oscr_trinity' 1000000100100000000-0010110010-1011-010100--200310000000-0-00003001000-00100----?0010000002011200000100001000000000000-00100100000-011110110010??110000110110--00010010000100001-0101100000000000011111101001110-0000110000110121000010100000001---------00--0101111
AcraII 1000000100100000000-0010110010-1011-010100--200310000000-0-00003001000-00100----?0010000002011200000100001000000000000-00100100000-011110110010??110000110110--00010010000100001-0101100000000000011111101001110-0000110000110121000010100000001---------00--0101111
'Bisra_trinity' 1000000100100000000-0010110010-1011-010100--200310000000-0-00003001000-00100----

?0010000002011200000100001000000000000-00100100000-
011110110010??110000110110--00010010000100001-
0101100000000000111111101001110-00001100001101210000101000001---------00-
-0101111
Bare 10000001001000000000-0010110010-1011-
010100--200310000000-0-00003001000-00100----
?0010000002011200000100001000000000000-00100100000-
011110110010??110000110110--00010010000100001-
0101100000000000111111101001110-00001100001101210000101000001---------00-
-0101111
Lheb 10000001001000000000-0010110010-1011-
010100--200310000000-0-00003001000-00100----
?0010000002011200000100001000000000000-00100100000-
011110110010??110000110110--00010010000100001-
0101100000000000111111101001110-00001100001101210000101000001---------00-
-0101111
LquiE 10000001001000000000-0010110010-1011-
010100--200310000000-0-00003001000-00100----
?0010000002011200000100001000000000000-00100100000-
011110110010??110000110110--00010010000100001-
0101100000000000111111101001110-00001100001101210000101000001---------00-
-0101111
Abal 10000001001000000000-0010110010-1011-
010100--200310000000-0-00003001000-00100----
?0010000002011200000100001000000000000-00100100000-
011110110010??110000110110--00010010000100001-
0101100000000000111111101001110-00001100001101210000101000001---------00-
-0101111
Bgig 10000001001000000000-0010110010-1011-
010100--200310000000-0-00003001000-00100----
?0010000002011200000100001000000000000-00100100000-
011110110010??110000110110--00010010000100001-
0101100000000000111111101001110-00001100001101210000101000001---------00-
-0101111
Grosphus 10000001001000000000-0010110010-1011-
010100--200310000000-0-00003001000-00100----
?0010000002011200000100001000000000000-00100100000-
011110110010??110000110110--00010010000100001-
0101100000000000111111101001110-00001100001101210000101000001---------00-
-0101111
ParabuthusSS 10000001001000000000-0010110010-
1011-010100--200310000000-0-00003001000-00100----
?0010000002011200000100001000000000000-00100100000-
011110110010??110000110110--00010010000100001-

01011000000000001111111010011 10-000011000011012100001010000001---------00-
-0101111
Uoli 1000000100100000000-0010110010-1011-
010100--200310000000-0-00003001000-00100----
?0010000002011200000100001000000000000-00100100000-
011110110010??110000110110--00010010000100001-
010110000000000011111110100 1110-000011000011012100001010000001---------00-
-0101111
Uvit 1000000100100000000-0010110010-1011-
010100--200310000000-0-00003001000-00100----
?0010000002011200000100001000000000000-00100100000-
011110110010??110000110110--00010010000100001-
0101100000000000111111101001110-000011000011012100001010000001---------00-
-0101111
Tsmi 1000000100100000000-0010110010-1011-
010100--200310000000-0-00003001000-00100----
?0010000002011200000100001000000000000-00100100000-
011110110010??110000110110--00010010000100001-
0101100000000000111111101001110-000011000011012100001010000001---------00-
-0101111
Tarc 1000000100100000000-0010110010-1011-
010100--200310000000-0-00003001000-00100----
?0010000002011200000100001000000000000-00100100000-
011110110010??110000110110--00010010000100001-
0101100000000000111111101001110-000011000011012100001010000001---------00-
-0101111
Tcos 1000000100100000000-0010110010-1011-
010100--200310000000-0-00003001000-00100----
?0010000002011200000100001000000000000-00100100000-
011110110010??110000110110--00010010000100001-
0101100000000000111111101001110-000011000011012100001010000001---------00-
-0101111
Tserrulatus 1000000100100000000-0010110010-1011-
010100--200310000000-0-00003001000-00100----
?0010000002011200000100001000000000000-00100100000-
011110110010??110000110110--00010010000100001-
0101100000000000111111101001110-000011000011012100001010000001---------00-
-0101111
Tlac 1000000100100000000-0010110010-1011-
010100--200310000000-0-00003001000-00100----
?0010000002011200000100001000000000000-00100100000-
011110110010??110000110110--00010010000100001-
0101100000000000111111101001110-000011000011012100001010000001---------00-
-0101111

Pdeb 1000000100100000000-0010110010-1011-010100--200310000000-0-00003001000-00100----?0010000002011200000100001000000000000-00100100000-011110110010??110000110110--00010010000100001-0101100000000000111111101001110-000011000011012100001010000001---------00--0101111

Jaga 1000000100100000000-0010110010-1011-010100--200310000000-0-00003001000-00100----?0010000002011200000100001000000000000-00100100000-011110110010??110000110110--00010010000100001-0101100000000000111111101001110-000011000011012100001010000001---------00--0101111

Jroc 1000000100100000000-0010110010-1011-010100--200310000000-0-00003001000-00100----?0010000002011200000100001000000000000-00100100000-011110110010??110000110110--00010010000100001-0101100000000000111111101001110-000011000011012100001010000001---------00--0101111

Hjun 1000000100100000000-0010110010-1011-010100--200310000000-0-00003001000-00100----?0010000002011200000100001000000000000-00100100000-011110110010??110000110110--00010010000100001-0101100000000000111111101001110-000011000011012100001010000001---------00--0101111

'Rgar_trinity' 1000000100100000000-0010110010-1011-010100--200310000000-0-00003001000-00100----?0010000002011200000100001000000000000-00100100000-011110110010??110000110110--00010010000100001-0101100000000000111111101001110-000011000011012100001010000001---------00--0101111

Ccar 1000000100100000000-0010110010-1011-010100--200310000000-0-00003001000-00100----?0010000002011200000100001000000000000-00100100000-011110110010??110000110110--00010010000100001-0101100000000000111111101001110-000011000011012100001010000001---------00--0101111

Chentzi 1000000100100000000-0010110010-1011-010100--200310000000-0-00003001000-00100----?0010000002011200000100001000000000000-00100100000-011110110010??110000110110--00010010000100001-0101100000000000111111101001110-000011000011012100001010000001---------00--0101111

'Centruroides_vittatus' 1000000100100000000-0010110010-1011-010100--200310000000-0-00003001000-00100----

?00100000020112000001000010000000000000-00100100000-
011110110010??110000110110--00010010000100001-
0101100000000000111111101001110-000011000011012100001010000001---------00-
-0101111
CscuSS 1000000100100000000-0010110010-1011-
010100--200310000000-0-00003001000-00100----
?00100000020112000001000010000000000000-00100100000-
011110110010??110000110110--00010010000100001-
0101100000000000111111101001110-000011000011012100001010000001---------00-
-0101111
Iurus 1000000100100000000-0010110010-1011-
010100--200310000000-0-00003001000-00100----
?00100000020112000001000010000000000000-00100100000-
011110110010??110000110110--00010010000100001-
0101100000000000111111101001110-000011000011012100001010000001---------00-
-1101111
Bcor 1000000100100000000-0010110010-1011-
010100--200310000000-0-00003001000-00100----
?00100000020112000001000010000000000000-00100100000-
011110110010??110000110110--00010010000100001-
0101100000000000111111101001110-000011000011012100001010000001---------00-
-1101111
'Bothriurus_burmeisteriSS' 1000000100100000000-0010110010-
1011-010100--200310000000-0-00003001000-00100----
?00100000020112000001000010000000000000-00100100000-
011110110010??110000110110--00010010000100001-
0101100000000000111111101001110-000011000011012100001010000001---------00-
-1101111
Centro 1000000100100000000-0010110010-1011-
010100--200310000000-0-00003001000-00100----
?00100000020112000001000010000000000000-00100100000-
011110110010??110000110110--00010010000100001-
0101100000000000111111101001110-000011000011012100001010000001---------00-
-1101111
Cerc 1000000100100000000-0010110010-1011-
010100--200310000000-0-00003001000-00100----
?00100000020112000001000010000000000000-00100100000-
011110110010??110000110110--00010010000100001-
0101100000000000111111101001110-000011000011012100001010000001---------00-
-1101111
Cque 1000000100100000000-0010110010-1011-
010100--200310000000-0-00003001000-00100----
?00100000020112000001000010000000000000-00100100000-
011110110010??110000110110--00010010000100001-

0101100000000000111111101001110-00001100001101210000101000000 1---------00--1101111
Csul 1000000100100000000-0010110010-1011-010100--200310000000-0-00003001000-00100----?0010000002011200000100001000000000000-00100100000-011110110010??110000110110--00010010000100001-0101100000000000111111101001110-000011000011012100001010000001---------00--1101111
Ckeyserlingii 1000000100100000000-0010110010-1011-010100--200310000000-0-00003001000-00100----?0010000002011200000100001000000000000-00100100000-011110110010??110000110110--00010010000100001-0101100000000000111111101001110-000011000011012100001010000001---------00--1101111
SuperSS 1000000100100000000-0010110010-1011-010100--200310000000-0-00003001000-00100----?0010000002011200000100001000000000000-00100100000-011110110010??110000110110--00010010000100001-0101100000000000111111101001110-000011000011012100001010000001---------00--1101111
AnuroctonusSS 1000000100100000000-0010110010-1011-010100--200310000000-0-00003001000-00100----?0010000002011200000100001000000000000-00100100000-011110110010??110000110110--00010010000100001-0101100000000000111111101001110-000011000011012100001010000001---------00--1101111
Brotheas 1000000100100000000-0010110010-1011-010100--200310000000-0-00003001000-00100----?0010000002011200000100001000000000000-00100100000-011110110010??110000110110--00010010000100001-0101100000000000111111101001110-000011000011012100001010000001---------00--1101111
ScorpiopsSS 1000000100100000000-0010110010-1011-010100--200310000000-0-00003001000-00100----?0010000002011200000100001000000000000-00100100000-011110110010??110000110110--00010010000100001-0101100000000000111111101001110-000011000011012100001010000001---------00--1101111
EuscorpiusSS 1000000100100000000-0010110010-1011-010100--200310000000-0-00003001000-00100----?0010000002011200000100001000000000000-00100100000-011110110010??110000110110--00010010000100001-0101100000000000111111101001110-000011000011012100001010000001---------00--1101111

Pdilutus 1000000100100000000-0010110010-1011-
010100--200310000000-0-00003001000-00100----
?00100000020112000001000010000000000000-00100100000-
011110110010??110000110110--00010010000100001-
010110000000000001111111101001110-00001100001101210000101000000 1---------00-
-1101111
'Mega_gertschi' 1000000100100000000-0010110010-
1011-010100--200310000000-0-00003001000-00100----
?00100000020112000001000010000000000000-00100100000-
011110110010??110000110110--00010010000100001-
010110000000000001111111101001110-000011000011012100001010000001---------00-
-1101111
MegacormusSS 1000000100100000000-0010110010-
1011-010100--200310000000-0-00003001000-00100----
?00100000020112000001000010000000000000-00100100000-
011110110010??110000110110--00010010000100001-
010110000000000001111111101001110-000011000011012100001010000001---------00-
-1101111
Umordax 1000000100100000000-0010110010-1011-
010100--200310000000-0-00003001000-00100----
?00100000020112000001000010000000000000-00100100000-
011110110010??110000110110--00010010000100001-
010110000000000001111111101001110-000011000011012100001010000001---------00-
-1101111
BelisariusSS 1000000100100000000-0010110010-1011-
010100--200310000000-0-00003001000-00100----
?00100000020112000001000010000000000000-00100100000-
011110110010??110000110110--00010010000100001-
010110000000000001111111101001110-000011000011012100001010000001---------00-
-11011--
Haztecus 1000000100100000000-0010110010-1011-
010100--200310000000-0-00003001000-00100----
?00100000020112000001000010000000000000-00100100000-
011110110010??110000110110--00010010000100001-
010110000000000001111111101001110-000011000011012100001010000001---------00-
-1101111
HadrSS 1000000100100000000-0010110010-1011-
010100--200310000000-0-00003001000-00100----
?00100000020112000001000010000000000000-00100100000-
011110110010??110000110110--00010010000100001-
010110000000000001111111101001110-000011000011012100001010000001---------00-
-1101111
Hconcolorous 1000000100100000000-0010110010-
1011-010100--200310000000-0-00003001000-00100----

?001000000201120000010000100000000000-00100100000-
011110110010??110000110110--00010010000100001-
010110000000000011111101001110-000011000011012100001010000001---------00-
-1101111
Hspadix 1000000100100000000-0010110010-1011-
010100--200310000000-0-00003001000-00100----
?001000000201120000010000100000000000-00100100000-
011110110010??110000110110--00010010000100001-
010110000000000011111101001110-000011000011012100001010000001---------00-
-1101111
Uhuachuca 1000000100100000000-0010110010-1011-
010100--200310000000-0-00003001000-00100----
?001000000201120000010000100000000000-00100100000-
011110110010??110000110110--00010010000100001-
010110000000000011111101001110-000011000011012100001010000001---------00-
-1101111
Kboggerti 1000000100100000000-0010110010-1011-
010100--200310000000-0-00003001000-00100----
?001000000201120000010000100000000000-00100100000-
011110110010??110000110110--00010010000100001-
010110000000000011111101001110-000011000011012100001010000001---------00-
-1101111
Papacheanus 1000000100100000000-0010110010-
1011-010100--200310000000-0-00003001000-00100----
?001000000201120000010000100000000000-00100100000-
011110110010??110000110110--00010010000100001-
010110000000000011111101001110-000011000011012100001010000001---------00-
-1101111
Vcashi 1000000100100000000-0010110010-1011-
010100--200310000000-0-00003001000-00100----
?001000000201120000010000100000000000-00100100000-
011110110010??110000110110--00010010000100001-
010110000000000011111101001110-000011000011012100001010000001---------00-
-1101111
Pbae 1000000100100000000-0010110010-1011-
010100--200310000000-0-00003001000-00100----
?001000000201120000010000100000000000-00100100000-
011110110010??110000110110--00010010000100001-
010110000000000011111101001110-000011000011012100001010000001---------00-
-1101111
Smes 1000000100100000000-0010110010-1011-
010100--200310000000-0-00003001000-00100----
?001000000201120000010000100000000000-00100100000-
011110110010??110000110110--00010010000100001-

0101100000000000111111101001110-0000110000110121000010100000001---------00--1101111
Svachoni 1000000100100000000-0010110010-1011-010100--200310000000-0-00003001000-00100----?00100000020112000001000010000000000000-00100100000-011110110010??110000110110--00010010000100001-0101100000000000111111101001110-0000110000110121000010100000001---------00--1101111
Vmexicanus 1000000100100000000-0010110010-1011-010100--200310000000-0-00003001000-00100----?00100000020112000001000010000000000000-00100100000-011110110010??110000110110--00010010000100001-0101100000000000111111101001110-0000110000110121000010100000001---------00--1101111
Sgertschi 1000000100100000000-0010110010-1011-010100--200310000000-0-00003001000-00100----?00100000020112000001000010000000000000-00100100000-011110110010??110000110110--00010010000100001-0101100000000000111111101001110-0000110000110121000010100000001---------00--1101111
Kacapulco 1000000100100000000-0010110010-1011-010100--200310000000-0-00003001000-00100----?00100000020112000001000010000000000000-00100100000-011110110010??110000110110--00010010000100001-0101100000000000111111101001110-0000110000110121000010100000001---------00--1101111
Kchamela 1000000100100000000-0010110010-1011-010100--200310000000-0-00003001000-00100----?00100000020112000001000010000000000000-00100100000-011110110010??110000110110--00010010000100001-0101100000000000111111101001110-0000110000110121000010100000001---------00--1101111
Mvariegatus 1000000100100000000-0010110010-1011-010100--200310000000-0-00003001000-00100----?00100000020112000001000010000000000000-00100100000-011110110010??110000110110--00010010000100001-0101100000000000111111101001110-0000110000110121000010100000001---------00--1101111
Pschwenkmeyeri 1000000100100000000-0010110010-1011-010100--200310000000-0-00003001000-00100----?00100000020112000001000010000000000000-00100100000-011110110010??110000110110--00010010000100001-0101100000000000111111101001110-0000110000110121000010100000001---------00--1101111

Pspinigerus 1000000100100000000-0010110010-1011-
010100--200310000000-0-00003001000-00100----
?0010000002011200000100001000000000000-00100100000-
011110110010??110000110110--00010010000100001-
010110000000000011111101001110-000011000011012100001010000001---------00-
-1101111
Kpurepecha 1000000100100000000-0010110010-1011-
010100--200310000000-0-00003001000-00100----
?0010000002011200000100001000000000000-00100100000-
011110110010??110000110110--00010010000100001-
010110000000000011111101001110-000011000011012100001010000001---------00-
-1101111
Ccoahuilae 1000000100100000000-0010110010-1011-
010100--200310000000-0-00003001000-00100----
?0010000002011200000100001000000000000-00100100000-
011110110010??110000110110--00010010000100001-
010110000000000011111101001110-000011000011012100001010000001---------00-
-1101111
Moccidentalis 1000000100100000000-0010110010-1011-
010100--200310000000-0-00003001000-00100----
?0010000002011200000100001000000000000-00100100000-
011110110010??110000110110--00010010000100001-
010110000000000011111101001110-000011000011012100001010000001---------00-
-1101111
Tintrepidus 1000000100100000000-0010110010-1011-
010100--200310000000-0-00003001000-00100----
?0010000002011200000100001000000000000-00100100000-
011110110010??110000110110--00010010000100001-
010110000000000011111101001110-000011000011012100001010000001---------00-
-1101111
Uyaschenkoi 1000000100100000000-0010110010-1011-
010100--200310000000-0-00003001000-00100----
?0010000002011200000100001000000000000-00100100000-
011110110010??110000110110--00010010000100001-
010110000000000011111101001110-000011000011012100001010000001---------00-
-1111111
'Uro_elongatus' 1000000100100000000-0010110010-1011-
010100--200310000000-0-00003001000-00100----
?0010000002011200000100001000000000000-00100100000-
011110110010??110000110110--00010010000100001-
010110000000000011111101001110-000011000011012100001010000001---------00-
-1111111
Urodacus 1000000100100000000-0010110010-1011-
010100--200310000000-0-00003001000-00100----

?0010000002011200000100001000000000000-00100100000-
011110110010??110000110110--00010010000100001-
01011000000000001111111101001110-0000110000110121000010100000001---------00-
-1111111
'Hormiops_davidovi' 1000000100100000000-0010110010-
1011-010100--200310000000-0-00003001000-00100----
?0010000002011200000100001000000000000-00100100000-
011110110010??110000110110--00010010000100001-
01011000000000001111111101001110-0000110000110121000010100000001---------00-
-1111111
'Hormurus_weberi' 1000000100100000000-0010110010-
1011-010100--200310000000-0-00003001000-00100----
?0010000002011200000100001000000000000-00100100000-
011110110010??110000110110--00010010000100001-
01011000000000001111111101001110-0000110000110121000010100000001---------00-
-1111111
Liocheles 1000000100100000000-0010110010-1011-
010100--200310000000-0-00003001000-00100----
?0010000002011200000100001000000000000-00100100000-
011110110010??110000110110--00010010000100001-
01011000000000001111111101001110-0000110000110121000010100000001---------00-
-1111111
'Liocheles_sp_KhaoDeng' 1000000100100000000-0010110010-
1011-010100--200310000000-0-00003001000-00100----
?0010000002011200000100001000000000000-00100100000-
011110110010??110000110110--00010010000100001-
01011000000000001111111101001110-0000110000110121000010100000001---------00-
-1111111
'Nebo_hierochonticus' 1000000100100000000-0010110010-
1011-010100--200310000000-0-00003001000-00100----
?0010000002011200000100001000000000000-00100100000-
011110110010??110000110110--00010010000100001-
01011000000000001111111101001110-0000110000110121000010100000001---------00-
-1111111
DiplocentrusSS 1000000100100000000-0010110010-
1011-010100--200310000000-0-00003001000-00100----
?0010000002011200000100001000000000000-00100100000-
011110110010??110000110110--00010010000100001-
01011000000000001111111101001110-0000110000110121000010100000001---------00-
-1111111
'Diplocentrus_zacatecanus' 1000000100100000000-0010110010-
1011-010100--200310000000-0-00003001000-00100----
?0010000002011200000100001000000000000-00100100000-
011110110010??110000110110--00010010000100001-

0101100000000000111111101001110-0000110000110121000010100000001---------00-
-1111111
Kolotl 1000000100100000000-0010110010-1011-
010100--200310000000-0-00003001000-00100----
?00100000020112000001000010000000000000-00100100000-
011110110010??110000110110--00010010000100001-
0101100000000000111111101001110-0000110000110121000010100000001---------00-
-1111111
'Kolotl_poncei' 1000000100100000000-0010110010-1011-
010100--200310000000-0-00003001000-00100----
?00100000020112000001000010000000000000-00100100000-
011110110010??110000110110--00010010000100001-
0101100000000000111111101001110-0000110000110121000010100000001---------00-
-1111111
'Iomachus_politus' 1000000100100000000-0010110010-
1011-010100--200310000000-0-00003001000-00100----
?00100000020112000001000010000000000000-00100100000-
011110110010??110000110110--00010010000100001-
0101100000000000111111101001110-0000110000110121000010100000001---------00-
-1111111
HadogenesSS 1000000100100000000-0010110010-
1011-010100--200310000000-0-00003001000-00100----
?00100000020112000001000010000000000000-00100100000-
011110110010??110000110110--00010010000100001-
0101100000000000111111101001110-0000110000110121000010100000001---------00-
-1111111
'Hadogenes_paucidens' 1000000100100000000-0010110010-
1011-010100--200310000000-0-00003001000-00100----
?00100000020112000001000010000000000000-00100100000-
011110110010??110000110110--00010010000100001-
0101100000000000111111101001110-0000110000110121000010100000001---------00-
-1111111
'Opistacanthus_asper' 1000000100100000000-0010110010-
1011-010100--200310000000-0-00003001000-00100----
?00100000020112000001000010000000000000-00100100000-
011110110010??110000110110--00010010000100001-
0101100000000000111111101001110-0000110000110121000010100000001---------00-
-1111111
'Opistacanthus_validus' 1000000100100000000-0010110010-
1011-010100--200310000000-0-00003001000-00100----
?00100000020112000001000010000000000000-00100100000-
011110110010??110000110110--00010010000100001-
0101100000000000111111101001110-0000110000110121000010100000001---------00-
-1111111

'Chiromachus_ochropus' 1000000100100000000-0010110010-1011-010100--200310000000-0-00003001000-00100----?0010000002011200000100001000000000000-00100100000-011110110010??110000110110--00010010000100001-0101100000000000111111101001110-00001100001101210000101000001---------00--1111111

OpisthacanthusSS 1000000100100000000-0010110010-1011-010100--200310000000-0-00003001000-00100----?0010000002011200000100001000000000000-00100100000-011110110010??110000110110--00010010000100001-0101100000000000111111101001110-00001100001101210000101000001---------00--1111111

'Palaeocheloctonus_pauliani' 1000000100100000000-0010110010-1011-010100--200310000000-0-00003001000-00100----?0010000002011200000100001000000000000-00100100000-011110110010??110000110110--00010010000100001-0101100000000000111111101001110-00001100001101210000101000001---------00--1111111

'Opistophthalmus_glabrifrons' 1000000100100000000-0010110010-1011-010100--200310000000-0-00003001000-00100----?0010000002011200000100001000000000000-00100100000-011110110010??110000110110--00010010000100001-0101100000000000111111101001110-00001100001101210000101000001---------00--1111111

PandinusSS 1000000100100000000-0010110010-1011-010100--200310000000-0-00003001000-00100----?0010000002011200000100001000000000000-00100100000-011110110010??110000110110--00010010000100001-0101100000000000111111101001110-00001100001101210000101000001---------00--1111111

'Opistophthalmus_walhbergi' 1000000100100000000-0010110010-1011-010100--200310000000-0-00003001000-00100----?0010000002011200000100001000000000000-00100100000-011110110010??110000110110--00010010000100001-0101100000000000111111101001110-00001100001101210000101000001---------00--1111111

'Opistophthalmus_boehmi' 1000000100100000000-0010110010-1011-010100--200310000000-0-00003001000-00100----?0010000002011200000100001000000000000-00100100000-011110110010??110000110110--00010010000100001-0101100000000000111111101001110-00001100001101210000101000001---------00--1111111

Sfuscus 1000000100100000000-0010110010-1011-010100--200310000000-0-00003001000-00100----

?0010000002011200000100001000000000000-00100100000-
011110110010??110000110110--00010010000100001-
010110000000000011111101001110-0000110000110121000010100000001---------00-
-1111111
'Cioa_Trinity'
1000000110000000010110000000001110000---20-01110010100000-
10000010010110100011000100111?20021100100011100010120000000000-
00100110000-01201011000000010000001110--0011101000112011-
1110011111??111110000010-1111110-000011011001113110100001011001---------
00---0-1111
'Cisr_Trinity' 1000000110000000010110000000001110000-
--20-01110010100000-
10000010010110100011000100111?20021100100011100010120000000000-
00100110000-01201011000000010000001110--0011101000112011-
1110011111??111110000010-1111110-000011011001113110100001011001---------
00---0-1111
'Dvar_IZ29740'
1000000110000000010110000000001110000---20-11110010100000-
10000010010110100011000100111?20021100100011100010120000000000-
00100110000-012010110000000100000001110--0011101000102011-
1110011111??111110000010-1111110-000011011001113110100001011001---------
00---0-1111
'Acor_IZ71252'
1000000110000000010110000000001110000---20-11110010100000-
10000010010110100011000100111?20021100100011100010120000000000-
00100110000-012010110000000100000001110--0011101000102011-
1110011111??111110000010-1111110-000011011001113110100001011001---------
00---0-1111
'Pmar_Trinity'
1000000110000000010110000000001110000---20-11110010100000-
10000010010110100011000100111?20021100100011100010120000000000-
00100110000-012010110000000100000001110--0011101000102011-
1110011111??111110000010-1111110-000011011001113110100001011001---------
00---0-1111
'Mastigoproctus_giganteus_new'
100000011?00001001011000000000111001010021---110010100000-0-01001101100-
00100----000111100021100100011000010120001000000-00100110010-
01201011100000110000001110--0001111000102011-
1110011111111111100000111111110-000012011001113100100001001001---------
00---0-1111
'Mlac_IZ71253'
100000011?00001001011000000000111001010021---110010100000-0-01001101100-
00100----000111100021100100011000010120001000000-00100110010-
012010111000001100000001110--0001111000102011-

1110011111111111100000111111110-0000120110011131001000010010001---------
00---0-1111
'Draculoides_vinei_IZ-46896' 0000000110-010-
00101100000000111001010121---110010100000-1111001000000-00100----
00011110002110010001100001012000100000-00100110010-00--2-
11100000110000011110--0001111000102011-?1100111111011112-2011111111110-
00001201100111310110000100?001---------00---0-1111
'Stenochrus_portoricensis_MCZ' 0000000110-010-
00101100000000111001010121---110010100000-1111001000000-00100----
00011110002110010001100001012000100000-00100110010-00--2-
11100000110000011110--0001111000102011-?1100111111011112-2011111111110-
00001201100111310110000100?001---------00---0-1111
'Stenochrus_sp_IZ-74289' 0000000110-010-
00101100000000111001010121---110010100000-1111001000000-00100----
00011110002110010001100001012000100000-00100110010-00--2-
11100000110000011110--0001111000102011-?1100111111011112-2011111111110-
00001201100111310110000100?001---------00---0-1111
Liphistius 1000000100000000010010000010011100001--
-20---110111110000-0-00000000000-00000----
00010000002110000010100001012000000000-00100111100-
01201111000011010000001110--0011001000112000-
00100110000000000000001101001110-01000-000001113111100001010001---------
0022-0-1111
'Liphistius_sp2'
1000000100000000001001000001001110001---20---110111110000-0-00000000000-
00000----00010000002110000010100001012000000000-00100111100-
01201111000011010000001110--0011001000112000-
00100110000000000000001101001110-01000-000001113111100001010001---------
0022-0-1111
'Sphodros_rufipes'
1000000100000000001001000001001?10001---20---110011110000-0-00000000000-
00000----000100000021100000101000010120000000001000100111100-
01201111000011110000001110--0011001000112000-
00100110000000000000001101001110-01000-000001113111100001010001---------
000?-0-1111
'Megahexura_fulva'
1000000100000000001001000001001?10001---20---110011110000-0-00000000000-
00000----000100000021100000101000010120000000001000100111100-
01201111000011110000001110--0011001000112000-
00100110000000000000001101001110-01000-000001113111100001010001---------
0002-0-1111
'Aliatypus_coylei'
1000000100000000001001000001001?10001---20---110011110000-0-00000000000-
00000----000100000021100000101000010120000000001000100111100-

01201111000011110000001110--0011001000112000-
00100110000000000000001101001110-01000-000001113111100001010001---------
0000-0-1111
'Antrodiaetus_unicolor'
100000010000000001001000001001?10001---20---110011110000-0-00000000000-
00000----00010000002110000010100001012000000001000100111100-
01201111000011110000001110--0011001000112000-
00100110000000000000001101001110-01000-000001113111100001010001---------
0001-0-1111
'Porrhothele_sp'
100000010000000001001000001001010001---20---110011110000-0-00000000000-
00000----00010000002110000010100001012000000001000100111100-
01201111000011110000001110--0011001000112000-
00100110000000000000001101001110-01000-000001113111100001010001---------
0001-0-1111
'Microhexura_montivaga'
100000010000000001001000001001010001---20---110011110000-0-00000000000-
00000----00010000002110000010100001012000000001000100111100-
01201111000011110000001110--0011001000112000-
00100110000000000000001101001110-01000-000001113111100001010001---------
0001-0-1111
'Macrothele_calpeiana'
100000010000000001001000001001010001---20---110011110000-0-00000000000-
00000----00010000002110000010100001012000000001000100111100-
01201111000011110000001110--0011001000112000-
00100110000000000000001101001110-01000-000001113111100001010001---------
0001-0-1111
'Paratropis_sp'
100000010000000001001000001001010001---20---110011110000-0-00000000000-
00000----00010000002110000010100001012000000001000100111100-
01201111000011110000001110--0011001000112000-
00100110000000000000001101001110-01000-000001113111100001010001---------
0001-0-1111
'Cyclocosmia_truncata'
100000010000000001001000001001010001---20---110011110000-0-00000000000-
00000----00010000002110000010100001012000000001000100111100-
01201111000011110000001110--0011001000112000-
00100110000000000000001101001110-01000-000001113111100001010001---------
0000-0-1111
'Hebestatis_theveneti'
100000010000000001001000001001010001---20---110011110000-0-00000000000-
00000----00010000002110000010100001012000000001000100111100-
01201111000011110000001110--0011001000112000-

00100110000000000000001101001110-01000-000001113111100001010001---------
0000-0-1111
'Idiops_bersebaensis'
1000000100000000001001000001001010001---20---110011110000-0-00000000000-
00000----000100000021100000101000010120000000001000100111100-
012011110000111100000001110--0011001000112000-
00100110000000000000001101001110-01000-000001113111100001010001---------
0000-0-1111
'Aptostichus_stephencolberti'
1000000100000000001001000001001010001---20---110011110000-0-00000000000-
00000----000100000021100000101000010120000000001000100111100-
012011110000111100000001110--0011001000112000-
00100110000000000000001101001110-01000-000001113111100001010001---------
0000-0-1111
'Promyrmekiaphila_clathrata'
1000000100000000001001000001001010001---20---110011110000-0-00000000000-
00000----000100000021100000101000010120000000001000100111100-
012011110000111100000001110--0011001000112000-
00100110000000000000001101001110-01000-000001113111100001010001---------
0000-0-1111
'Brachythele_longitarsis'
1000000100000000001001000001001010001---20---110011110000-0-00000000000-
00000----000100000021100000101000010120000000001000100111100-
012011110000111100000001110--0011001000112000-
00100110000000000000001101001110-01000-000001113111100001010001---------
0001-0-1111
'Pionothele_sp'
1000000100000000001001000001001010001---20---110011110000-0-00000000000-
00000----000100000021100000101000010120000000001000100111100-
012011110000111100000001110--0011001000112000-
00100110000000000000001101001110-01000-000001113111100001010001---------
0000-0-1111
'Stanwellia_sp'
1000000100000000001001000001001010001---20---110011110000-0-00000000000-
00000----000100000021100000101000010120000000001000100111100-
012011110000111100000001110--0011001000112000-
00100110000000000000001101001110-01000-000001113111100001010001---------
0001-0-1111
'Damarchus_sp'
1000000100000000001001000001001010001---20---110011110000-0-00000000000-
00000----000100000021100000101000010120000000001000100111100-
012011110000111100000001110--0011001000112000-
00100110000000000000001101001110-01000-000001113111100001010001---------
0001-0-1111

'Trichopelma_laselva'
1000000100000000001001000001001010001---20---110011110000-0-00000000000-
00000----0001000000211000001010000101200000000100010011 1100-
01201111000011110000001110--0011001000112000-
001001100000000000000001101001110-01000-0000011131111000010100 01---------
0001-0-1111
'Acanthoscurria_geniculata'
1000000100000000001001000001001010001---20---110011110000-0-00000000000-
00000----0001000000211000001010000101200000000100010011 1100-
01201111000011110000001110--0011001000112000-
001001100000000000000001101001110-01000-0000011131111000010100 01---------
0001-0-1111
'A_hentzi_pooled'
1000000100000000001001000001001010001---20---110011110000-0-00000000000-
00000----0001000000211000001010000101200000000100010011 1100-
01201111000011110000001110--0011001000112000-
001001100000000000000001101001110-01000-0000011131111000010100 01---------
0001-0-1111
'Aphonopelma_iviei'
1000000100000000001001000001001010001---20---110011110000-0-00000000000-
00000----0001000000211000001010000101200000000100010011 1100-
01201111000011110000001110--0011001000112000-
001001100000000000000001101001110-01000-0000011131111000010100 01---------
0001-0-1111
'Filistata_insidiatrix'
1000000100000000001001000001001?10001---20---110211110000-0-00000000000-
00000----0001000000211000001010000101200000000100010011 1100-
01201111000011010001011110--0011001000102000-
001001000000000000000001101001110-01000-0000011131111000010100 01---------
0000-0-1111
Kukulcaniahibernalis2
1000000100000000001001000001001?10001---20---110211110000-0-00000000000-
00000----0001000000211000001010000101200000000100010011 1100-
01201111000011010001011110--0011001000102000-
001001000000000000000001101001110-01000-0000011131111000010100 01---------
0000-0-1111
'Hypochilus_gertschi'
1000000100000000001001000001001?10001---20---110211110000-0-00000000000-
00000----0001000000211000001010000101200000000100010011 1100-
01201111000011010001011110--0011001000102000-
001001000000000000000001101001110-01000-0000011131111000010100 01---------
0000-0-1111
'Hypochilus_pococki'
1000000100000000001001000001001?10001---20---110211110000-0-00000000000-

00000----00010000002110000010100001012000000001000100111100-
01201111000011010001011110--0011001000102000-
0010010000000000000000001101001110-01000-000001113111100001010001---------
0000-0-1111
'Calponia_harrisonfordi'
100000010000000001001000001001?10001---20---110211110000-0-00000000000-
00000----00010000002110000010100001012000000001000100111100-
01201111000011010001011110--0011001000102000-
0010010000000000000000001101001110-01000-000001113111100001010001---------
0000-0-1111
'Segestria_sp'
100000010000000001001000001001?10001---20---110211110000-0-00000000000-
00000----00010000002110000010100001012000000001000100111100-
01201111000011010001011110--0011001000102000-
0010010000000000000000001101001110-01000-000001113111100001010001---------
0000-0-1111
'Maoriata_sp'
100000010000000001001000001001?10001---20---110211110000-0-00000000000-
00000----00010000002110000010100001012000000001000100111100-
01201111000011010001011110--0011001000102000-
0010010000000000000000001101001110-01000-000001113111100001010001---------
0000-0-1111
'Orsolobidae_sp'
100000010000000001001000001001?10001---20---110211110000-0-00000000000-
00000----00010000002110000010100001012000000001000100111100-
01201111000011010001011110--0011001000102000-
0010010000000000000000001101001110-01000-000001113111100001010001---------
0000-0-1111
Dysdera
100000010000000001001000001001?10001---20---110211110000-0-00000000000-
00000----00010000002110000010100001012000000001000100111100-
01201111000011010001011110--0011001000102000-
0010010000000000000000001101001110-01000-000001113111100001010001---------
0000-0-1111
'Ischnothyreus_sp'
100000010000000001001000001001?10001---20---110211110000-0-00000000000-
00000----00010000002110000010100001012000000001000100111100-
01201111000011010001011110--0011001000102000-
0010010000000000000000001101001110-01000-000001113111100001010001---------
0000-0-1111
'Opopaea_sp'
100000010000000001001000001001?10001---20---110211110000-0-00000000000-
00000----00010000002110000010100001012000000001000100111100-
01201111000011010001011110--0011001000102000-

00100100000000000000001101001110-01000-000001113111100001010001---------
0000-0-1111
'Diguetia_sp'
100000010000000001001000001001?10001---20---110211110000-0-00000000000-
00000----00010000002110000010100001012000000001000100111100-
01201111000011010001011110--0011001000102000-
00100100000000000000001101001110-01000-000001113111100001010001---------
0000-0-1111
'Pholcus_manueli'
100000010000000001001000001001?10001---20---110211110000-0-00000000000-
00000----00010000002110000010100001012000000001000100111100-
01201111000011010001011110--0011001000102000-
00100100000000000000001101001110-01000-000001113111100001010001---------
0000-0-1111
'Pholcus_phalangioides'
100000010000000001001000001001?10001---20---110211110000-0-00000000000-
00000----00010000002110000010100001012000000001000100111100-
01201111000011010001011110--0011001000102000-
00100100000000000000001101001110-01000-000001113111100001010001---------
0000-0-1111
'Loxosceles_deserta'
100000010000000001001000001001?10001---20---110211110000-0-00000000000-
00000----00010000002110000010100001012000000001000100111100-
01201111000011010001011110--0011001000102000-
00100100000000000000001101001110-01000-000001113111100001010001---------
0000-0-1111
'Drymusa_sp'
100000010000000001001000001001?10001---20---110211110000-0-00000000000-
00000----00010000002110000010100001012000000001000100111100-
01201111000011010001011110--0011001000102000-
00100100000000000000001101001110-01000-000001113111100001010001---------
0000-0-1111
'Periegops_suterii'
100000010000000001001000001001?10001---20---110211110000-0-00000000000-
00000----00010000002110000010100001012000000001000100111100-
01201111000011010001011110--0011001000102000-
00100100000000000000001101001110-01000-000001113111100001010001---------
0000-0-1111
'Ochyrocera_sp'
100000010000000001001000001001?10001---20---110211110000-0-00000000000-
00000----00010000002110000010100001012000000001000100111100-
01201111000011010001011110--0011001000102000-
00100100000000000000001101001110-01000-000001113111100001010001---------
0000-0-1111

'Scytodes_globula'
1000000100000000010010000001001?10001---20---110211110000-0-00000000000-00000----00010000002110000010100001012000000001000100111100-012011110000110100010111110--0011001000102000-00100100000000000000000001101001110-01000-0000011131111000010100001---------0000-0-1111
'Scytodes_thoracica'
1000000100000000010010000001001?10001---20---110211110000-0-00000000000-00000----00010000002110000010100001012000000001000100111100-012011110000110100010111110--0011001000102000-00100100000000000000000001101001110-01000-0000011131111000010100001---------0000-0-1111
Archoleptoneta
1000000100000000010010000001001?10001---20---110211110000-0-00000000000-00000----00010000002110000010100001012000000001000100111100-012011110000110100010111110--0011001000102000-00100100000000000000000001101001110-01000-0000011131111000010100001---------0000-0-1111
Calileptoneta
1000000100000000010010000001001?10001---20---110211110000-0-00000000000-00000----00010000002110000010100001012000000001000100111100-012011110000110100010111110--0011001000102000-00100100000000000000000001101001110-01000-0000011131111000010100001---------0000-0-1111
Austrochilus
1000000100000000010010000001001?10001---20---110211110000-0-00000000000-00000----00010000002110000010100001012000000001000100111100-012011110000110100010111110--0011001000102000-00100100000000000000000001101001110-01000-0000011131111000010100001---------0000-0-1111
'Hickmania_clean'
1000000100000000010010000001001?10001---20---110211110000-0-00000000000-00000----00010000002110000010100001012000000001000100111100-012011110000110100010111110--0011001000102000-00100100000000000000000001101001110-01000-0000011131111000010100001---------0000-0-1111
'Progradungula_GOOD'
1000000100000000010010000001001?10001---20---110211110000-0-00000000000-00000----00010000002110000010100001012000000001000100111100-012011110000110100010111110--0011001000102000-00100100000000000000000001101001110-01000-0000011131111000010100001---------0000-0-1111
'Tarlina_sp' 1000000100000000010010000001001?10001---20---110211110000-0-00000000000-00000----

00010000002110000010100001012000000001000100111100-
01201111000011010001011110--0011001000102000-
00100100000000000000001101001110-01000-000001113111100001010001---------
0000-0-1111
'Gradungula_sorenseni'
100000010000000001001000001001?10001---20---110211110000-0-00000000000-
00000----00010000002110000010100001012000000001000100111100-
01201111000011010001011110--0011001000102000-
00100100000000000000001101001110-01000-000001113111100001010001---------
0000-0-1111
'Pianoa_isolata'
100000010000000001001000001001?10001---20---110211110000-0-00000000000-
00000----00010000002110000010100001012000000001000100111100-
01201111000011010001011110--0011001000102000-
00100100000000000000001101001110-01000-000001113111100001010001---------
0000-0-1111
'Austrarchaea_sp'
100000010000000001001000001001?10001---20---110211110000-0-00000000000-
00000----00010000002110000010100001012000000001000100111100-
01201111000011010001011110--0011001000102000-
00100100000000000000001101001110-01000-000001113111100001010001---------
0000-0-1111
'Mecysmauchenius_clean'
100000010000000001001000001001?10001---20---110211110000-0-00000000000-
00000----00010000002110000010100001012000000001000100111100-
01201111000011010001011110--0011001000102000-
00100100000000000000001101001110-01000-000001113111100001010001---------
0000-0-1111
'Huttonia_palpimanoides'
100000010000000001001000001001?10001---20---110211110000-0-00000000000-
00000----00010000002110000010100001012000000001000100111100-
01201111000011010001011110--0011001000102000-
00100100000000000000001101001110-01000-000001113111100001010001---------
0000-0-1111
'Othiotops_birabeni'
100000010000000001001000001001?10001---20---110211110000-0-00000000000-
00000----00010000002110000010100001012000000001000100111100-
01201111000011010001011110--0011001000102000-
00100100000000000000001101001110-01000-000001113111100001010001---------
0000-0-1111
'Palpimanus_gibbulus'
100000010000000001001000001001?10001---20---110211110000-0-00000000000-
00000----00010000002110000010100001012000000001000100111100-
01201111000011010001011110--0011001000102000-

0010010000000000000000001101001110-01000-000001113111100001010001---------
0000-0-1111
'Stegodyphus_sp'
100000010000000001001000001001?10001---20---110211110000-0-00000000000-
00000----0001000000211000001010000101200000000010001001111100-
01201111000011010001011110--0011001000102000-
0010010000000000000000001101001110-01000-000001113111100001010001---------
0100-0-1111
'Megadictyna_thilenii'
100000010000000001001000001001?10001---20---110211110000-0-00000000000-
00000----0001000000211000001010000101200000000010001001111100-
01201111000011010001011110--0011001000102000-
0010010000000000000000001101001110-01000-000001113111100001010001---------
0100-0-1111
Nicodamidae2
100000010000000001001000001001?10001---20---110211110000-0-00000000000-
00000----0001000000211000001010000101200000000010001001111100-
01201111000011010001011110--0011001000102000-
0010010000000000000000001101001110-01000-000001113111100001010001---------
0100-0-1111
'Nicodamidae_sp1'
100000010000000001001000001001?10001---20---110211110000-0-00000000000-
00000----0001000000211000001010000101200000000010001001111100-
01201111000011010001011110--0011001000102000-
0010010000000000000000001101001110-01000-000001113111100001010001---------
0100-0-1111
'Mysmenella_sp'
100000010000000001001000001001?10001---20---110211110000-0-00000000000-
00000----0001000000211000001010000101200000000010001001111100-
01201111000011010001011110--0011001000102000-
0010010000000000000000001101001110-01000-000001113111100001010001---------
0100-0-1111
'Novanapis_sp'
100000010000000001001000001001?10001---20---110211110000-0-00000000000-
00000----0001000000211000001010000101200000000010001001111100-
01201111000011010001011110--0011001000102000-
0010010000000000000000001101001110-01000-000001113111100001010001---------
0100-0-1111
'Anelosimus_clean'
100000010000000001001000001001?10001---20---110211110000-0-00000000000-
00000----0001000000211000001010000101200000000010001001111100-
01201111000011010001011110--0011001000102000-
0010010000000000000000001101001110-01000-000001113111100001010001---------
0100-0-1111

Latrodectus
1000000100000000001001000001001?10001---20---110211110000-0-00000000000-
00000----000100000002110000010100001012000000001000100111100-
012011110000110100010111110--0011001000102000-
001001000000000000000001101001110-01000-000001113111100001010001---------
0100-0-1111
'Euryopis_sp'
1000000100000000001001000001001?10001---20---110211110000-0-00000000000-
00000----000100000002110000010100001012000000001000100111100-
012011110000110100010111110--0011001000102000-
001001000000000000000001101001110-01000-000001113111100001010001---------
0100-0-1111
'Parasteatoda_tepidariorum'
1000000100000000001001000001001?10001---20---110211110000-0-00000000000-
00000----000100000002110000010100001012000000001000100111100-
012011110000110100010111110--0011001000102000-
001001000000000000000001101001110-01000-000001113111100001010001---------
0100-0-1111
'Pararchaea_alba'
1000000100000000001001000001001?10001---20---110211110000-0-00000000000-
00000----000100000002110000010100001012000000001000100111100-
012011110000110100010111110--0011001000102000-
001001000000000000000001101001110-01000-000001113111100001010001---------
0100-0-1111
'Malkara_clean'
1000000100000000001001000001001?10001---20---110211110000-0-00000000000-
00000----000100000002110000010100001012000000001000100111100-
012011110000110100010111110--0011001000102000-
001001000000000000000001101001110-01000-000001113111100001010001---------
0100-0-1111
'Malkaridae_GH09'
1000000100000000001001000001001?10001---20---110211110000-0-00000000000-
00000----000100000002110000010100001012000000001000100111100-
012011110000110100010111110--0011001000102000-
001001000000000000000001101001110-01000-000001113111100001010001---------
0100-0-1111
'Perissopmeros_sp'
1000000100000000001001000001001?10001---20---110211110000-0-00000000000-
00000----000100000002110000010100001012000000001000100111100-
012011110000110100010111110--0011001000102000-
001001000000000000000001101001110-01000-000001113111100001010001---------
0100-0-1111
'Australomimetus_clean'
1000000100000000001001000001001?10001---20---110211110000-0-00000000000-

00000----0001000000211000001010000101200000000010001001111100-
01201111000011010001011110--0011001000102000-
0010010000000000000000001101001110-01000-000001113111100001010001---------
0100-0-1111
'Ero_leonina'
1000000100000000010010000001001?10001---20---110211110000-0-00000000000-
00000----0001000000211000001010000101200000000010001001111100-
01201111000011010001011110--0011001000102000-
0010010000000000000000001101001110-01000-000001113111100001010001---------
0100-0-1111
'Arkys_sp' 1000000100000000010010000001001?10001-
--20---110211110000-0-00000000000-00000----
0001000000211000001010000101200000000010001001111100-
01201111000011010001011110--0011001000102000-
0010010000000000000000001101001110-01000-000001113111100001010001---------
0100-0-1111
Demadiana
1000000100000000010010000001001?10001---20---110211110000-0-00000000000-
00000----0001000000211000001010000101200000000010001001111100-
01201111000011010001011110--0011001000102000-
0010010000000000000000001101001110-01000-000001113111100001010001---------
0100-0-1111
Leucage
1000000100000000010010000001001?10001---20---110211110000-0-00000000000-
00000----0001000000211000001010000101200000000010001001111100-
01201111000011010001011110--0011001000102000-
0010010000000000000000001101001110-01000-000001113111100001010001---------
0100-0-1111
'Meta_ovalis_good'
1000000100000000010010000001001?10001---20---110211110000-0-00000000000-
00000----0001000000211000001010000101200000000010001001111100-
01201111000011010001011110--0011001000102000-
0010010000000000000000001101001110-01000-000001113111100001010001---------
0100-0-1111
'Nanometa_sp'
1000000100000000010010000001001?10001---20---110211110000-0-00000000000-
00000----0001000000211000001010000101200000000010001001111100-
01201111000011010001011110--0011001000102000-
0010010000000000000000001101001110-01000-000001113111100001010001---------
0100-0-1111
'Tetragnatha_tantalus'
1000000100000000010010000001001?10001---20---110211110000-0-00000000000-
00000----0001000000211000001010000101200000000010001001111100-
01201111000011010001011110--0011001000102000-

0010010000000000000000001101001110-01000-000001113111100001010001---------
0100-0-1111
'Cepheia_clean'
10000001000000000100100000100 1?10001---20---110211110000-0-00000000000-
00000----000100000021100000101000010120000000010001001111 00-
012011110000110100010111 10--0011001000102000-
0010010000000000000000001101001110-01000-000001113111100001010001---------
0100-0-1111
'Mysmena_leichardti'
10000001000000000100100000100 1?10001---20---110211110000-0-00000000000-
00000----000100000021100000101000010120000000010001001111 00-
012011110000110100010111 10--0011001000102000-
0010010000000000000000001101001110-01000-000001113111100001010001---------
0100-0-1111
'Microdipoena_guttata'
10000001000000000100100000100 1?10001---20---110211110000-0-00000000000-
00000----000100000021100000101000010120000000010001001111 00-
012011110000110100010111 10--0011001000102000-
0010010000000000000000001101001110-01000-000001113111100001010001---------
0100-0-1111
'Spinanapis_sp'
10000001000000000100100000100 1?10001---20---110211110000-0-00000000000-
00000----000100000021100000101000010120000000010001001111 00-
012011110000110100010111 10--0011001000102000-
0010010000000000000000001101001110-01000-000001113111100001010001---------
0100-0-1111
'Tekelloides_sp2'
10000001000000000100100000100 1?10001---20---110211110000-0-00000000000-
00000----000100000021100000101000010120000000010001001111 00-
012011110000110100010111 10--0011001000102000-
0010010000000000000000001101001110-01000-000001113111100001010001---------
0100-0-1111
'Laminacauda_clean'
10000001000000000100100000100 1?10001---20---110211110000-0-00000000000-
00000----000100000021100000101000010120000000010001001111 00-
012011110000110100010111 10--0011001000102000-
0010010000000000000000001101001110-01000-000001113111100001010001---------
0100-0-1111
Frontinella 10000001000000000100100000100 1?10001-
--20---110211110000-0-00000000000-00000----
000100000021100000101000010120000000010001001111 00-
012011110000110100010111 10--0011001000102000-
0010010000000000000000001101001110-01000-000001113111100001010001---------
0100-0-1111

'Novafroneta_sp'
100000010000000001001000001001?10001---20---110211110000-0-00000000000-
00000----00010000002110000010100001012000000001000100111100-
012011110000110100010111110--0011001000102000-
001001000000000000000001101001110-01000-000001113111100001010001---------
0100-0-1111
'Synotaxus_sp'
100000010000000001001000001001?10001---20---110211110000-0-00000000000-
00000----00010000002110000010100001012000000001000100111100-
012011110000110100010111110--0011001000102000-
001001000000000000000001101001110-01000-000001113111100001010001---------
0100-0-1111
'Nesticus_bishopi'
100000010000000001001000001001?10001---20---110211110000-0-00000000000-
00000----00010000002110000010100001012000000001000100111100-
012011110000110100010111110--0011001000102000-
001001000000000000000001101001110-01000-000001113111100001010001---------
0100-0-1111
'Nesticus_cooperi'
100000010000000001001000001001?10001---20---110211110000-0-00000000000-
00000----00010000002110000010100001012000000001000100111100-
012011110000110100010111110--0011001000102000-
001001000000000000000001101001110-01000-000001113111100001010001---------
0100-0-1111
'Meringa_sp'
100000010000000001001000001001?10001---20---110211110000-0-00000000000-
00000----00010000002110000010100001012000000001000100111100-
012011110000110100010111110--0011001000102000-
001001000000000000000001101001110-01000-000001113111100001010001---------
0100-0-1111
'Runga_sp'
100000010000000001001000001001?10001---20---110211110000-0-00000000000-
00000----00010000002110000010100001012000000001000100111100-
012011110000110100010111110--0011001000102000-
001001000000000000000001101001110-01000-000001113111100001010001---------
0100-0-1111
'Physoglenes_clean'
100000010000000001001000001001?10001---20---110211110000-0-00000000000-
00000----00010000002110000010100001012000000001000100111100-
012011110000110100010111110--0011001000102000-
001001000000000000000001101001110-01000-000001113111100001010001---------
0100-0-1111
'Physoglenidae_clean'
100000010000000001001000001001?10001---20---110211110000-0-00000000000-

00000----000100000021100000101000010120000000001000100111100-
01201111000011010001011110--0011001000102000-
00100100000000000000001101001110-01000-000001113111100001010001---------
0100-0-1111
'Ogulnius_sp'
1000000100000000001001000001001?10001---20---110211110000-0-00000000000-
00000----000100000021100000101000010120000000001000100111100-
01201111000011010001011110--0011001000102000-
00100100000000000000001101001110-01000-000001113111100001010001---------
0100-0-1111
'Baalzebub_sp'
1000000100000000001001000001001?10001---20---110211110000-0-00000000000-
00000----000100000021100000101000010120000000001000100111100-
01201111000011010001011110--0011001000102000-
00100100000000000000001101001110-01000-000001113111100001010001---------
0100-0-1111
'Theridiosoma_gemmosum'
1000000100000000001001000001001?10001---20---110211110000-0-00000000000-
00000----000100000021100000101000010120000000001000100111100-
01201111000011010001011110--0011001000102000-
00100100000000000000001101001110-01000-000001113111100001010001---------
0100-0-1111
'Theridiosoma_savannum'
1000000100000000001001000001001?10001---20---110211110000-0-00000000000-
00000----000100000021100000101000010120000000001000100111100-
01201111000011010001011110--0011001000102000-
00100100000000000000001101001110-01000-000001113111100001010001---------
0100-0-1111
'Trichonephila_clavipes_Genome'
1000000100000000001001000001001?10001---20---110211110000-0-00000000000-
00000----000100000021100000101000010120000000001000100111100-
01201111000011010001011110--0011001000102000-
00100100000000000000001101001110-01000-000001113111100001010001---------
0100-0-1111
'Micrathena_gracilis'
1000000100000000001001000001001?10001---20---110211110000-0-00000000000-
00000----000100000021100000101000010120000000001000100111100-
01201111000011010001011110--0011001000102000-
00100100000000000000001101001110-01000-000001113111100001010001---------
0100-0-1111
'Verrucosa_arenata'
1000000100000000001001000001001?10001---20---110211110000-0-00000000000-
00000----000100000021100000101000010120000000001000100111100-
01201111000011010001011110--0011001000102000-

00100100000000000000001101001110-01000-000001113111100001010001---------
0100-0-1111

'Cyrtophora_sp'
1000000100000000010010000001001?10001---20---110211110000-0-00000000000-
00000----0001000000211000001010000010120000000001000100111100-
01201111000011010001011110--0011001000102000-
00100100000000000000001101001110-01000-000001113111100001010001---------
0100-0-1111

Neoscona
1000000100000000010010000001001?10001---20---110211110000-0-00000000000-
00000----0001000000211000001010000010120000000001000100111100-
01201111000011010001011110--0011001000102000-
00100100000000000000001101001110-01000-000001113111100001010001---------
0100-0-1111

'Gasteracantha_hasselti'
1000000100000000010010000001001?10001---20---110211110000-0-00000000000-
00000----0001000000211000001010000010120000000001000100111100-
01201111000011010001011110--0011001000102000-
00100100000000000000001101001110-01000-000001113111100001010001---------
0100-0-1111

'Macracantha_arcuata'
1000000100000000010010000001001?10001---20---110211110000-0-00000000000-
00000----0001000000211000001010000010120000000001000100111100-
01201111000011010001011110--0011001000102000-
00100100000000000000001101001110-01000-000001113111100001010001---------
0100-0-1111

'Menneus_sp'
1000000100000000010010000001001?10001---20---110211110000-0-00000000000-
00000----0001000000211000001010000010120000000001000100111100-
01201111000011010001011110--0011001000102000-
00100100000000000000001101001110-01000-000001113111100001010001---------
0100-0-1111

'Deinopis_clean'
1000000100000000010010000001001?10001---20---110211110000-0-00000000000-
00000----0001000000211000001010000010120000000001000100111100-
01201111000011010001011110--0011001000102000-
00100100000000000000001101001110-01000-000001113111100001010001---------
0100-0-1111

'Deinopis_longipes'
1000000100000000010010000001001?10001---20---110211110000-0-00000000000-
00000----0001000000211000001010000010120000000001000100111100-
01201111000011010001011110--0011001000102000-
00100100000000000000001101001110-01000-000001113111100001010001---------
0100-0-1111

'Tamopsis_sp'
1000000100000000001001000001001?10001---20---110211110000-0-00000000000-
00000----0001000000211000001010000101200000000100010011100-
0120111100001101000101110--0011001000102000-
0010010000000000000000011010011 10-01000-000001113111100001010001---------
0100-0-1111
'Oecobius_navus'
1000000100000000001001000001001?10001---20---110211110000-0-00000000000-
00000----0001000000211000001010000101200000000100010011100-
0120111100001101000101110--0011001000102000-
001001000000000000000001101001110-01000-000001113111100001010001---------
0100-0-1111
'Oecobius_sp'
1000000100000000001001000001001?10001---20---110211110000-0-00000000000-
00000----0001000000211000001010000101200000000100010011100-
0120111100001101000101110--0011001000102000-
001001000000000000000001101001110-01000-000001113111100001010001---------
0100-0-1111
'Zosis_sp' 1000000100000000001001000001001?10001-
--20---110211100000-0-00000000000-00000----
0001000000211000001010000101200000000100010011100-
0120111100001101000101110--0011001000102000-
001001000000000000000001101001110-01000-000001113111100001010001---------
0100-0-1111
'Philoponella_herediae'
1000000100000000001001000001001?10001---20---110211100000-0-00000000000-
00000----0001000000211000001010000101200000000100010011100-
0120111100001101000101110--0011001000102000-
001001000000000000000001101001110-01000-000001113111100001010001---------
0100-0-1111
'Uloborus_glomosus'
1000000100000000001001000001001?10001---20---110211100000-0-00000000000-
00000----0001000000211000001010000101200000000100010011100-
0120111100001101000101110--0011001000102000-
001001000000000000000001101001110-01000-000001113111100001010001---------
0100-0-1111
'Cybaeodamus_taim'
1000000100000000001001000001001?10001---20---110211110000-0-00000000000-
00000----0001000000211000001010000101200000000100010011100-
0120111100001101000101110--0011001000102000-
001001000000000000000001101001110-01000-000001113111100001010001---------
0100-0-1111
'Forsterella_sp'
1000000100000000001001000001001?10001---20---110211110000-0-00000000000-

00000----000100000021100000101000010120000000010001001111100-
0120111100001101000101111 0--0011001000102000-
001001000000000000000001101001110-01000-00000111311110000101000 1---------
0100-0-1111
'Caayguara_clean'
1000000100000000001001000001001?10001---20---110211110000-0-00000000000-
00000----000100000021100000101000010120000000010001001111100-
01201111000011010001011110--0011001000102000-
001001000000000000000001101001110-01000-000001113111100001010001---------
0100-0-1111
Amaurobius
1000000100000000001001000001001?10001---20---110211110000-0-00000000000-
00000----000100000021100000101000010120000000010001001111100-
01201111000011010001011110--0011001000102000-
001001000000000000000001101001110-01000-000001113111100001010001---------
0100-0-1111
'Callobius_sp'
1000000100000000001001000001001?10001---20---110211110000-0-00000000000-
00000----000100000021100000101000010120000000010001001111100-
01201111000011010001011110--0011001000102000-
001001000000000000000001101001110-01000-000001113111100001010001---------
0100-0-1111
'Neoramia_sp'
1000000100000000001001000001001?10001---20---110211110000-0-00000000000-
00000----000100000021100000101000010120000000010001001111100-
01201111000011010001011110--0011001000102000-
001001000000000000000001101001110-01000-000001113111100001010001---------
0100-0-1111
Amphinecta
1000000100000000001001000001001?10001---20---110211110000-0-00000000000-
00000----000100000021100000101000010120000000010001001111100-
01201111000011010001011110--0011001000102000-
001001000000000000000001101001110-01000-000001113111100001010001---------
0100-0-1111
'Metaltella_simoni'
1000000100000000001001000001001?10001---20---110211110000-0-00000000000-
00000----000100000021100000101000010120000000010001001111100-
01201111000011010001011110--0011001000102000-
001001000000000000000001101001110-01000-000001113111100001010001---------
0100-0-1111
Cambridgea
1000000100000000001001000001001?10001---20---110211110000-0-00000000000-
00000----000100000021100000101000010120000000010001001111100-
01201111000011010001011110--0011001000102000-

00100100000000000000001101001110-01000-000001113111100001010001---------
0100-0-1111
'Paramatachia_sp'
1000000100000000001001000001001?10001---20---110211110000-0-00000000000-
00000----00010000002110000010100001012000000001000100111100-
012011110000110100010111110--0011001000102000-
00100100000000000000001101001110-01000-000001113111100001010001---------
0100-0-1111
'Agelenopsis_emertoni_cdhitest.fasta.transdecoder'1000000100000000001001000001001?10001---20---110211110000-0-00000000000-00000----
00010000002110000010100001012000000001000100111100-
012011110000110100010111110--0011001000102000-
00100100000000000000001101001110-01000-000001113111100001010001---------
0100-0-1111
'Agelenopsis_pennsylvanica'
1000000100000000001001000001001?10001---20---110211110000-0-00000000000-
00000----00010000002110000010100001012000000001000100111100-
012011110000110100010111110--0011001000102000-
00100100000000000000001101001110-01000-000001113111100001010001---------
0100-0-1111
'Calymmaria_clean'
1000000100000000001001000001001?10001---20---110211110000-0-00000000000-
00000----00010000002110000010100001012000000001000100111100-
012011110000110100010111110--0011001000102000-
00100100000000000000001101001110-01000-000001113111100001010001---------
0100-0-1111
'Calymmaria_persica'
1000000100000000001001000001001?10001---20---110211110000-0-00000000000-
00000----00010000002110000010100001012000000001000100111100-
012011110000110100010111110--0011001000102000-
00100100000000000000001101001110-01000-000001113111100001010001---------
0100-0-1111
'Cicurina_travisae'
1000000100000000001001000001001?10001---20---110211110000-0-00000000000-
00000----00010000002110000010100001012000000001000100111100-
012011110000110100010111110--0011001000102000-
00100100000000000000001101001110-01000-000001113111100001010001---------
0100-0-1111
'Cicurina_vobora'
1000000100000000001001000001001?10001---20---110211110000-0-00000000000-
00000----00010000002110000010100001012000000001000100111100-
012011110000110100010111110--0011001000102000-
00100100000000000000001101001110-01000-000001113111100001010001---------
0100-0-1111

'Homalonychus_theologus'
1000000100000000001001000001001?10001---20---110211110000-0-00000000000-
00000----00010000002110000010100001012000000001000100111100-
012011110000110100010111110--0011001000102000-
0010010000000000000000001101001110-01000-000001113111100001010001---------
0100-0-1111
'Tengella_sp'
1000000100000000001001000001001?10001---20---110211110000-0-00000000000-
00000----00010000002110000010100001012000000001000100111100-
012011110000110100010111110--0011001000102000-
0010010000000000000000001101001110-01000-000001113111100001010001---------
0100-0-1111
'Uliodon_sp'
1000000100000000001001000001001?10001---20---110211110000-0-00000000000-
00000----00010000002110000010100001012000000001000100111100-
012011110000110100010111110--0011001000102000-
0010010000000000000000001101001110-01000-000001113111100001010001---------
0100-0-1111
'Misumenoides_formosipes'
1000000100000000001001000001001?10001---20---110211110000-0-00000000000-
00000----00010000002110000010100001012000000001000100111100-
012011110000110100010111110--0011001000102000-
0010010000000000000000001101001110-01000-000001113111100001010001---------
0100-0-1111
'Sidymella_sp'
1000000100000000001001000001001?10001---20---110211110000-0-00000000000-
00000----00010000002110000010100001012000000001000100111100-
012011110000110100010111110--0011001000102000-
0010010000000000000000001101001110-01000-000001113111100001010001---------
0100-0-1111
'Anahita_punctulata'
1000000100000000001001000001001?10001---20---110211110000-0-00000000000-
00000----00010000002110000010100001012000000001000100111100-
012011110000110100010111110--0011001000102000-
0010010000000000000000001101001110-01000-000001113111100001010001---------
0100-0-1111
'Peucetia_longipalpis'
1000000100000000001001000001001?10001---20---110211110000-0-00000000000-
00000----00010000002110000010100001012000000001000100111100-
012011110000110100010111110--0011001000102000-
0010010000000000000000001101001110-01000-000001113111100001010001---------
0100-0-1111
'Dolomedes_trinity'
1000000100000000001001000001001?10001---20---110211110000-0-00000000000-

00000----0001000000211000001010000101200000000010001001111100-
01201111000011010001011110--0011001000102000-
0010010000000000000000001101001110-01000-000001113111100001010001---------
0100-0-1111

'Pisaurina_mira'
1000000100000000001001000001001?10001---20---110211110000-0-00000000000-
00000----0001000000211000001010000101200000000010001001111100-
01201111000011010001011110--0011001000102000-
0010010000000000000000001101001110-01000-000001113111100001010001---------
0100-0-1111

'Cupiennius_sp'
1000000100000000001001000001001?10001---20---110211110000-0-00000000000-
00000----0001000000211000001010000101200000000010001001111100-
01201111000011010001011110--0011001000102000-
0010010000000000000000001101001110-01000-000001113111100001010001---------
0100-0-1111

Allocosa 1000000100000000001001000001001?10001-
--20---110211110000-0-00000000000-00000----
0001000000211000001010000101200000000010001001111100-
01201111000011010001011110--0011001000102000-
0010010000000000000000001101001110-01000-000001113111100001010001---------
0100-0-1111

'Schizocosa_rovneri'
1000000100000000001001000001001?10001---20---110211110000-0-00000000000-
00000----0001000000211000001010000101200000000010001001111100-
01201111000011010001011110--0011001000102000-
0010010000000000000000001101001110-01000-000001113111100001010001---------
0100-0-1111

'Habronattus_signatus'
1000000100000000001001000001001?10001---20---110211110000-0-00000000000-
00000----0001000000211000001010000101200000000010001001111100-
01201111000011010001011110--0011001000102000-
0010010000000000000000001101001110-01000-000001113111100001010001---------
0100-0-1111

'Habronattus_ustulatus'
1000000100000000001001000001001?10001---20---110211110000-0-00000000000-
00000----0001000000211000001010000101200000000010001001111100-
01201111000011010001011110--0011001000102000-
0010010000000000000000001101001110-01000-000001113111100001010001---------
0100-0-1111

'Teminius_sp'
1000000100000000001001000001001?10001---20---110211110000-0-00000000000-
00000----0001000000211000001010000101200000000010001001111100-
01201111000011010001011110--0011001000102000-

0010010000000000000000001101001110-01000-000001113111100001010001---------
0100-0-1111
'Karaops_raveni'
1000000100000000001001000001001?10001---20---110211110000-0-00000000000-
00000----00010000002110000010100001012000000000100010011 1100-
012011110000110100010111 10--0011001000102000-
0010010000000000000000001101001110-01000-000001113111100001010001---------
0100-0-1111
'Falconina_gracilis'
1000000100000000001001000001001?10001---20---110211110000-0-00000000000-
00000----000100000021100000101000010120000000001000100111100-
01201111000011010001011110--0011001000102000-
0010010000000000000000001101001110-01000-000001113111100001010001---------
0100-0-1111
'Nyssus_sp'
1000000100000000001001000001001?10001---20---110211110000-0-00000000000-
00000----000100000021100000101000010120000000001000100111100-
01201111000011010001011110--0011001000102000-
0010010000000000000000001101001110-01000-000001113111100001010001---------
0100-0-1111
'Clubiona_clean'
1000000100000000001001000001001?10001---20---110211110000-0-00000000000-
00000----000100000021100000101000010120000000001000100111100-
01201111000011010001011110--0011001000102000-
0010010000000000000000001101001110-01000-000001113111100001010001---------
0100-0-1111
'Rebilus_sp'
1000000100000000001001000001001?10001---20---110211110000-0-00000000000-
00000----000100000021100000101000010120000000001000100111100-
01201111000011010001011110--0011001000102000-
0010010000000000000000001101001110-01000-000001113111100001010001---------
0100-0-1111
'Liocranidae_GOOD'
1000000100000000001001000001001?10001---20---110211110000-0-00000000000-
00000----000100000021100000101000010120000000001000100111100-
01201111000011010001011110--0011001000102000-
0010010000000000000000001101001110-01000-000001113111100001010001---------
0100-0-1111
'Trachelas_tranquillus'
1000000100000000001001000001001?10001---20---110211110000-0-00000000000-
00000----000100000021100000101000010120000000001000100111100-
01201111000011010001011110--0011001000102000-
0010010000000000000000001101001110-01000-000001113111100001010001---------
0100-0-1111

'Sergiolus_capulatus'
1000000100000000001001000001001?10001---20---110211110000-0-00000000000-00000----00010000002110000010100001012000000001000100111100-012011110000110100010111110--0011001000102000-001001000000000000000001101001110-01000-000001113111100001010001---------0100-0-1111
'Molycria_sp'
1000000100000000001001000001001?10001---20---110211110000-0-00000000000-00000----00010000002110000010100001012000000001000100111100-012011110000110100010111110--0011001000102000-001001000000000000000001101001110-01000-000001113111100001010001---------0100-0-1111
Anzacia 1000000100000000001001000001001?10001---20---110211110000-0-00000000000-00000----00010000002110000010100001012000000001000100111100-012011110000110100010111110--0011001000102000-001001000000000000000001101001110-01000-000001113111100001010001---------0100-0-1111
'Lampona_sp'
1000000100000000001001000001001?10001---20---110211110000-0-00000000000-00000----00010000002110000010100001012000000001000100111100-012011110000110100010111110--0011001000102000-001001000000000000000001101001110-01000-000001113111100001010001---------0100-0-1111
Goniotarbus 1000?0011?000000???????????0???00-000--?????????????????-???00?0??????????????????010???002?1???00?0100???040010?010???0010-???0???113?1?????00?????021------????????????????????????????????????????????????????0000??????????????????????????---------0?---??????
Bornatarbus 1000?0011?000000???????????0???00-000--?????????????????-???00?0??????????????????010???002?1???00?0100???040010?000???0010-???0???113?1?????00?????021------????????????????????????????????????????????????????0000??????????????????????????---------0?---??????

;

END;
